# On-site ribosome remodeling by locally synthesized ribosomal proteins in axons

**DOI:** 10.1101/500033

**Authors:** Toshiaki Shigeoka, Max Koppers, Hovy Ho-Wai Wong, Julie Qiaojin Lin, Asha Dwivedy, Janaina de Freitas Nascimento, Roberta Cagnetta, Francesca van Tartwijk, Florian Ströhl, Jean-Michel Cioni, Mark Carrington, Clemens F. Kaminski, William A. Harris, Hosung Jung, Christine E. Holt

**Author notes:** Correspondence should be addressed to C.E.H. or T.S. H.H.-W.W, and J.Q.L contributed equally.

## Abstract

Ribosomes are known to be assembled in the nucleolus, yet recent studies have revealed robust enrichment and translation of mRNAs encoding ribosomal proteins (RPs) in axons, far away from neuronal cell bodies. Using subcellular proteomics and live-imaging, we show that locally synthesized RPs incorporate into axonal ribosomes in a nucleolus-independent fashion. We revealed that axonal RP translation is regulated through a novel sequence motif, CUIC, that forms a RNA-loop structure in the region immediately upstream of the initiation codon. Inhibition of axonal CUIC-regulated RP translation leads to defects in local translation activity and axon branching, demonstrating the physiological relevance of the axonal ribosome remodeling. These results indicate that axonal translation supplies cytoplasmic RPs to maintain/modify local ribosomal function far from the nucleolus.

## INTRODUCTION

RNA localization and local translation play key roles in the assembly and maintenance of neuronal connections (Campbell and Holt, 2001; Holt and Schuman, 2013; Wu et al., 2005). Recent genome-wide studies on the axonal transcriptome revealed that thousands of mRNAs are localized to the axon. A consistent but unexpected finding of these studies is the robust enrichment of mRNAs that encode ribosomal proteins (RPs), protein components of ribosomes. Axons are long neuronal processes that carry out many vital specific cellular functions far from their cell bodies, including translation, and must therefore maintain their protein synthetic machinery in good order. However, because most eukaryotic ribosome assembly is well known to occur in the nucleolus (Fromont-Racine et al., 2003; Lastick and McConkey, 1976), the physiological function of RP-coding mRNAs in a neuronal subcellular compartment far distant from the nucleus was enigmatic. RP-coding mRNAs have been abundantly detected in axons of a variety of neuron types, such as retinal ganglion cells (RGCs) (Zivraj et al., 2010), sympathetic neurons (Andreassi et al., 2010) and motor neurons (Saal et al., 2014) by using several different methods including microarray, SAGE and RNA-seq (Briese et al., 2016; Cajigas et al., 2012; Gioio et al., 2004; Gumy et al., 2011; Moccia et al., 2003; Taylor et al., 2009; Zivraj et al., 2010). The axonal localization of RP-coding mRNAs cannot simply be explained by Brownian diffusion of mRNAs from the soma since several studies showed that these transcripts are significantly enriched in the axon compared to the cell body (Andreassi et al., 2010; Saal et al., 2014), suggesting the presence of mechanisms that selectively target the RP-coding transcripts to the axon. Furthermore, recent studies provide evidence that RP-coding mRNAs are robustly translated in RGC axons both *in vivo* (Shigeoka et al., 2016) and *in vitro* (Cagnetta et al., 2018) raising the possibility that locally supplied RPs serve to support axonal function.

The eukaryotic ribosome is a macromolecular machine composed of 4 ribosomal RNA (rRNA) molecules and ~80 different RPs. The eukaryotic RPs are synthesized in the cytoplasm and commonly shipped into the nucleus for assembly into ribosomal subunits in the nucleolus although a few ribosomal proteins are added to the ribosome in the cytoplasm, such as Rpl24, Rpl10 and Rplp0 (Panse and Johnson, 2010). In contrast to the classical view that the ribosome is an invariant, monolithic molecular complex, several lines of evidence implicate heterogeneity and tissue-specificity of the RP composition of ribosomes. In spite of the house-keeping role of the ribosome, some RPs show highly tissue-specific expression patterns in vertebrates (Bortoluzzi et al., 2001). Furthermore, mutations or knockdown of certain RPs results in tissue-specific defects, many of which occur in the central nervous system (CNS). In zebrafish, translational inhibition of 21 RP mRNAs in whole embryos produces tissue-specific defects with 7 cases showing specific brain phenotypes (Uechi et al., 2006). Mice which carry a mutation in *rpl24* have severe axon degeneration in the optic nerve but few other CNS defects (Rice et al., 1997), and mice harboring a mutation in *rpl38* (Tail-short (Ts) mutants) display hearing loss, skeletal malformations and impaired fetal erythropoiesis (Morgan, 1950; Noben-Trauth and Latoche, 2011). Although the tissue specific phenotypes may partly be due to extra-ribosomal functions of RPs (Warner and McIntosh, 2009) and/or the difference in transcriptomes among tissues, technological advances in quantitative mass spectrometry (MS) have provided direct evidence for heterogeneity in ribosomal RP composition (Shi et al., 2017; Slavov et al., 2015). These studies uncovered that the stoichiometry among core RPs in the ribosome is variable and depends on the translational status of the ribosome. Furthermore, it has also been shown that the presence of certain RPs in the ribosome affects the target specificity of translation, suggesting a role of RPs in functional specialization of ribosomes (Kondrashov et al., 2011; Shi et al., 2017). These results strongly suggest the presence of ribosomes with variable core RP(s) in eukaryotic cells, but the mechanism that establishes the heterogeneity in RP composition remains unknown.

A number of previous studies call into question the widely held view of the ribosome as a stable molecular machine whose components remain unchangeable during its lifetime. For example, several RPs in the ribosome have higher turnover rates than other RP components, suggesting the possibility that individual RPs in the ribosome are replaced by free cytoplasmic (extra-ribosomal) RPs (Lastick and McConkey, 1976; Mathis et al., 2017). Furthermore, a potential role for free RPs in the maintenance of ribosomes in axons has been implicated in a mouse model of spinal muscular atrophy (SMA) (Bernabo et al., 2017). The depletion of the survival motor neuron (SMN) protein, an RNA-binding protein that associates with RP-coding mRNAs (Rage et al., 2013), caused a significant decrease in translation levels of RP-coding mRNAs in mouse motor neuron axons. Although the causal relationship remains uncertain, SMN depletion also leads to a 27% reduction in the number of ribosomes in the axons (Bernabo et al., 2017). These studies prompted us to ask whether axonal ribosomes incorporate locally synthesized RPs to support and/or modify the ribosome function far from the cell body although the limited amount of axonal material make this a technically challenging question to address.

In this study, we explore intra-ribosomal roles of axonally synthesized RPs using a range of technical approaches including live imaging, *in vivo* gene knockdown, bioinformatics, nascent protein labeling and mass spectrometry-based proteomics. We found that axonal translation of RPs coordinately peaks at the axon branching stage in RGCs *in vivo,* and their translation is regulated by a branch-promoting factor, Netrin-1, through a novel loop structure-forming sequence motif, CUIC, that is shared by ~70% of RP-coding mRNAs. Using nascent protein labeling and proteomic mass spectrometry analysis on ribosomes isolated from pure axons, together with live-imaging approaches, we show that axonally synthesized RPs are physically incorporated into axonal ribosomes in a nucleolus-independent fashion. Furthermore, we demonstrate the physiological importance of the axonal ribosome remodeling by showing that inhibition of axonal RP translation leads to a significant decrease in the level of axonal mRNA translation and severe axon branching defects *in vivo.* Our results provide direct evidence that cue-induced axonal translation supplies cytoplasmic RPs to support axonal ribosome function and axon arborization, and suggest a novel view that the ribosome can be dynamically remodeled ‘on-site’ through the cytoplasmic incorporation of locally-synthesized RPs.

## RESULTS

### Axonal synthesis of RPs is coordinately regulated *in vivo* and *in vitro*

First, we performed RNA-seq to analyze the transcriptome of RGC axons, which were isolated using Boyden chambers (Zheng et al., 2001) with a 1 um pore filter. Consistent with previous studies, ribosome-related gene ontology (GO) terms are robustly enriched in the axonal transcriptome (Figure 1A). We confirmed that mRNAs for >80 % of RPs were detected in RGC axons. Next, we investigated the developmental profile of axonal RP synthesis to estimate their physiological roles from their expression pattern. Analysis of our genome-wide translatome data of mouse RGC axons *in vivo* (Shigeoka et al., 2016) uncovered that the axonal translation of most RP-coding mRNAs peaks during the branching/synaptogenesis stage (P0.5) and declines thereafter in a synchronous manner (Figure 1B). We detected the axonal translation of 33 RP-coding mRNAs at P0.5 but only 5 RP mRNAs at later stages at a significant level. The pattern of translational changes of RP-coding mRNAs during the postnatal period (P0.5-P7.5) is significantly different from the other translated mRNAs (Figure 1B, lower panel), suggesting the presence of a mechanism that co-regulates the synthesis of many different RPs in axons. Since axon branching is regulated by extrinsic cues such as Netrin-1, Sema3A (Semaphorin 3A) and BDNF (brain derived nerve growth factor) (Alsina et al., 2001; Cioni et al., 2013; Dent et al., 2004; Lebrand et al., 2004; Manitt et al., 2009; Tang and Kalil, 2005), we hypothesized that axonal RP synthesis may be coordinately regulated by such cues. To test this, we analyzed a proteomics data set of the cue-induced nascent (newly synthesized) proteome in cultured *Xenopus laevis* RGC axons (Cagnetta et al., 2018). In this study, RGC axons were grown in physical separation from their cell bodies by culturing whole embryonic eyes in a Boyden chamber (Figure S1A). The intact nature of the eyes permits only the RGC axons to grow out of the eye (through the optic nerve) onto the transfilter and to extend through the 1 μm pores onto the reverse side thus providing a high purity source of axonal material. We then analyzed the newly synthesized proteins with and without cue-stimulation using stable isotope labeling by amino acids in cell culture combined with single-pot solid-phase sample preparation (pSILAC-SP3) and mass spectrometry (Figure S1A)(Cagnetta et al., 2018). We performed a GO enrichment analysis for groups of proteins whose translation is significantly promoted or suppressed by one of three branch-regulating factors, Netrin-1, Sema3A and BDNF (Alsina et al., 2001; Dent et al., 2004; Lebrand et al., 2004). This analysis revealed that RPs are particularly enriched in the group of proteins whose translation is promoted by Netrin-1, even after just 5 min of stimulation (Figure 1C, Figure S1B and Supplementary Table). To validate this finding, axons were stimulated with Netrin-1 for 5 min with or without a protein synthesis inhibitor, cycloheximide, and immunostained for Rps4x and Rps14. Consistent with the pSILAC-SP3 results, quantification of immunofluorescence (QIF) indicated a protein synthesis-dependent increase of the RP protein levels in axonal growth cones in response to Netrin-1 (Figure 1D and 1E).

**Figure 1.**
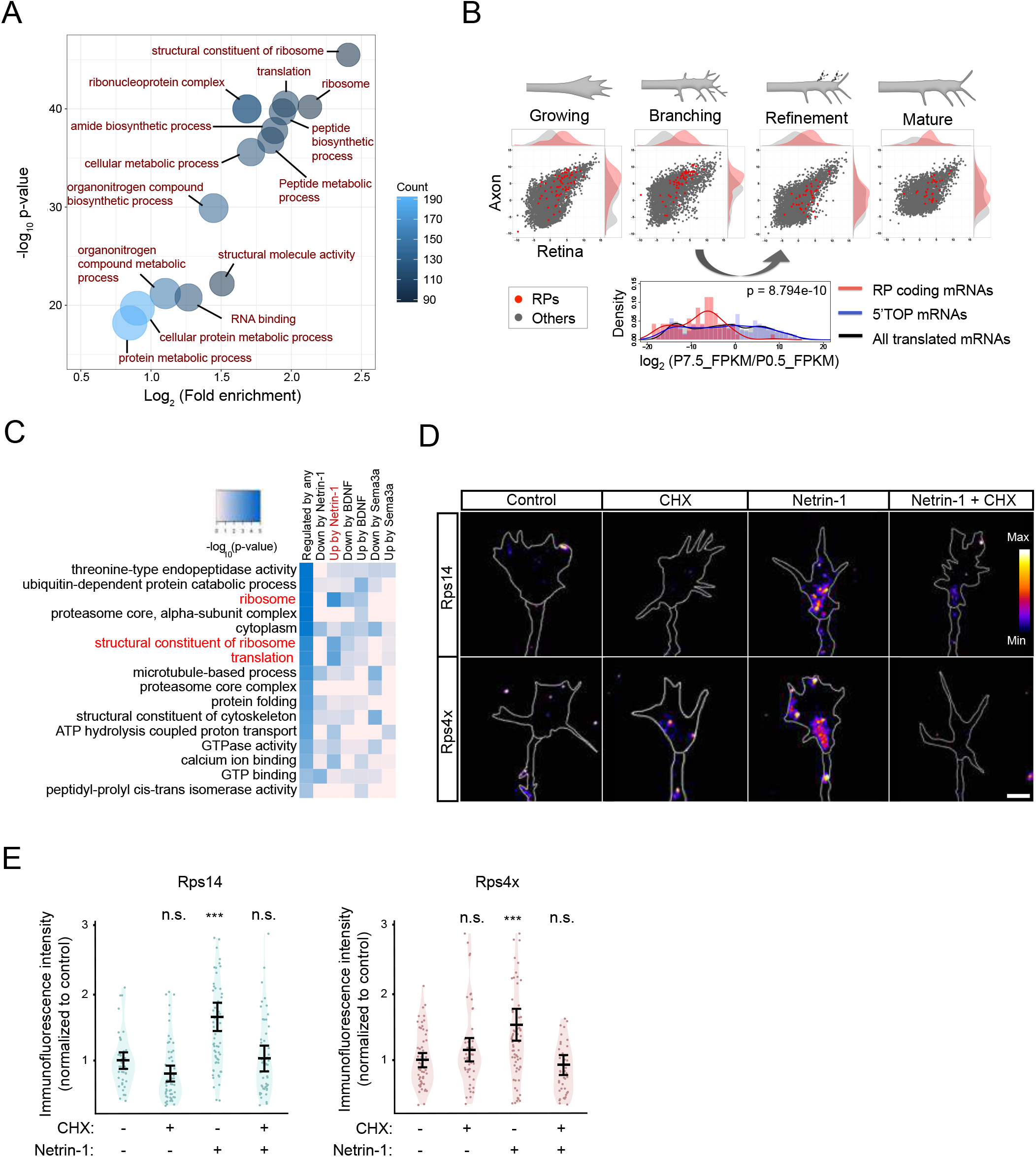
Axonally synthesis of RPs are coordinately regulated *in vivo* and *in vitro*. **(A)** Enrichment of GO terms in the frog RGC axon transcriptome, analyzed using DAVID. **(B)** Scatter plots with density plots (upper) showing relative abundance (FPKM) of translated mRNAs coding for RPs (red) and other proteins (grey) in the mouse RGC axon (y-axis) and retina (x-axis), which are obtained by Axon-TRAP system *in vivo.* The histogram (lower) shows the distribution of the ratio of abundance of two consecutive stages (Refinement/Branching). **(C)** Enrichment (Fisher’s p) of GO terms in gene groups whose translation is increased (“Up”) or decreased (“Down’) by cues. “Regulated by any” includes genes changed by any of three cues. Relative position of CUIC and RNA-secondary structure of 5’ UTRs of mouse RP coding mRNAs. In the heatmap, 5’ UTR sequences, in which each nucleotide is colored by the predicted secondary structure, are aligned by the position of CUIC. **(D-E)** Images **(D)** and plots **(E)** showing the result of quantitative immunofluorescence (qIF) for Rps4x (*n* = 59(Control), 50(CHX), 70(Netrin-1), 44(Netrin-1 + CHX)) and Rps14 (*n* = 42(Control), 59(CHX), 66(Netrin-1), 64(Netrin-1 +CHX)) in growth cones with or without Netrin-1 (5 min) /cycloheximide (CHX) treatment (bars = average with 95% CI, one-way ANOVA with Bonferroni’s Multiple Comparison test compared to the control, *** p≤0.001).

### Axonal translation of RP-coding mRNAs is induced by Netrin-1 through a loop structure-forming sequence motif upstream of the initiation codon

The coordinated regulation of axonal translation of RP-coding mRNAs led us to infer that these mRNAs have some common *cis* regulatory element(s). To explore this possibility, we performed a *de novo* motif discovery for mouse 5’ and 3’ UTR sequences of all RP-coding mRNAs (Heinz et al., 2010). Although we could not find any significantly enriched motif in 3’ UTRs, we identified a sequence motif of 8 nucleotides in the 5’ UTR (Figure S2A). We found that ~70% of RP-coding mRNAs share this novel 5’ UTR sequence motif, YYYYTTYC, which in most cases (>90%) is immediately (20-80nt) upstream of the initiation codon of RP-coding mRNAs (Figure 2A and S2B). The motif is highly conserved among animal species, especially at the TTYC “core” sequence (Figure S2C). Therefore, we named this YYYYTTYC motif as “Cis-element Upstream of the Initiation Codon (CUIC)”. Predicted RNA secondary structure of 5’ UTRs of RP-coding mRNAs shows that the nucleotide region around the CUIC motif tends to form a single-stranded loop, which may allow *trans* factor(s) to recognize this motif (Figure 2A and S2D and Supplementary Table). The single-stranded loop around the CUIC motif is also well conserved among animal species (Figure 2B). GO enrichment analysis revealed that not only ribosome related GO terms but also those linked to neuron morphogenesis, such as “neuron projection morphogenesis” and “neuron projection differentiation” are enriched in all CUIC-containing genes in the mouse genome (Figure 2C), suggesting a potential role of the CUIC motif in axon projection and branching. Consistent with this, CUIC-containing genes have significantly higher translation levels than average in the axonal translatome (Figure S2E) and they are significantly enriched in the axonally translated mRNAs identified by our Axon-TRAP analysis (Figure S2F). Pyrimidine-tract sequences in the 5’ UTR, such as the 5’ terminal oligopyrimidine (TOP)-like (Levy et al., 1991; Thoreen et al., 2012) and Pyrimidine-Rich Translational Element (PRTE) (Hsieh et al., 2012), are known to regulate translation. We therefore investigated the effect of the CUIC motif on axonal mRNA translation by performing fluorescence recovery after photobleaching (FRAP) experiments using a Venus fluorescent protein expressed from mRNAs with and without the CUIC motif in the UTR. We tested the UTR of Rps4x, the RP that showed the most significant increase of axonal translation after Netrin-1 stimulation (Figure 2D, Figure S1B). While no significant difference was observed in the FRAP signal in basal conditions, the addition of Netrin-1 caused a significantly higher recovery of fluorescence in the reporter containing the full length 5’ UTR, compared to the CUIC-deleted or 5’ UTR-deleted reporter constructs (Figure 2E and 2F), indicating a higher translation rate of the CUIC-containing mRNA. Consistent with these measurements, an independent single molecule translational imaging technique (Ifrim et al., 2015; Strohl et al., 2017; Tatavarty et al., 2012) revealed that a significantly higher number of translation events in axonal growth cones was observed from the CUIC-containing RP reporter when compared to the CUIC-deleted reporter construct in the Netrin-1 stimulated condition (Figure 2G). Together, these data provide evidence that the CUIC motif is, at least partially, responsible for the Netrin-1 induced axonal translation of RPs.

**Figure 2.**
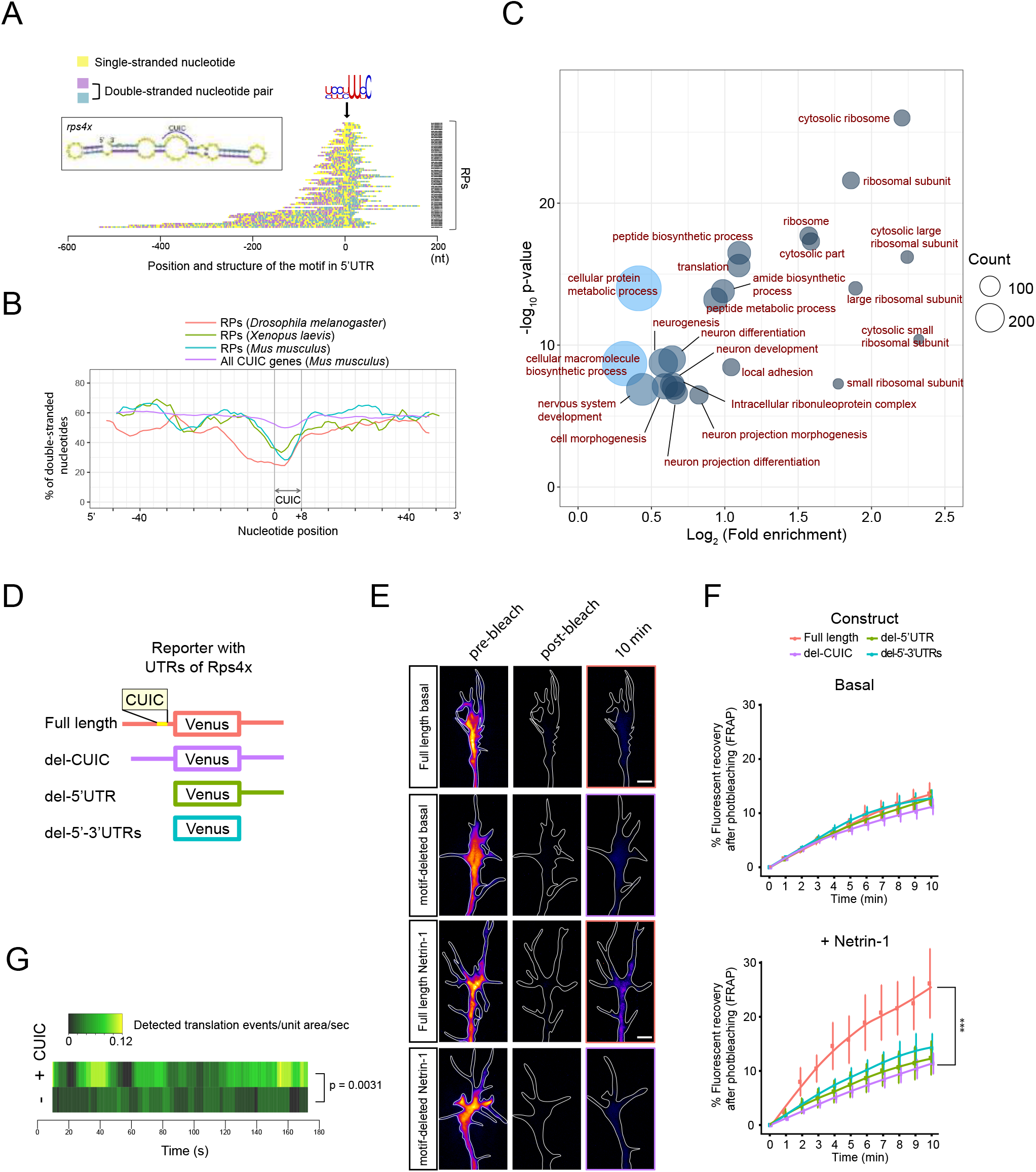
Axonal translation of RP-coding mRNAs is induced by Netrin-1 through a loop structure-forming sequence motif upstream of the initiation codon. **(A)** Relative position of CUIC and RNA-secondary structure of 5’ UTRs of mouse RP coding mRNAs. In the heatmap, 5’ UTR sequences, in which each nucleotide is colored by the predicted secondary structure, are aligned by the position of CUIC. **(B)** Average fraction of double-stranded nucleotides around CUIC in 5’ UTRs of all CUIC-containing mouse genes and RPs (moving average, 7 nucleotides window). **(C)** Enrichment of GO terms (GOTERM_BP_5 and GOTERM_CC_5) of all CUIC-containing mouse genes, analyzed using DAVID. **(D-F)** Venus reporter constructs **(D)** and images **(E)** and plots **(F)** of relative fluorescence recovery of Venus reporter constructs after photobleaching (error bar = SEM) in RGC growth cones. \*\*\**p* < 0.0001; two-way ANOVA comparing Full-UTRs (*n* = 8) with Del-motif (*n* = 12). (G) Heatmap showing moving average (20s window) of count of detected translation events per unit area per second with the Netrin-1 stimulation in a single molecule translation imaging. p-value: Mann–Whitney U test between Full-UTRs (*n* = 11) and Del-motif (*n* = 12) using total count in each growth cone.

### Axonal RP synthesis provides a cytoplasmic pool of free extra-ribosomal RPs

The coordinated regulation of axonally synthesized RPs suggests that they are likely to have a common ribosome-related function rather than a disparate variety of extra-ribosomal roles (Warner and McIntosh, 2009) in axons. However, since most ribosomal proteins are thought to be assembled into ribosomes exclusively in the nucleolus, it seemed puzzling that ribosomal proteins are synthesized in the distal axon, far away from the nucleolus (Fromont-Racine et al., 2003).One possibility is that the retrograde transport of axonally synthesized RPs allows the assembly of these into ribosomes in the nucleolus. Another possibility is that these locally synthesized RPs are incorporated into axonal ribosomes. To distinguish these possibilities, we first focused on the abundance of RPs outside of ribosomes (free cytoplasmic RPs) in the distal axon because robust axonal RP translation should result in the accumulation of free RPs in the axon unless these are retrogradely transported toward the soma, or quickly degraded. To detect free RPs in the axon, we performed sucrose density gradient fractionation of the axon lysate, generated using the Boyden chamber, and analyzed RP levels in ribosome-free fractions (Figure S3A). Mass-spectrometry analysis of fractionated whole-brain (control) samples showed that RPs are significantly enriched in the mono/polysome fraction but almost completely depleted in ribosome-free fractions (Figure 3A, 3B and S3B). In contrast, analysis of axon samples showed a robust accumulation of free RPs in ribosome-free fractions (Figure 3A and 3B). These results suggest that most axonally synthesized RPs remain in the axon.

**Figure 3.**
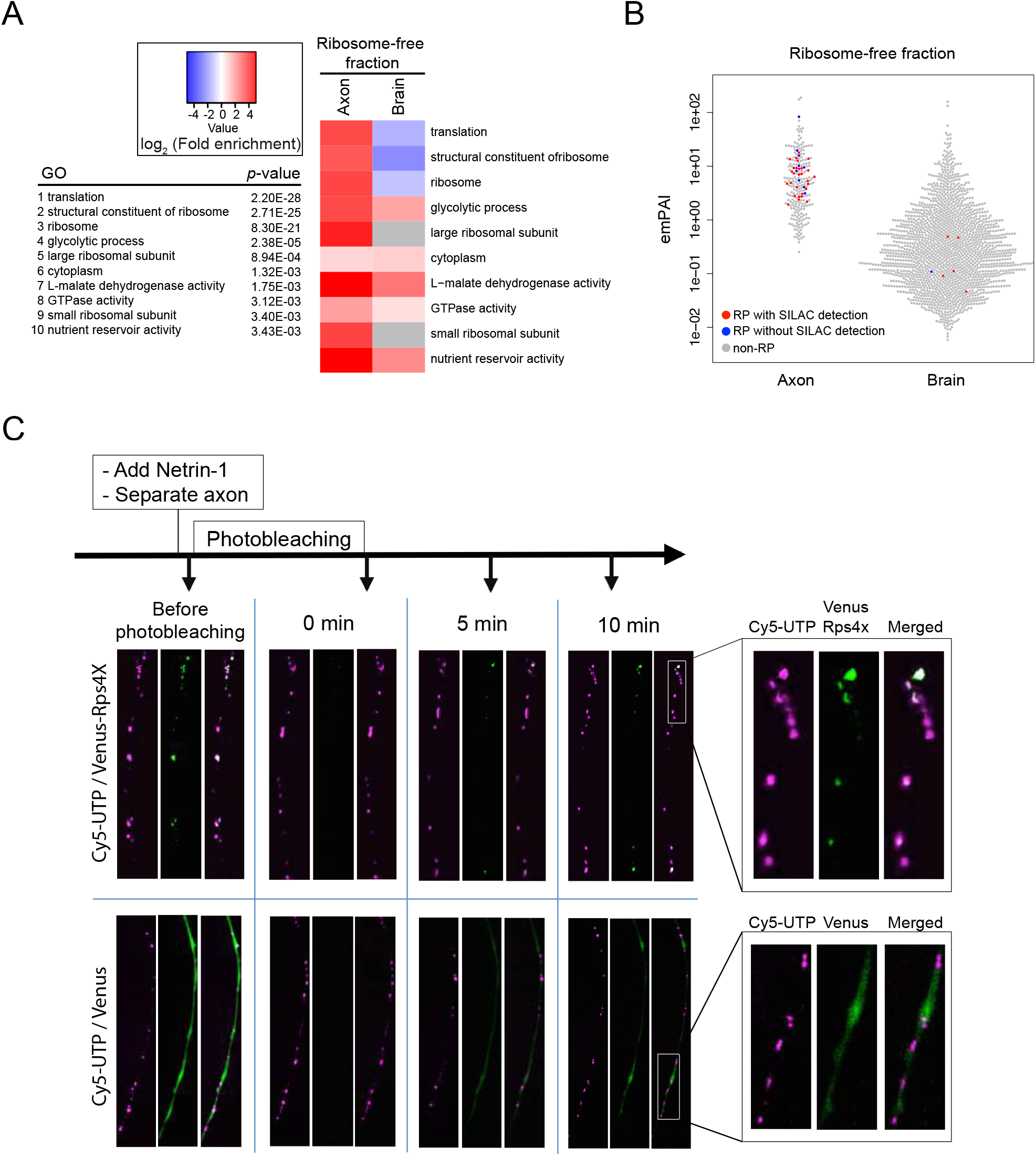
Axonally synthesized RPs remain in the axon and co-localize with ribosome-containing granules. **(A)** GO terms enriched in the ribosome-free fraction of the axon sample. Left: Ranking of p-value in the axon, Right: Fold enrichment values in the axon and the brain sample. **(B)** Relative abundance of proteins in the ribosome-free fraction (Red: RPs detected by SILAC, Blue: RPs not detected by SILAC, Grey: non-RP proteins). **(C)** Live imaging of Cy5-UTP (red) and UTR-Rps4x-Venus fusion or UTR-Venus (green) reporter constructs before and (0, 5 and 10 min) after photobleaching of green fluorescence.

### Axonally synthesized RPs co-localize with ribosome-containing granules

The fact that locally synthesized RPs remain in the axon favors the hypothesis that axonal RP synthesis provides a cytoplasmic pool of free RPs to support the ribosome function through axonal incorporation/exchange of ribosome components. Since the variable RP stoichiometry in the ribosomes suggested that a significant fraction of ribosomes is deficient in certain core RP component(s) in eukaryotic cells (Shi et al., 2017; Slavov et al., 2015), we reasoned that the interaction between these deficient ribosomes and cytoplasmic free RPs may serve to repair or modify the ribosome. Furthermore, our proteome analysis on isolated *Xenopus* RGC axons detected 133 nucleolus/ribosome assembly factors, more than 50% of whose mRNAs are detected in the axonal transcriptome (Supplementary table), suggesting the possibility that some of these auxiliary factors may support the local ribosomal incorporation of RPs. To investigate the axonal incorporation of locally synthesized RPs, we examined the co-localization of axonally synthesized RPs with ribosomes using a modified FRAP approach and dual-channel time-lapse imaging. In this experiment, we introduced a cDNA encoding Venus-Rps4x, a Venus-tagged RP fusion protein, and Cy5-UTP, which marks ribosome/RNA-rich granules (Wong et al., 2017), into RGC axons. Venus-Rps4x was expressed in RGC axons by eye electroporation in embryos injected with Cy5-UTP. The blastomere-injected Cy5-UTP broadly labels all RNA, including rRNA and mRNA, and is useful for distinguishing ribosome/RNA-rich granules in live axons (Wong et al., 2017). We then examined the co-localization between Cy5 fluorescence and the recovered Venus fluorescence after photobleaching the signal from pre-existing proteins to investigate the interaction of axonally synthesized Rps4x with a ribosome-containing granule. Before photobleaching, we observed that the fluorescence signals of Venus-Rps4x, but not the Venus-only control, were co-localized with Cy5 fluorescence, thus demonstrating that Cy5-UTP labeled ribosome-containing granules. Next, we cut the axons to exclude the possibility of newly synthesized Venus proteins being transported from the soma. Immediately after that, we added Netrin-1 to stimulate Rps4x synthesis, and then performed the photobleaching of the Venus fluorescence with 488-nm laser light. We found that the recovered Venus-Rps4x signal, but not control Venus signal, exhibits co-localization and co-movement with Cy5-labeled RNA granules 10 minutes after photobleaching in severed somaless RGC axons (Figure 3C), suggesting a rapid association of axonally synthesized Rps4x with ribosome-containing granules.

### Axonally synthesized RPs are incorporated into the ribosome in a nucleolus-independent manner

To obtain biochemical evidence for the axonal incorporation of newly synthesized RPs into the ribosome, we developed a novel method combining axon-pSILAC (Cagnetta et al., 2018) with axonal ribosome purification. In this method, we first labeled newly synthesized proteins with heavy amino acids (AAs) in somaless axons cultured in a Boyden chamber. This method of axon isolation was confirmed to contain pure axons and to exclude soma, nuclei and dendrites (Cagnetta et al., 2018). Approximately 2000 eyes were cultured for each experiment to obtain sufficient axonal material (~40 μg protein typical yield). Then, axonal ribosomes were purified using a sucrose cushion and ultracentrifugation, followed by identification of newly synthesized proteins with mass spectrometry (Figure 4A). We performed the analysis on axonal samples that were labeled with heavy amino acids for 3 hours after soma removal. We added Netrin-1 together with heavy amino acids to stimulate the axonal translation of RPs. We also tested this method on control eye samples labeled with heavy amino acids for 48 hours. Mass-spectrometry analysis revealed that ~93% of all detected proteins in the axonal ribosome samples are in a single gene-network cluster that includes RPs and proteins that interact with the ribosome, further validating the ribosome purification procedure (Figure 4B, Figure S4A). Strikingly, 21 RPs (26% of total number of RPs) labeled with heavy amino acids were detected in the axonal ribosome samples, suggesting that a considerable number of axonally synthesized RPs are incorporated into axonal ribosomes (Figure 4B). We also found that the ratios of labeled/unlabeled RP peptides in the axonal ribosome sample are highly variable among RPs (Figure 4C). Since the labeled/unlabeled ratio is expected to reflect the incorporation efficiency of newly synthesized RPs, this result suggests that the efficiency of axonal incorporation/replacement is significantly different among RP species. These results contrast with the relatively constant ratios found in whole cells (the eye sample) where most of the incorporation events are due to the *de novo* ribosome assembly in the nucleoli (Figure 4C). Remarkably, some RPs, such as Rps4x and Rps14, showed much higher labeled/unlabeled ratios in axons than those observed in whole cells (Figure 4C). This result excludes the possibility that the detection of labeled RPs in the axonal ribosome samples is caused by the contamination of our axon sample with whole cells because if labeled RPs in the axonal ribosome samples were sourced from RPs assembled into the ribosome in the nucleus of cells contaminating the axon culture, these cells, labeled for 3 hours, would give rise to lower labeled/unlabeled ratios of RP peptides than 48 hours labeled cells (the eye sample). We confirmed that neither puromycin nor RNase A/T1 treatment reduced this incorporation (Figure 4D and 4E), eliminating the possibility that the signal was due to the detection of nascent polypeptides, or due to the association of free extra-ribosomal RPs with mRNAs. This is also supported by the fact that the position of heavy peptides was not biased toward the N-terminus (Figure 4F). Finally, analysis of the location of labeled RPs in the eukaryotic ribosome structure (Klinge et al., 2012) revealed an enrichment of axonally synthesized RPs in the solvent exposed sides compared to the subunit interface, raising the possibility that axonal RP incorporation takes place mainly on the surface of mRNA-binding 80S ribosomes (Figure 4G and Figure S4B).

**Figure 4.**
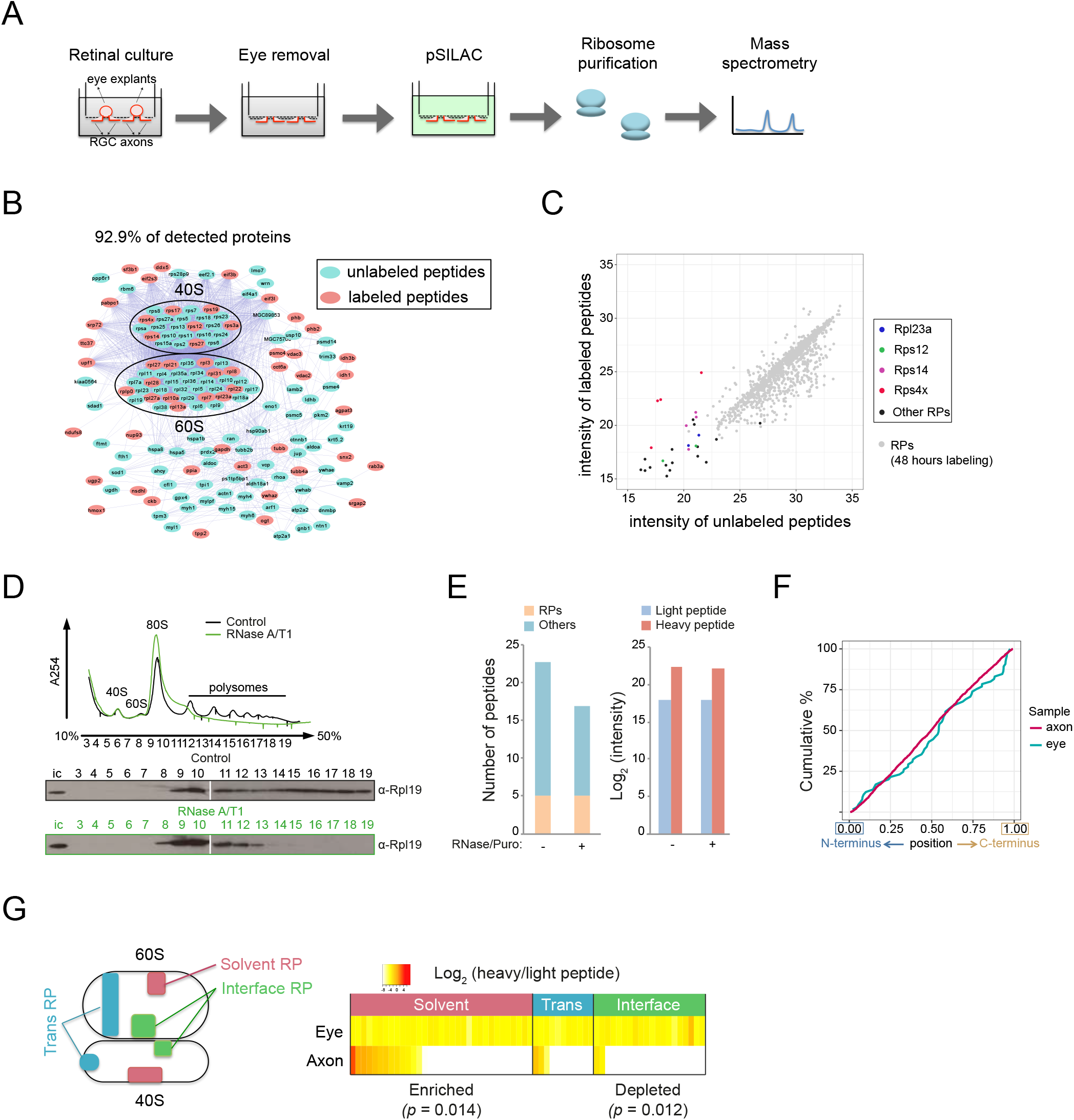
Axonally synthesized RPs are incorporated into the ribosome in a nucleolus-independent manner. **(A)** Diagram showing the strategy of SILAC and axonal ribosome purification. **(B)** Scatter plot showing abundance (log_2_ (intensity)) of labeled and unlabeled RP peptides detected in any of the biological replicates. **(C)** Gene network analysis of proteins detected in the axonal ribosome sample. Red nodes highlight proteins with SILAC-labeled peptides. **(D)** UV absorbance (A254) profile and immunoblot detection of Rpl19 in each fraction of samples with and without RNase A/T1 treatment. **(E)** Left panel shows the average number of peptides (RPs: pink, other proteins: green) detected in the puro/RNase-treated and untreated axonal ribosome sample. Right panel shows the comparison of intensity of a peptide from Rps4x. **(F)** Cumulative percentage of relative position of labeled peptides detected in ribosome samples. **(G)** Heatmap showing the number and ratio of intensity (heavy/light) of RPs detected in the eye and axonal ribosome samples. RPs were classified based on their position in the ribosomal subunits as depicted in the left diagram. One-tailed p-values were calculated by Fisher’s exact test using the number of detected RPs in each group.

Taken together, these findings suggest that locally synthesized RPs are incorporated into axonal ribosomes in a nucleolus-independent manner.

### Locally synthesized Rps4x is required to maintain ribosome function in axons

To ask whether axonally synthesized RPs incorporated into local ribosomes are important for their ability to synthesize proteins, we focused on Rps4x. Our imaging and proteomics analysis of the axonal ribosome revealed that Rps4x is robustly incorporated into ribosomes with higher efficiency than the other RPs (Figure 4C), consistent with the fact that Rps4x protein synthesis is induced most effectively by Netrin-1 stimulation of axons (Figure S1B). Furthermore, a recent study uncovered that the stoichiometry of Rps4x in the ribosome is significantly changed by the translational status in yeast and in mouse embryonic stem cells with the highest bias scores among all RPs, indicating that a considerable fraction of ribosomes in eukaryotic cells is deficient of Rps4x (Slavov et al., 2015). These results prompted us to hypothesize that the axonal translation and incorporation of Rps4x is particularly crucial for the ribosome function in the axon. To evaluate the impact of axonal Rps4x synthesis on ribosome function, we first performed axon-specific inhibition of Rps4x synthesis using a microfluidic chamber which isolates the axonal compartment from the cell body compartment while enabling visualization of the axons (Taylor et al., 2005). Taking advantage of the fluidic isolation of the solution in the chamber with a lower medium volume, we delivered FITC-conjugated *rps4x* morpholinos only into the axon chamber (Figure 5A).

**Figure 5.**
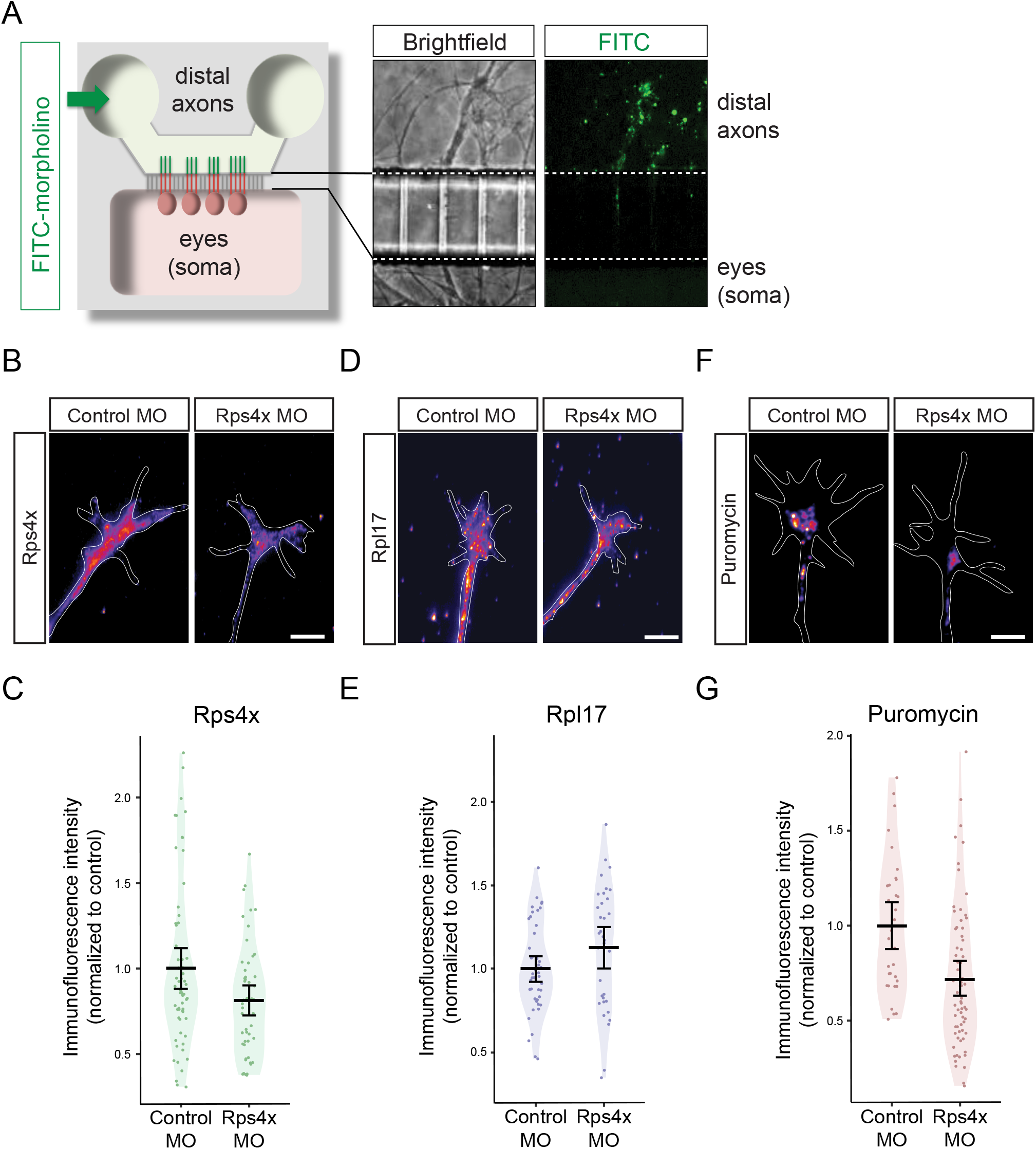
Locally synthesized Rps4x is required to maintain ribosome function in axons. **(A)** Diagram (left) and image of RGC axons with FITC-morpholino in a microfluidic chamber **(B-G)** Images **(B, D, F)** and plots **(C, F, G)** of average, 95% CI and distribution of normalized levels of Rps4x. *p* = 0.01454 (Welch t-test), n = 62 (Cont. MO), 52 (Rps4x MO) **(B-C)**, Rpl17. *p* = 0.09168 (Welch t-test), n = 46(Cont. MO), 34 (Rps4x MO) (d-E) and puromycin incorporation *p* = 0.0007783(Welch t-test), *p* = 0.0003976 (Mann-Whitney U test), *n* = 32 (Cont.), 66 (Rps4x) **(F-G)**.

We confirmed that Rps4x levels were decreased only in axons but not in the soma of the compartmentalized cultures (Figure 5B, 5C and S5). We also found that the level of another RP, Rpl17, is not changed by the *rps4x* morpholinos (Figure 5D and 5E), suggesting that the total ribosome level is not changed. To examine the effect of the inhibition of axonal Rps4x synthesis on ribosome function, we visualized and quantified newly synthesized proteins using pulse-labeling with a low concentration of puromycin, a structural analog of aminoacylated-tRNA and an anti-puromycin antibody (Starck et al., 2004; tom Dieck et al., 2015). Strikingly, we found that the *rps4x* morpholinos delivered to the axonal compartment, but not a control morpholino, significantly reduces the level of puromycin-labeled newly synthesized proteins in growth cones (Figure 5F and 5G). These data show that inhibition of axonal *rps4x* translation decreases the global translation level in the axon, suggesting a crucial role of axonally synthesized Rps4x in the axonal ribosome function.

### Axonal synthesis of Rps4x is crucial for axon branching *in vivo*

The last question we wanted to address in this study was whether newly synthesized RPs incorporated into axonal ribosomes are critical for axonal branching as our analysis showed that axonal synthesis of RPs is promoted by a branch-promoting factor, Netrin-1, and it peaks at the branching stage in mouse RGC axons *in vivo* (Figure 1). Moreover, a previous study demonstrated that exposed intact brains treated with the protein synthesis inhibitors cycloheximide and anisomycin show significantly (40%–70%) reduced arbor dynamics, suggesting that local translation plays a crucial role in proper arbor morphogenesis (Wong et al., 2017). We therefore asked whether the intra-ribosomal function of axonally synthesized Rps4x is critical for proper axon branching *in vivo.* Taking advantage of an *in vivo* system to visualize single *Xenopus* RGC axons (Wong and Holt, 2018; Wong et al., 2017), we tested the effect of inhibiting Rps4x synthesis on axon branching using morpholino-based knockdown (Figure 6A). *In vivo* electroporation of a morpholino that inhibits *Rps4x* translation into stage 28 eyes significantly reduced the number, length and complexity of RGC axon branches in the midbrain optic tectum at stage 45 (Figure 6B-F and Figure S6). To visualize the dynamic processes underlying these phenotypes, we performed *in vivo* live imaging of axons in the intact brain approximately 2.5 days following morpholino electroporation into the eye at stage 28. Filopodial and branching dynamics were both significantly reduced by inhibition of *Rps4x* translation at stage 41-43 (Figure 7A and Figure S7). Importantly, these branching phenotypes were rescued by a morpholino-resistant *Rps4x* cDNA with full length UTRs, but not by a cDNA lacking either the entire 5’ UTR or just the CUIC motif, suggesting that the CUIC-mediated regulation of Rps4x synthesis is crucial for axon branching (Figure 6B-F and Figure S6). To examine whether these phenotypes were caused by the inhibition of local axonal rather than somal translation of *Rps4x,* we delivered the morpholino directly into arborizing RGC axons in the tectum by electroporation at stage 41–43. Similar to the global inhibition of *Rps4x* translation in RGCs, local inhibition of axonal *Rps4x* translation abolished the net increases in both filopodia and branches (Figure 7A and Figure S7). To evaluate the extent of secondary effects due to morpholino-electroporated tectal (post-synaptic) cells, we delivered the morpholino into the tectum before the arrival of RGC axons (stages 35/36–37/38) and subsequently visualized the branching dynamics of axons after tectal entry at stage 41–43. No significant differences were observed in branching dynamics, indicating that inhibition of Rps4x synthesis in tectal cells is not responsible for the observed phenotypes (Figure 7A and Figure S7). Together, these results indicate that CUIC-mediated axonal translation of *Rps4x* is critical for proper arbor architecture of RGC axons *in vivo.*

**Figure 6.**
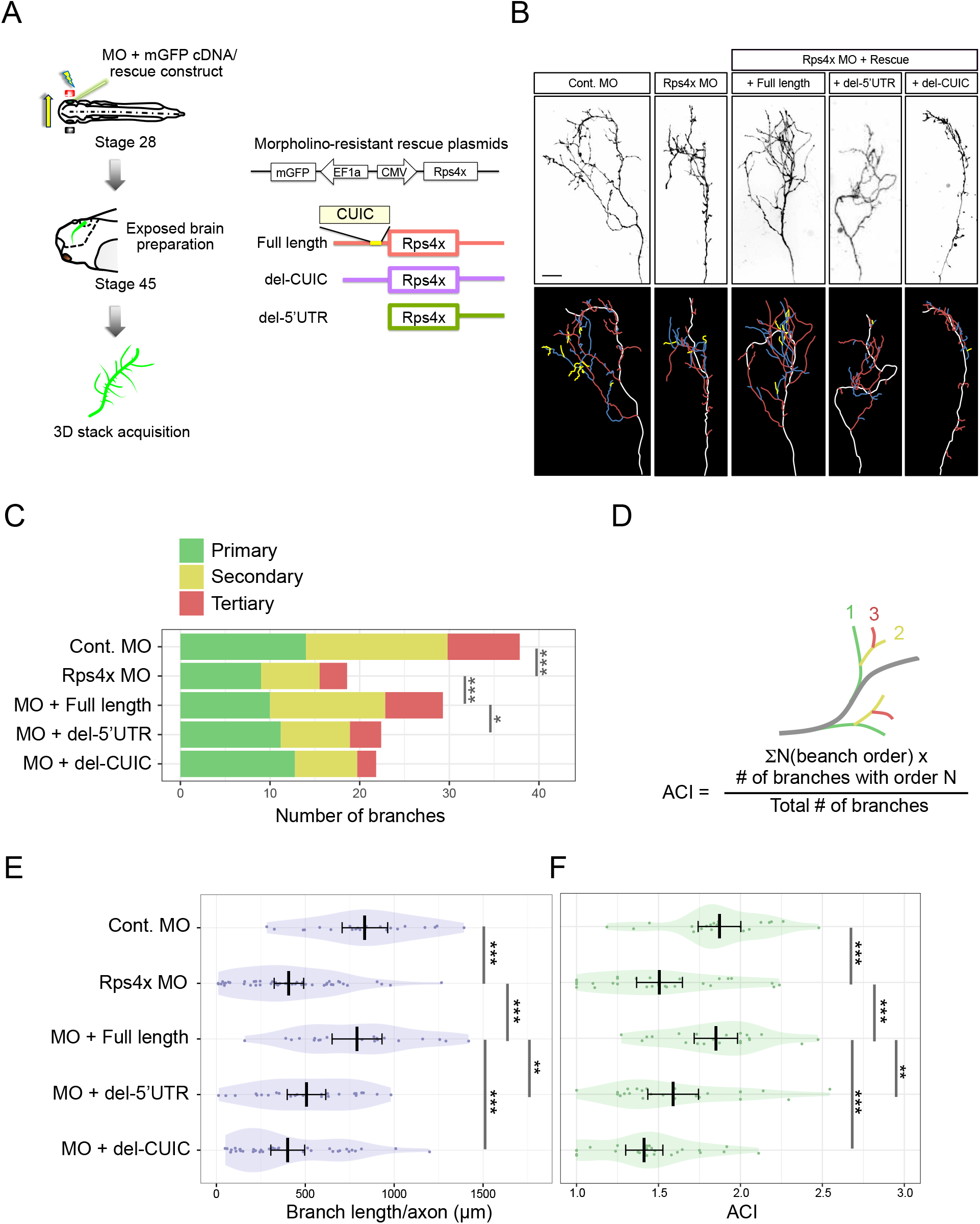
CUIC-mediated translation of Rps4x is crucial for axon branching *in vivo*. **(A)** Diagram for eye electroporation and live image acquisition of axon arbors. Dual promoter construct used for rescue condition. The rescue expression is driven by the CMV promoter and the GFP is driven by the EF1a promoter. **(B)** Lateral view of a single RGC axon in the tectum with color-coded images of axon shaft (white), primary (red), secondary (blue) and tertiary (yellow) branches. **(C)** A bar graph showing the number of branches in Control, Rps4x morphants and Rps4x morphants with different rescue constructs (primary: F4,146=5.3, p=0.0005; secondary: F4,146=12.8, p<0.0001; tertiary: F4,146=7.6, p<0.0001; total: F4,146=11.14,p<0.0001). (D) Formulation of axon complexity index (ACI). **(E-F)** Average, 95% CI and distribution of total branch length **(E)** and ACI **(F)** in the embryos electroporated with a morpholino/rescue construct (one-way ANOVA with Two-stage step-up method of Benjamini, Krieger and Yekutiei multiple comparisons test). *n* = 21 (Cont. MO), 47 (Cont. MO), 21 (MO+WT), 25 (MO+del-5’ UTR), 37 (MO+del-CUIC)

**Figure 7.**
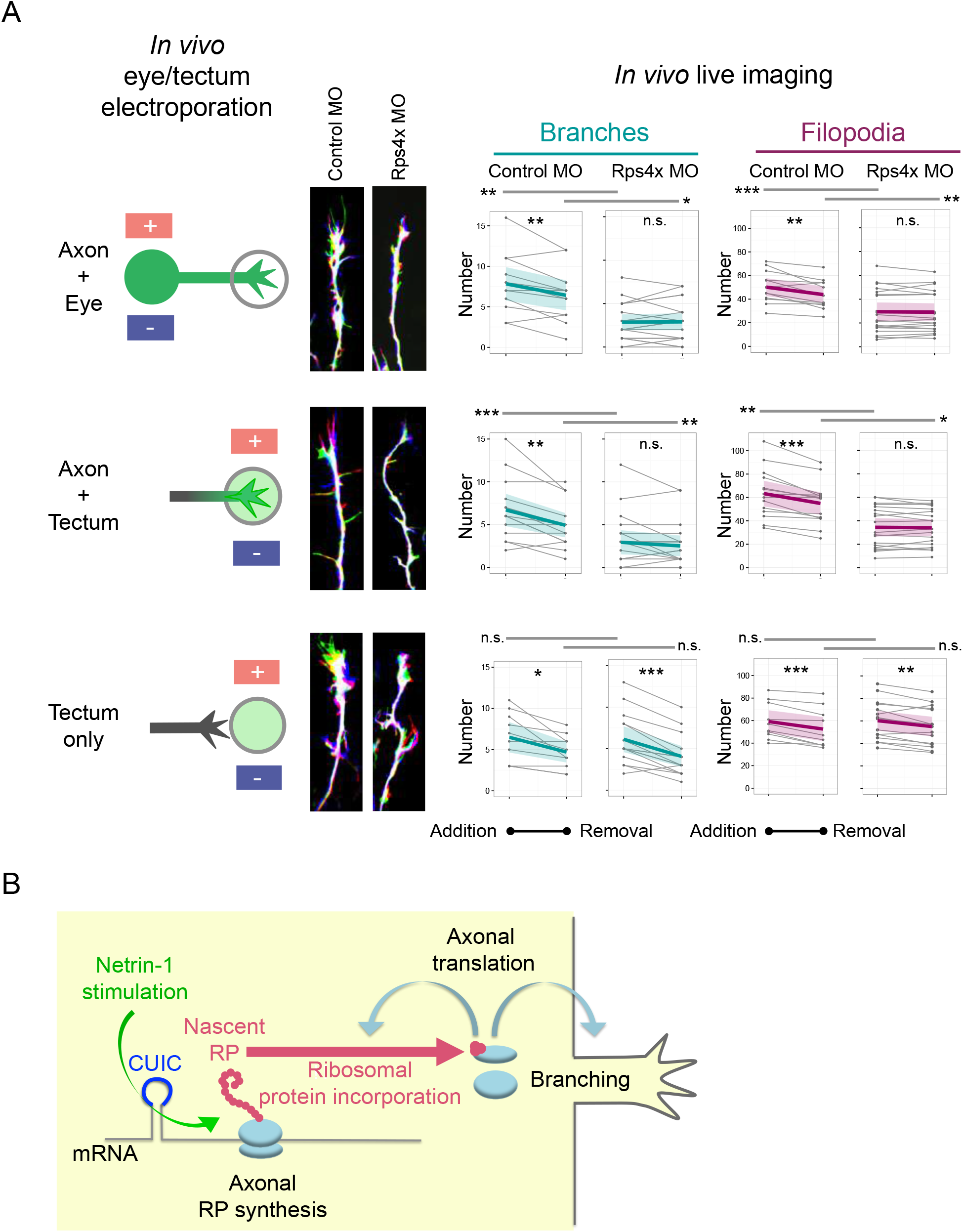
Axonal translation of Rps4x is crucial for axon branching *in vivo*. **(A)** Axon branching in Control MO- and rps4x MO-positive axons after eye electroporation (upper, n = 12 (Cont.), 20 (Rps4x)) and tectum electroporation at stages 41-43 (middle, n = 14 (Cont.), 19 (Rps4x)) and at stages stage 35-38 (lower, n = 10 (Cont.), 15 (Rps4x)). Images show merged overlay of 3 time-points (0’, 5’ and 10’ in blue, red and green respectively). Line graphs show number of added/removed branches/filopodia (paired and unpaired t test). **(B)** A model in which axonal RP synthesis supports the ribosome function and axonal branching.

## DISCUSSION

Our results provide evidence that locally synthesized RPs in RGC axons are incorporated into cytosolic ribosomes and are required for the axonal ribosome function. As axons must maintain cellular function at a long distance from their somas, our findings suggest that they have adopted a nucleolus-independent mechanism for the on-site repair and/or modification of their ribosomes through the local supply of newly synthesized RPs. Previous quantitative proteomics studies demonstrated that the stoichiometry of core components in ribosomes is highly variable, thereby indicating that a considerable fraction of ribosomes in eukaryotic cells does not contain the complete set of RPs (Shi et al., 2017; Slavov et al., 2015).

Although we cannot entirely rule out the possibility of *de novo* assembly of ribosomes in axons, a more plausible hypothesis is that the free, extra-ribosomal RPs in axons are incorporated into these specialized or defective ribosomes. It is thus possible that axonal RP synthesis plays an important role in ribosome maintenance, particularly for the ribosome components at the solvent-exposed sides that may be prone to damage (Figure 7B). It is also possible that the incorporation of axonal RPs modifies the ribosomal composition and structure to generate “specialized’’ ribosomes (Genuth and Barna, 2018; Kondrashov et al., 2011; Shi et al., 2017; Xue et al., 2015) which are tuned to preferentially translate specific mRNAs that regulate axon development and physiological function. Although the amount of protein obtained from axonal ribosomes in our system was not enough for the quantitative analysis of RP composition through proteomics or cryo-EM based methods, the comparison of the RP composition between axonal ribosomes and somal ribosomes will be an important objective for future research.

We uncovered that free extra-ribosomal RPs are highly enriched in axons compared to the whole cell (Figure 3A). This may partly explain the much higher frequencies of the ribosomal incorporation of free RPs in Netrin-1 stimulated axons (Figure 4) than those predicted from previous turnover studies performed on whole cells (Dice and Schimke, 1972; Lastick and McConkey, 1976). When considering the ribosomal incorporation of extra-ribosomal RPs as a form of protein-RNA interaction, the incorporation rate should depend on the concentration of free extra-ribosomal RPs. As Netrin-1 induced axonal RP translation may significantly increase the concentration of free RPs in axons, particularly in the case of Rps4x (Figure S1B), it may elicit an acute increase in ribosome remodeling in axons (Figure 4). An important question related to this is whether the ribosomal incorporation of RP in axons needs any catalytic activity of trans-acting auxiliary factors. In prokaryotes, a factor-free *in vitro* assembly of functional ribosomal subunits has been successfully demonstrated by bringing together purified rRNAs and RPs (Pulk et al., 2010; Sykes and Williamson, 2009), but this has not yet been reported in eukaryotes. Accumulating evidence suggests that a large number of non-ribosomal factors (>200) and small nucleolar RNAs are involved in the *de novo* ribosome assembly in eukaryotic cells, but most of these factors are related to rRNA processing or transport of RPs/pre-ribosomes, while only a few factors are known to support the incorporation of RPs (Kressler et al., 2010). Compared to the *de novo* ribosome biogenesis, which comprises the processing and folding of the pre-rRNA and its sequential assembly with ribosomal proteins, the cytoplasmic binding of a free RP to a deficient ribosome is a much simpler process. The axonal incorporation of most RPs can be explained simply by the affinity and interface stability between the rRNAs and RPs since most RPs have a small molecular size, making them easy-to-fold, and show affinity for rRNAs (Ulbrich and Wool, 1978). It is also possible that some ribosome assembly factors in axons play auxiliary roles in the ribosomal incorporation of some RPs, particular in case of large core RPs. Indeed, we identified a considerable number of ribosome assembly factors in the axonal proteome, transcriptome and/or translatome. For example, one of the axonally synthesized RPs detected in the axonal ribosomes is Rps3, a large and core component of the 40S subunit, and our transcriptome analysis revealed that RGC axons contain mRNA encoding Ltv1, a nucleolar protein whose function is required for Rps3 assembly onto ribosomes during *de novo* ribosome biogenesis (Loar et al., 2004; Seiser et al., 2006). This implicates the possibility that Ltv1 may play a supportive role in the axonal incorporation of Rps3. To fully understand the mechanisms underlying axonal RP incorporation, further work on the axonally-detected ribosome assembly factors is needed.

We identified a new motif, CUIC, which is a form of pyrimidine-rich element (PRE) with an 8 nucleotide core sequence, YYYYTTYC, in the region immediately upstream of the initiation codon of many axonally translated mRNAs coding for RPs. This motif in the RP-coding mRNAs is crucial not only for Netrin-1 induced axonal RP translation *in vitro* but also for axonal RP function in axon arborization *in vivo.* Furthermore, our analysis on RNA secondary structure revealed that this motif tends to form a single strand loop structure, suggesting that a single strand RNA binding protein (RBP) may be involved in the translational regulation of RPs. It has been known that the 5’ oligo-pyrimidine (TOP) sequence plays an important role in the translational control of mRNAs, including RP-coding mRNAs, downstream of the mTOR signaling pathway (Levy et al., 1991). Since the mouse expressed sequence tag (EST) showed that 5’ end of the CUIC-containing RP mRNAs is highly variable, there is a possibility that an alternative transcription start site (TSS) or an endonucleolytic cleavage site immediately upstream of CUIC generates the 5’ TOP-like mRNAs. Previous studies suggested that the RBP La-related protein 1 (LARP1), an RNA-binding protein (RBP), associates with 5’ TOP mRNAs to mediate the mTOR signal (Lahr et al., 2015), and another recent report uncovered that 5’ UTR LARP1 binding sites rarely overlapped with 5’ TOP sequences, but instead it binds predominantly at the 3’-most end of 5’ UTRs of 5’ TOP mRNAs (Hong et al., 2017). Together with our findings, these observations suggest an interesting possibility that the binding of RBP(s), such as LARP1, to the CUIC sequence controls the translation initiation at the initiation codon located immediately downstream of the CUIC sequence.

In this study, we uncovered crucial functions of locally synthesized Rps4x in axonal mRNA translation and in proper axon branching *in vivo.* A recent proteomic study showed that Rps4x is the most significantly sub-stoichiometric of all RPs in the polysomal ribosomes of eukaryotic cells (Slavov et al., 2015), suggesting the possibility that, compared to Rps4x-containing ribosomes, Rps4x-deficient ribosomes have less translation elongation activity, which may cause a polysomal traffic jam on the mRNA (Chu et al., 2014). Although we cannot completely exclude the possibility that some unknown extra-ribosomal function of Rps4x is responsible for the axon branching phenotype, we propose that the intra-ribosomal function of Rps4x is more likely to explain the phenotype observed after axonal inhibition of Rps4x synthesis since several lines of evidence suggest that the process of axon branching is highly dependent on local translation. Indeed, a previous study showed that RNA granules dock at sites of branch emergence and invade stabilized branches, and that acute inhibition of axonal translation by protein synthesis inhibitors causes a very similar axon branching defect as observed after Rps4x knockdown (Wong et al., 2017). Our previous Axon-TRAP study also suggested the importance of axonal translation in axon arborization (Shigeoka et al., 2016). We compared the axonally translated mRNAs at four different stages, E17.5, P0.5, P7.5 and adult, and we found that the number of axonally translated mRNAs is highest at the branching stage (P0.5), when approximately 70% of all axonally translated mRNAs were detected. These results suggest a critical role of axonal protein synthesis in determining the architecture of axon arbors and presynapses *in vivo,* and axonal RP translation induced by Netrin-1 and the CUIC motif may make axonal ribosomes competent for the extensive protein synthesis during axon arborization and synaptogenesis (Figure 7B). Further studies are needed to provide a fuller understanding of how RP mRNA translation in the pre-synaptic compartment contributes to these processes which are fundamental to the establishment of neural circuits.

## EXPERIMENTAL MODEL AND SUBJECT DETAILS

### *Xenopus laevis* Embryos

*Xenopus laevis* eggs were fertilized *in vitro* and embryos were raised in 0.1x Modified Barth’s Saline (MBS; 8.8mM NaCl, 0.1 mM KCl, 0.24mM NaHCO3, 0.1 mM HEPES, 82μM MgSO4, 33μM Ca(NO_3_)_2_, 41 μM CaCl_2_) at 14-20°C and staged according to the tables of Nieuwkoop and Faber (Nieuwkoop and Faber, 1994). All animal experiments were approved by the University of Cambridge Ethical Review Committee in compliance with the University of Cambridge Animal Welfare Policy. This research has been regulated under the Animals (Scientific Procedures) Act 1986 Amendment Regulations 2012 following ethical review by the University of Cambridge Animal Welfare and Ethical Review Body (AWERB).

### Primary *Xenopus* Retinal Cultures

Eye primordia were dissected from Tricaine Methanesulfonate (MS222) (Sigma-Aldrich) anesthetized embryos of either sex at stage 35/36 and cultured on 10μg/ml poly-L-lysine (Sigma-Aldrich)- and 10μg/ml laminin (Sigma-Aldrich)-coated glass bottom dishes (MatTek) in 60% L-15 medium (ThermoFisher), 1x Antibiotic-Antimycotic (ThermoFisher) at 20°C for 24-48 hours. 10-20 eye primordia (from 5-10 embryos) were cultured per dish and, typically, 2-3 dishes were used per experimental condition for each biological replicate.

## METHOD DETAILS

### Motif Analysis

All sequences of mouse cDNAs were retrieved from BioMart at Ensembl (GRCm38, ensemble Genes 91). *De novo* motif analysis was performed using HOMER version 3.0 with custom FASTA files containing all 5’ UTR sequences of mouse RP-coding cDNAs. For the analysis of CUIC-containing genes, we selected all genes whose mRNA contains the 8mer nucleotides (T/C)(T/C)(T/C)(T/C)TT(T/C)C located less than 100nt upstream of the initiation codon. The secondary structure analysis of UTRs was performed using performed using RNAfold in the ViennaRNA package version 2.4.3 with default settings. The conservation among species of all CUIC-containing RP mRNAs were calculated from phastCons60way.UCSC.mm10.

### Plasmid Construction

To construct Venus reporter plasmids used in FRAP and single molecule translation imaging, Venus cDNA and 5’ / 3’ UTR of mouse Rps4x (NM_009094) were integrated into BamHI-XbaI sites of pCS2+ (University of Michigan, Ann Arbor, Mich.). 8 nucleotides (CTCTTTCC) in the 5’ UTR were deleted in the “Del-motif” construct. To generate Rps4x-Venus fusion constructs, Venus and *X.laevis* Rps4x.S sequences were inserted into the BamHI-RcoRI sites of pCS2+. In these constructs, a CMV promoter drives the expression of 5’ UTR(Rps4x.S)-Venus-linker (Gly-Gly-Ser-Gly-Gly-Gly-Ser-Gly)-CDS (Rps4x.S, NM_001097003.1)-3’ UTR(Rps4x.S). Because frog 5’ UTR sequences of Rps4x.S in public databases could be truncated, we used a sequence of actually transcribed mRNA in frog embryos, which is obtained from a previously published RNA-seq data (GSM1948717) mapped to genome Xla.v91: 5’-cgcgctctcttcctgccagagttcagcgcgcactctttatcccggcgggaccggaaggaggaggtcttttcc-3’. To construct the plasmids for the rescue experiment of the morpholino phenotype, we replaced the β-actin cDNA of the morpholino insensitive β-actin/mGFP dual promoter construct(Wong et al., 2017) with 5’ UTR-CDS-3’ UTR of Rps4x.S in which silent mutations (ATGGCTCGCGGACCGAAGAAGC => ATGGCACGGGGCCCCAAAAAAC) were introduced to avoid morpholino binding. In the “del-CUIC” rescue construct, both of the two motifs (TCTCTTCC and TCTTTTCC) in the 5’ UTR were removed.

### Axon Culture and SILAC

Replicates in each experiment using *Xenopus laevis* in this study were obtained from different batches of embryos. We performed the axon cultures on a Boyden chamer device (1um pore, Falcon™ Cell Culture Inserts, 10289270/353102, Thermo fisher scientific) coated on both sides of the membrane with poly-L-lysine (10μg/ml) and only on the bottom side with laminin (10μg/ml). Eyes of stage 34-37 embryos of X. *laevis* were cultured on the filter of the chamber in 60% L15 medium containing penicillin streptomycin fungizone (Gibco) at room temperature for 48hrs. Then, we treated the eyes with lysine-, and arginine-free L15 (60%) medium for 1hr. After the removal of soma, the axons were cultured in L15 depletion medium containing “heavy” amino acids (84μg/ml [13C6,15N4] l-arginine, 146μg/ml [13C6,15N2] l-lysine (Silantes, Germany)) and Netrin-1 (600 ng/ml; R&D systems) for 3hrs. Soma removal was confirmed by DAPI staining. For the preparation of control eye samples, we cultured the dissected eyes in L15 depletion medium containing “heavy” amino acids at room temperature for 48hrs. Lysis of axons was performed using 500μl Lysis buffer (9mM Tris-HCl pH 7.4, 270mM KCl, 9 mM MgCl2, 1 % n-octylglycoside (Sigma-Aldrich),100μg/ml cycloheximide (Sigma-Aldrich), 0.5mM DTT, EDTA-free protease inhibitor cocktail (Roche) and SUPERase In RNase Inhibitor (Ambion)). Lysates were centrifuged at 16.000g at 4°C for 15 min and the supernatant was transferred to a cold 1.5ml tube. For the puromycin/RNase treated sample, axons were treated with 200μM puromycin for 15min before lysis and lysates were treated with 10μg/μl RNase A (Ambion) and 250U RNase T1 (Ambion) for 15 min.

### RNA-seq analysis of *Xenopus laevis* axons

To obtain the axonal translatome we performed axon cultures as described above. After 48 hours, we removed the soma and and lysed the axons in RLT buffer (Qiagen) containing β-mercaptoethanol. RNA was then extracted using the RNeasy Mini kit (Qiagen) followed by incolumn DNase I treatment to remove genomic DNA contamination. We then amplified cDNA using a method developed for single cell transcriptomics (Tang et al., 2009) with minor modifications (Shigeoka et al., 2016). The cDNA library preparation was performed using a KAPA Hyperprep kit (Roche). The cDNA libraries were subjected to a RNA-seq run on Next-seq 500 instrument (Illumina) using the 150 cycles high output kit (Illumina).

### Polysome Fractionation

For density gradient fractionation, lysate was layered on a sucrose gradient (10-50%) and ultracentrifugation was performed using a Beckman SW-40Ti rotor and Beckman Optima L-100 XP ultracentrifuge, with a speed of 35,000 rpm at 4°C for 160 min. Fractionations and UV absorbance profiling were carried out using Density Gradient Fractionation System (Teledyne ISCO). For sucrose cushioning, the lysate was layered on a 20% sucrose solution (20% sucrose, 10 mM Tris-HCl pH 7.4, 300 mM KCl,10 mM MgCl2), which contains a high concentration of KCl to avoid the binding of proteins to ribosomes. Then, ultracentrifugation was performed using a Beckman SW-55Ti rotor and Beckman Optima L-100 XP ultracentrifuge, with a speed of 41,000rpm at 4°C for 120 min.

### Quantitative PCR

RNA from fractionated samples was isolated using the RNeasy mini kit (Qiagen) and reverse transcribed into cDNA using random hexamers and the SuperScript III First-strand synthesis kit (Thermo Fisher Scientific). Triplicate reactions for qPCR were prepared using this cDNA and Quantitect SYBR Green PCR kit (Qiagen) according to manufacturer’s instructions. Plates were centrifuged shortly and run on a LightCycler 480 machine (Roche) using the following PCR conditions: denaturation step for 15 seconds at 94°C; annealing step for 30 seconds at 60°C; extension step for 30 seconds at 72°C. The following primers were used for qPCR: *18S rRNA* 5’-GTAACCCGCTGAACCCCGTT-3’ and 5’CCATCCAATCGGTAGTAGCG-3’; *28S rRNA* 5’-CTGTCAAACCGTAACGCAGG-3’ and 5’CTGACTTAGAGGCGTTCAGTCA-3’.

### Axonal Morpholino Introduction

Modified microfluidic chambers (Xona microfluidics, SOC150) were pre-coated with poly-L-lysine (10μg/ml) and with laminin (10μg/ml). Eyes dissected from the stage 30-33 X. *laevis* embryos were plated in the open chamber of SOC150. RGC axons were grown in 60% L15 medium containing μg/ul penicillin streptomycin fungizone at room temperature for 48 hrs. For the morpholino introduction, we prepared two solutions: diluted transfection reagent (2μl NeuroPORTER Transfection Reagent (Sigma-Aldrich) with 5.5μ L15 (60%)) and morpholino solution (2.5μl of 1mM morpholino oligonucleotide (mixture of 5’-CTTTTTCGGTCCACGAGCCATTTTC-3’ (against Rps4x.L) and 5’-TTCTTCGGTCCGCGAGCCATG-3’ (against Rps4x.S) with 5ul L15 (60%)). We mixed 7.5μl morpholino solution with 7.5μl diluted transfection reagent and incubated the mixture at room temperature for 5 min. We added the 15μl of mixture directly to 200μl of the growing medium in the axon chamber and incubate it at room temperature for 24 hours.

### Immunohistochemistry

After detaching microfluidic chambers from the glass bottom dishes, *Xenopus* retinal cultures were fixed in 2% formaldehyde/7,5% sucrose in PBS for 20 min at 20°C. The fixed cultures were steamed for 10min in sodium citrate buffer for antigen retrieval for ribosomal protein staining. Subsequently, they were permeabilized for 3-5 min in 0.1% Triton-X-100 in PBS and blocked with 5% heat-inactivated goat serum in PBS for 30 min at 20°C. Primary antibodies were incubated overnight at 4°C, followed by Alexa Fluor-conjugated secondary antibodies for 45 min at 20°C in dark. Cultures were mounted in FluorSave (Calbiochem). Antibodies were used at the following dilutions. Primary antibodies: rabbit anti-Rps4X (Proteintech, Cat#14799-1-AP; RRID: AB_2238567, 1:200), 1:200 rabbit anti-Rpl17 (Proteintech, Cat#14121-1-AP; RRID: AB_2253985, 1:200). Secondary antibodies: goat antirabbit Alexa Fluor 594 (Abcam, Cat#ab150080; RRID: AB_2650602, 1:1000).

For puromycin labelling, culture medium in the axonal compartment was replaced with 200μl of culture medium containing 600ng/ml Netrin-1 (Sigma) and 10μg/ml puromycin (Sigma). After 20min, the axonal compartment was washed once with fresh culture medium before detaching the microfluidic chamber from the dish. The retinal culture was immediately fixed in 2% formaldehyde/7,5% sucrose in PBS for 20 min at 20°C, permeabilized for 3-5 min in 0.1% Triton-X-100 in PBS, blocked with 5% heat-inactivated goat serum in PBS for 30 min at 20°C and then labelled with Alexa Fluor 647-conjugated mouse anti-puromycin antibody (Millipore, Cat#MABE343-AF647, 1:250) overnight at 4°C. Cultures were mounted in FluorSave (Calbiochem). Randomly selected non-collapsed growth cones were imaged at 60x on a Nikon Eclipse TE2000-U inverted microscope equipped with an EMCCD camera. Exposure time was kept constant and below gray-scale pixel saturation.

### Blastomere Microinjection

Embryos were microinjected with Cy5-UTP at 100μM in a total volume of 5nl (PerkinElmer) into both of the dorsal blastomeres at 4-or 8-cell stage. Embryos were first de-jellied in 2% Cysteine (Sigma-Aldrich) in 1x MBS (pH 8.0), washed 3 times in 0.1x MBS and aligned on a grid in 4% Ficoll (Sigma-Aldrich) in 0.1x MBS with 1X antibiotic-antimyotic (Thermo Fisher Scientific). Injections were performed using glass capillary needles (outer diameter: 1.0mm; inner diameter: 0.5mm; Harvard Apparatus) and a pressurized microinjector (Picospritzer, General Valve).

### Fluorescence Recovery After Photobleaching

Retinal cultures for FRAP assays were obtained from eyes of stage 34 embryos expressing one of the four Venus constructs indicated in Figure 2D or Venus-Rps4X and Venus-only for Cy5-UTP colocalization experiments, which were introduced by eye-targeted electroporation at stage 26. FRAP imaging was performed on an Olympus IX81 inverted microscope equipped with a PerkinElmer Spinning Disk UltraVIEW VoX and a 60x (NA, 1.30) Olympus silicone oil immersion objective. Images were acquired with an ORCA-Flash4.0 V2 CMOS camera (Hamamatsu) using Volocity software (PerkinElmer). Photobleaching was performed using an UltraVIEW PhotoKinesis device (PerkinElmer). The photobleached area was manually defined so that growth cones and >20μm of the axon shaft were bleached (thus reducing likelihood of fluorescence recovery resulting from Venus diffusion from unbleached areas of the axon shaft). Photobleaching was performed at 85-90% laser power (488 nm laser line) with 20–30 bleach cycles. Time-lapse images were captured at 1 min intervals using a 488 nm laser line at 20% laser power, together with phase contrast images for the corresponding time point. Exposure time was adjusted to avoid pixel saturation.

### Single Molecule Translation Imaging

Embryos at stage 26 were electroporated with plasmids and left in 0.1X MBS to continue to develop. Venus-expressing eyes from electroporated embryos at stage 34 were dissected and cultured. After 24 hours, a non-collapsed fluorescent growth cone was randomly selected and, prior to the bleaching step, imaged with low irradiance (<2W/cm^2^) in both fluorescence and bright field mode to generate an outline image. The growth cone was then photobleached for 10-30s with an irradiance of 1.5 kW/cm2 to eliminate the fluorescence. A reduced laser intensity of 0.3 kW/cm2 was used to ensure survival of the axons while simultaneously bleaching newly synthesized Venus proteins. The flash-like recovery of Venus fluorescence recorded with an exposure time of 200 milliseconds for 180 seconds. After that another bright field image was taken to check for survival. Retracted growth cones were excluded from analysis. In Netrin-1-stimulated conditions, 600ng/ml of Netrin-1 was bath applied immediately before the photobleaching step. All imaging steps were performed under epifluorescence illumination. An EM gain of 200 was used on the EMCCD camera to ensure single molecule sensitivity. The field of illumination was twice the size of the imaged field of view to bleach diffusing or transported fluorescent proteins from outside the growth cone before entering the field of view. Imaging was performed on a custom-made inverted single-molecule fluorescence microscope built around a commercial microscope frame (Olympus IX73). The illumination laser wavelength was at 488nm (Coherent Sapphire) for excitation of the YFP derivate Venus in combination with a 525/645 emission filter (Semrock) and a dichroic beam splitter (Chroma ZT405/488/561/640rpc). The laser beam was circularly polarized to excite fluorescent proteins homogeneously regardless of their orientation. The microscope was equipped with an EM-CCD camera (Andor iXon Ultra 897) with effective pixel size on the sample of 118 nm. A 100**x** NA=1.49 oil immersion TIRF objective (Olympus) was used.

### *In Vivo* Knockdown and Imaging

Targeted eye electroporation was performed as previously described(Falk et al., 2007; Wong et al., 2017). Stage 28 embryos were anesthetized in 0.4mg/ml MS222 in 1X MBS. The retinal primordium was injected with electroporation mixture, followed by electric pulses of 50ms duration at 1000ms intervals, delivered at 18V (please refer to the list below for the mixture and the number of electric pulses delivered for each experiment). The embryos were recovered and raised in 0.1X MBS until the desired embryonic stage for experiment.

1. Mature axon visualization (Figure 6B, 6E, 6F and Extended Figure 6): 1μg/μl of pCS2+mGFP (or 1μg/μl of pCS2+mGFP/MO resistant Rps4x rescue dual promoter construct cDNA for rescue experiments), 0.5mM control/Rps4x MO; 1 pulse.
2. Axon branching dynamics (Figure 7 and Extended Figure 7): 1μg/μl of pCS2+mGFP or 1μg/μl of pCS2+mRFP, 0.5mM control/Rps4x MO; 4 pulses

For tectum electroporation (Figure 7 and Extended Figure 7), the lateral surface of the hemisphere of the brain contralateral to the eye labeled with mRFP (electroporated at Stage 28 as described above) was exposed by careful removal of overlying eye and epidermis. 8X 18V electric pulses of 50ms duration at 1000ms intervals were delivered immediately after the 1mM control/Rps4x MO was locally ejected at the vicinity of the target area. The procedure was repeated once to ensure efficient delivery of the MO(Wong et al., 2017). Embryos were lightly anaesthetized with 0.4mg/ml MS222 in 1xMBS. The lateral surface of the brain contralateral to the electroporated eye was exposed by carefully removing the overlying epidermis and the contralateral eye. The electroporated eyes were also surgically removed to prevent somal contribution of proteins in Figure 2C and Extended Figure 5. Embryos were mounted in an oxygenated chamber created with Permanox slides (Sigma-Aldrich) and Gene Frame (ThermoFisher) and bathed in 1xMBS with 0.1mg/ml MS222, for visualization with fluorescence microscopy. Imaging was performed using 40X (NA 1.25) or 60X UPLSAPO objectives (NA 1.3) with a PerkinElmer Spinning Disk UltraVIEW ERS, Olympus IX81 inverted spinning disk confocal microscope. Z-stack intervals of 1-2μm were employed for acquiring images with Volocity (PerkinElmer).

Proteins were resolved by SDS-PAGE on NuPage 4-12% Bis-Tris gels (Invitrogen) and transferred to nitrocellulose membrane (Bio-Rad). The blots were blocked in milk and then incubated with primary antibodies in milk overnight at 4°C. After washing 3 times with TBS-T the blots were incubated with HRP-conjugated secondary antibodies for 1 hour at RT, washed again for 3 times in TBS-T, followed by ECL-based detection (Pierce ECL plus, Thermo Scientific). The following primary antibodies were used for Western blot analysis: mouse anti-Rpl19 (Abcam, Cat#ab58328; RRID: AB_945305, 1:1000), mouse anti-Rps23 (Abcam, Cat#ab57644; RRID: AB_945314, 1:1000), rabbit anti-Rps4X (Proteintech, Cat#14799-1-AP; RRID: AB_2238567, 1:1000), rabbit β-catenin (Sigma-Aldrich, Cat#C2206; RRID: AB_476831, 1:8000).

### Mass Spectrometry

1D gel bands were transferred into a 96-well PCR plate. The bands were cut into 1mm2 pieces, destained, reduced (DTT) and alkylated (iodoacetamide) and subjected to enzymatic digestion with chymotrypsin overnight at 37°C. After digestion, the supernatant was pipetted into a sample vial and loaded onto an autosampler for automated LC-MS/MS analysis. All LC-MS/MS experiments were performed using a Dionex Ultimate 3000 RSLC nanoUPLC (Thermo Fisher Scientific Inc, Waltham, MA, USA) system and a Q Exactive Orbitrap mass spectrometer (Thermo Fisher Scientific Inc, Waltham, MA, USA). Separation of peptides was performed by reverse-phase chromatography at a flow rate of 300nL/min and a Thermo Scientific reverse-phase nano Easy-spray column (Thermo Scientific PepMap C18, 2μm particle size, 100A pore size, 75μm i.d. × 50cm length). Peptides were loaded onto a precolumn (Thermo Scientific PepMap 100 C18, 5μm particle size, 100A pore size, 300μm i.d. × 5mm length) from the Ultimate 3000 autosampler with 0.1% formic acid for 3 minutes at a flow rate of 10μL/min. After this period, the column valve was switched to allow elution of peptides from the pre-column onto the analytical column. Solvent A was water + 0.1% formic acid and solvent B was 80% acetonitrile, 20% water + 0.1% formic acid. The linear gradient employed was 2-40% B in 30 minutes. The LC eluant was sprayed into the mass spectrometer by means of an Easy-Spray source (Thermo Fisher Scientific Inc.). All m/z values of eluting ions were measured in an Orbitrap mass analyzer, set at a resolution of 70000 and was scanned between m/z 380-1500. Data-dependent scans (Top 20) were employed to automatically isolate and generate fragment ions by higher energy collisional dissociation (HCD, NCE:25%) in the HCD collision cell and measurement of the resulting fragment ions was performed in the Orbitrap analyser, set at a resolution of 17500. Singly charged ions and ions with unassigned charge states were excluded from being selected for MS/MS and a dynamic exclusion window of 20 seconds was employed. Raw data were processed using Maxquant (version 1.6.1.0) (Cox and Mann, 2008) with default settings. MS/MS spectra were searched against the X. *laevis* protein sequences from Xenbase (xlaevisProtein.fasta). Enzyme specificity was set to trypsin/P, allowing a maximum of two missed cleavages. The minimal peptide length allowed was set to seven amino acids. Global false discovery rates for peptide and protein identification were set to 1%. The match-between runs and re-quantify options were enabled.

## QUANTIFICATION AND STATISTICAL ANALYSIS

### Statistics

The n number for each experiment, details of statistical analysis and software are described in the figure legends or main text. Statistical analyses used in this study include one-way ANOVA, Welch’s t-test, Mann-Whitney U test and Fisher’s exact Test. Statistical significance is defined as, n.s., not significant, *P<0.05, **P<0.01, ***P<0.001. Statistical analysis was performed using R version 3.2.2 or Prism (GraphPad).

### Bioinformatics Analysis

We analyzed the developmental change of level of all mRNAs translated in the mouse RGC axons in data set (GSE79352). The sequence reads were mapped to the mouse genome (mm10) using TopHat 2 version 2.0.12 with default settings, except for the “--read-realign-edit-dist 0” option. Transcript assembly and estimation of FPKM (Fragments Per Kilobase of transcript per Million fragments sequenced) values were performed using Cufflinks version 2.2.1. For RNA-seq analysis of frog RGC axons, the sequence reads were mapped to the X. laevis v9.2 genome (Xenbase) using HISAT2 2.1.0 with default settings. Transcript assembly and estimation of FPKM (Fragments Per Kilobase of transcript per Million fragments sequenced) values were performed using Cufflinks version 2.1.1. We only analyzed genes showing FPKM > 1 in both of two replicates. For analysis of SILAC, we first extracted all proteins that show a significant change (FDR < 0.01) by stimulation of any of three molecular cues, Netrin-1, Sema3a and BDNF. Then, we performed a GO enrichment analysis using DAVID 6.8 with default settings for BP-direct, MF-direct and CC-direct categories for each group of proteins whose translations are increased or decreased by each of three cues. We used all detected proteins as the background of the enrichment calculation.

### Quantification of Immunofluorescence

For quantitation of fluorescence intensity, the growth cone outline was traced on the phase contrast image using Volocity version 6.0.1 (PerkinElmer), then superimposed on the fluorescent image. The software calculated the fluorescent intensity within the growth cone, giving a measurement of pixel intensity per unit area. The growth cone outline was then placed in an adjacent area clear of cellular material to record the background fluorescent intensity. This reading was subtracted from the growth cone reading, yielding the background-corrected intensity.

### FRAP analysis

Quantification of fluorescence intensity was performed using Volocity software (PerkinElmer). At each time point, the outline of the growth cone was traced using phase contrast images. Mean gray values from the 488-channel were subsequently calculated as mean pixel intensity per unit area within the specified region of interest (ROI). This ROI was then placed in an adjacent area clear of cellular material to record the background fluorescent intensity. This reading was subtracted from the growth cone reading, yielding the background-corrected intensity. Unhealthy axons exhibiting signs of photo-toxicity after FRAP (characterized by blebbing, growth cone collapse and/or retraction) were excluded from analysis. In addition, only growth cones of axons extending more than 100μm from the eye explant were quantified to reduce effects of somal diffusion. Relative fluorescent recovery (R) at each time point was calculated by the formula: Rx = (Ix – Ipost) / (Ipre – Ipost). Where, Ix = normalised fluorescent intensity of the growth cone ROI at time point ‘x’, Ipre = normalised fluorescent intensity before photobleaching and Ipost = normalised fluorescent intensity immediately after photobleaching (t=0mins). Significance was tested using a two-way ANOVA.

### Single Molecule Translation Imaging Analysis

Translation event counting was performed by manual counting supplemented with the previously reported software-assisted automated event detection(Strohl et al., 2017), where localisations of individual protein translation events were retrieved using maximum likelihood estimation with a Gaussian model fit via the software package rapidSTORM. A threshold of ~6700 ADC, corresponding to ~500 photons per localization, was applied to filter out noise and non-Venus blinking events. This threshold was found by manual selection of Venus flashes and determination of the “average” photon budget of a single emitting Venus molecule. The tracking option of rapidSTORM was used to recombine photons emanating from the same Venus protein over multiple frames. All events in a small area around the growth cones were included for analysis due to the high mobility of filopodia. Results are normalised to the growth cone area, thus given as event/s/μm^2^.

### Branching Analysis

A filopodium was defined as a protrusion with length <5μm while a branch was defined as a protrusion with length >5μm (Drinjakovic et al., 2010; Hornberg et al., 2013; Kalous et al., 2014; Wong et al., 2017). Data were analyzed in PRISM 7 (GraphPad). Data are presented as mean and error bars represent SEM. ‘n’ represents the number of axons. *p<0.01, **p<0.01, ***p<0.001, #p<0.05, ##p<0.01, ###p<0.001. Details of statistic results such as p values, degree of freedom, and t/F values are presented in the figure legends. For axon arbor analysis, 3D projection of axon arbors acquired at 40X were semi-automatically traced through the z-axis using the Simple Neurite Tracer plugin(Longair et al., 2011) in Fiji. The resulting traces were then analyzed for the number and the length of axon branches as well as the Axon Complexity Index (ACI)(Marshak et al., 2007). These measured parameters were compared using one-way ANOVA with the Two-stage step-up method of Benjamini, Krieger and Yekutieli multiple comparisons test. Cumulative distribution curves of total branch number represent least-squares fits to a Gaussian function and were compared using Extra sum-of-squares F test. The proportions of simple (ACI<1.4) and complex (ACI≥1.4) arbors in different groups were compared using Fisher’s exact test (Drinjakovic et al., 2010). For analysis of branching dynamics, the numbers of protrusions added and removed were counted on the terminal 50μm of mGFP/mRFP-labeled RGC axons for 10 min (imaged at an interval of 30s) (Wong et al., 2017). The addition and removal of protrusions were then compared statistically. A paired t test was used for intragroup comparison and unpaired t test was used for intergroup comparisons.

### Western Blot Analysis

Developed films from Western blot detection were scanned and imported into FIJI. The color was inverted and the background corrected signals for Rps4x and β-catenin were measured. Measured Rps4x levels were then normalized to β-catenin to obtain a rationmetric readout. A paired t test was used to assess differences in Rps4X protein levels between control MO and Rps4X MO samples (*n* = 3 independent experiments).

## DATA AND SOFTWARE AVAILABILITY

The mass spectrometry proteomics data have been deposited to the ProteomeXchange Consortium via the PRIDE partner repository with the dataset identifier PXD011032.

## Author contributions

T.S., C.E.H. and H.J. designed the experiments. T.S. wrote the manuscript with support from M.K., H.H.-W.W., C.E.H and W.A.H. H.H.-W.W. performed *in vivo* knock down and branching analysis. J.Q.L. performed FRAP. J.Q.L., F.v.T, F.S. and C.F.K performed single molecule translation imaging. J.Q.L, T.S., and M.K., performed *in vitro* knock-down. T.S., M.K., J.M.C. and A.D. carried out *in vitro* retinal culture and SILAC. M.K., T.S., J.d.F.N., M.C. performed fractionation of axon samples. J.Q.L., M.K., and R.C. performed immunostaining. M.K. and T.S. performed biochemical experiments. T.S. performed bioinformatics analysis. C.E.H. and T.S. supervised the project.

## Acknowledgements

We apologize to authors of key papers, which we could not cite due to space limitations. We would like to thank J.K. Mooslehner for technical assistance. This work was supported by Wellcome Trust Programme Grant (085314/Z/08/Z), European Research Council Advanced Grant (322817) to C.E.H. and by the Netherlands Organization for Scientific Research (NWO Rubicon 019.161LW.033) to M.K. and the ‘Canada First Research Excellence Fund awarded to McGill University for the Healthy Brains for Healthy Lives initiative’ to H.H.-W.W.

## Supplementary Materials

Materials and Methods

Figures S1-S7

Supplementary Tables

