## Supplemental figures for "On-site ribosome remodeling by locally synthesized ribosomal proteins in axons"

### Supplemental Figure 1, related to Figure 1

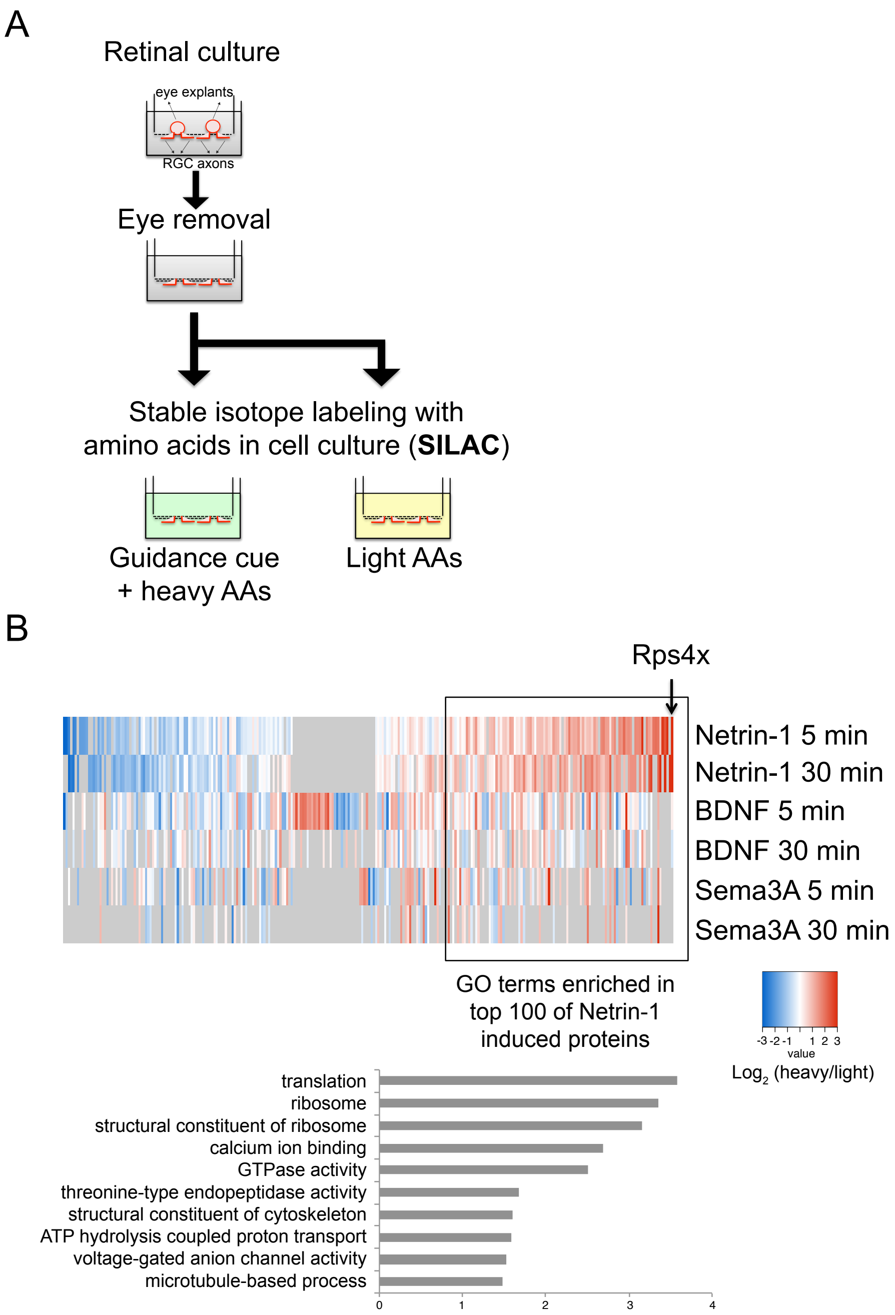


**Figure S1, Related to Figure 1**

(**A**) Diagram showing the strategy of the axon-pSILAC technique. (**B**) The heatmap (upper panel) shows log2 fold changes of levels of axonal protein synthesis that are increased(red) or decreased(blue) by treatment with three different molecular cues. This includes all proteins whose translation level is significantly changed by any of three cues. Rps4x shows the strongest increase of axonal translation after Netrin-1 treatment. The lower bar graph shows the result of a GO enrichment analysis (CC_direct, BP-direct and MF-direct) for the top 100 of Netrin-1-induced proteins using DAVID.

### Supplemental Figure 2, related to Figure 2

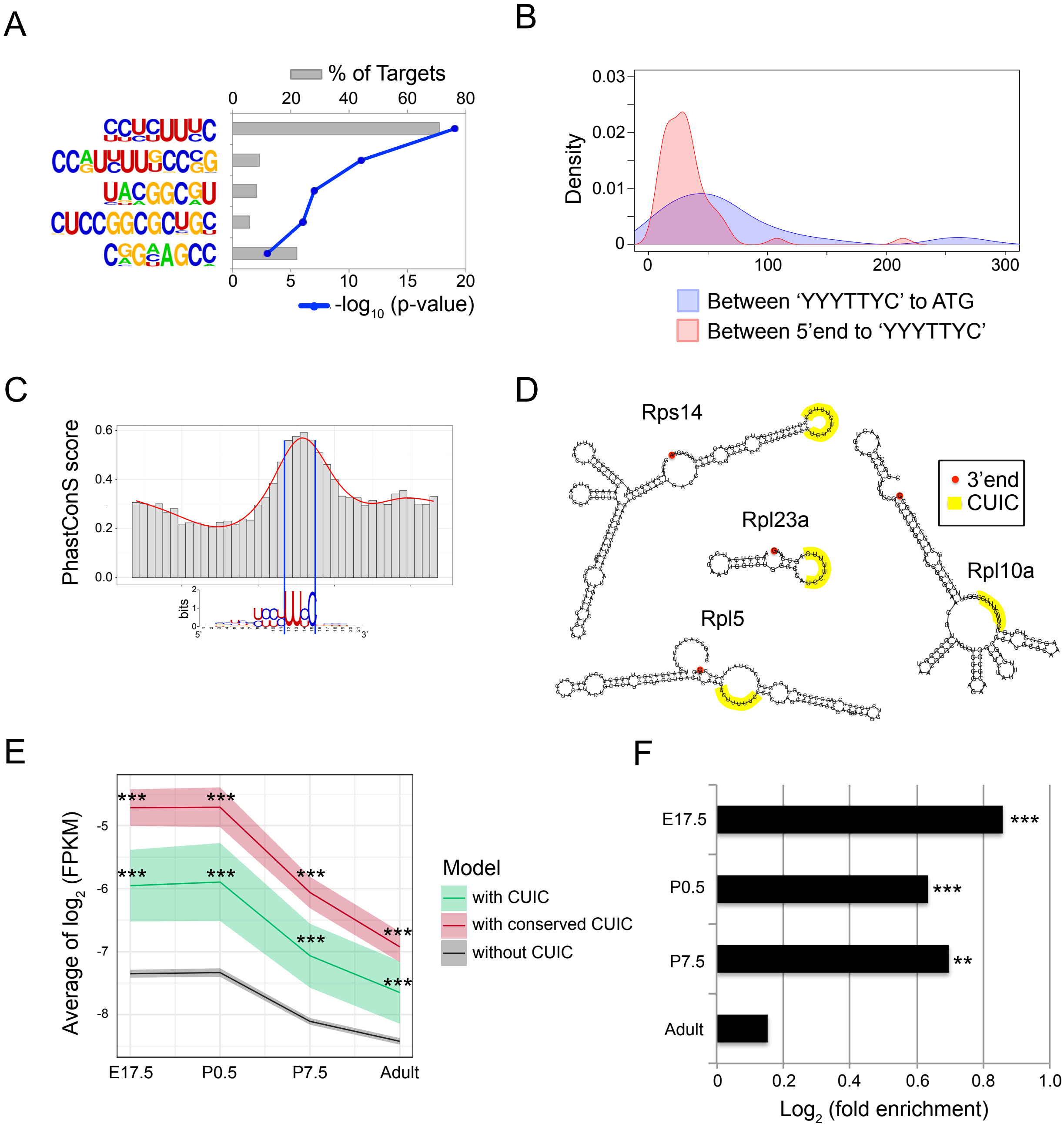


**Figure S2, Related to Figure 2**
(**A**) A graph showing the result of de novo motif finding for the 5' UTR of mouse RPs using HOMER. Grey bars show the percentage of motif-containing targets and the blue line represents the p-values. (**B**) Kernel density estimation of lengths between 5' end and CUIC, and between CUIC and the initiation codon. (**C**) Average interspecies conservation scores using PhastCons (mm10.60way.PhastCons) around the CUIC motif in mouse RP-coding mRNAs. (**D**) Examples of RNA secondary structures of CUIC containing 5' UTRs of RPs. (**E**) The graph shows the average and 95% CI of relative abundance (log2(FPKM)) of all genes with or without CUIC in mouse RGC axons of 4 different developmental stages (***p<0.001, Kolmogorov–Smirnov test, compared to "without CUIC"). (**F**) A bar graph representing the enrichment (log2(fold enrichment)) of CUIC motif containing genes in axonally translated genes (DEG) (*p<0.05, ** p<0.01, *** p<0.001, Fisher's exact test).

### Supplemental Figure 3, related to Figure 3

A

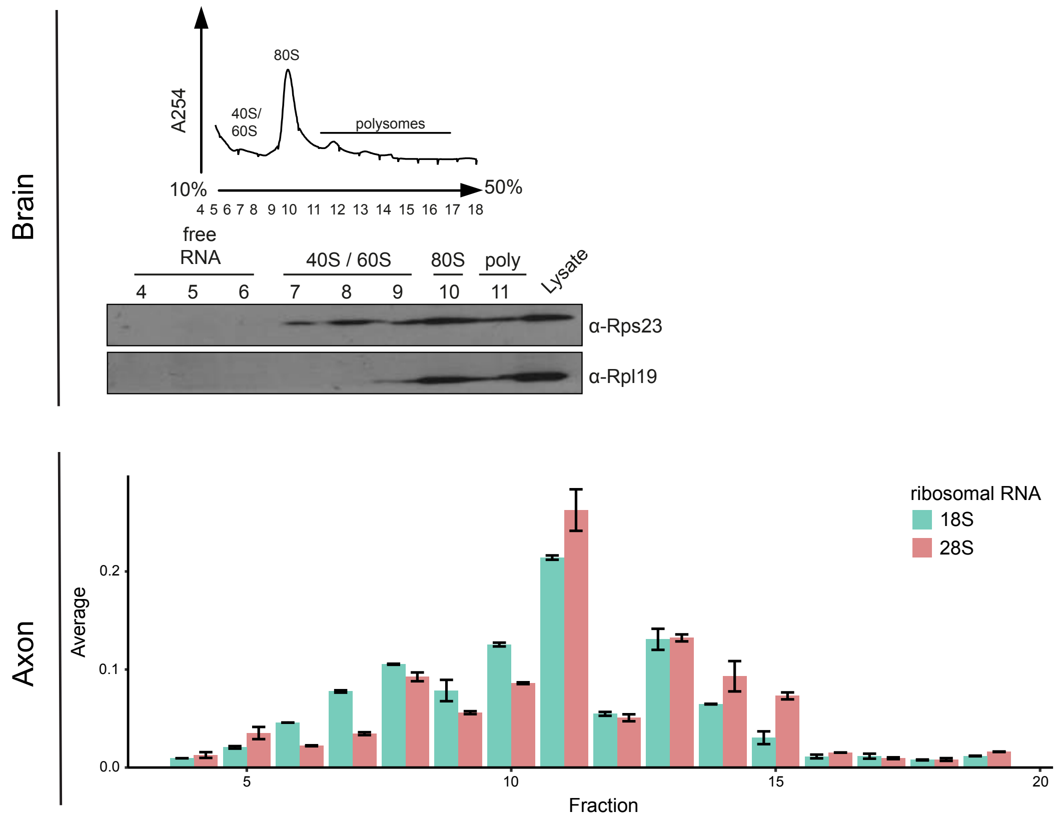


B

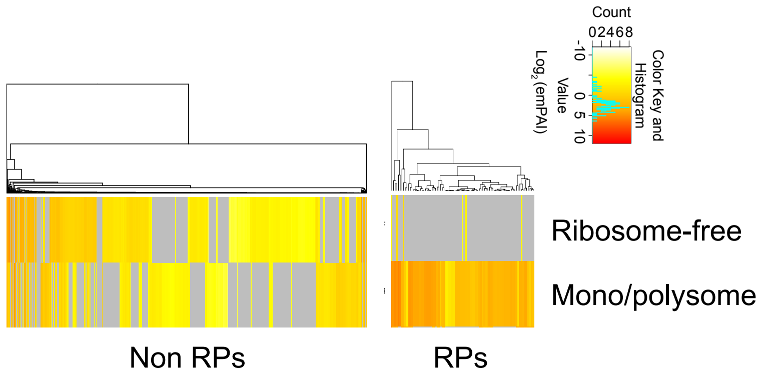


**Figure S3, Related to Figure 3**

(**A**) UV absorbance (A254) profile, immunoblot detection (Rps23 and Rpl19) and qRT-PCR detection of rRNAs in each fraction of the axon and whole brain sample. (**B**) Relative abundance of RPs and the other proteins detected in the mono/polysome fractions and the ribosome free fractions of whole brain sample.

### Supplemental Figure 4, related to Figure 4

A

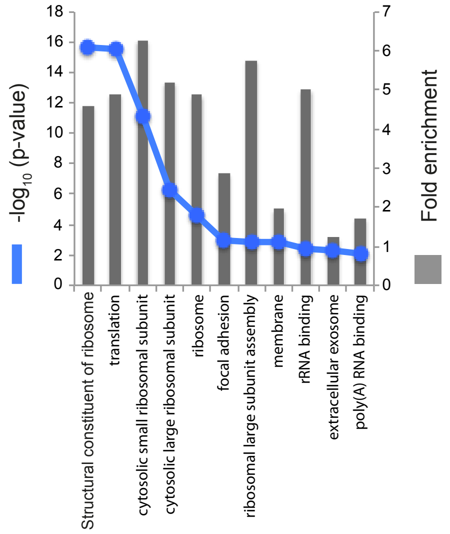


B

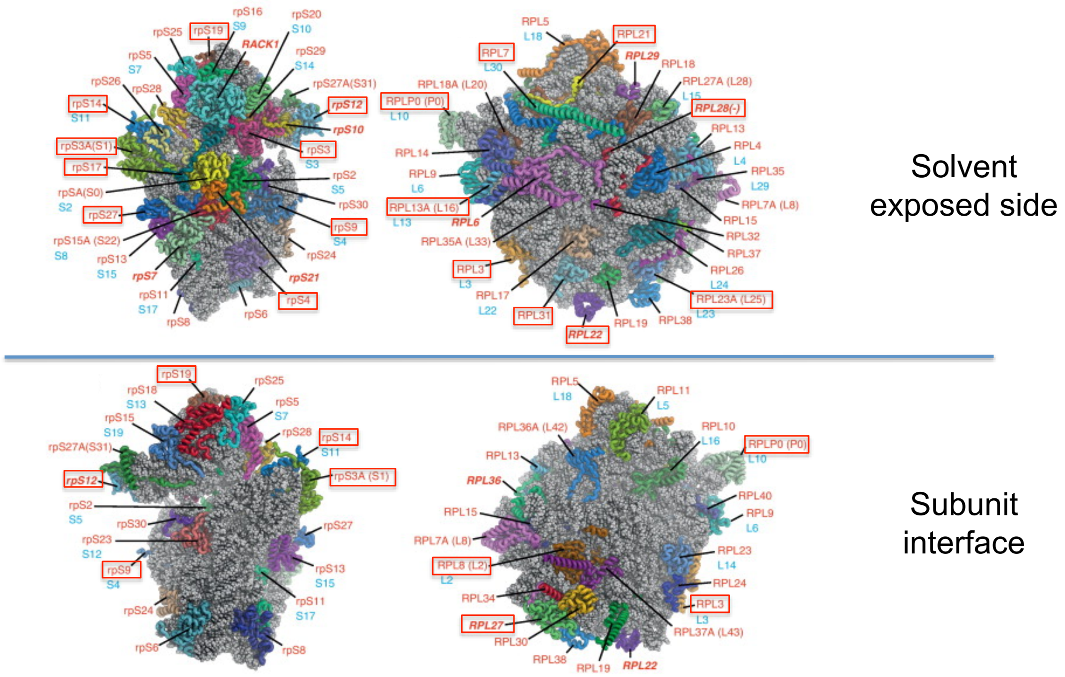


**Figure S4, Related to Figure 4**
(**A**) Enrichment (Blue line: Fisher's p, bars: fold enrichment) of GO terms in newly synthesized proteins in axons. (**B**) Structure of ribosomal subunits(Klinge et al., 2012) and positions of axonally incorporated RPs.

Supplemental Figure 5, related to Figure 5

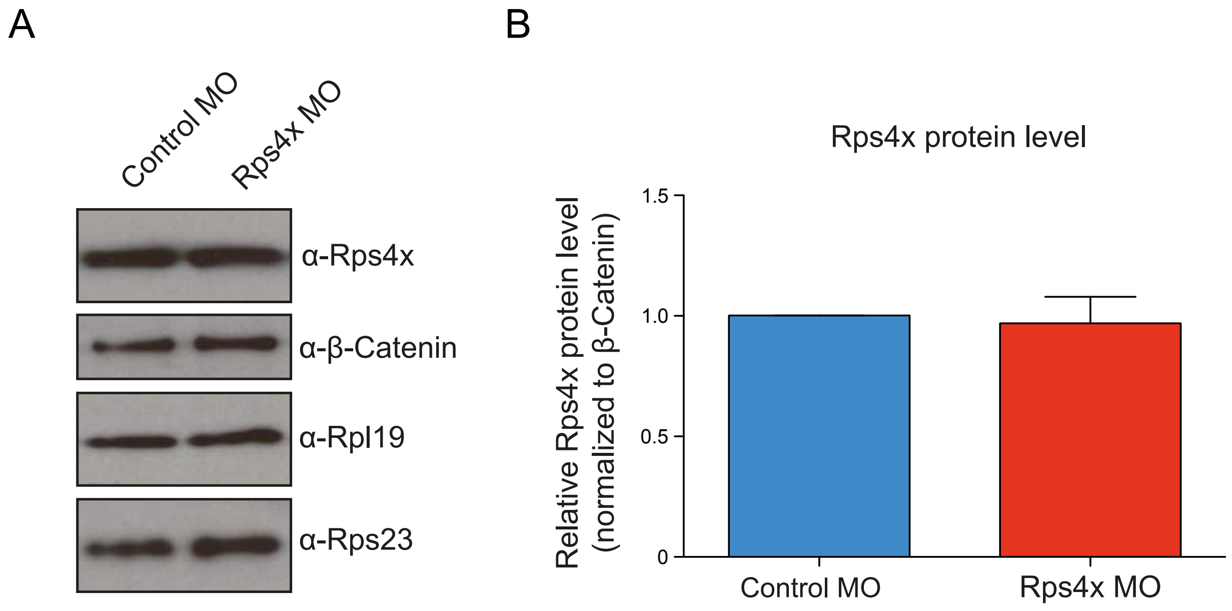


**Figure S5, Related to Figure 5**
(**A**) Western blots of protein samples isolated from the somal compartment from Control MO or Rps4X MO treated samples. (**B**) Bar graph showing Western blot quantification of Rps4x protein levels, normalized to β-Catenin protein levels. Error bars = SEM.

Supplemental Figure 6, related to Figure 6

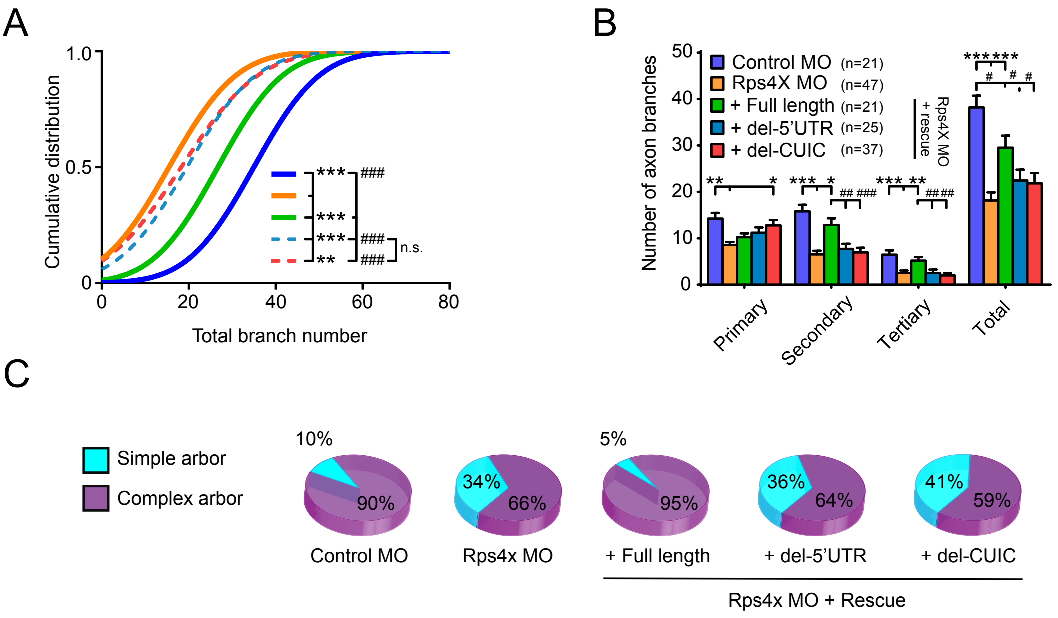


**Figure S6, Related to Figure 6**

(**A**) The cumulative distribution of total branch number (Extra sum-of-squares F test). (**B**) The proportion of branches in the Rps4x MO condition (primary:$F_{4,146}$=9.8, p<0.0001; secondary: $F_{4,146}$=5.7, p=0.0003; tertiary: $F_{4,146}$=4.6, p=0.002). (**C**) The percentage of complex arbor (ACI≥1.4) was reduced in Rps4x MO condition. Error bars represent SEM. Vs. rps4X MO: *p<0.05, **p<0.01, ***p<0.001, vs. Rps4X full length rescue: #p<0.05, ##p<0.01, ###p<0.001 (one-way ANOVA with Two-stage step-up method of Benjamini, Krieger and Yekutieli multiple comparisons test, n = 21 (Cont. MO), 47 (Cont. MO), 21 (MO+WT), 25 (MO+del-5' UTR), 37 (MO+del-CUIC) (c, d, e and g)).

### Supplemental Figure 7, related to Figure 7

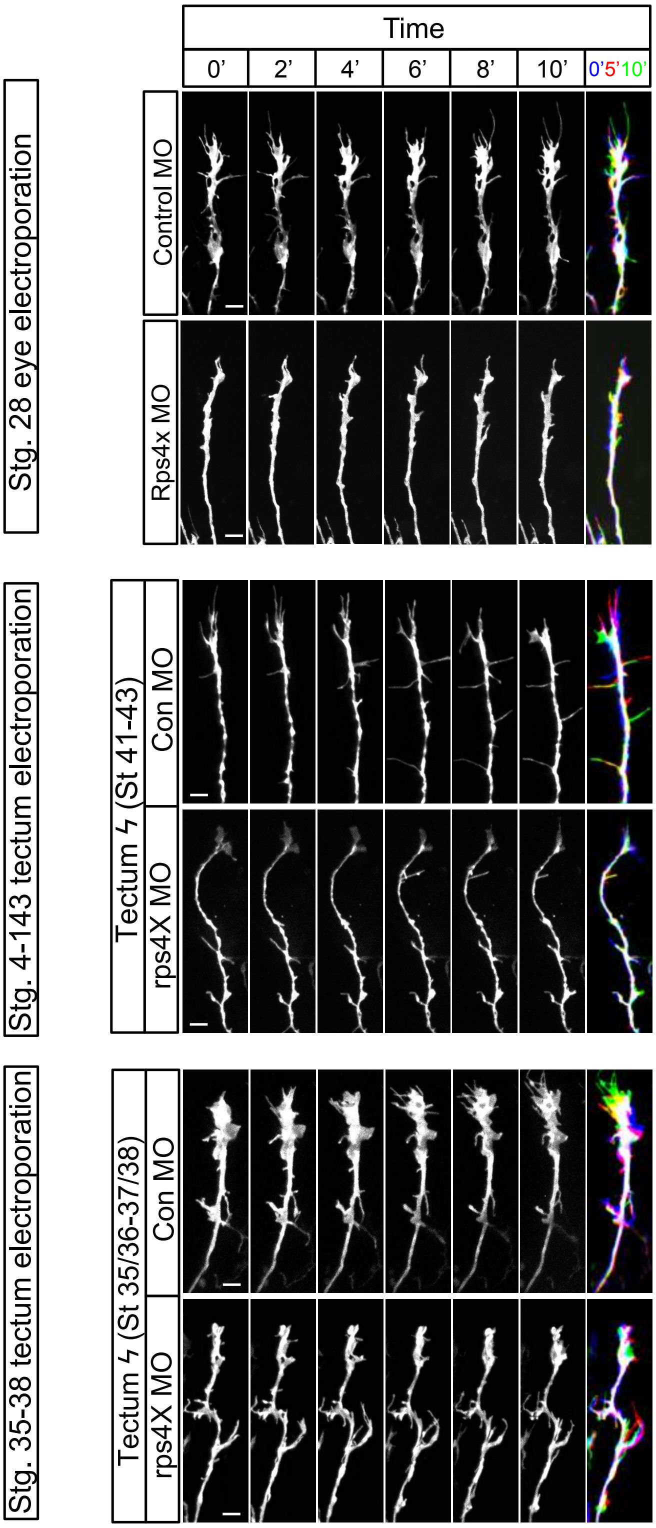


**Figure S7, Related to Figure 7**
Timelapse images of axonal branching in the tectum after eye electroporation (upper: stage 28) and tectum electroporation (middle: stages 41-43, lower: stage 35-38). scale bar = 20μm.
