## Supplemental tables for "On-site ribosome remodeling by locally synthesized ribosomal proteins in axons"

| **Table** | |
|---|---|
| CUIC_genes | All mouse genes with the CUIC motif and RNA secondary structures |
| Regulated_protein_syntesis | *X. laevis* proteins whose translation was significantly changed by any cue-stimulation tested |
| Enriched_GOs | GO terms enriched in gene groups whose translation was signifiacntly regulatd by cue stimulations |
| Axonal_RibosomeAssemblyFactors | All genes annotated with GO termes "nucleolus" or "ribosome biogenesis" that are detected in the axonal transcritome or proteome |

**CUIC_genes** All mouse genes with the CUIC motif and RNA secondary structures

structure_5UTR_in_RNAfold: Open parentheses: base is paired to another base ahead it, Closed parenthesees: indicate that a base is paired to another base behind it. Periods (or dots) indicate an unpaired base.

| Gene_name | gene | transcript | position | structure |
|---|---|---|---|---|
| 1110059E24Rik | ENSMUSG00000035171 | ENSMUST00000038830 | 214 | ) ) ) ) ) ) ) . |
| 1190007I07Rik | ENSMUSG00000063320 | ENSMUST00000079648 | 192 | . . . . . . . . |
| 1700007K09Rik | ENSMUSG00000030858 | ENSMUST00000188899 | 107 | . ) ) ) ) ) ) ) |
| 1700018B08Rik | ENSMUSG00000031809 | ENSMUST00000034265 | 181 | . . . . ( ( ( ( |
| 1700020A23Rik | ENSMUSG00000027409 | ENSMUST00000028898 | 207 | ( ( ( ( ( . . . |
| 1700029H14Rik | ENSMUSG00000031452 | ENSMUST00000134023 | 80 | ) ) ) ) ) ) . ) |
| 1700029H14Rik | ENSMUSG00000031452 | ENSMUST00000187391 | 80 | ) ) ) ) ) ) . ) |
| 1700066B19Rik | ENSMUSG00000073598 | ENSMUST00000097617 | 45 | ( ( ( ( ( ( . . |
| 1810062G17Rik | ENSMUSG00000027713 | ENSMUST00000029268 | 134 | . . . . . ) ) ) |
| 2210407C18Rik | ENSMUSG00000037145 | ENSMUST00000048801 | 3 | . . . . ( ( ( ( |
| 2310007B03Rik | ENSMUSG00000034159 | ENSMUST00000043718 | 196 | . . . . . . . . |
| 2310007B03Rik | ENSMUSG00000034159 | ENSMUST00000143419 | 205 | . . . . . . . ( |
| 2310061I04Rik | ENSMUSG00000050705 | ENSMUST00000148721 | 75 | . . ( ( ( ( . . |
| 2700049A03Rik | ENSMUSG00000034601 | ENSMUST00000149564 | 894 | . . ( ( ( ( ( ( |
| 3830417A13Rik | ENSMUSG00000031179 | ENSMUST00000033522 | 212 | . ) ) ) ) ) ) ) |
| 4922502D21Rik | ENSMUSG00000047720 | ENSMUST00000051283 | 178 | ( . . . . . . . |
| 4930467E23Rik | ENSMUSG00000096265 | ENSMUST00000098909 | 75 | ( ( ( ( ( ( ( ( |
| 4930503B20Rik | ENSMUSG00000090202 | ENSMUST00000049703 | 89 | . . ) ) ) ) ) ) |
| 4930522H14Rik | ENSMUSG00000060491 | ENSMUST00000082306 | 54 | . . ) ) ) . . ) |
| 4930550L24Rik | ENSMUSG00000046180 | ENSMUST00000062542 | 116 | ) ) ) ) ) ) ) . |
| 4930568D16Rik | ENSMUSG00000026882 | ENSMUST00000028243 | 60 | ) ) ) ) ) ) ) . |
| 4930578C19Rik | ENSMUSG00000037358 | ENSMUST00000044188 | 31 | . . . . . . . . |
| 4933408B17Rik | ENSMUSG00000049357 | ENSMUST00000056932 | 1183 | ) ) ) ) ) ) . . |
| 5330417C22Rik | ENSMUSG00000040412 | ENSMUST00000106625 | 13 | . . . . . . . . |
| 5530400C23Rik | ENSMUSG00000055594 | ENSMUST00000048459 | 24 | ) . ) ) ) ) ) ) |
| 6430531B16Rik | ENSMUSG00000073795 | ENSMUST00000097970 | 30 | ) ) ) . . . ) ) |
| 6820408C15Rik | ENSMUSG00000032680 | ENSMUST00000039961 | 411 | . . . . . ) ) ) |
| 9130019O22Rik | ENSMUSG00000030823 | ENSMUST00000049052 | 113 | ( . ( . . . ( ( |
| 9830107B12Rik | ENSMUSG00000073386 | ENSMUST00000063481 | 258 | . . . . . . . . |
| A1cf | ENSMUSG00000052595 | ENSMUST00000075838 | 61 | . . . . ) ) ) ) |
| A530016L24Rik | ENSMUSG00000043122 | ENSMUST00000057465 | 226 | ( ( ( ( ( ( ( ( |
| A530064D06Rik | ENSMUSG00000043939 | ENSMUST00000053612 | 228 | ) ) ) ) . . ) ) |
| A530064D06Rik | ENSMUSG00000043939 | ENSMUST00000027764 | 241 | ) ) ) ) . . ) ) |
| AA986860 | ENSMUSG00000042510 | ENSMUST00000039323 | 295 | . ) ) ) ) ) ) ) |
| Aak1 | ENSMUSG00000057230 | ENSMUST00000089519 | 466 | . ) ) ) ) ) ) ) |
| AB124611 | ENSMUSG00000057191 | ENSMUST00000086361 | 12 | . . . . . . . . |
| Abat | ENSMUSG00000057880 | ENSMUST00000115839 | 113 | ) ) ) ) ) ) ) ) |
| Abat | ENSMUSG00000057880 | ENSMUST00000065987 | 169 | ) ) ) ) ) ) ) ) |
| Abcb1a | ENSMUSG00000040584 | ENSMUST00000047753 | 82 | . ( ( ( ( ( ( ( |
| Abcb7 | ENSMUSG00000031333 | ENSMUST00000033695 | 5 | . . . . . . . . |
| Abcd2 | ENSMUSG00000055782 | ENSMUST00000069511 | 163 | ( . . . ( ( ( ( |
| Abhd14b | ENSMUSG00000042073 | ENSMUST00000048527 | 262 | . . . . . . . . |
| Abtb1 | ENSMUSG00000030083 | ENSMUST00000032169 | 65 | . ) ) ) . . ) ) |

| | | | | |
|---|---|---|---|---|
| Abtb2 | ENSMUSG00000032724 | ENSMUST00000076212 | 341 | ( ( . ( ( ( ( ( |
| AC121905.1 | ENSMUSG00000106923 | ENSMUST00000202173 | 39 | ) ) . . . ) ) ) |
| Acad11 | ENSMUSG00000090150 | ENSMUST00000047799 | 10 | ( . . . . . . . |
| Acd | ENSMUSG00000038000 | ENSMUST00000042608 | 29 | ( ( ( ( ( . . . |
| Acin1 | ENSMUSG00000022185 | ENSMUST00000126166 | 37 | ( ( ( . . ( ( ( |
| Acin1 | ENSMUSG00000022185 | ENSMUST00000141453 | 88 | ( ( ( . . ( ( ( |
| Acot13 | ENSMUSG00000006717 | ENSMUST00000006900 | 159 | ) . . . . . . . |
| Acsl1 | ENSMUSG00000018796 | ENSMUST00000110372 | 136 | . . . . . . ( ( |
| Actl7b | ENSMUSG00000070980 | ENSMUST00000095080 | 33 | ( . . ( ( ( ( ( |
| Actr1a | ENSMUSG00000025228 | ENSMUST00000040270 | 27 | ( ( ( ( ( ( ( . |
| Acvr1 | ENSMUSG00000026836 | ENSMUST00000112599 | 211 | . . . . . . . . |
| Acvr1 | ENSMUSG00000026836 | ENSMUST00000112601 | 377 | ) ) ) ) . ) ) ) |
| Acvrl1 | ENSMUSG00000000530 | ENSMUST00000120028 | 211 | ) ) ) . . ) . ) |
| Adam11 | ENSMUSG00000020926 | ENSMUST00000068150 | 104 | . . . . . . . . |
| Adam11 | ENSMUSG00000020926 | ENSMUST00000103081 | 104 | . . . . . . . . |
| Adam1b | ENSMUSG00000062438 | ENSMUST00000079368 | 385 | . . ( ( ( ( ( ( |
| Adam28 | ENSMUSG00000014725 | ENSMUST00000022642 | 64 | . . . ) ) ) ) ) |
| Adam28 | ENSMUSG00000014725 | ENSMUST00000111072 | 64 | . . . ) ) ) ) ) |
| Adam28 | ENSMUSG00000014725 | ENSMUST00000224039 | 64 | . . . ) ) ) ) ) |
| Adamts17 | ENSMUSG00000058145 | ENSMUST00000098382 | 1 | . . . . . . . . |
| Adamts8 | ENSMUSG00000031994 | ENSMUST00000068135 | 237 | . . . . . . . . |
| Adamtsl2 | ENSMUSG00000036040 | ENSMUST00000091233 | 152 | . ( ( ( ( ( ( . |
| Adat2 | ENSMUSG00000019808 | ENSMUST00000019944 | 274 | . . . . . ) ) ) |
| Adcy3 | ENSMUSG00000020654 | ENSMUST00000020984 | 140 | . . . . ) ) ) ) |
| Adcy3 | ENSMUSG00000020654 | ENSMUST00000152065 | 522 | . . . . ) ) ) ) |
| Adcyap1 | ENSMUSG00000024256 | ENSMUST00000064775 | 185 | . . ) ) ) ) . . |
| Adgrg1 | ENSMUSG00000031785 | ENSMUST00000093271 | 257 | ) ) ) ) ) ) ) ) |
| Adgrg7 | ENSMUSG00000022755 | ENSMUST00000023437 | 142 | . . . . . . . . |
| Adm | ENSMUSG00000030790 | ENSMUST00000033054 | 118 | . . . ) ) ) ) ) |
| Adora3 | ENSMUSG00000000562 | ENSMUST00000164730 | 214 | ) ) . . . . ) ) |
| Adrb3 | ENSMUSG00000031489 | ENSMUST00000081438 | 746 | ) . . ) ) ) ) ) |
| Adss | ENSMUSG00000015961 | ENSMUST00000016105 | 18 | ) . . . . . . . |
| Aebp1 | ENSMUSG00000020473 | ENSMUST00000102923 | 120 | ( ( ( . . . . . |
| Afm | ENSMUSG00000029369 | ENSMUST00000113179 | 40 | ) . . . . . . . |
| Aga | ENSMUSG00000031521 | ENSMUST00000033920 | 45 | . . . . . . . . |
| Aga | ENSMUSG00000031521 | ENSMUST00000211424 | 45 | . . . . . . . . |
| Agmo | ENSMUSG00000050103 | ENSMUST00000049874 | 324 | . ( . . . . . . |
| Agxt2 | ENSMUSG00000089678 | ENSMUST00000110542 | 39 | . . . . . . . . |
| AI464131 | ENSMUSG00000046312 | ENSMUST00000054920 | 126 | ( ( ( ( ( . . . |
| AI467606 | ENSMUSG00000045165 | ENSMUST00000056288 | 139 | ) ) ) ) ) ) . ) |
| Ak1 | ENSMUSG00000026817 | ENSMUST00000113278 | 90 | . . ( ( ( . . . |
| Ak6 | ENSMUSG00000078941 | ENSMUST00000022135 | 361 | ) ) ) ) ) . ) ) |
| Akap4 | ENSMUSG00000050089 | ENSMUST00000115751 | 63 | ( ( ( . . . . . |
| Akap9 | ENSMUSG00000040407 | ENSMUST00000044492 | 112 | ) ) . . . . . . |
| Akr1a1 | ENSMUSG00000028692 | ENSMUST00000030455 | 312 | ) ) ) . . . . . |
| Alg9 | ENSMUSG00000032059 | ENSMUST00000034561 | 346 | ) . ) ) ) . . . |
| Alkbh5 | ENSMUSG00000042650 | ENSMUST00000044250 | 375 | ( ( ( ( ( ( ( ( |
| Alms1 | ENSMUSG00000063810 | ENSMUST00000072018 | 81 | . . . . . . . . |
| Alox5ap | ENSMUSG00000060063 | ENSMUST00000071130 | 277 | ) ) ) ) ) ) ) ) |
| Alpl | ENSMUSG00000028766 | ENSMUST00000030551 | 193 | ( ( ( ( . ( ( ( |
| Ambn | ENSMUSG00000029288 | ENSMUST00000031226 | 28 | . . . ) ) ) ) ) |
| Ambn | ENSMUSG00000029288 | ENSMUST00000198265 | 48 | . . . ) ) ) ) ) |
| Ammecr1l | ENSMUSG00000041915 | ENSMUST00000115808 | 431 | . . . . . . . . |
| Ampd1 | ENSMUSG00000070385 | ENSMUST00000090715 | 36 | ) ) . . . . . . |
| Amph | ENSMUSG00000021314 | ENSMUST00000200466 | 5 | . . . . . . . . |
| Amph | ENSMUSG00000021314 | ENSMUST00000003345 | 151 | . . . . . . . . |
| Anapc11 | ENSMUSG00000025135 | ENSMUST00000093140 | 308 | ) ) . . . ) ) ) |
| Angptl1 | ENSMUSG00000033544 | ENSMUST00000027885 | 654 | . . . . . . ( ( |

| | | | | |
|---|---|---|---|---|
| Ankhd1 | ENSMUSG00000024483 | ENSMUST00000155329 | 35 | . ) ) ) ) ) ) ) |
| Ankrd17 | ENSMUSG00000055204 | ENSMUST00000081914 | 166 | . ) ) ) ) ) . ) |
| Ankrd17 | ENSMUSG00000055204 | ENSMUST00000014421 | 515 | ( ( . ( . . . . |
| Ankrd44 | ENSMUSG00000052331 | ENSMUST00000179030 | 141 | . ) ) ) ( ( ( ( |
| Ankrd6 | ENSMUSG00000040183 | ENSMUST00000084750 | 208 | . . . . . . . . |
| Ankrd6 | ENSMUSG00000040183 | ENSMUST00000084749 | 337 | ) . . . . . . . |
| Ankrd6 | ENSMUSG00000040183 | ENSMUST00000035719 | 447 | . . ) ) ) ) ) ) |
| Anln | ENSMUSG00000036777 | ENSMUST00000040912 | 138 | . ( ( ( . . . . |
| Anp32e | ENSMUSG00000015749 | ENSMUST00000171368 | 187 | ( ( ( ( ( ( ( ( |
| Anp32e | ENSMUSG00000015749 | ENSMUST00000015893 | 234 | ( ( ( ( ( ( ( ( |
| Anp32e | ENSMUSG00000015749 | ENSMUST00000165307 | 266 | ( ( ( ( ( ( ( ( |
| Anpep | ENSMUSG00000039062 | ENSMUST00000049004 | 52 | . . . . . . ) ) |
| Antxr2 | ENSMUSG00000029338 | ENSMUST00000031281 | 318 | ) ) ) . . . . . |
| Anxa2 | ENSMUSG00000032231 | ENSMUST00000034756 | 123 | . . . . . . . . |
| Anxa4 | ENSMUSG00000029994 | ENSMUST00000113675 | 89 | . . . . ( ( ( ( |
| Ap2a1 | ENSMUSG00000060279 | ENSMUST00000107857 | 8 | ( . . . . . . . |
| Ap2a1 | ENSMUSG00000060279 | ENSMUST00000166972 | 78 | . . . . ) ) ) ) |
| Ap5b1 | ENSMUSG00000049562 | ENSMUST00000096318 | 350 | . . ( ( . ( ( ( |
| Apbb1 | ENSMUSG00000037032 | ENSMUST00000191011 | 93 | . . . . . . . . |
| Aph1a | ENSMUSG00000015750 | ENSMUST00000056710 | 94 | . . . . . . ) ) |
| Aph1a | ENSMUSG00000015750 | ENSMUST00000015894 | 180 | . . . . . . ) ) |
| Apoc2 | ENSMUSG00000002992 | ENSMUST00000142352 | 216 | ) ) ) ) ) ) ) ) |
| Apoc3 | ENSMUSG00000032081 | ENSMUST00000118649 | 86 | . . ) ) ) ) ) ) |
| Apoc4 | ENSMUSG00000074336 | ENSMUST00000003071 | 30 | ( ( ( ( ( ( ( ( |
| Apol7b | ENSMUSG00000068252 | ENSMUST00000089469 | 68 | ) ) ) ) . . . . |
| Apol7c | ENSMUSG00000044309 | ENSMUST00000062562 | 79 | ( ( ( . . . . . |
| Apol7e | ENSMUSG00000071716 | ENSMUST00000096358 | 84 | ) ) ) ) . . . . |
| Apold1 | ENSMUSG00000090698 | ENSMUST00000167323 | 286 | ( ( ( ( . . . ( |
| Arc | ENSMUSG00000022602 | ENSMUST00000023268 | 132 | ( ( ( ( ( . . . |
| Arc | ENSMUSG00000022602 | ENSMUST00000110009 | 134 | . . . . . . ) ) |
| Arfgap3 | ENSMUSG00000054277 | ENSMUST00000067215 | 51 | . . . . . . . . |
| Arfip2 | ENSMUSG00000030881 | ENSMUST00000131446 | 200 | ) ) ) ) . . ) ) |
| Arhgap15 | ENSMUSG00000049744 | ENSMUST00000112824 | 1 | ( ( ( ( ( ( ( . |
| Arhgap24 | ENSMUSG00000057315 | ENSMUST00000073302 | 539 | ) ) ) ) ) ) ) ) |
| Arhgap25 | ENSMUSG00000030047 | ENSMUST00000101197 | 65 | ( ( ( ( ( . . . |
| Arhgap39 | ENSMUSG00000033697 | ENSMUST00000036176 | 97 | ) ) . ( ( ( ( ( |
| Arhgap39 | ENSMUSG00000033697 | ENSMUST00000077821 | 184 | ( ( ( ( ( ( ( ( |
| Arhgap9 | ENSMUSG00000040345 | ENSMUST00000219511 | 121 | . . . . . . . . |
| Arhgdib | ENSMUSG00000030220 | ENSMUST00000032344 | 22 | ( . . . . . . ( |
| Arhgef1 | ENSMUSG00000040940 | ENSMUST00000117419 | 73 | ) ) . . . . ) ) |
| Arhgef18 | ENSMUSG00000004568 | ENSMUST00000004684 | 119 | ) ) ) ) ) ) . ) |
| Arhgef26 | ENSMUSG00000036885 | ENSMUST00000079300 | 1090 | . . ) ) ) ) ) ) |
| Arid3c | ENSMUSG00000066224 | ENSMUST00000150809 | 25 | ( ( ( ( . . . . |
| Arid3c | ENSMUSG00000066224 | ENSMUST00000171251 | 56 | ( ( ( ( . . . . |
| Arl14 | ENSMUSG00000098207 | ENSMUST00000183126 | 14 | ( ( ( ( ( ( ( ( |
| Arl15 | ENSMUSG00000042348 | ENSMUST00000091201 | 100 | . . . . . . . . |
| Arl6 | ENSMUSG00000022722 | ENSMUST00000023405 | 185 | . ) ) ) ) ) ) ) |
| Arl6ip4 | ENSMUSG00000029404 | ENSMUST00000031351 | 318 | . . . . . . . ) |
| Armc10 | ENSMUSG00000038525 | ENSMUST00000072896 | 140 | . . ( ( ( . . . |
| Armc5 | ENSMUSG00000042178 | ENSMUST00000044660 | 703 | ( . ( ( ( ( ( ( |
| Armc6 | ENSMUSG00000002343 | ENSMUST00000019679 | 328 | . . . . . . . . |
| Armcx3 | ENSMUSG00000049047 | ENSMUST00000081834 | 430 | ) ) ) ) ) ) ) . |
| Armcx4 | ENSMUSG00000049804 | ENSMUST00000124226 | 521 | ) ) ) ) ) ) ) ) |
| Armcx6 | ENSMUSG00000050394 | ENSMUST00000052431 | 370 | ) ) ) ) . . ) ) |
| Armt1 | ENSMUSG00000061759 | ENSMUST00000095893 | 136 | ( ( ( ( . . . . |
| Arnt2 | ENSMUSG00000015709 | ENSMUST00000085077 | 219 | ) ) ) . ) ) ) ) |
| Arntl | ENSMUSG00000055116 | ENSMUST00000210074 | 196 | ) ) ) ) ) . . . |
| Arntl | ENSMUSG00000055116 | ENSMUST00000047321 | 511 | ) . ) ) ) ) ) ) |

| | | | | |
|---|---|---|---|---|
| Arpc2 | ENSMUSG00000006304 | ENSMUST00000006467 | 279 | . ) ) ) ) ) ) ) |
| Arpc3 | ENSMUSG00000029465 | ENSMUST00000102525 | 17 | ( ( ( . . . . . |
| Arr3 | ENSMUSG00000060890 | ENSMUST00000113769 | 74 | ( ( ( . ( ( ( ( |
| Arrdc2 | ENSMUSG00000002910 | ENSMUST00000002989 | 258 | . . . . . . . . |
| Arx | ENSMUSG00000035277 | ENSMUST00000046565 | 160 | . . ( ( ( ( ( . |
| Asb10 | ENSMUSG00000038204 | ENSMUST00000048302 | 96 | . . . . . . . . |
| Ascc1 | ENSMUSG00000044475 | ENSMUST00000050516 | 37 | . ) ) ) . . ) ) |
| Asgr2 | ENSMUSG00000040963 | ENSMUST00000102572 | 83 | ( . . . . . . . |
| Asic1 | ENSMUSG00000023017 | ENSMUST00000228185 | 115 | ( ( ( ( ( . ( ( |
| Asph | ENSMUSG00000028207 | ENSMUST00000108333 | 122 | ) ) ) ) ) ) ) ) |
| Asph | ENSMUSG00000028207 | ENSMUST00000108335 | 122 | ) ) ) ) ) ) ) ) |
| Asph | ENSMUSG00000028207 | ENSMUST00000108334 | 122 | ) ) ) ) ) ) ) ) |
| Asph | ENSMUSG00000028207 | ENSMUST00000103004 | 122 | ) ) ) ) ) ) ) ) |
| Asph | ENSMUSG00000028207 | ENSMUST00000108339 | 243 | ) ) ) ) ) ) ) ) |
| Astn2 | ENSMUSG00000028373 | ENSMUST00000084496 | 67 | . . . . . . . . |
| Astn2 | ENSMUSG00000028373 | ENSMUST00000068214 | 195 | . . . . . . . . |
| Asxl3 | ENSMUSG00000045215 | ENSMUST00000097655 | 360 | ) ) ) ) ) . ) ) |
| Atcay | ENSMUSG00000034958 | ENSMUST00000047408 | 378 | ) ) ) . . . . ) |
| Atp11b | ENSMUSG00000037400 | ENSMUST00000029257 | 182 | ) ) ) ) ) . ) ) |
| Atp1a1 | ENSMUSG00000033161 | ENSMUST00000036493 | 215 | . . . ( ( . . . |
| Atp2b2 | ENSMUSG00000030302 | ENSMUST00000101045 | 522 | . . . . . . . . |
| Atp2b2 | ENSMUSG00000030302 | ENSMUST00000089003 | 550 | ) . ) ) ) ) ) ) |
| Atp2b2 | ENSMUSG00000030302 | ENSMUST00000101044 | 559 | ) ) . . . . . . |
| Atp2c1 | ENSMUSG00000032570 | ENSMUST00000038118 | 321 | . . . . . . . . |
| Atp5h | ENSMUSG00000034566 | ENSMUST00000043931 | 16 | ( ( ( ( ( ( ( . |
| Atp5o | ENSMUSG00000022956 | ENSMUST00000023677 | 91 | ) . ) ) ) . . . |
| Atp6ap2 | ENSMUSG00000031007 | ENSMUST00000033313 | 52 | ) ) ) . . . . . |
| Atp6v0c | ENSMUSG00000024121 | ENSMUST00000024932 | 157 | . . . . . . . ) |
| Atpaf2 | ENSMUSG00000042709 | ENSMUST00000108721 | 115 | . ) ) ) ) ) ) ) |
| Atxn2l | ENSMUSG00000032637 | ENSMUST00000040202 | 224 | . . . . . . . . |
| Axl | ENSMUSG00000002602 | ENSMUST00000002677 | 136 | ) ) ) . ) ) ) ) |
| Axl | ENSMUSG00000002602 | ENSMUST00000085948 | 154 | ) ) ) . ) ) ) ) |
| B3gat1 | ENSMUSG00000045994 | ENSMUST00000161115 | 122 | . . . . . . . . |
| B3gnt8 | ENSMUSG00000059479 | ENSMUST00000076034 | 97 | ( ( ( . . . . . |
| B430306N03Rik | ENSMUSG00000043740 | ENSMUST00000049614 | 246 | ) ) . . . . . . |
| B9d2 | ENSMUSG00000063439 | ENSMUST00000108403 | 214 | ) ) ) ) ) ) ) ) |
| Bach2 | ENSMUSG00000040270 | ENSMUST00000171600 | 81 | ( ( . . . . . . |
| Baiap2 | ENSMUSG00000025372 | ENSMUST00000026436 | 347 | ) ) . . . . . . |
| Baiap2 | ENSMUSG00000025372 | ENSMUST00000075180 | 347 | ) ) . . . . . . |
| Baiap2 | ENSMUSG00000025372 | ENSMUST00000103021 | 347 | ) ) . . . . . . |
| Banp | ENSMUSG00000025316 | ENSMUST00000026354 | 119 | ) . ) ) . ) ) ) |
| Banp | ENSMUSG00000025316 | ENSMUST00000170857 | 155 | ) . ) ) . ) ) ) |
| Banp | ENSMUSG00000025316 | ENSMUST00000093078 | 207 | ) ) ) ) ) ) ) ) |
| Batf2 | ENSMUSG00000039699 | ENSMUST00000045042 | 55 | ) ) ) ) ) ) ) ) |
| Baz2b | ENSMUSG00000026987 | ENSMUST00000112550 | 202 | . ) ) ) ) ) ) ) |
| BB287469 | ENSMUSG00000079031 | ENSMUST00000110149 | 484 | . . . . . . ) ) |
| Bbc3 | ENSMUSG00000002083 | ENSMUST00000002152 | 215 | ) ) ) ) ) ) ) ) |
| BC048679 | ENSMUSG00000061877 | ENSMUST00000073406 | 0 | ( ( ( ( ( ( ( . |
| BC048679 | ENSMUSG00000061877 | ENSMUST00000144156 | 0 | ( ( ( ( ( ( ( . |
| Bckdhb | ENSMUSG00000032263 | ENSMUST00000190166 | 115 | ) ) ) . . ) ) ) |
| Bcl11a | ENSMUSG00000000861 | ENSMUST00000109514 | 132 | ( . . . . . . . |
| Bcl11a | ENSMUSG00000000861 | ENSMUST00000109516 | 188 | ( . . . . . . . |
| Bcl11a | ENSMUSG00000000861 | ENSMUST00000000881 | 249 | ( . . . . . . . |
| Bcl2 | ENSMUSG00000057329 | ENSMUST00000189999 | 283 | ( ( ( ( ( . . . |
| Bcl2 | ENSMUSG00000057329 | ENSMUST00000112751 | 1378 | ) . . . . . . . |
| Bcl7a | ENSMUSG00000029438 | ENSMUST00000031391 | 148 | ) ) ) . . . . . |
| Bcs1l | ENSMUSG00000026172 | ENSMUST00000113733 | 167 | ) ) ) ) ) ) ) ) |
| Bcs1l | ENSMUSG00000026172 | ENSMUST00000027358 | 250 | ) ) ) ) ) ) . . |

| | | | | |
|---|---|---|---|---|
| Best3 | ENSMUSG00000020169 | ENSMUST00000020378 | 468 | ) . ) ) . ) ) ) |
| Bet1 | ENSMUSG00000032757 | ENSMUST00000049166 | 129 | ) . ) . . ) ) ) |
| Bglap3 | ENSMUSG00000074489 | ENSMUST00000075523 | 205 | . ) ) ) ) . ) ) |
| Bhlhe40 | ENSMUSG00000030103 | ENSMUST00000032194 | 208 | . . . . . ) ) ) |
| Bicd1 | ENSMUSG00000003452 | ENSMUST00000111513 | 372 | . . . . . . . . |
| Bicd1 | ENSMUSG00000003452 | ENSMUST00000086829 | 404 | . . . . . . . . |
| Birc3 | ENSMUSG00000032000 | ENSMUST00000013949 | 268 | . ) ) ) ) ) ) ) |
| Bivm | ENSMUSG00000041684 | ENSMUST00000114709 | 396 | ) . . . ) ) ) . |
| Bmi1 | ENSMUSG00000026739 | ENSMUST00000028071 | 442 | . ) ) ) ) ) ) ) |
| Bmp1 | ENSMUSG00000022098 | ENSMUST00000022693 | 86 | ) ) ) ) . . ) ) |
| Bmp2 | ENSMUSG00000027358 | ENSMUST00000028836 | 1120 | ) . . . . ( ( ( |
| Bmp3 | ENSMUSG00000029335 | ENSMUST00000200388 | 597 | ( ( ( . . . . ( |
| Bmp3 | ENSMUSG00000029335 | ENSMUST00000031278 | 609 | ( ( ( . . . . ( |
| Bmp5 | ENSMUSG00000032179 | ENSMUST00000012281 | 673 | ( ( ( . . ( ( ( |
| Bmpr2 | ENSMUSG00000067336 | ENSMUST00000087435 | 1214 | . . . . . . . . |
| Bmt2 | ENSMUSG00000042742 | ENSMUST00000045235 | 47 | ( ( ( ( ( ( ( ( |
| Bpi | ENSMUSG00000052922 | ENSMUST00000065039 | 105 | . . . . . . ) ) |
| Bpifb6 | ENSMUSG00000068009 | ENSMUST00000088955 | 71 | ( ( . . . . . . |
| Braf | ENSMUSG00000002413 | ENSMUST00000002487 | 113 | . . . . . . . . |
| Brca1 | ENSMUSG00000017146 | ENSMUST00000017290 | 92 | ) ) ) ) ( ( ( ( |
| Bri3bp | ENSMUSG00000037905 | ENSMUST00000049040 | 56 | . . . . . ( ( ( |
| Btbd11 | ENSMUSG00000020042 | ENSMUST00000105306 | 67 | . ) ) ) ) ) ) ) |
| Btbd11 | ENSMUSG00000020042 | ENSMUST00000105307 | 467 | . . . . ) ) ) ) |
| Btbd2 | ENSMUSG00000003344 | ENSMUST00000003434 | 512 | . . . . . . . . |
| Btd | ENSMUSG00000021900 | ENSMUST00000090147 | 13 | ( ( . . . . ( ( |
| Btn1a1 | ENSMUSG00000000706 | ENSMUST00000041674 | 67 | . ) ) ) . ) ) ) |
| Btnl1 | ENSMUSG00000062638 | ENSMUST00000080254 | 266 | . . . . ) ) ) ) |
| Bzw1 | ENSMUSG00000051223 | ENSMUST00000050552 | 294 | ( . . . . . . . |
| C1qtnf2 | ENSMUSG00000046491 | ENSMUST00000057679 | 102 | ) ) ) ) ) ) ) . |
| C2 | ENSMUSG00000024371 | ENSMUST00000025230 | 33 | . ( ( ( ( ( ( ( |
| C3 | ENSMUSG00000024164 | ENSMUST00000024988 | 87 | . . . . . . . . |
| C330021F23Rik | ENSMUSG00000065952 | ENSMUST00000136592 | 622 | . ) . ) ) ) ) ) |
| Cacna1c | ENSMUSG00000051331 | ENSMUST00000078320 | 190 | ) ) ) ) ) ) ) ( |
| Cacna1c | ENSMUSG00000051331 | ENSMUST00000112790 | 237 | ) ) ) . . . ( ( |
| Calm1 | ENSMUSG00000001175 | ENSMUST00000110082 | 259 | ) ) ) . ) ) ) ) |
| Calr4 | ENSMUSG00000028558 | ENSMUST00000030285 | 39 | ) ) . . . . . . |
| Calr4 | ENSMUSG00000028558 | ENSMUST00000106629 | 361 | . . . . . ) ) ) |
| Calr4 | ENSMUSG00000028558 | ENSMUST00000106631 | 613 | . . . . . ) ) ) |
| Camk1d | ENSMUSG00000039145 | ENSMUST00000044009 | 231 | . . . ) ) ) ) ) |
| Capn1 | ENSMUSG00000024942 | ENSMUST00000025891 | 130 | ) ) ) ) . . ) ) |
| Capn2 | ENSMUSG00000026509 | ENSMUST00000068505 | 132 | . ( ( . . . . . |
| Capn3 | ENSMUSG00000079110 | ENSMUST00000028749 | 273 | . . . . . . . . |
| Capn3 | ENSMUSG00000079110 | ENSMUST00000110721 | 273 | . . . . . . . . |
| Capn9 | ENSMUSG00000031981 | ENSMUST00000093033 | 4 | . . . ( ( ( ( ( |
| Caprin1 | ENSMUSG00000027184 | ENSMUST00000111147 | 133 | ( ( ( . . . ( ( |
| Capza1 | ENSMUSG00000070372 | ENSMUST00000094028 | 73 | ) ) . . . . ( ( |
| Car10 | ENSMUSG00000056158 | ENSMUST00000107863 | 882 | ) ) . . . . ) ) |
| Cartpt | ENSMUSG00000021647 | ENSMUST00000224142 | 378 | ) ) ) . . ) ) ) |
| Casd1 | ENSMUSG00000015189 | ENSMUST00000015333 | 285 | ) ) ) ) ) ) ) ) |
| Caskin2 | ENSMUSG00000034471 | ENSMUST00000041684 | 473 | ) . . ) . ) ) ) |
| Cav1 | ENSMUSG00000007655 | ENSMUST00000007799 | 43 | ( . . . . . . . |
| Ccdc136 | ENSMUSG00000029769 | ENSMUST00000181464 | 210 | ) . ) ) ) ) ) ) |
| Ccdc136 | ENSMUSG00000029769 | ENSMUST00000180829 | 210 | ) . ) ) ) ) ) ) |
| Ccdc27 | ENSMUSG00000039492 | ENSMUST00000047207 | 15 | . . . ( ( ( ( ( |
| Ccdc54 | ENSMUSG00000050685 | ENSMUST00000062439 | 198 | ) ) ) ) ) ) ) ) |
| Ccdc60 | ENSMUSG00000043913 | ENSMUST00000050178 | 438 | ) ) ) ) ) ) ) ) |
| Ccdc65 | ENSMUSG00000003354 | ENSMUST00000003444 | 112 | . . . . . . . . |
| Ccdc83 | ENSMUSG00000030617 | ENSMUST00000107221 | 34 | ) . ) ) ) ) . ) |

| | | | | |
|---|---|---|---|---|
| Ccdc83 | ENSMUSG00000030617 | ENSMUST00000040413 | 48 | ).)))).. |
| Ccdc88b | ENSMUSG00000047810 | ENSMUST00000113440 | 8 | .((((... |
| Ccl2 | ENSMUSG00000035385 | ENSMUST00000000193 | 74 | ))))))). |
| Ccl24 | ENSMUSG00000004814 | ENSMUST00000004936 | 25 | .....))) |
| Ccl3 | ENSMUSG00000000982 | ENSMUST00000001008 | 28 | ........ |
| Ccl7 | ENSMUSG00000035373 | ENSMUST00000021011 | 58 | ........ |
| Ccr6 | ENSMUSG00000040899 | ENSMUST00000177568 | 111 | )....... |
| Ccr6 | ENSMUSG00000040899 | ENSMUST00000180103 | 202 | ...))).) |
| Ccr6 | ENSMUSG00000040899 | ENSMUST00000164411 | 330 | ...))).) |
| Ccr6 | ENSMUSG00000040899 | ENSMUST00000097418 | 532 | )..)..)) |
| Ccr9 | ENSMUSG00000029530 | ENSMUST00000168910 | 80 | .......) |
| Cct2 | ENSMUSG00000034024 | ENSMUST00000047672 | 34 | ........ |
| Cct5 | ENSMUSG00000022234 | ENSMUST00000022842 | 40 | ........ |
| Cd207 | ENSMUSG00000034783 | ENSMUST00000037882 | 7 | (((((((. |
| Cd247 | ENSMUSG00000005763 | ENSMUST00000027849 | 32 | (....... |
| Cd247 | ENSMUSG00000005763 | ENSMUST00000005907 | 66 | (....... |
| Cd247 | ENSMUSG00000005763 | ENSMUST00000161971 | 74 | (....... |
| Cd276 | ENSMUSG00000035914 | ENSMUST00000165365 | 545 | ))...))) |
| Cd2bp2 | ENSMUSG00000042502 | ENSMUST00000166791 | 42 | .......) |
| Cd300lg | ENSMUSG00000017309 | ENSMUST00000017453 | 0 | ...((((. |
| Cd300lg | ENSMUSG00000017309 | ENSMUST00000107164 | 0 | ...((((. |
| Cd300lg | ENSMUSG00000017309 | ENSMUST00000107163 | 0 | ...((((. |
| Cd33 | ENSMUSG00000004609 | ENSMUST00000205503 | 31 | (((((... |
| Cd37 | ENSMUSG00000030798 | ENSMUST00000098461 | 40 | ........ |
| Cd38 | ENSMUSG00000029084 | ENSMUST00000030964 | 284 | )))))))) |
| Cd44 | ENSMUSG00000005087 | ENSMUST00000060516 | 104 | ........ |
| Cd44 | ENSMUSG00000005087 | ENSMUST00000099673 | 119 | ........ |
| Cd44 | ENSMUSG00000005087 | ENSMUST00000005218 | 234 | ........ |
| Cd46 | ENSMUSG00000016493 | ENSMUST00000162650 | 5 | ........ |
| Cd55b | ENSMUSG00000026401 | ENSMUST00000119432 | 33 | ......)) |
| Cd55b | ENSMUSG00000026401 | ENSMUST00000112488 | 33 | ......)) |
| Cd5l | ENSMUSG00000015854 | ENSMUST00000015998 | 34 | ))...... |
| Cd68 | ENSMUSG00000018774 | ENSMUST00000018918 | 14 | ((((.((( |
| Cdc14b | ENSMUSG00000033102 | ENSMUST00000109769 | 110 | ........ |
| Cdc40 | ENSMUSG00000038446 | ENSMUST00000044166 | 189 | (....... |
| Cdc42ep3 | ENSMUSG00000036533 | ENSMUST00000068958 | 460 | ...))))) |
| Cdca2 | ENSMUSG00000048922 | ENSMUST00000163100 | 293 | ).)))... |
| Cdcp2 | ENSMUSG00000047636 | ENSMUST00000062495 | 743 | .))))))) |
| Cdh16 | ENSMUSG00000031881 | ENSMUST00000211903 | 66 | ...))))) |
| Cdh16 | ENSMUSG00000031881 | ENSMUST00000212882 | 148 | ((((.... |
| Cdh16 | ENSMUSG00000031881 | ENSMUST00000163783 | 199 | ..(((((( |
| Cdh19 | ENSMUSG00000047216 | ENSMUST00000094626 | 187 | (((..... |
| Cdh7 | ENSMUSG00000026312 | ENSMUST00000027542 | 162 | (((((((( |
| Cdh7 | ENSMUSG00000026312 | ENSMUST00000172005 | 313 | ......)) |
| Cdhr2 | ENSMUSG00000034918 | ENSMUST00000037145 | 54 | ).)).)). |
| Cdhr5 | ENSMUSG00000025497 | ENSMUST00000080654 | 31 | ((...... |
| Cdhr5 | ENSMUSG00000025497 | ENSMUST00000167263 | 43 | ....)))) |
| Cdk11b | ENSMUSG00000029062 | ENSMUST00000067081 | 24 | (.((.((( |
| Cdk12 | ENSMUSG00000003119 | ENSMUST00000107538 | 0 | (((((((( |
| Cdk12 | ENSMUSG00000003119 | ENSMUST00000003203 | 21 | ....)))) |
| Cdk12 | ENSMUSG00000003119 | ENSMUST00000107539 | 276 | ....)))) |
| Cdk13 | ENSMUSG00000041297 | ENSMUST00000223490 | 363 | ((((.... |
| Cdkal1 | ENSMUSG00000006191 | ENSMUST00000091674 | 78 | (((..((( |
| Cdkal1 | ENSMUSG00000006191 | ENSMUST00000006353 | 89 | (((..((( |
| Cdkl2 | ENSMUSG00000029403 | ENSMUST00000113143 | 479 | (....... |
| Cdkn1b | ENSMUSG00000003031 | ENSMUST00000003115 | 468 | ......)) |
| Cdkn2c | ENSMUSG00000028551 | ENSMUST00000063531 | 197 | )))))))) |
| Ceacam5 | ENSMUSG00000008789 | ENSMUST00000081907 | 172 | (((((((( |

| | | | | |
|---|---|---|---|---|
| Cela3b | ENSMUSG00000023433 | ENSMUST00000102522 | 13 | . . . . . . . . |
| Celf2 | ENSMUSG00000002107 | ENSMUST00000182706 | 61 | ) . ) . . . . ) |
| Celf2 | ENSMUSG00000002107 | ENSMUST00000183209 | 121 | ) . ) ) ) ) ) ) |
| Cep19 | ENSMUSG00000035790 | ENSMUST00000115168 | 391 | . . . . . . ) ) |
| Cep295nl | ENSMUSG00000076433 | ENSMUST00000103024 | 207 | . . . ) ) ) ) ) |
| Ces1c | ENSMUSG00000057400 | ENSMUST00000034189 | 2 | . . . . . . . . |
| Cfap20 | ENSMUSG00000031796 | ENSMUST00000034249 | 226 | . . . . . . ( ( |
| Cfl1 | ENSMUSG00000056201 | ENSMUST00000209469 | 125 | . ) ) ) ) ) ) ) |
| Cfp | ENSMUSG00000001128 | ENSMUST00000001156 | 24 | . . . . . . ) ) |
| Chd5 | ENSMUSG00000005045 | ENSMUST00000030775 | 278 | . . . . . . ) ) |
| Chd5 | ENSMUSG00000005045 | ENSMUST00000164662 | 278 | . . . . . . ) ) |
| Chga | ENSMUSG00000021194 | ENSMUST00000021610 | 131 | . . . . . . . . |
| Chic1 | ENSMUSG00000031327 | ENSMUST00000116547 | 14 | . . . . . . . . |
| Chl1 | ENSMUSG00000030077 | ENSMUST00000066905 | 215 | ) . ) ) ) ) ) ) |
| Chmp3 | ENSMUSG00000053119 | ENSMUST00000059462 | 98 | ( ( ( ( ( ( ( ( |
| Chp1 | ENSMUSG00000014077 | ENSMUST00000014221 | 90 | ( ( ( ( ( ( ( . |
| Chrnb3 | ENSMUSG00000031492 | ENSMUST00000079463 | 129 | ( ( . ( ( ( ( ( |
| Chrnb3 | ENSMUSG00000031492 | ENSMUST00000060943 | 200 | ( ( . ( ( ( ( ( |
| Cisd3 | ENSMUSG00000078695 | ENSMUST00000107583 | 138 | . . . . . . . . |
| Cish | ENSMUSG00000032578 | ENSMUST00000168260 | 253 | . . . . . . . . |
| Cish | ENSMUSG00000032578 | ENSMUST00000085102 | 937 | . . . . . . . . |
| CK137956 | ENSMUSG00000028813 | ENSMUST00000030614 | 103 | . . ) ) ) ) ) . |
| Clca1 | ENSMUSG00000028255 | ENSMUST00000029919 | 6 | ( . ( ( ( ( ( ( |
| Clcn5 | ENSMUSG00000004317 | ENSMUST00000004428 | 142 | ( ( ( . ( ( ( ( |
| Cldn16 | ENSMUSG00000038148 | ENSMUST00000161053 | 324 | . . . . . . . . |
| Cldn17 | ENSMUSG00000055811 | ENSMUST00000069549 | 102 | ( . ( ( ( . . . |
| Cldn19 | ENSMUSG00000066058 | ENSMUST00000084309 | 65 | . . . . . . . . |
| Cldnd2 | ENSMUSG00000038973 | ENSMUST00000040227 | 57 | . . . ) ) ) ) ) |
| Clec12a | ENSMUSG00000053063 | ENSMUST00000065289 | 411 | ) ) ) ) ) ) . . |
| Clip4 | ENSMUSG00000024059 | ENSMUST00000024854 | 79 | ( ( ( ( ( . ( ( |
| Clrn3 | ENSMUSG00000050866 | ENSMUST00000053716 | 136 | . . ) ) ) ) ) ) |
| Cltc | ENSMUSG00000047126 | ENSMUST00000103186 | 181 | . . . . . . . . |
| Cmc2 | ENSMUSG00000014633 | ENSMUST00000148235 | 70 | . . . . . . ) ) |
| Cmtm3 | ENSMUSG00000031875 | ENSMUST00000034343 | 72 | ) ) ) ) ) ) ) ) |
| Cmtm5 | ENSMUSG00000040759 | ENSMUST00000227441 | 99 | . ( ( ( . . . . |
| Cmtm5 | ENSMUSG00000040759 | ENSMUST00000037814 | 335 | ) ) ) ) ) . ) ) |
| Cnga4 | ENSMUSG00000030897 | ENSMUST00000033187 | 35 | ( ( ( ( ( . . . |
| Cnppd1 | ENSMUSG00000033159 | ENSMUST00000190679 | 166 | ) ) ) ) ) . ) ) |
| Cnr1 | ENSMUSG00000044288 | ENSMUST00000057188 | 542 | ) ) ) ) ) ) ( ( |
| Cnrip1 | ENSMUSG00000044629 | ENSMUST00000058159 | 260 | . . . . . . . . |
| Cntn6 | ENSMUSG00000030092 | ENSMUST00000089215 | 386 | ) ) ) ) ) ) ) ) |
| Col13a1 | ENSMUSG00000058806 | ENSMUST00000105453 | 342 | ) ) ) ) ) ) ) ) |
| Col13a1 | ENSMUSG00000058806 | ENSMUST00000105454 | 377 | ) ) ) ) ) ) ) ) |
| Col22a1 | ENSMUSG00000079022 | ENSMUST00000159993 | 411 | ) ) ) ) . . . . |
| Col24a1 | ENSMUSG00000028197 | ENSMUST00000029848 | 545 | . . . . . . ) ) |
| Col3a1 | ENSMUSG00000026043 | ENSMUST00000087883 | 158 | ) ) ) ) ) ) ) ) |
| Colca2 | ENSMUSG00000079559 | ENSMUST00000114427 | 194 | ) ) ) ) ) ) ) ) |
| Copb1 | ENSMUSG00000030754 | ENSMUST00000033012 | 125 | ) ) ) ) ) . . . |
| Cops3 | ENSMUSG00000019373 | ENSMUST00000019517 | 61 | ) ) ( ( ( ( ( ( |
| Coro1a | ENSMUSG00000030707 | ENSMUST00000032949 | 116 | ( ( ( ( ( ( ( ( |
| Coro6 | ENSMUSG00000020836 | ENSMUST00000108391 | 4 | . . . . . . . . |
| Cox11 | ENSMUSG00000020544 | ENSMUST00000020851 | 1 | . . . . . . ( ( |
| Cox14 | ENSMUSG00000023020 | ENSMUST00000023761 | 143 | ) ) ) ) ) ) ) ) |
| Cox6b2 | ENSMUSG00000051811 | ENSMUST00000182111 | 161 | . . . . . . . . |
| Cox7a2l | ENSMUSG00000024248 | ENSMUST00000025095 | 2 | ( ( ( ( . . . . |
| Cox7a2l | ENSMUSG00000024248 | ENSMUST00000167741 | 2 | ( ( ( ( . . . . |
| Cox7c | ENSMUSG00000017778 | ENSMUST00000131011 | 179 | . . . ) ) ) ) ) |
| Cox8c | ENSMUSG00000043319 | ENSMUST00000053611 | 33 | ) . . . . . . . |

| | | | | |
|---|---|---|---|---|
| Cpa6 | ENSMUSG00000042501 | ENSMUST00000035577 | 162 | ........ |
| Cpeb3 | ENSMUSG00000039652 | ENSMUST00000079754 | 79 | (((((((. |
| Cped1 | ENSMUSG00000062980 | ENSMUST00000115383 | 844 | ))))))). |
| Cpne4 | ENSMUSG00000032564 | ENSMUST00000057742 | 586 | ........ |
| Cpne6 | ENSMUSG00000022212 | ENSMUST00000074225 | 270 | ..)))).. |
| Cpne8 | ENSMUSG00000052560 | ENSMUST00000014777 | 26 | ........ |
| Cpne8 | ENSMUSG00000052560 | ENSMUST00000064391 | 70 | ........ |
| Cps1 | ENSMUSG00000025991 | ENSMUST00000027144 | 52 | ..)))).. |
| Cptp | ENSMUSG00000029073 | ENSMUST00000030950 | 287 | ((((.... |
| Cpxcr1 | ENSMUSG00000072995 | ENSMUST00000101269 | 134 | ((((..(. |
| Cr2 | ENSMUSG00000026616 | ENSMUST00000082321 | 38 | ........ |
| Crebl2 | ENSMUSG00000032652 | ENSMUST00000046303 | 188 | )))))))) |
| Crisp1 | ENSMUSG00000025431 | ENSMUST00000026498 | 9 | ((.(.... |
| Crisp3 | ENSMUSG00000025433 | ENSMUST00000026499 | 9 | ........ |
| Crot | ENSMUSG00000003623 | ENSMUST00000003720 | 256 | .))).... |
| Crtap | ENSMUSG00000032431 | ENSMUST00000084881 | 11 | (....... |
| Crx | ENSMUSG00000041578 | ENSMUST00000044434 | 141 | )))).... |
| Cryba4 | ENSMUSG00000066975 | ENSMUST00000112383 | 10 | ........ |
| Cryz | ENSMUSG00000028199 | ENSMUST00000029850 | 516 | ..)))))) |
| Cs | ENSMUSG00000005683 | ENSMUST00000005826 | 150 | ..(((... |
| Csf3r | ENSMUSG00000028859 | ENSMUST00000106162 | 279 | )))((((( |
| Csnk1a1 | ENSMUSG00000024576 | ENSMUST00000165123 | 423 | ((((.... |
| Csnk2b | ENSMUSG00000024387 | ENSMUST00000174024 | 15 | ...))))) |
| Csnk2b | ENSMUSG00000024387 | ENSMUST00000025246 | 128 | ........ |
| Csnka2ip | ENSMUSG00000068167 | ENSMUST00000089279 | 468 | ....)))) |
| Cstf3 | ENSMUSG00000027176 | ENSMUST00000028599 | 98 | ........ |
| Ctif | ENSMUSG00000052928 | ENSMUST00000165559 | 252 | (....... |
| Ctnna3 | ENSMUSG00000060843 | ENSMUST00000105440 | 64 | ........ |
| Ctnnd1 | ENSMUSG00000034101 | ENSMUST00000066177 | 128 | ))))))). |
| Ctnnd1 | ENSMUSG00000034101 | ENSMUST00000036811 | 192 | ........ |
| Ctnnd1 | ENSMUSG00000034101 | ENSMUST00000111697 | 358 | )))).... |
| Ctnnd1 | ENSMUSG00000034101 | ENSMUST00000111691 | 383 | )))).... |
| Ctnnd1 | ENSMUSG00000034101 | ENSMUST00000067232 | 412 | )))).... |
| Cts3 | ENSMUSG00000074870 | ENSMUST00000054702 | 145 | ))).)))) |
| Ctsg | ENSMUSG00000040314 | ENSMUST00000015583 | 54 | .....((( |
| Ctsh | ENSMUSG00000032359 | ENSMUST00000034915 | 225 | )))))))) |
| Ctu1 | ENSMUSG00000038888 | ENSMUST00000038332 | 84 | .))))... |
| Cul5 | ENSMUSG00000032030 | ENSMUST00000166367 | 178 | ..)))))) |
| Cul5 | ENSMUSG00000032030 | ENSMUST00000034529 | 182 | ..)))))) |
| Cux1 | ENSMUSG00000029705 | ENSMUST00000176778 | 601 | ........ |
| Cx3cr1 | ENSMUSG00000052336 | ENSMUST00000064165 | 73 | .))..))) |
| Cxxc4 | ENSMUSG00000044365 | ENSMUST00000181904 | 510 | ........ |
| Cyp11b2 | ENSMUSG00000022589 | ENSMUST00000167634 | 10 | .....))) |
| Cyp19a1 | ENSMUSG00000032274 | ENSMUST00000034811 | 11 | ........ |
| Cyp2j5 | ENSMUSG00000052520 | ENSMUST00000030299 | 108 | .((((((. |
| Cyp4f14 | ENSMUSG00000024292 | ENSMUST00000054174 | 153 | (((..... |
| Cyp51 | ENSMUSG00000001467 | ENSMUST00000001507 | 372 | ........ |
| D5Ertd577e | ENSMUSG00000070677 | ENSMUST00000094593 | 100 | ....)))) |
| D5Ertd579e | ENSMUSG00000029190 | ENSMUST00000031091 | 252 | ((((.... |
| D630039A03Rik | ENSMUSG00000052117 | ENSMUST00000063816 | 447 | ........ |
| Dach1 | ENSMUSG00000055639 | ENSMUST00000069334 | 158 | ........ |
| Dach1 | ENSMUSG00000055639 | ENSMUST00000071533 | 158 | ........ |
| Dag1 | ENSMUSG00000039952 | ENSMUST00000166905 | 209 | (....... |
| Dag1 | ENSMUSG00000039952 | ENSMUST00000191899 | 306 | )))).))) |
| Dag1 | ENSMUSG00000039952 | ENSMUST00000080435 | 318 | (....... |
| Dag1 | ENSMUSG00000039952 | ENSMUST00000171412 | 383 | ...((((( |
| Dcaf6 | ENSMUSG00000026571 | ENSMUST00000027856 | 209 | ........ |
| Dclre1a | ENSMUSG00000025077 | ENSMUST00000182276 | 559 | )))).))) |

| | | | | |
|---|---|---|---|---|
| Dclre1c | ENSMUSG00000026648 | ENSMUST00000115066 | 183 | ) . ( ( ( ( ( . |
| Dcx | ENSMUSG00000031285 | ENSMUST00000112851 | 94 | . . . ) . . ) ) |
| Dcx | ENSMUSG00000031285 | ENSMUST00000112856 | 109 | . . . ) . . ) ) |
| Dcx | ENSMUSG00000031285 | ENSMUST00000033642 | 201 | . . . ) . . ) ) |
| Ddb2 | ENSMUSG00000002109 | ENSMUST00000028696 | 79 | ( ( . . . . . . |
| Ddc | ENSMUSG00000020182 | ENSMUST00000109659 | 68 | ) ) ) ) ) ) ) . |
| Ddt | ENSMUSG00000001666 | ENSMUST00000001716 | 10 | . . . . . . . . |
| Ddx3y | ENSMUSG00000069045 | ENSMUST00000091190 | 94 | . . . . . . . ) |
| Ddx50 | ENSMUSG00000020076 | ENSMUST00000020270 | 34 | . . ( ( ( ( ( ( |
| Ddx54 | ENSMUSG00000029599 | ENSMUST00000031598 | 368 | . . . . . ) ) ) |
| Defb50 | ENSMUSG00000058568 | ENSMUST00000078879 | 3 | ( . . . . . . ( |
| Dennd4b | ENSMUSG00000042404 | ENSMUST00000098914 | 74 | ) ) . ) ) ) ) ) |
| Dhh | ENSMUSG00000023000 | ENSMUST00000023737 | 240 | ) . . . . . . . |
| Dhrs3 | ENSMUSG00000066026 | ENSMUST00000154208 | 347 | . . . . . . . . |
| Dhrs7c | ENSMUSG00000033044 | ENSMUST00000168612 | 25 | . . . . . ) ) ) |
| Dhx15 | ENSMUSG00000029169 | ENSMUST00000199321 | 104 | ) . ) ) . . ) ) |
| Dhx15 | ENSMUSG00000029169 | ENSMUST00000031061 | 129 | ) . . ( ( ( ( ( |
| Dkk1 | ENSMUSG00000024868 | ENSMUST00000025803 | 249 | . . . . . . . . |
| Dlc1 | ENSMUSG00000031523 | ENSMUST00000163663 | 373 | ) . . . . ) ) ) |
| Dlgap4 | ENSMUSG00000061689 | ENSMUST00000099145 | 528 | . . . . . . . . |
| Dlx1 | ENSMUSG00000041911 | ENSMUST00000037119 | 1797 | . . . ( ( ( . . |
| Dlx2 | ENSMUSG00000023391 | ENSMUST00000024159 | 340 | ) ) ) ) ) ) ) ) |
| Dmc1 | ENSMUSG00000022429 | ENSMUST00000023065 | 145 | ) ) ) ) ) . . . |
| Dmd | ENSMUSG00000045103 | ENSMUST00000114000 | 204 | ) ) ) ) ) ) ) ) |
| Dmrtc2 | ENSMUSG00000011349 | ENSMUST00000011493 | 64 | ( ( ( . . . . . |
| Dnaic2 | ENSMUSG00000034706 | ENSMUST00000069325 | 346 | ( ( . . . . ( ( |
| Dnaja1 | ENSMUSG00000028410 | ENSMUST00000030118 | 120 | ) ) ) . . . ) ) |
| Dnaja2 | ENSMUSG00000031701 | ENSMUST00000034138 | 49 | ( ( ( . . . . . |
| Dnajb1 | ENSMUSG00000005483 | ENSMUST00000212300 | 259 | ) ) . ) ) . . . |
| Dnajc16 | ENSMUSG00000040697 | ENSMUST00000038014 | 292 | ) . ( ( ( ( ( ( |
| Dnajc18 | ENSMUSG00000024350 | ENSMUST00000025208 | 22 | . . ( ( ( . ( ( |
| Dnajc6 | ENSMUSG00000028528 | ENSMUST00000106933 | 46 | . . ) ) . . ) ) |
| Dnajc9 | ENSMUSG00000021811 | ENSMUST00000022345 | 114 | ) ) ) . ) ) ) ) |
| Dnmt3b | ENSMUSG00000027478 | ENSMUST00000103150 | 392 | ) ) . . . . . . |
| Dnmt3b | ENSMUSG00000027478 | ENSMUST00000088976 | 392 | ) ) . . . . . . |
| Dnmt3b | ENSMUSG00000027478 | ENSMUST00000081628 | 393 | ) ) . . . . . . |
| Dnmt3b | ENSMUSG00000027478 | ENSMUST00000109774 | 393 | ) ) . . . . . . |
| Doc2a | ENSMUSG00000052301 | ENSMUST00000064110 | 552 | ( ( ( ( ( ( ( ( |
| Dock9 | ENSMUSG00000025558 | ENSMUST00000040700 | 19 | . . . . . . ( ( |
| Dock9 | ENSMUSG00000025558 | ENSMUST00000212181 | 19 | . . . . . . ( ( |
| Dppa2 | ENSMUSG00000072419 | ENSMUST00000097175 | 8 | . . . . . . . ) |
| Dppa4 | ENSMUSG00000058550 | ENSMUST00000050705 | 53 | . . ) ) ) ) ) ) |
| Dppa4 | ENSMUSG00000058550 | ENSMUST00000096045 | 53 | . . ) ) ) ) ) ) |
| Dpysl2 | ENSMUSG00000022048 | ENSMUST00000022629 | 254 | . . . . . . . ( |
| Dtwd2 | ENSMUSG00000024505 | ENSMUST00000025383 | 27 | ( ( ( ( . . . . |
| Dtwd2 | ENSMUSG00000024505 | ENSMUST00000163590 | 27 | ( ( ( ( . . . . |
| Duoxa1 | ENSMUSG00000027224 | ENSMUST00000110537 | 256 | . . . . . . . . |
| Dus3l | ENSMUSG00000007603 | ENSMUST00000007747 | 24 | . . . . . . ) . |
| Dus4l | ENSMUSG00000020648 | ENSMUST00000020977 | 141 | . . . . . . . ) |
| Dusp19 | ENSMUSG00000027001 | ENSMUST00000028384 | 297 | ) ) ) ) . ) ) ) |
| Dut | ENSMUSG00000027203 | ENSMUST00000051605 | 246 | ) ) ) . . . . . |
| Dync1h1 | ENSMUSG00000018707 | ENSMUST00000018851 | 134 | . . . . . . ) ) |
| Dync2h1 | ENSMUSG00000047193 | ENSMUST00000048417 | 68 | ) . . . ) ) . . |
| Dyrk1a | ENSMUSG00000022897 | ENSMUST00000023614 | 99 | ) ) ) ) . . ) ) |
| Dyrk1a | ENSMUSG00000022897 | ENSMUST00000119878 | 554 | ) ) ) ) ) ) ) ) |
| Dzip1l | ENSMUSG00000037784 | ENSMUST00000112886 | 122 | . . . . ) ) ) ) |
| Dzip1l | ENSMUSG00000037784 | ENSMUST00000078367 | 130 | ) ) . . . . . . |
| E330014E10Rik | ENSMUSG00000072813 | ENSMUST00000071182 | 100 | ( ( ( ( . . . . |

| | | | | |
|---|---|---|---|---|
| Ears2 | ENSMUSG00000030871 | ENSMUST00000033159 | 156 | . . . . ) ) ) ) |
| Ebag9 | ENSMUSG00000022339 | ENSMUST00000022964 | 329 | . ) ) ) ) ) ) ) |
| Ebf1 | ENSMUSG00000057098 | ENSMUST00000081265 | 805 | ) ) ) ) ) ) ) . |
| Ebf1 | ENSMUSG00000057098 | ENSMUST00000109268 | 805 | ) ) ) ) ) ) ) . |
| Ebf1 | ENSMUSG00000057098 | ENSMUST00000101326 | 805 | ) ) ) ) ) ) ) . |
| Eci2 | ENSMUSG00000021417 | ENSMUST00000171229 | 13 | ) ) . . . . . . |
| Eci3 | ENSMUSG00000021416 | ENSMUST00000021853 | 15 | . ( ( ( ( ( ( . |
| Eda2r | ENSMUSG00000034457 | ENSMUST00000037353 | 213 | ) ) ) ) ) ) ) ) |
| Edn1 | ENSMUSG00000021367 | ENSMUST00000021796 | 387 | . . . . . . . . |
| Eef1a1 | ENSMUSG00000037742 | ENSMUST00000042235 | 52 | ) ) ) ) ) ) ) ) |
| Eef1b2 | ENSMUSG00000025967 | ENSMUST00000129339 | 307 | . . . . . . ) ) |
| Eef1g | ENSMUSG00000071644 | ENSMUST00000052248 | 151 | . ( ( ( . . . . |
| Eef2 | ENSMUSG00000034994 | ENSMUST00000047864 | 14 | ( . . . . . . . |
| Efemp2 | ENSMUSG00000024909 | ENSMUST00000070118 | 242 | ) ) ( ( ( ( ( . |
| Egflam | ENSMUSG00000042961 | ENSMUST00000058593 | 166 | ( . . . . . . . |
| Egflam | ENSMUSG00000042961 | ENSMUST00000096494 | 213 | ( . . . . . . . |
| Eid1 | ENSMUSG00000091337 | ENSMUST00000164756 | 66 | . . . . . . . . |
| Eif1a | ENSMUSG00000057561 | ENSMUST00000078079 | 342 | ) ) ) ) ) ) ) ) |
| Eif2a | ENSMUSG00000027810 | ENSMUST00000029387 | 222 | ) ) ) ) ) ) ) ) |
| Eif2b4 | ENSMUSG00000029145 | ENSMUST00000114603 | 204 | ) ) ) ) ) ) . . |
| Eif2s3x | ENSMUSG00000035150 | ENSMUST00000050328 | 56 | . ) ) . . . . . |
| Eif2s3y | ENSMUSG00000069049 | ENSMUST00000091197 | 57 | . . . . . . . . |
| Eif3c | ENSMUSG00000030738 | ENSMUST00000032992 | 8 | . . . . . . . . |
| Eif3d | ENSMUSG00000016554 | ENSMUST00000100484 | 91 | . . . . . . . . |
| Eif3e | ENSMUSG00000022336 | ENSMUST00000022960 | 10 | ( . . . . . . . |
| Eif3h | ENSMUSG00000022312 | ENSMUST00000022925 | 43 | . . . . . . . . |
| Eif3i | ENSMUSG00000028798 | ENSMUST00000102593 | 3 | . . . . . . . . |
| Eif3k | ENSMUSG00000053565 | ENSMUST00000066070 | 5 | . . ( ( ( ( ( . |
| Eif3k | ENSMUSG00000053565 | ENSMUST00000208616 | 55 | ) ) . ) ) ) ) ) |
| Eif3m | ENSMUSG00000027170 | ENSMUST00000028592 | 36 | . . . . . ) ) ) |
| Eif4e2 | ENSMUSG00000026254 | ENSMUST00000113231 | 128 | . . ) ) ) ) ) ) |
| Elavl4 | ENSMUSG00000028546 | ENSMUST00000102722 | 346 | ) ) ) ) ) ) ) ) |
| Elavl4 | ENSMUSG00000028546 | ENSMUST00000106597 | 346 | ) ) ) ) ) ) ) ) |
| Elf3 | ENSMUSG00000003051 | ENSMUST00000185752 | 27 | . . . . . . . . |
| Elf3 | ENSMUSG00000003051 | ENSMUST00000003135 | 157 | ) ) ) ) ) ) ) ) |
| Elp5 | ENSMUSG00000018565 | ENSMUST00000108594 | 250 | ) ) ) ) ) . . . |
| Emc6 | ENSMUSG00000047260 | ENSMUST00000054952 | 49 | ( ( ( ( ( ( ( ( |
| Emg1 | ENSMUSG00000004268 | ENSMUST00000004379 | 38 | . . . . . ) ) ) |
| Eml3 | ENSMUSG00000071647 | ENSMUST00000096241 | 300 | . . . . . . . ) |
| Eno1 | ENSMUSG00000063524 | ENSMUST00000080926 | 355 | . ) ) ) ) ) ) ) |
| Enpp3 | ENSMUSG00000019989 | ENSMUST00000020169 | 61 | ( ( ( ( ( ( ( ( |
| Enpp4 | ENSMUSG00000023961 | ENSMUST00000024757 | 123 | ) ) ) ) ) ) ) . |
| Ensa | ENSMUSG00000038619 | ENSMUST00000037983 | 49 | . . . . . . . ) |
| Ensa | ENSMUSG00000038619 | ENSMUST00000058230 | 58 | . . . . . . . ) |
| Ep300 | ENSMUSG00000055024 | ENSMUST00000068387 | 1197 | . . ) ) ) ) ) ) |
| Epha5 | ENSMUSG00000029245 | ENSMUST00000053733 | 346 | ) ) ) ) ) ) ) . |
| Ephb3 | ENSMUSG00000005958 | ENSMUST00000006112 | 345 | ) . . . . . . . |
| Eps15 | ENSMUSG00000028552 | ENSMUST00000102729 | 112 | . . . . . . . . |
| Epsti1 | ENSMUSG00000022014 | ENSMUST00000022591 | 165 | . . . ) ) ) ) ) |
| Eras | ENSMUSG00000031160 | ENSMUST00000033500 | 154 | . . . . . . . . |
| Erbb2 | ENSMUSG00000062312 | ENSMUST00000058295 | 97 | ) ) ) ) ) ) ) ) |
| Erc1 | ENSMUSG00000030172 | ENSMUST00000183703 | 522 | ) ) ) ) ) . . . |
| Ercc4 | ENSMUSG00000022545 | ENSMUST00000023206 | 209 | ( ( ( ( ( ( ( ( |
| Ercc6l | ENSMUSG00000051220 | ENSMUST00000056904 | 85 | . . . . ) ) ) ) |
| Ergic3 | ENSMUSG00000005881 | ENSMUST00000006035 | 77 | ) ) . ) ) ) ) ) |
| Erich2 | ENSMUSG00000075302 | ENSMUST00000100041 | 249 | ) . . ) ) ) ) ) |
| Erich3 | ENSMUSG00000078161 | ENSMUST00000051862 | 173 | ) . ) ) ) ) . ( |
| Erp44 | ENSMUSG00000028343 | ENSMUST00000030028 | 52 | . . . . . . . . |

| | | | | |
|---|---|---|---|---|
| Esd | ENSMUSG00000021996 | ENSMUST00000022573 | 54 | ) . . . . ) ) ) |
| Esp23 | ENSMUSG00000096697 | ENSMUST00000178532 | 220 | ) ) . ) ) ) ) ) |
| Esrp2 | ENSMUSG00000084128 | ENSMUST00000115979 | 190 | . ) ) ) ) . ) ) |
| Etl4 | ENSMUSG00000036617 | ENSMUST00000045555 | 475 | . . . ( ( ( ( . |
| Etl4 | ENSMUSG00000036617 | ENSMUST00000114614 | 475 | . . . ( ( ( ( . |
| Ets1 | ENSMUSG00000032035 | ENSMUST00000050797 | 239 | . . . ( ( ( . . |
| Ets1 | ENSMUSG00000032035 | ENSMUST00000034534 | 266 | . . . ( ( ( . . |
| Etv6 | ENSMUSG00000030199 | ENSMUST00000081028 | 388 | . . ) ) ) ) ) ) |
| Evl | ENSMUSG00000021262 | ENSMUST00000109854 | 125 | ) . . . . ) ) ) |
| Eya1 | ENSMUSG00000025932 | ENSMUST00000027066 | 586 | ) . ) ) ) . . ) |
| Eya1 | ENSMUSG00000025932 | ENSMUST00000168081 | 621 | ) . ) ) ) . . ) |
| Fabp6 | ENSMUSG00000020405 | ENSMUST00000020672 | 15 | . . . ) ) . . ) |
| Faim2 | ENSMUSG00000023011 | ENSMUST00000023750 | 400 | . . . ( ( ( ( . |
| Fam102a | ENSMUSG00000039157 | ENSMUST00000048375 | 681 | . . . . . . . . |
| Fam167b | ENSMUSG00000050493 | ENSMUST00000052835 | 102 | . . . . . . . . |
| Fam193a | ENSMUSG00000037210 | ENSMUST00000180376 | 233 | ) ) . . . . . . |
| Fam19a1 | ENSMUSG00000059187 | ENSMUST00000122120 | 381 | ( ( ( . . . . . |
| Fam20a | ENSMUSG00000020614 | ENSMUST00000020938 | 494 | ) . ) ) ) ) ) ) |
| Fam221a | ENSMUSG00000047115 | ENSMUST00000121903 | 65 | ) ) ) . . . . . |
| Fam221a | ENSMUSG00000047115 | ENSMUST00000060561 | 149 | ) ) ) ) ) ) ) . |
| Fam46c | ENSMUSG00000044468 | ENSMUST00000061455 | 214 | ) ) . ) ) ) ) ) |
| Fam49a | ENSMUSG00000020589 | ENSMUST00000069005 | 118 | ) ) ) . ) ) ) ) |
| Fam49a | ENSMUSG00000020589 | ENSMUST00000069066 | 229 | ) ) ) ) ) ) . ) |
| Fam57b | ENSMUSG00000058966 | ENSMUST00000207020 | 337 | ) ) ) ) ) . . ) |
| Fam58b | ENSMUSG00000049489 | ENSMUST00000059468 | 39 | . . . . . . ) ) |
| Fam71b | ENSMUSG00000020401 | ENSMUST00000063166 | 37 | ( ( ( ( . . . . |
| Fam72a | ENSMUSG00000055184 | ENSMUST00000068613 | 777 | . . . . . . . . |
| Fap | ENSMUSG00000000392 | ENSMUST00000102732 | 144 | ) ) ) ) . . . . |
| Far2 | ENSMUSG00000030303 | ENSMUST00000111607 | 100 | ) ) ) ) ( ( ( ( |
| Far2 | ENSMUSG00000030303 | ENSMUST00000032443 | 257 | ( ( ( . ( ( ( ( |
| Farp1 | ENSMUSG00000025555 | ENSMUST00000026635 | 374 | . . . . . . . . |
| Fasl | ENSMUSG00000000817 | ENSMUST00000000834 | 188 | . ) ) ) ) ) . ) |
| Fastkd5 | ENSMUSG00000079043 | ENSMUST00000110262 | 139 | ) ) . . ) ) ) ) |
| Fau | ENSMUSG00000038274 | ENSMUST00000178310 | 62 | ( ( . ( . . . . |
| Fau | ENSMUSG00000038274 | ENSMUST00000043074 | 62 | . . . . . . . . |
| Fau | ENSMUSG00000038274 | ENSMUST00000179142 | 263 | . . . . . . . . |
| Faxc | ENSMUSG00000028246 | ENSMUST00000029908 | 174 | ( ( ( . . . ( ( |
| Fbf1 | ENSMUSG00000020776 | ENSMUST00000103031 | 581 | . . ( ( ( ( ( ( |
| Fbl | ENSMUSG00000046865 | ENSMUST00000042405 | 18 | ( ( ( . ( ( ( ( |
| Fbxl15 | ENSMUSG00000025226 | ENSMUST00000026256 | 204 | ) ) ) ) ( ( ( ( |
| Fbxl8 | ENSMUSG00000033313 | ENSMUST00000036221 | 373 | . . . . . . . . |
| Fbxo33 | ENSMUSG00000035329 | ENSMUST00000043204 | 229 | . . . . . . . . |
| Fbxo39 | ENSMUSG00000070388 | ENSMUST00000108504 | 108 | ) ) ) ) ( ( ( ( |
| Fbxo5 | ENSMUSG00000019773 | ENSMUST00000019907 | 209 | . . . . . . . . |
| Fbxo7 | ENSMUSG00000001786 | ENSMUST00000130320 | 8 | ( ( ( ( . . . . |
| Fcgrt | ENSMUSG00000003420 | ENSMUST00000003512 | 387 | . . ) ) ) . . . ) |
| Fcrl1 | ENSMUSG00000059994 | ENSMUST00000072480 | 52 | ( ( ( ( . . ( ( |
| Fcrl1 | ENSMUSG00000059994 | ENSMUST00000163661 | 54 | ( ( ( ( . . ( ( |
| Ffar1 | ENSMUSG00000044453 | ENSMUST00000052700 | 23 | ( ( ( ( . ( ( ( |
| Fgf14 | ENSMUSG00000025551 | ENSMUST00000026631 | 308 | ) . . . . . . . |
| Fgf23 | ENSMUSG00000000182 | ENSMUST00000000186 | 34 | ( ( ( ( ( ( . . |
| Fhit | ENSMUSG00000060579 | ENSMUST00000161302 | 51 | . . . . . . . . |
| Fibin | ENSMUSG00000074971 | ENSMUST00000099626 | 342 | . ( ( ( ( ( ( ( |
| Fkbp1a | ENSMUSG00000032966 | ENSMUST00000044011 | 181 | ) ) ) ) ) ) ) . |
| Flnb | ENSMUSG00000025278 | ENSMUST00000052678 | 119 | ) ) . . . ) ) ) |
| Flrt2 | ENSMUSG00000047414 | ENSMUST00000057324 | 731 | . . . . . . . ) |
| Fmo6 | ENSMUSG00000095576 | ENSMUST00000178465 | 72 | . . . ) ) ) ) ) |
| Fmo9 | ENSMUSG00000026560 | ENSMUST00000148677 | 63 | . . . ( ( ( ( ( |

| | | | | |
|---|---|---|---|---|
| Fmod | ENSMUSG00000041559 | ENSMUST00000048183 | 249 | ) ) . . . . . . |
| Fnbp1 | ENSMUSG00000075415 | ENSMUST00000113552 | 116 | ) ) ) ) ) ) ) ) |
| Fnbp1 | ENSMUSG00000075415 | ENSMUST00000113559 | 120 | . . . ) ) ) ) ) |
| Fnbp1 | ENSMUSG00000075415 | ENSMUST00000113564 | 134 | ) ) ) ) ) ) ) ) |
| Fnbp1 | ENSMUSG00000075415 | ENSMUST00000113562 | 173 | ) ) ) ) ) ) ) . |
| Fnbp1 | ENSMUSG00000075415 | ENSMUST00000113560 | 185 | ) ) ) ) ) ) ) ) |
| Fnbp4 | ENSMUSG00000008200 | ENSMUST00000013759 | 14 | ( ( ( . . . ( ( |
| Foxc2 | ENSMUSG00000046714 | ENSMUST00000054691 | 357 | ) ) ) . . . . ( |
| Foxd4 | ENSMUSG00000051490 | ENSMUST00000058600 | 455 | . . . . . . . . |
| Foxh1 | ENSMUSG00000033837 | ENSMUST00000037824 | 28 | . . . . . ( ( ( |
| Fpr1 | ENSMUSG00000045551 | ENSMUST00000061516 | 3 | ( ( . . ( ( ( ( |
| Frmd4a | ENSMUSG00000026657 | ENSMUST00000091497 | 121 | ) ) ) ) ) ) ) ) |
| Fscn2 | ENSMUSG00000025380 | ENSMUST00000026445 | 134 | ) ) ) . ) ) ) ) |
| Fsip1 | ENSMUSG00000027344 | ENSMUST00000028821 | 119 | ( ( . ( ( ( . . |
| Fstl4 | ENSMUSG00000036264 | ENSMUST00000036796 | 194 | . . . . . . . . |
| Ftmt | ENSMUSG00000024510 | ENSMUST00000025388 | 59 | ( ( ( ( ( . . . |
| Fubp1 | ENSMUSG00000028034 | ENSMUST00000166984 | 97 | ) ) ) ) . ) ) ) |
| Fubp3 | ENSMUSG00000026843 | ENSMUST00000113482 | 137 | ) ) ) ) ) ) ) ) |
| Fut7 | ENSMUSG00000036587 | ENSMUST00000041654 | 434 | . . . . . . . . |
| Fut7 | ENSMUSG00000036587 | ENSMUST00000114278 | 454 | . . . ) ) ) ) . |
| Fxr1 | ENSMUSG00000027680 | ENSMUST00000001620 | 140 | . . . ) ) ) . ) |
| Fzd7 | ENSMUSG00000041075 | ENSMUST00000114246 | 493 | ) ) ) . ) ) ) ) |
| G0s2 | ENSMUSG00000009633 | ENSMUST00000009777 | 208 | ) ) ) . . . . . |
| Gabpb2 | ENSMUSG00000038766 | ENSMUST00000136139 | 455 | ) ) ) ) ) ) . ) |
| Gabra5 | ENSMUSG00000055078 | ENSMUST00000068456 | 301 | ) ) ) ) ) ) ) ) |
| Gabrq | ENSMUSG00000031344 | ENSMUST00000114553 | 235 | ) ) ) ) . . . ) |
| Gabrr3 | ENSMUSG00000074991 | ENSMUST00000114341 | 31 | . . . ( ( ( ( ( |
| Galc | ENSMUSG00000021003 | ENSMUST00000021390 | 67 | . . . . . . . . |
| Galk2 | ENSMUSG00000027207 | ENSMUST00000028636 | 178 | . . . . ) ) ) ) |
| Galr3 | ENSMUSG00000114755 | ENSMUST00000058004 | 24 | ) ) . . ( ( ( ( |
| Gapvd1 | ENSMUSG00000026867 | ENSMUST00000102800 | 564 | ) ) ) . . . . . |
| Gas7 | ENSMUSG00000033066 | ENSMUST00000108680 | 70 | ) ) . . . . . . |
| Gas7 | ENSMUSG00000033066 | ENSMUST00000108682 | 235 | . . . . . . ) ) |
| Gata2 | ENSMUSG00000015053 | ENSMUST00000015197 | 525 | ) ) ) ) ) ) ) ) |
| Gcm2 | ENSMUSG00000021362 | ENSMUST00000021791 | 185 | ( ( ( ( ( . ( ( |
| Gdap1l1 | ENSMUSG00000017943 | ENSMUST00000109420 | 45 | ( . . . . . . . |
| Gdf5 | ENSMUSG00000038259 | ENSMUST00000040162 | 225 | . . . ) ) ) . . |
| Gdi2 | ENSMUSG00000021218 | ENSMUST00000223396 | 223 | ) ) . . . . . . |
| Gdpd5 | ENSMUSG00000035314 | ENSMUST00000037528 | 953 | ( ( ( ( ( ( ( ( |
| Ggt5 | ENSMUSG00000006344 | ENSMUST00000072217 | 272 | ) ) . . ) ) ) ) |
| Gid4 | ENSMUSG00000018415 | ENSMUST00000070681 | 352 | . . . . . . ) ) |
| Gja1 | ENSMUSG00000050953 | ENSMUST00000068581 | 114 | . . . . ( ( ( ( |
| Gjc3 | ENSMUSG00000056966 | ENSMUST00000077119 | 183 | ) ) ) ) ) ) . . |
| Glra2 | ENSMUSG00000018589 | ENSMUST00000058787 | 471 | ) ) ) . . . ) ) |
| Glrx2 | ENSMUSG00000018196 | ENSMUST00000129653 | 50 | . . ( ( ( ( ( ( |
| Glrx2 | ENSMUSG00000018196 | ENSMUST00000145969 | 338 | ( . . . . ( ( ( |
| Gls | ENSMUSG00000026103 | ENSMUST00000114513 | 131 | . . . . ) ) ) ) |
| Gls | ENSMUSG00000026103 | ENSMUST00000114510 | 140 | . . . . ) ) ) ) |
| Glt8d2 | ENSMUSG00000020251 | ENSMUST00000020485 | 163 | ( . . . . . . . |
| Gm10334 | ENSMUSG00000071517 | ENSMUST00000095999 | 69 | . . . . . . . . |
| Gm10681 | ENSMUSG00000095388 | ENSMUST00000058728 | 31 | . . . . . . . ( |
| Gm11213 | ENSMUSG00000084782 | ENSMUST00000210529 | 8 | . . . . . . . . |
| Gm11541 | ENSMUSG00000056008 | ENSMUST00000069852 | 196 | . . ) ) ) . . . |
| Gm11992 | ENSMUSG00000040978 | ENSMUST00000043285 | 515 | ) ) ) ) . . . . |
| Gm13040 | ENSMUSG00000070616 | ENSMUST00000169056 | 2 | . ( ( ( ( ( . . |
| Gm13306 | ENSMUSG00000073877 | ENSMUST00000108018 | 99 | . . . . . . . . |
| Gm136 | ENSMUSG00000071015 | ENSMUST00000095129 | 191 | . . . . ( ( ( ( |
| Gm1527 | ENSMUSG00000074655 | ENSMUST00000099170 | 137 | ) ) ) ) ) ) ) ) |

| | | | | |
|---|---|---|---|---|
| Gm15319 | ENSMUSG00000074449 | ENSMUST00000084046 | 83 | (((((((( |
| Gm16513 | ENSMUSG00000095996 | ENSMUST00000201663 | 96 | )..))))) |
| Gm17727 | ENSMUSG00000090738 | ENSMUST00000171898 | 185 | .)))))). |
| Gm2016 | ENSMUSG00000072905 | ENSMUST00000110147 | 480 | ......)) |
| Gm21119 | ENSMUSG00000095294 | ENSMUST00000178438 | 83 | (((((((( |
| Gm21319 | ENSMUSG00000095724 | ENSMUST00000164517 | 489 | ))..)))) |
| Gm3139 | ENSMUSG00000095074 | ENSMUST00000202642 | 136 | ((((.... |
| Gm3183 | ENSMUSG00000095954 | ENSMUST00000201703 | 136 | ((((.... |
| Gm3286 | ENSMUSG00000079423 | ENSMUST00000201629 | 82 | )..))))) |
| Gm3286 | ENSMUSG00000079423 | ENSMUST00000200932 | 125 |))).)))) |
| Gm4027 | ENSMUSG00000092019 | ENSMUST00000164838 | 483 | ........ |
| Gm4297 | ENSMUSG00000081218 | ENSMUST00000119285 | 1563 | )))))))) |
| Gm43786 | ENSMUSG00000107252 | ENSMUST00000201722 | 156 | .....((( |
| Gm44805 | ENSMUSG00000109350 | ENSMUST00000150569 | 72 | ))..)))) |
| Gm45837 | ENSMUSG00000030653 | ENSMUST00000084894 | 12 | (.....(( |
| Gm5615 | ENSMUSG00000074448 | ENSMUST00000098900 | 16 | ..((.... |
| Gm5662 | ENSMUSG00000079029 | ENSMUST00000101168 | 480 | ........ |
| Gm5934 | ENSMUSG00000084063 | ENSMUST00000120734 | 1563 | )......( |
| Gm648 | ENSMUSG00000064016 | ENSMUST00000081133 | 148 | ((((.... |
| Gm7694 | ENSMUSG00000102752 | ENSMUST00000179801 | 40 | .......) |
| Gm8300 | ENSMUSG00000079034 | ENSMUST00000110152 | 492 | ...))))) |
| Gm8369 | ENSMUSG00000058470 | ENSMUST00000079855 | 82 | .))))).. |
| Gm9573 | ENSMUSG00000090588 | ENSMUST00000164502 | 115 | .....))) |
| Gmfg | ENSMUSG00000060791 | ENSMUST00000078845 | 26 | ..))))). |
| Gmip | ENSMUSG00000036246 | ENSMUST00000036074 | 65 | (....... |
| Gng4 | ENSMUSG00000021303 | ENSMUST00000021734 | 346 | ))...... |
| Gpat2 | ENSMUSG00000046338 | ENSMUST00000062211 | 24 | ((...... |
| Gpat3 | ENSMUSG00000029314 | ENSMUST00000031255 | 271 | .....))) |
| Gpc2 | ENSMUSG00000029510 | ENSMUST00000161827 | 140 | ))(((((( |
| Gpi1 | ENSMUSG00000036427 | ENSMUST00000038027 | 76 | ......(( |
| Gpn2 | ENSMUSG00000028848 | ENSMUST00000030661 | 17 | )))...(( |
| Gpr150 | ENSMUSG00000045509 | ENSMUST00000056130 | 152 | .....((( |
| Gpr165 | ENSMUSG00000031210 | ENSMUST00000033554 | 433 |))))).)) |
| Gpr17 | ENSMUSG00000052229 | ENSMUST00000064016 | 151 | ........ |
| Gpr33 | ENSMUSG00000035148 | ENSMUST00000040161 | 56 | )))))))) |
| Gpr39 | ENSMUSG00000026343 | ENSMUST00000027581 | 332 | )))).))) |
| Gpr61 | ENSMUSG00000046793 | ENSMUST00000062028 | 649 | )))))... |
| Gpr65 | ENSMUSG00000021886 | ENSMUST00000075072 | 462 | (.((((.. |
| Gpr75 | ENSMUSG00000043999 | ENSMUST00000109430 | 234 | ((((((.. |
| Gpr85 | ENSMUSG00000048216 | ENSMUST00000060442 | 1146 | )))))).) |
| Gpx2 | ENSMUSG00000042808 | ENSMUST00000082431 | 111 | ((...((( |
| Gramd2 | ENSMUSG00000074259 | ENSMUST00000098661 | 204 | .))))))) |
| Grap2 | ENSMUSG00000042351 | ENSMUST00000043149 | 208 | .....))) |
| Grb2 | ENSMUSG00000059923 | ENSMUST00000106497 | 258 | )))))))) |
| Grb2 | ENSMUSG00000059923 | ENSMUST00000021090 | 284 | ))))).)) |
| Gria1 | ENSMUSG00000020524 | ENSMUST00000094179 | 52 | )))))))) |
| Gria1 | ENSMUSG00000020524 | ENSMUST00000036315 | 198 | )))))))) |
| Gria3 | ENSMUSG00000001986 | ENSMUST00000076349 | 171 | )))))))) |
| Grk4 | ENSMUSG00000052783 | ENSMUST00000001112 | 482 | ..)))..) |
| Grk4 | ENSMUSG00000052783 | ENSMUST00000074651 | 482 | ..)))..) |
| Grm2 | ENSMUSG00000023192 | ENSMUST00000023959 | 110 | )).))))) |
| Grm5 | ENSMUSG00000049583 | ENSMUST00000155358 | 91 | ....)))) |
| Grm5 | ENSMUSG00000049583 | ENSMUST00000125009 | 481 | ........ |
| Gse1 | ENSMUSG00000031822 | ENSMUST00000034279 | 607 | .))))))) |
| Gsn | ENSMUSG00000026879 | ENSMUST00000201185 | 106 | .......( |
| Gsta4 | ENSMUSG00000032348 | ENSMUST00000034903 | 77 | ...((((( |
| Gtf2a1 | ENSMUSG00000020962 | ENSMUST00000021345 | 402 | ..))).)) |
| Gtf2f1 | ENSMUSG00000002658 | ENSMUST00000002733 | 95 | .))))))) |

| | | | | |
|---|---|---|---|---|
| Gtpbp3 | ENSMUSG00000007610 | ENSMUST00000007754 | 622 | ))))..). |
| Gtsf1l | ENSMUSG00000070708 | ENSMUST00000094653 | 229 | ))...((( |
| Guca1b | ENSMUSG00000023979 | ENSMUST00000024774 | 78 | )))))))) |
| Gucy1b3 | ENSMUSG00000028005 | ENSMUST00000029635 | 54 | .....))) |
| Gucy1b3 | ENSMUSG00000028005 | ENSMUST00000193597 | 58 | ...((((( |
| Gypc | ENSMUSG00000090523 | ENSMUST00000174000 | 88 | )))..... |
| Gypc | ENSMUSG00000090523 | ENSMUST00000174459 | 142 | )))..... |
| Gzmb | ENSMUSG00000015437 | ENSMUST00000015581 | 33 | ...((((( |
| Gzmc | ENSMUSG00000079186 | ENSMUST00000015585 | 14 | ...((((( |
| Gzmd | ENSMUSG00000059256 | ENSMUST00000082093 | 17 | ........ |
| Gzme | ENSMUSG00000022156 | ENSMUST00000089549 | 10 | .((((... |
| Gzmf | ENSMUSG00000015441 | ENSMUST00000022757 | 0 | ...((((( |
| Gzmm | ENSMUSG00000054206 | ENSMUST00000020549 | 356 | ......)) |
| H1fx | ENSMUSG00000044927 | ENSMUST00000056403 | 220 | ........ |
| H2afz | ENSMUSG00000037894 | ENSMUST00000174561 | 83 | ))...))) |
| H2afz | ENSMUSG00000037894 | ENSMUST00000041045 | 169 | )))))))) |
| Has2 | ENSMUSG00000022367 | ENSMUST00000050544 | 477 | ....)))) |
| Haus3 | ENSMUSG00000079555 | ENSMUST00000060049 | 196 | (((((... |
| Hcfc1 | ENSMUSG00000031386 | ENSMUST00000033761 | 324 | .))))))) |
| Hdac5 | ENSMUSG00000008855 | ENSMUST00000008999 | 734 | ....)))) |
| Hdac7 | ENSMUSG00000022475 | ENSMUST00000121514 | 131 | ......)) |
| Hdx | ENSMUSG00000034551 | ENSMUST00000113422 | 85 | )))..... |
| Hdx | ENSMUSG00000034551 | ENSMUST00000038472 | 85 | ((((...( |
| Hes2 | ENSMUSG00000028940 | ENSMUST00000030782 | 93 | )).))))) |
| Hexb | ENSMUSG00000021665 | ENSMUST00000022169 | 29 | (((....( |
| Hexim1 | ENSMUSG00000048878 | ENSMUST00000053063 | 637 | .)))))). |
| Hif1a | ENSMUSG00000021109 | ENSMUST00000021530 | 292 | (((((... |
| Hist1h1a | ENSMUSG00000049539 | ENSMUST00000055770 | 4 | ........ |
| Hist1h2ad | ENSMUSG00000071478 | ENSMUST00000090776 | 7 | (((((((( |
| Hist1h2bh | ENSMUSG00000114456 | ENSMUST00000224359 | 5 | ........ |
| Hist1h3c | ENSMUSG00000069310 | ENSMUST00000091752 | 66 | ...)))).. |
| Hist1h3d | ENSMUSG00000099583 | ENSMUST00000105105 | 26 | ........ |
| Hist3h2a | ENSMUSG00000078851 | ENSMUST00000108817 | 13 | ..((.... |
| Hmgcr | ENSMUSG00000021670 | ENSMUST00000022176 | 55 | ))))(((( |
| Hmgcs1 | ENSMUSG00000093930 | ENSMUST00000224188 | 23 | ((((((.( |
| Hmgcs1 | ENSMUSG00000093930 | ENSMUST00000179869 | 41 | ......)) |
| Hmgn5 | ENSMUSG00000031245 | ENSMUST00000033597 | 94 | ))...))) |
| Hnf1b | ENSMUSG00000020679 | ENSMUST00000108113 | 96 | (((((((. |
| Hnf1b | ENSMUSG00000020679 | ENSMUST00000108114 | 156 | )).((((( |
| Hnf1b | ENSMUSG00000020679 | ENSMUST00000021016 | 175 | ...((((( |
| Hnrnpa1 | ENSMUSG00000046434 | ENSMUST00000036004 | 19 | ..(((((. |
| Hnrnpa1 | ENSMUSG00000046434 | ENSMUST00000087351 | 19 | ..(((((. |
| Hnrnpc | ENSMUSG00000060373 | ENSMUST00000227458 | 102 | )))))((( |
| Hnrnpc | ENSMUSG00000060373 | ENSMUST00000227242 | 104 | )))))((( |
| Hnrnpc | ENSMUSG00000060373 | ENSMUST00000228748 | 106 | )))))((( |
| Hnrnpc | ENSMUSG00000060373 | ENSMUST00000111610 | 127 | )))))... |
| Hnrnpul1 | ENSMUSG00000040725 | ENSMUST00000206832 | 26 | ....(((( |
| Homez | ENSMUSG00000057156 | ENSMUST00000142283 | 158 | )))))..) |
| Hopx | ENSMUSG00000059325 | ENSMUST00000113453 | 99 | ..)))))) |
| Hoxa2 | ENSMUSG00000014704 | ENSMUST00000014848 | 108 | (((((((. |
| Hoxb8 | ENSMUSG00000056648 | ENSMUST00000052650 | 975 | ........ |
| Hoxb9 | ENSMUSG00000020875 | ENSMUST00000000010 | 48 | ........ |
| Hscb | ENSMUSG00000043510 | ENSMUST00000056937 | 2 | ........ |
| Hsd17b12 | ENSMUSG00000027195 | ENSMUST00000028619 | 52 | ........ |
| Hsd3b4 | ENSMUSG00000095143 | ENSMUST00000196861 | 31 | .......( |
| Hspa12a | ENSMUSG00000025092 | ENSMUST00000066285 | 67 | ))))...) |
| Hspa4 | ENSMUSG00000020361 | ENSMUST00000020630 | 206 | (..(...( |
| Htr1a | ENSMUSG00000021721 | ENSMUST00000022235 | 579 | )))))))) |

| | | | | |
|---|---|---|---|---|
| Htr1b | ENSMUSG00000049511 | ENSMUST00000051005 | 2 | ( ( ( ( . . . ( |
| Htr2c | ENSMUSG00000041380 | ENSMUST00000036303 | 603 | ) . ) ) . . . ( |
| Hyal4 | ENSMUSG00000029680 | ENSMUST00000031691 | 173 | ) ) ) ) ) . . . |
| Hydin | ENSMUSG00000059854 | ENSMUST00000043141 | 53 | ) ) ) . ( ( ( ( |
| Id1 | ENSMUSG00000042745 | ENSMUST00000038368 | 32 | . . . . . . . . |
| Id3 | ENSMUSG00000007872 | ENSMUST00000008016 | 349 | ) ) ) ) . . . . |
| Idi1 | ENSMUSG00000058258 | ENSMUST00000169314 | 259 | . . ( ( ( . . ( |
| Ifitm1 | ENSMUSG00000025491 | ENSMUST00000026564 | 175 | ) ) ) ) ) ) ) ) |
| Ift20 | ENSMUSG00000001105 | ENSMUST00000128788 | 352 | ) . ) ) ) ) ) ) |
| Ift27 | ENSMUSG00000016637 | ENSMUST00000016781 | 257 | ) ) . ) ) ) ) ) |
| Ift46 | ENSMUSG00000002031 | ENSMUST00000002099 | 366 | ) ) ) ) ) ) ) ) |
| Ift88 | ENSMUSG00000040040 | ENSMUST00000122063 | 77 | ) ) ) ) . . . . |
| Igbp1 | ENSMUSG00000031221 | ENSMUST00000033570 | 62 | . ) ) . . . . . |
| Igf1 | ENSMUSG00000020053 | ENSMUST00000105300 | 31 | . . . . . . . . |
| Igf1 | ENSMUSG00000020053 | ENSMUST00000095360 | 221 | ) . . . . . . . |
| Igf2 | ENSMUSG00000048583 | ENSMUST00000000033 | 71 | . . . . . . . . |
| Igf2bp3 | ENSMUSG00000029814 | ENSMUST00000031838 | 475 | ) ) . . . ) ) ) |
| Igfbp2 | ENSMUSG00000039323 | ENSMUST00000120564 | 130 | ) ) ) ) ) ) ) . |
| Igfbp5 | ENSMUSG00000026185 | ENSMUST00000027377 | 682 | . . . . . ( ( ( |
| Igsf1 | ENSMUSG00000031111 | ENSMUST00000114893 | 5 | . . . ( ( ( ( ( |
| Igsf3 | ENSMUSG00000042035 | ENSMUST00000043983 | 737 | ) ) ) ) ) ) ) ) |
| Igsf6 | ENSMUSG00000035004 | ENSMUST00000047194 | 19 | ( ( ( ( ( ( ( . |
| Ikbkap | ENSMUSG00000028431 | ENSMUST00000030140 | 117 | ( ( ( . . . . . |
| Ikzf2 | ENSMUSG00000025997 | ENSMUST00000027146 | 121 | . . . . . . . . |
| Il1f5 | ENSMUSG00000026983 | ENSMUST00000028360 | 253 | ) ) ) ) . . . ( |
| Imp3 | ENSMUSG00000032288 | ENSMUST00000034827 | 8 | . . . . . . . . |
| Impact | ENSMUSG00000024423 | ENSMUST00000025290 | 35 | ( ( ( ( . . . . |
| Inca1 | ENSMUSG00000057054 | ENSMUST00000108542 | 47 | ) ) ) ) . . . . |
| Inca1 | ENSMUSG00000057054 | ENSMUST00000108543 | 355 | ) ) ) ) . ) ) . |
| Ino80d | ENSMUSG00000040865 | ENSMUST00000165066 | 213 | ( ( ( . ( ( ( ( |
| Inpp5e | ENSMUSG00000026925 | ENSMUST00000114090 | 294 | . . ) ) ) ) ) ) |
| Inpp5e | ENSMUSG00000026925 | ENSMUST00000145701 | 559 | ) ) ) ) ) ) ) ) |
| Insig2 | ENSMUSG00000003721 | ENSMUST00000159085 | 162 | . . ) ) ) ) ) ) |
| Insig2 | ENSMUSG00000003721 | ENSMUST00000071064 | 169 | . . . . . . . . |
| Insig2 | ENSMUSG00000003721 | ENSMUST00000003818 | 405 | . . . . . . . . |
| Insm1 | ENSMUSG00000068154 | ENSMUST00000089257 | 326 | ) ) . . . . . . |
| Insrr | ENSMUSG00000005640 | ENSMUST00000029711 | 426 | ( ( ( ( . . . . |
| Ints1 | ENSMUSG00000029547 | ENSMUST00000200393 | 102 | ) ) . . . . . ) |
| Ip6k2 | ENSMUSG00000032599 | ENSMUST00000085018 | 170 | . . ) ) ) ) ) ) |
| Ip6k2 | ENSMUSG00000106672 | ENSMUST00000201336 | 170 | . . ) ) ) ) ) ) |
| Ipcef1 | ENSMUSG00000064065 | ENSMUST00000105617 | 71 | . . . ( ( ( ( ( |
| Ippk | ENSMUSG00000021385 | ENSMUST00000021817 | 87 | . . . . . . . . |
| Iqcf5 | ENSMUSG00000066382 | ENSMUST00000085113 | 7 | ( . ( ( ( ( ( . |
| Iqsec1 | ENSMUSG00000034312 | ENSMUST00000101151 | 236 | . . . ) ) ) ) ) |
| Ireb2 | ENSMUSG00000032293 | ENSMUST00000034843 | 39 | . . . . . . . . |
| Irf2bpl | ENSMUSG00000034168 | ENSMUST00000038422 | 813 | ( ( . . . . . . |
| Ist1 | ENSMUSG00000031729 | ENSMUST00000034164 | 45 | . . . ) ) ) ) ) |
| Itgal | ENSMUSG00000030830 | ENSMUST00000106306 | 92 | ) ) ) ) ) ) ) ) |
| Itgal | ENSMUSG00000030830 | ENSMUST00000117762 | 123 | ) ) ) ) ) ) ) ) |
| Itgam | ENSMUSG00000030786 | ENSMUST00000106242 | 45 | ) ) ) . . . . . |
| Itgam | ENSMUSG00000030786 | ENSMUST00000064821 | 74 | ) ) ) ) ) . ) ) |
| Itgb3 | ENSMUSG00000020689 | ENSMUST00000021028 | 4 | . . . . . . . ( |
| Itm2a | ENSMUSG00000031239 | ENSMUST00000033591 | 92 | ) ) ) ) ) ) ) ) |
| Jak2 | ENSMUSG00000024789 | ENSMUST00000065796 | 355 | . . . . ( ( ( . |
| Jph3 | ENSMUSG00000025318 | ENSMUST00000026357 | 18 | . . . . . . . . |
| Kap | ENSMUSG00000032758 | ENSMUST00000048032 | 97 | . . . . . . . . |
| Kars | ENSMUSG00000031948 | ENSMUST00000034426 | 26 | ( ( ( ( ( ( . ( |
| Kcna4 | ENSMUSG00000042604 | ENSMUST00000037012 | 1159 | . . . . . ) ) ) |

| | | | | |
|---|---|---|---|---|
| Kcnf1 | ENSMUSG00000051726 | ENSMUST00000170580 | 606 | ( . . . . ( ( ( |
| Kcng1 | ENSMUSG00000074575 | ENSMUST00000099069 | 53 | . ) ) ) ) ) . . |
| Kcnh5 | ENSMUSG00000034402 | ENSMUST00000042299 | 743 | ) . . ) ) ) . ) |
| Kcnh8 | ENSMUSG00000035580 | ENSMUST00000039366 | 16 | . . . . ( ( ( ( |
| Kcnip1 | ENSMUSG00000053519 | ENSMUST00000065970 | 273 | ) ) . . . . . . |
| Kcnip1 | ENSMUSG00000053519 | ENSMUST00000109340 | 331 | ) ) . . . . . . |
| Kcnip3 | ENSMUSG00000079056 | ENSMUST00000103215 | 100 | ) ) ) ) . ) ) . |
| Kcnip4 | ENSMUSG00000029088 | ENSMUST00000175660 | 261 | . . . . . . . . |
| Kcnj11 | ENSMUSG00000096146 | ENSMUST00000211674 | 316 | ) ) ) ) . ) ) ) |
| Kcnk10 | ENSMUSG00000033854 | ENSMUST00000110113 | 1 | ( ( . . . . . ( |
| Kcnk10 | ENSMUSG00000033854 | ENSMUST00000221240 | 386 | ) . . . . . . . |
| Kcnk10 | ENSMUSG00000033854 | ENSMUST00000221305 | 468 | . . . . . . ) ) |
| Kcnk4 | ENSMUSG00000024957 | ENSMUST00000025908 | 198 | ) ( ( ( ( ( ( ( |
| Kcns3 | ENSMUSG00000043673 | ENSMUST00000164495 | 338 | . . . . . . ) ) |
| Kcns3 | ENSMUSG00000043673 | ENSMUST00000055673 | 401 | ) ) ) ) . ) ) ) |
| Kctd11 | ENSMUSG00000046731 | ENSMUST00000050555 | 1107 | ( ( ( ( . . . . |
| Kctd20 | ENSMUSG00000005936 | ENSMUST00000168507 | 130 | ) ) . . . . . ) |
| Kctd9 | ENSMUSG00000034327 | ENSMUST00000152243 | 615 | ) ) ) ) ) ) ) ( |
| Kdm2b | ENSMUSG00000029475 | ENSMUST00000046073 | 129 | ( ( ( ( . . ( ( |
| Kdm4c | ENSMUSG00000028397 | ENSMUST00000030102 | 107 | ( ( ( . . . . . |
| Kdm4d | ENSMUSG00000053914 | ENSMUST00000058796 | 630 | ) ) ) . ) ) ) ) |
| Kel | ENSMUSG00000029866 | ENSMUST00000031899 | 196 | ) ) ) . ( ( ( ( |
| Kera | ENSMUSG00000019932 | ENSMUST00000105286 | 411 | . ) ) ) ) . . ) |
| Kif26a | ENSMUSG00000021294 | ENSMUST00000128402 | 137 | . . . . . . . . |
| Kif26b | ENSMUSG00000026494 | ENSMUST00000161017 | 415 | . . . . . . . . |
| Kif2c | ENSMUSG00000028678 | ENSMUST00000065896 | 109 | ) ) ) ) ) ) ) ) |
| Kif5a | ENSMUSG00000074657 | ENSMUST00000217895 | 113 | . . . ( ( ( ( . |
| Kif5a | ENSMUSG00000074657 | ENSMUST00000099172 | 381 | ) . . . ) ) ) ) |
| Kifc5b | ENSMUSG00000024301 | ENSMUST00000078961 | 156 | ) ) . ) ) ) ) ) |
| Kirrel3 | ENSMUSG00000032036 | ENSMUST00000190549 | 344 | ) ) ) ) ) ) ) ) |
| Kiss1 | ENSMUSG00000102367 | ENSMUST00000178033 | 23 | ) . . . ) ) ) ) |
| Klc4 | ENSMUSG00000003546 | ENSMUST00000003642 | 4 | ( . . ( ( ( ( ( |
| Klhdc9 | ENSMUSG00000045259 | ENSMUST00000061878 | 121 | ) ) ) ) ) ) ) . |
| Klhl15 | ENSMUSG00000043929 | ENSMUST00000113908 | 347 | . . . . . . ) ) |
| Klhl15 | ENSMUSG00000043929 | ENSMUST00000113916 | 439 | ( ( ( ( ( . . . |
| Klhl4 | ENSMUSG00000025597 | ENSMUST00000040504 | 48 | . . . ) ) ) ) ) |
| Klhl4 | ENSMUSG00000025597 | ENSMUST00000113371 | 48 | . . . ) ) ) ) ) |
| Klk10 | ENSMUSG00000030693 | ENSMUST00000014058 | 84 | . . . . . . ) ) |
| Klk8 | ENSMUSG00000064023 | ENSMUST00000085461 | 466 | ( ( ( ( ( . . . |
| Klra1 | ENSMUSG00000079853 | ENSMUST00000032288 | 481 | ) ) ) ) ) ) ) . |
| Kmt5a | ENSMUSG00000049327 | ENSMUST00000059580 | 21 | ) ) . . . . . . |
| Kntc1 | ENSMUSG00000029414 | ENSMUST00000031366 | 111 | ) ) ) ) ) ) ) ) |
| Krcc1 | ENSMUSG00000053012 | ENSMUST00000168700 | 397 | ) ) ) ) ) ) . ) |
| Krit1 | ENSMUSG00000000600 | ENSMUST00000171023 | 90 | ) ) ) ) ) ) ) ) |
| Krt1 | ENSMUSG00000046834 | ENSMUST00000023790 | 14 | ) ) ) ) ) ) . . |
| Krt14 | ENSMUSG00000045545 | ENSMUST00000007272 | 15 | . . . . . . . . |
| Krt17 | ENSMUSG00000035557 | ENSMUST00000080893 | 15 | . . . . . . . . |
| Krt19 | ENSMUSG00000020911 | ENSMUST00000007317 | 229 | ) ) ) ) ) ) ) ) |
| Krt82 | ENSMUSG00000049548 | ENSMUST00000023713 | 16 | . . . . . . . . |
| Krtcap2 | ENSMUSG00000042747 | ENSMUST00000168900 | 140 | ) . . . . . . . |
| Krtcap2 | ENSMUSG00000042747 | ENSMUST00000040888 | 503 | . . . ) ) ) ) ) |
| Ksr2 | ENSMUSG00000061578 | ENSMUST00000180430 | 771 | . . . . . . . . |
| Kti12 | ENSMUSG00000073775 | ENSMUST00000102738 | 44 | ( . . ( ( ( ( ( |
| L1td1 | ENSMUSG00000087166 | ENSMUST00000154279 | 141 | . . . . ( ( ( ( |
| Lag3 | ENSMUSG00000030124 | ENSMUST00000032217 | 303 | . . . . . . . . |
| Lamb1 | ENSMUSG00000002900 | ENSMUST00000002979 | 84 | ) ) . ) . ) ) ) |
| Lancl1 | ENSMUSG00000026000 | ENSMUST00000113979 | 261 | . ) ) ) ) ) ) ) |
| Lat2 | ENSMUSG00000040751 | ENSMUST00000036362 | 110 | ) ) ) ) ) ) ) ) |

| | | | | |
|---|---|---|---|---|
| Lat2 | ENSMUSG00000040751 | ENSMUST00000077636 | 145 | ) ) ) ) ) ) ) ) |
| Lats1 | ENSMUSG00000040021 | ENSMUST00000165952 | 281 | . ) ) ) . . . . |
| Lcmt1 | ENSMUSG00000030763 | ENSMUST00000033025 | 94 | ) . ) ) . ) ) ) |
| Lctl | ENSMUSG00000032401 | ENSMUST00000034969 | 205 | . ( ( ( ( . . ( |
| Ldb1 | ENSMUSG00000025223 | ENSMUST00000056931 | 631 | ) ) ) ) ) ) ) ) |
| Ldhb | ENSMUSG00000030246 | ENSMUST00000032373 | 95 | ) ) ) ) . . . . |
| Ldhc | ENSMUSG00000030851 | ENSMUST00000014545 | 20 | . . . . . . ( ( |
| Lefty1 | ENSMUSG00000038793 | ENSMUST00000037361 | 69 | . . . . . . . . |
| Leng9 | ENSMUSG00000043432 | ENSMUST00000058358 | 612 | ) ( ( ( ( . ( ( |
| Lhfpl2 | ENSMUSG00000045312 | ENSMUST00000054274 | 282 | . ) ) ) . . . ) |
| Lif | ENSMUSG00000034394 | ENSMUST00000040750 | 78 | . . . . ) ) ) ) |
| Lime1 | ENSMUSG00000090077 | ENSMUST00000048077 | 962 | ) ) ) ) ) ) ) ( |
| Lin7a | ENSMUSG00000019906 | ENSMUST00000020057 | 148 | . . . . . . . . |
| Lipi | ENSMUSG00000032948 | ENSMUST00000062721 | 86 | ) ) ) ) . . ) ) |
| Lmbrd2 | ENSMUSG00000039704 | ENSMUST00000227556 | 158 | ) . ( ( ( ( ( ( |
| Lmo1 | ENSMUSG00000036111 | ENSMUST00000036992 | 465 | ( ( . . . . . ( |
| Lmod1 | ENSMUSG00000048096 | ENSMUST00000059352 | 134 | . ) ) ) ) . ) ) |
| Lmod3 | ENSMUSG00000044086 | ENSMUST00000095655 | 145 | ( ( ( ( ( ( ( ( |
| Lmtk3 | ENSMUSG00000062044 | ENSMUST00000072580 | 51 | . . . . . ( ( ( |
| Lmtk3 | ENSMUSG00000062044 | ENSMUST00000120005 | 206 | ) . ) ) ) ) ) ) |
| Lonrf3 | ENSMUSG00000016239 | ENSMUST00000016383 | 159 | ) ) ) ) . . . ) |
| Lor | ENSMUSG00000043165 | ENSMUST00000058150 | 15 | . . . . . . . . |
| Lox | ENSMUSG00000024529 | ENSMUST00000025409 | 328 | ) ) . . . . . ) |
| Lpgat1 | ENSMUSG00000026623 | ENSMUST00000110855 | 307 | . . . . . . . ) |
| Lrrc18 | ENSMUSG00000041673 | ENSMUST00000120866 | 269 | . ) ) . . ) ) ) |
| Lrrc18 | ENSMUSG00000041673 | ENSMUST00000038956 | 285 | . . ) ) ) ) ) ) |
| Lrrc4 | ENSMUSG00000049939 | ENSMUST00000062304 | 81 | . . . . . . ) ) |
| Lrrc49 | ENSMUSG00000047766 | ENSMUST00000114032 | 11 | ( ( ( ( ( ( . . |
| Lrrc49 | ENSMUSG00000047766 | ENSMUST00000166168 | 11 | ( ( ( ( ( ( . . |
| Lrrc75b | ENSMUSG00000046807 | ENSMUST00000051129 | 174 | . . . . . . . . |
| Lsm10 | ENSMUSG00000050188 | ENSMUST00000055575 | 162 | . . ) ) ) ) ) ) |
| Lsm4 | ENSMUSG00000031848 | ENSMUST00000034311 | 123 | ) ) . . . . . . |
| Ltbp1 | ENSMUSG00000001870 | ENSMUST00000112514 | 36 | . . . . . . . . |
| Ltbp1 | ENSMUSG00000001870 | ENSMUST00000112516 | 137 | . . . . . . . . |
| Ltn1 | ENSMUSG00000052299 | ENSMUST00000039449 | 52 | ) ) ) ) ) ) ) . |
| Luc7l2 | ENSMUSG00000029823 | ENSMUST00000057692 | 290 | ) ) . ) ) ) ) ) |
| Luc7l2 | ENSMUSG00000029823 | ENSMUST00000161538 | 290 | ) ) . ) ) ) ) ) |
| Ly6h | ENSMUSG00000022577 | ENSMUST00000023241 | 104 | ) ) ) ) ) . . . |
| Ly6h | ENSMUSG00000022577 | ENSMUST00000065417 | 181 | ) ) ) ) ) . . ) |
| Lypd6 | ENSMUSG00000050447 | ENSMUST00000112712 | 164 | . . . ) ) . . ) |
| Lypla1 | ENSMUSG00000025903 | ENSMUST00000027036 | 53 | ) ) ) ) ) ) ) ) |
| Lyrm1 | ENSMUSG00000030922 | ENSMUST00000208202 | 124 | . . . . . . . . |
| Lyrm1 | ENSMUSG00000030922 | ENSMUST00000106517 | 384 | ) ) ) ) . . ) ) |
| Mab21l1 | ENSMUSG00000056947 | ENSMUST00000075422 | 385 | ) ) ) ) ) ) ) ) |
| Macf1 | ENSMUSG00000028649 | ENSMUST00000082108 | 109 | ) ) ) . . . ( ( |
| Macf1 | ENSMUSG00000028649 | ENSMUST00000097897 | 109 | ) ) ) . . . ( ( |
| Maf1 | ENSMUSG00000022553 | ENSMUST00000023212 | 388 | . . ( ( ( ( . . |
| Maf1 | ENSMUSG00000022553 | ENSMUST00000161527 | 446 | ) ) ( ( ( ( . . |
| Mageb18 | ENSMUSG00000067649 | ENSMUST00000113955 | 358 | . ) ) ) . ) ) ) |
| Magi1 | ENSMUSG00000045095 | ENSMUST00000204347 | 545 | . . . . . ( ( ( |
| Magi2 | ENSMUSG00000040003 | ENSMUST00000088516 | 190 | . ( ( ( ( ( . . |
| Mak | ENSMUSG00000021363 | ENSMUST00000021792 | 216 | ) ) . . . . . . |
| Mak | ENSMUSG00000021363 | ENSMUST00000070193 | 319 | ) ) ) ) ) ) ) ) |
| Mak | ENSMUSG00000021363 | ENSMUST00000165087 | 321 | ) ) ) ) ) ) ) ) |
| Mamdc2 | ENSMUSG00000033207 | ENSMUST00000036069 | 193 | . . . ) ) . . . |
| Map2k1 | ENSMUSG00000004936 | ENSMUST00000005066 | 321 | . . . . . . . . |
| Mapk10 | ENSMUSG00000046709 | ENSMUST00000112847 | 525 | . . . ( ( ( ( . |
| Mapk10 | ENSMUSG00000046709 | ENSMUST00000170792 | 525 | . . . ( ( ( ( . |

| | | | | |
|---|---|---|---|---|
| March10 | ENSMUSG00000078627 | ENSMUST00000049995 | 303 | ) ) ) ) ) ) ) ) |
| March10 | ENSMUSG00000078627 | ENSMUST00000100332 | 303 | ) ) ) ) ) ) ) ) |
| March11 | ENSMUSG00000022269 | ENSMUST00000152841 | 98 | . . . . . . . . |
| March11 | ENSMUSG00000022269 | ENSMUST00000140840 | 98 | . . . . . . . . |
| March9 | ENSMUSG00000040502 | ENSMUST00000040307 | 344 | ) . ) ) ) ) ) ) |
| Mark2 | ENSMUSG00000024969 | ENSMUST00000051711 | 149 | ) ) ) ) ) ) ) ) |
| Mark2 | ENSMUSG00000024969 | ENSMUST00000164205 | 522 | ) ) ) ) ) ) ) ) |
| Mark2 | ENSMUSG00000024969 | ENSMUST00000032557 | 542 | ) ) ) ) ) ) ) ) |
| Mark4 | ENSMUSG00000030397 | ENSMUST00000085715 | 299 | ( ( ( . . . . . |
| Mars2 | ENSMUSG00000046994 | ENSMUST00000061334 | 11 | ) . . . . ( ( ( |
| Matk | ENSMUSG00000004933 | ENSMUST00000117488 | 329 | ( . . . . . . . |
| Matn2 | ENSMUSG00000022324 | ENSMUST00000022947 | 195 | ) ) . . . ) ) ) |
| Max | ENSMUSG00000059436 | ENSMUST00000110395 | 95 | ) ) ) ) . . . . |
| Max | ENSMUSG00000059436 | ENSMUST00000082136 | 111 | ) ) ) ) ) ) ) . |
| Mb21d1 | ENSMUSG00000032344 | ENSMUST00000070742 | 98 | . . . . . . . . |
| Mbl1 | ENSMUSG00000037780 | ENSMUST00000225792 | 73 | ) ) . . . ) ) ) |
| Mbnl1 | ENSMUSG00000027763 | ENSMUST00000192607 | 391 | . . . . . . . . |
| Mbnl1 | ENSMUSG00000027763 | ENSMUST00000194069 | 930 | . . . ) ) ) ) ) |
| Mbnl1 | ENSMUSG00000027763 | ENSMUST00000099087 | 1029 | . . . . . . . . |
| Mccc2 | ENSMUSG00000021646 | ENSMUST00000022148 | 94 | ) ) . . . . . ) |
| Mcl1 | ENSMUSG00000038612 | ENSMUST00000037947 | 39 | . . . . . . . . |
| Mcoln3 | ENSMUSG00000036853 | ENSMUST00000039450 | 107 | ) . . . . . . . |
| Mcts1 | ENSMUSG00000000355 | ENSMUST00000000365 | 95 | ) ) ) . . . . . |
| Mdc1 | ENSMUSG00000061607 | ENSMUST00000082337 | 92 | ) . ) ) ) ) . . |
| Med23 | ENSMUSG00000019984 | ENSMUST00000092646 | 52 | . . . . . . . . |
| Mef2d | ENSMUSG00000001419 | ENSMUST00000107559 | 198 | ) ) . . . . . . |
| Mef2d | ENSMUSG00000001419 | ENSMUST00000107558 | 198 | ) ) . . . . . . |
| Mef2d | ENSMUSG00000001419 | ENSMUST00000001455 | 398 | ) ) ) ) ) ) ) ) |
| Mei1 | ENSMUSG00000068117 | ENSMUST00000188048 | 291 | ) ) ) ) . . ) ) |
| Meis1 | ENSMUSG00000020160 | ENSMUST00000068264 | 659 | . . . . ) ) ) ) |
| Meis1 | ENSMUSG00000020160 | ENSMUST00000185131 | 670 | . . . . ) ) ) ) |
| Metap2 | ENSMUSG00000036112 | ENSMUST00000180840 | 675 | ) ) ) . . . ) ) |
| Mettl23 | ENSMUSG00000090266 | ENSMUST00000106370 | 282 | ) ) ) . . . . . |
| Mfge8 | ENSMUSG00000030605 | ENSMUST00000107409 | 65 | ) ) . . ) ) ) ) |
| Mfge8 | ENSMUSG00000030605 | ENSMUST00000032825 | 72 | ) ) . . ) ) ) ) |
| Mfn2 | ENSMUSG00000029020 | ENSMUST00000105716 | 102 | ) . ) ) . ) ) ) |
| Mfn2 | ENSMUSG00000029020 | ENSMUST00000105715 | 321 | ) . ) ) . ) ) ) |
| Mfsd11 | ENSMUSG00000020818 | ENSMUST00000021173 | 167 | ) ) ) ) ) ) ) ) |
| Mfsd6l | ENSMUSG00000048329 | ENSMUST00000053211 | 56 | . . . . . ) ) . |
| Mgam | ENSMUSG00000068587 | ENSMUST00000071535 | 103 | . . . ) ) ) ) ) |
| Mgat2 | ENSMUSG00000043998 | ENSMUST00000060579 | 462 | . . . ( ( ( ( ( |
| Mgat4c | ENSMUSG00000019888 | ENSMUST00000020039 | 112 | ) ) ) ) ) ) ) ) |
| Mgat4c | ENSMUSG00000019888 | ENSMUST00000163753 | 239 | ) ) . ( ( ( ( ( |
| Mgat4c | ENSMUSG00000019888 | ENSMUST00000179929 | 652 | ( ( ( ( ( ( ( ( |
| Mid1 | ENSMUSG00000035299 | ENSMUST00000036753 | 40 | . . . . ( ( ( ( |
| Mindy3 | ENSMUSG00000026767 | ENSMUST00000155530 | 117 | ( ( . . . ( ( ( |
| Mindy3 | ENSMUSG00000026767 | ENSMUST00000144645 | 134 | ( ( . . . ( ( ( |
| Mindy3 | ENSMUSG00000026767 | ENSMUST00000028105 | 152 | ( ( . . . ( ( ( |
| Mlip | ENSMUSG00000032355 | ENSMUST00000034910 | 76 | ) ) ) . . . . . |
| Mllt11 | ENSMUSG00000053192 | ENSMUST00000065482 | 551 | . . . . ( ( ( ( |
| Mlst8 | ENSMUSG00000024142 | ENSMUST00000070888 | 74 | . . . . . . . . |
| Mlst8 | ENSMUSG00000024142 | ENSMUST00000179163 | 79 | . . . . . . . . |
| Mmp16 | ENSMUSG00000028226 | ENSMUST00000029881 | 170 | . . . ( ( ( ( ( |
| Mmp23 | ENSMUSG00000029061 | ENSMUST00000030937 | 120 | ) ) ) ) ) . . . |
| Mmrn2 | ENSMUSG00000041445 | ENSMUST00000111908 | 60 | . . ( ( . . . . |
| Mogat1 | ENSMUSG00000012187 | ENSMUST00000012331 | 12 | ( ( ( ( ( ( ( . |
| Mok | ENSMUSG00000056458 | ENSMUST00000070565 | 135 | ) ) ) ) ) ) ) ) |
| Mospd3 | ENSMUSG00000037221 | ENSMUST00000111007 | 140 | . . . . . . . . |

| | | | | |
|---|---|---|---|---|
| Mpeg1 | ENSMUSG00000046805 | ENSMUST00000081035 | 174 | ) ) ) ) ) ) ) . |
| Mre11a | ENSMUSG00000031928 | ENSMUST00000115632 | 70 | . ( ( ( ( ( ( ( |
| Mre11a | ENSMUSG00000031928 | ENSMUST00000034405 | 92 | ( ( ( ( ( . . . |
| Mrpl17 | ENSMUSG00000030879 | ENSMUST00000124482 | 78 | . . . . ) ) ) ) |
| Mrpl23 | ENSMUSG00000037772 | ENSMUST00000038675 | 137 | ) ) ) ) ) ) ) ) |
| Mrpl36 | ENSMUSG00000021607 | ENSMUST00000022098 | 93 | . . . ) ) ) ) ) |
| Mrpl41 | ENSMUSG00000036850 | ENSMUST00000045604 | 167 | . ) ) ) ) ) ) ) |
| Mrpl44 | ENSMUSG00000026248 | ENSMUST00000027464 | 83 | ) ) ) ) . . ) ) |
| Mrpl53 | ENSMUSG00000030037 | ENSMUST00000113938 | 49 | . . . . . . . . |
| Mrps9 | ENSMUSG00000060679 | ENSMUST00000057208 | 21 | . . . . . . . . |
| Mrrf | ENSMUSG00000026887 | ENSMUST00000028250 | 328 | ) . ) ) ) . . . |
| Ms4a18 | ENSMUSG00000094584 | ENSMUST00000177684 | 57 | ( ( ( ( ( ( . . |
| Ms4a4b | ENSMUSG00000056290 | ENSMUST00000035258 | 72 | ) . . . ) ) ) ) |
| Ms4a5 | ENSMUSG00000054523 | ENSMUST00000067673 | 47 | ) ) . . ) ) ) ) |
| Msantd3 | ENSMUSG00000039693 | ENSMUST00000064807 | 168 | ) ) ) ) ) ) ) ) |
| Msh5 | ENSMUSG00000007035 | ENSMUST00000007250 | 211 | ) ) ) ) ) ) ) ) |
| Msl1 | ENSMUSG00000052915 | ENSMUST00000037915 | 262 | ) ) . . . . . . |
| Msl2 | ENSMUSG00000066415 | ENSMUST00000085177 | 477 | . . . . . ) ) ) |
| Mt4 | ENSMUSG00000031757 | ENSMUST00000034207 | 14 | . . . . . . . . |
| Mterf4 | ENSMUSG00000026273 | ENSMUST00000027492 | 32 | . . ) ) ) ) ) ) |
| Mtif2 | ENSMUSG00000020459 | ENSMUST00000093239 | 376 | . . . . . ) ) ) |
| Mtif3 | ENSMUSG00000016510 | ENSMUST00000016654 | 207 | ( ( ( ( . . . . |
| Muc1 | ENSMUSG00000042784 | ENSMUST00000041142 | 37 | . . . ) ) . . . |
| Mum1 | ENSMUSG00000020156 | ENSMUST00000020365 | 317 | . . . . . . ) ) |
| Musk | ENSMUSG00000057280 | ENSMUST00000084578 | 126 | . . . ) ) ) ) ) |
| Musk | ENSMUSG00000057280 | ENSMUST00000098057 | 126 | . . . ) ) ) ) ) |
| Musk | ENSMUSG00000057280 | ENSMUST00000102893 | 126 | . . . ) ) ) ) ) |
| Musk | ENSMUSG00000057280 | ENSMUST00000177951 | 126 | . . . ) ) ) ) ) |
| Mustn1 | ENSMUSG00000042485 | ENSMUST00000040715 | 345 | ) ) ) ) ) . . ( |
| Myadm | ENSMUSG00000068566 | ENSMUST00000096744 | 156 | . . . . ) ) ) ) |
| Myadm | ENSMUSG00000068566 | ENSMUST00000164553 | 168 | . . . . ) ) ) ) |
| Myadm | ENSMUSG00000068566 | ENSMUST00000203566 | 277 | . . . . ) ) ) ) |
| Myadm | ENSMUSG00000097829 | ENSMUST00000180998 | 277 | . . . . ) ) ) ) |
| Myadm | ENSMUSG00000068566 | ENSMUST00000203328 | 305 | . . . . ) ) ) ) |
| Mycl | ENSMUSG00000028654 | ENSMUST00000030407 | 398 | ) ) ) ) . . ) ) |
| Myct1 | ENSMUSG00000046916 | ENSMUST00000051809 | 56 | ) ) ) . . . ) ) |
| Myl3 | ENSMUSG00000059741 | ENSMUST00000079784 | 67 | ) ) ) . . ) ) ) |
| Myl4 | ENSMUSG00000061086 | ENSMUST00000106956 | 129 | ) ) ) . . . . . |
| Myl6b | ENSMUSG00000039824 | ENSMUST00000026428 | 36 | ( ( . . . ( ( . |
| Mylk2 | ENSMUSG00000027470 | ENSMUST00000028970 | 135 | . . ( ( ( ( ( ( |
| Mylpf | ENSMUSG00000030672 | ENSMUST00000032910 | 107 | ) ) ) . . . . . |
| Mymk | ENSMUSG00000009214 | ENSMUST00000009358 | 94 | . . . . . . . . |
| Mymx | ENSMUSG00000079471 | ENSMUST00000169137 | 78 | ) ) ) ) ) ) . . |
| Myo15 | ENSMUSG00000042678 | ENSMUST00000081823 | 10 | ( ( ( . . . . . |
| Myo1g | ENSMUSG00000020437 | ENSMUST00000003459 | 48 | . . . ( ( ( ( ( |
| Myog | ENSMUSG00000026459 | ENSMUST00000027730 | 53 | ) . . . ) ) ) ) |
| Myoz3 | ENSMUSG00000049173 | ENSMUST00000056533 | 68 | . . . . ) ) ) ) |
| Myt1 | ENSMUSG00000010505 | ENSMUST00000081125 | 0 | . . . . . . . . |
| Mzt2 | ENSMUSG00000022671 | ENSMUST00000023351 | 405 | ) ) ) ) . . . . |
| N4bp2l2 | ENSMUSG00000029655 | ENSMUST00000118316 | 56 | . . . . . . . . |
| Naa10 | ENSMUSG00000031388 | ENSMUST00000114387 | 33 | . . . . . . . . |
| Naa10 | ENSMUSG00000031388 | ENSMUST00000114389 | 69 | . . . . . . . ) |
| Naa10 | ENSMUSG00000031388 | ENSMUST00000033763 | 89 | . ) ) ) . . . ) |
| Naa25 | ENSMUSG00000042719 | ENSMUST00000042163 | 14 | ( ( ( ( ( . . . |
| Naca | ENSMUSG00000061315 | ENSMUST00000092048 | 9 | . . ( ( . . . . |
| Naca | ENSMUSG00000061315 | ENSMUST00000073868 | 69 | . . . . . . . . |
| Naip2 | ENSMUSG00000078945 | ENSMUST00000117913 | 83 | ( ( ( ( ( ( ( ( |
| Nanog | ENSMUSG00000012396 | ENSMUST00000012540 | 190 | . . . . . . . . |

| | | | | |
|---|---|---|---|---|
| Nanos1 | ENSMUSG00000072437 | ENSMUST00000088237 | 217 | ) ) ) ) ) ) ) ) |
| Nap1l3 | ENSMUSG00000055733 | ENSMUST00000079490 | 307 | . . . . . . ) ) |
| Naprt | ENSMUSG00000022574 | ENSMUST00000023237 | 152 | ) ) ) ) ) ) ) ) |
| Ncapd2 | ENSMUSG00000038252 | ENSMUST00000043848 | 141 | . . . . . . . . |
| Ndfip1 | ENSMUSG00000024425 | ENSMUST00000025293 | 114 | . . . . . . . . |
| Ndnf | ENSMUSG00000049001 | ENSMUST00000054351 | 479 | ) ) ) ) ) ) ) ) |
| Ndrg1 | ENSMUSG00000005125 | ENSMUST00000005256 | 110 | . . . . ( ( ( ( |
| Ndrg4 | ENSMUSG00000036564 | ENSMUST00000080666 | 78 | . . . . . . . . |
| Ndrg4 | ENSMUSG00000036564 | ENSMUST00000073139 | 104 | . ) ) . . ) ) ) |
| Ndufaf1 | ENSMUSG00000027305 | ENSMUST00000028768 | 253 | . . . . . . ) ) |
| Ndufb5 | ENSMUSG00000027673 | ENSMUST00000127477 | 69 | . . . . . . . . |
| Ndufs2 | ENSMUSG00000013593 | ENSMUST00000013737 | 4 | ( . . . . . . . |
| Nectin1 | ENSMUSG00000032012 | ENSMUST00000034510 | 633 | ) ) ) ) ) ) ) ) |
| Nedd4 | ENSMUSG00000032216 | ENSMUST00000034740 | 135 | . . . . . . . . |
| Nefl | ENSMUSG00000022055 | ENSMUST00000022639 | 20 | . . . . . . . . |
| Nes | ENSMUSG00000004891 | ENSMUST00000090973 | 23 | . . . . . . . . |
| Neu3 | ENSMUSG00000035239 | ENSMUST00000036331 | 214 | . . . . . ( ( ( |
| Nfat5 | ENSMUSG00000003847 | ENSMUST00000077440 | 20 | . . . . . . . . |
| Nfkbid | ENSMUSG00000036931 | ENSMUST00000046177 | 405 | . . . . . ) ) ) |
| Nhsl1 | ENSMUSG00000039835 | ENSMUST00000037341 | 547 | ) ) ) ) ) . . . |
| Nipsnap3a | ENSMUSG00000015242 | ENSMUST00000188045 | 74 | . . . . . . . . |
| Nkg7 | ENSMUSG00000004612 | ENSMUST00000070518 | 156 | ) ) . ) ) ) ) ) |
| Nkx2-1 | ENSMUSG00000001496 | ENSMUST00000178477 | 17 | . . . . . . . . |
| Nkx2-9 | ENSMUSG00000058669 | ENSMUST00000072631 | 206 | . . . . . . . . |
| Nlrp3 | ENSMUSG00000032691 | ENSMUST00000101148 | 130 | ( ( ( ( ( ( . . |
| Nlrp4b | ENSMUSG00000034087 | ENSMUST00000047809 | 108 | . ( ( ( ( . . . |
| Nlrp4b | ENSMUSG00000034087 | ENSMUST00000117413 | 150 | ) ) ) . . . . . |
| Nlrp4c | ENSMUSG00000034690 | ENSMUST00000037728 | 109 | ) . . . . ) ) ) |
| Nlrp4e | ENSMUSG00000045693 | ENSMUST00000076470 | 72 | . . . . . . . . |
| Nlrx1 | ENSMUSG00000032109 | ENSMUST00000169651 | 99 | ( . . ( ( ( ( ( |
| Nmnat2 | ENSMUSG00000042751 | ENSMUST00000043313 | 33 | . . . . . . ( ( |
| Noc3l | ENSMUSG00000024999 | ENSMUST00000025963 | 63 | . . . . . . . ) |
| Nosip | ENSMUSG00000003421 | ENSMUST00000003513 | 19 | ) ) . . . ( ( ( |
| Nosip | ENSMUSG00000003421 | ENSMUST00000107829 | 46 | ) ) ) ) ) ( ( ( |
| Nox4 | ENSMUSG00000030562 | ENSMUST00000032781 | 144 | ( . ( ( ( ( ( ( |
| Noxo1 | ENSMUSG00000019320 | ENSMUST00000019464 | 973 | ) ) ) . . . . . |
| Npc2 | ENSMUSG00000021242 | ENSMUST00000021668 | 251 | ) ) ) . . ) ) ) |
| Npr2 | ENSMUSG00000028469 | ENSMUST00000030191 | 152 | ( ( ( . . . . . |
| Nprl2 | ENSMUSG00000010057 | ENSMUST00000010201 | 60 | ) ) ) ) . . . ) |
| Nptn | ENSMUSG00000032336 | ENSMUST00000085651 | 88 | ( ( ( ( . . . . |
| Nr2f1 | ENSMUSG00000069171 | ENSMUST00000125176 | 8 | . . . . . . . . |
| Nr2f1 | ENSMUSG00000069171 | ENSMUST00000091458 | 1099 | ) ) ) ) ) ) ) ) |
| Nr5a1 | ENSMUSG00000026751 | ENSMUST00000028084 | 173 | ) ) ) . . ( ( ( |
| Nr5a2 | ENSMUSG00000026398 | ENSMUST00000027649 | 61 | ( ( . ( . . . . |
| Nrbp2 | ENSMUSG00000075590 | ENSMUST00000019516 | 173 | . . . . . ( ( ( |
| Nrk | ENSMUSG00000052854 | ENSMUST00000064937 | 224 | ( ( ( ( . . . . |
| Nrros | ENSMUSG00000052384 | ENSMUST00000115165 | 46 | . . . . . . ) ) |
| Ntmt1 | ENSMUSG00000026857 | ENSMUST00000041830 | 223 | ) ) . . ) ) ) ) |
| Ntn1 | ENSMUSG00000020902 | ENSMUST00000021284 | 189 | ) ) ) ) ) ) ) ) |
| Ntrk3 | ENSMUSG00000059146 | ENSMUST00000039438 | 111 | . . . . . . . . |
| Ntrk3 | ENSMUSG00000059146 | ENSMUST00000039431 | 537 | . . . . . . . . |
| Nudt12 | ENSMUSG00000024228 | ENSMUST00000025065 | 132 | ( ( ( ( ( ( ( ( |
| Nudt3 | ENSMUSG00000024213 | ENSMUST00000025050 | 226 | ) . ( ( . . . . |
| Nudt5 | ENSMUSG00000025817 | ENSMUST00000026927 | 338 | . . . . . . . . |
| Oasl1 | ENSMUSG00000041827 | ENSMUST00000031540 | 32 | . . . . . . . . |
| Ocel1 | ENSMUSG00000002396 | ENSMUST00000002469 | 283 | . . . . . ) ) ) |
| Ocel1 | ENSMUSG00000002396 | ENSMUST00000110051 | 324 | . . . . . ) ) ) |
| Ocm | ENSMUSG00000029618 | ENSMUST00000031622 | 19 | . . . . . . . . |

| | | | | |
|---|---|---|---|---|
| Odf1 | ENSMUSG00000061923 | ENSMUST00000081966 | 168 | . . . . . . . ( |
| Odf3b | ENSMUSG00000047394 | ENSMUST00000049968 | 31 | . . . ) ) ) ) ) |
| Ogfr | ENSMUSG00000049401 | ENSMUST00000029087 | 201 | ( ( ( . . . . . |
| Ogt | ENSMUSG00000034160 | ENSMUST00000044475 | 134 | ( . . . . . . . |
| Ola1 | ENSMUSG00000027108 | ENSMUST00000028517 | 84 | . . . . . . . . |
| Olfr1157 | ENSMUSG00000075143 | ENSMUST00000099841 | 184 | ) ) ) . . ) ) ) |
| Olfr1167 | ENSMUSG00000100899 | ENSMUST00000099832 | 37 | . . . . . . . ) |
| Olfr1186 | ENSMUSG00000082882 | ENSMUST00000121619 | 22 | . . . . . . . . |
| Olfr1228 | ENSMUSG00000099486 | ENSMUST00000190757 | 57 | . ) ) ) ) ) . . |
| Olfr1229 | ENSMUSG00000075095 | ENSMUST00000099788 | 9 | . . . . . . . . |
| Olfr1262 | ENSMUSG00000051313 | ENSMUST00000061701 | 9 | ( ( ( ( . . . . |
| Olfr1283 | ENSMUSG00000109322 | ENSMUST00000090328 | 10 | ( . ( ( ( . . . |
| Olfr1295 | ENSMUSG00000108919 | ENSMUST00000207786 | 27 | . . . . . . . . |
| Olfr1321 | ENSMUSG00000067971 | ENSMUST00000088876 | 11 | . . . . . . . . |
| Olfr1335 | ENSMUSG00000066061 | ENSMUST00000219094 | 58 | . . . . . . . . |
| Olfr1346 | ENSMUSG00000054938 | ENSMUST00000056144 | 7 | . . . . . . . . |
| Olfr1352 | ENSMUSG00000046493 | ENSMUST00000058991 | 207 | ) ) ) . . ) ) . |
| Olfr1367 | ENSMUSG00000045508 | ENSMUST00000059216 | 6 | . . . . . . . . |
| Olfr1377 | ENSMUSG00000061952 | ENSMUST00000075177 | 29 | . . ) ) . . ) ) |
| Olfr1404 | ENSMUSG00000049456 | ENSMUST00000056592 | 3 | ( . . . . . . . |
| Olfr1410 | ENSMUSG00000063583 | ENSMUST00000217316 | 916 | ) ) . . ) ) ) ) |
| Olfr1419 | ENSMUSG00000067545 | ENSMUST00000087857 | 69 | . ) ) ) ) ) ) ) |
| Olfr1437 | ENSMUSG00000096436 | ENSMUST00000052558 | 197 | ) ) ) . ) ) ) ) |
| Olfr145 | ENSMUSG00000066748 | ENSMUST00000086062 | 642 | . . . . . . . . |
| Olfr146 | ENSMUSG00000058820 | ENSMUST00000073671 | 55 | ) ) . . . . . ) |
| Olfr1495 | ENSMUSG00000047207 | ENSMUST00000061669 | 22 | . . . . . . . . |
| Olfr1496 | ENSMUSG00000048356 | ENSMUST00000219412 | 47 | ) ) . . . ) ) ) |
| Olfr1507 | ENSMUSG00000059887 | ENSMUST00000073571 | 49 | . . . . ) ) ) ) |
| Olfr172 | ENSMUSG00000071510 | ENSMUST00000095991 | 46 | ) ) ) ) . . . . |
| Olfr187 | ENSMUSG00000043357 | ENSMUST00000207673 | 115 | ( ( ( . . . . . |
| Olfr19 | ENSMUSG00000048101 | ENSMUST00000057886 | 7 | ( ( . . . . . . |
| Olfr203 | ENSMUSG00000068182 | ENSMUST00000215893 | 165 | . ) ) ) ) ) . . |
| Olfr223 | ENSMUSG00000048919 | ENSMUST00000061293 | 66 | ) ) ) . . . . . |
| Olfr235 | ENSMUSG00000060049 | ENSMUST00000073507 | 34 | . ( ( . . . . . |
| Olfr313 | ENSMUSG00000070438 | ENSMUST00000082220 | 31 | . . . . . . . ( |
| Olfr316 | ENSMUSG00000107677 | ENSMUST00000203731 | 21 | ( ( . . . . . ( |
| Olfr318 | ENSMUSG00000108265 | ENSMUST00000189911 | 6 | . . . . . . . ( |
| Olfr434 | ENSMUSG00000059411 | ENSMUST00000076752 | 16 | . . . . . . . . |
| Olfr47 | ENSMUSG00000061210 | ENSMUST00000078057 | 16 | . . . . . . . . |
| Olfr50 | ENSMUSG00000111021 | ENSMUST00000112950 | 189 | . . . . . . . . |
| Olfr619 | ENSMUSG00000073944 | ENSMUST00000098196 | 36 | ( ( ( ( . . . . |
| Olfr620 | ENSMUSG00000045132 | ENSMUST00000052152 | 151 | ) ) ) ) ) ) ) . |
| Olfr644 | ENSMUSG00000110012 | ENSMUST00000077417 | 83 | ) ) ) ) . . ) ) |
| Olfr645 | ENSMUSG00000051340 | ENSMUST00000057104 | 297 | ( ( ( ( ( . . . |
| Olfr685 | ENSMUSG00000047794 | ENSMUST00000051355 | 66 | ) ) ) ) . ) ) ) |
| Olfr725 | ENSMUSG00000068437 | ENSMUST00000089844 | 84 | . . ) ) ) ) ) . |
| Olfr727 | ENSMUSG00000059488 | ENSMUST00000079142 | 11 | . . ) ) ) ) ) . |
| Olfr767 | ENSMUSG00000059762 | ENSMUST00000082131 | 55 | ( ( . . . . . . |
| Olfr769 | ENSMUSG00000042801 | ENSMUST00000050915 | 41 | . . ( ( ( ( . . |
| Olfr78 | ENSMUSG00000043366 | ENSMUST00000168007 | 314 | ) ) ) . ) ) ) ) |
| Olfr78 | ENSMUSG00000043366 | ENSMUST00000060187 | 1107 | ) ) ) . ) ) ) ) |
| Olfr785 | ENSMUSG00000111732 | ENSMUST00000204402 | 114 | . . . . . . . . |
| Olfr825 | ENSMUSG00000058084 | ENSMUST00000076814 | 90 | ) . . . . . . . |
| Olfr837 | ENSMUSG00000110621 | ENSMUST00000212482 | 41 | . . . . . . . . |
| Olfr866 | ENSMUSG00000050803 | ENSMUST00000062248 | 83 | . . . . . ) ) ) |
| Olfr878 | ENSMUSG00000066747 | ENSMUST00000086061 | 36 | ) . . . . . . . |
| Olfr889 | ENSMUSG00000096356 | ENSMUST00000217286 | 151 | . . ( ( ( ( . . |
| Olfr902 | ENSMUSG00000049334 | ENSMUST00000050733 | 48 | ( ( ( ( ( . . . |

| | | | | |
|---|---|---|---|---|
| Olfr913 | ENSMUSG00000059189 | ENSMUST00000081095 | 298 | . . . . ) ) ) ) |
| Olfr926 | ENSMUSG00000064333 | ENSMUST00000078289 | 12 | . . . . . . . ( |
| Olfr976 | ENSMUSG00000047352 | ENSMUST00000169307 | 22 | . . . . . . . . |
| Olfr987 | ENSMUSG00000075223 | ENSMUST00000099929 | 30 | . . . . . . . . |
| Olfr988 | ENSMUSG00000075222 | ENSMUST00000099928 | 30 | . . . . . . . . |
| Oosp1 | ENSMUSG00000041857 | ENSMUST00000048214 | 82 | ( ( ( . . . . ( |
| Opa1 | ENSMUSG00000038084 | ENSMUST00000038867 | 60 | ) ) ) ) ) ) . . |
| Opa1 | ENSMUSG00000038084 | ENSMUST00000160597 | 106 | ) ) ) ) ) ) . . |
| Oprl1 | ENSMUSG00000027584 | ENSMUST00000108763 | 205 | . . . . ) ) ) ) |
| Orc6 | ENSMUSG00000031697 | ENSMUST00000034132 | 125 | . ) ) ) ) ) ) . |
| Orm1 | ENSMUSG00000039196 | ENSMUST00000030044 | 1 | . . . . . . . . |
| Orm2 | ENSMUSG00000061540 | ENSMUST00000075341 | 1 | . . . . . . . . |
| Orm3 | ENSMUSG00000028359 | ENSMUST00000006687 | 1 | . . . . . . . ( |
| Oscar | ENSMUSG00000054594 | ENSMUST00000039507 | 24 | ) ) ) . . . ) ) |
| Oscar | ENSMUSG00000054594 | ENSMUST00000108645 | 48 | ) ) ) . . . ) ) |
| Otof | ENSMUSG00000062372 | ENSMUST00000114747 | 60 | ) . . ) ) ) ) ) |
| Otof | ENSMUSG00000062372 | ENSMUST00000074171 | 90 | ) ) ) ) ) ) ) ) |
| Otud3 | ENSMUSG00000041161 | ENSMUST00000097830 | 83 | ( . ( . . . . . |
| Oxgr1 | ENSMUSG00000044819 | ENSMUST00000058213 | 492 | . . ( ( ( ( ( . |
| P2ry13 | ENSMUSG00000036362 | ENSMUST00000040622 | 44 | ( . . . . . . . |
| P4ha2 | ENSMUSG00000018906 | ENSMUST00000019050 | 212 | ) ) ) ) ) ) ) ) |
| P4ha2 | ENSMUSG00000018906 | ENSMUST00000093107 | 251 | . . ) ) ) ) ) ) |
| Pacrg | ENSMUSG00000037196 | ENSMUST00000041463 | 352 | ( ( ( ( ( . ( ( |
| Pacs1 | ENSMUSG00000024855 | ENSMUST00000025786 | 208 | ) . . ) ) ) ) ) |
| Pafah2 | ENSMUSG00000037366 | ENSMUST00000105870 | 7 | . ( ( ( ( ( ( ( |
| Paics | ENSMUSG00000029247 | ENSMUST00000031160 | 472 | . . . . . . . ( |
| Pak3 | ENSMUSG00000031284 | ENSMUST00000112864 | 454 | ( ( ( ( . . . . |
| Palm2 | ENSMUSG00000090053 | ENSMUST00000102904 | 180 | ) . . ) ) ) ) ) |
| Pan3 | ENSMUSG00000029647 | ENSMUST00000031651 | 365 | ) . ) ) ) ) . ) |
| Pank1 | ENSMUSG00000033610 | ENSMUST00000112460 | 145 | . . . . . . . . |
| Paqr7 | ENSMUSG00000037348 | ENSMUST00000095074 | 128 | ) . . ) ) . ) ) |
| Paqr7 | ENSMUSG00000037348 | ENSMUST00000081525 | 351 | ) ) ) ) . . ( ( |
| Parp12 | ENSMUSG00000038507 | ENSMUST00000038398 | 116 | ( . . . . . . . |
| Parp3 | ENSMUSG00000023249 | ENSMUST00000112479 | 150 | . . . . ) ) ) ) |
| Parp3 | ENSMUSG00000023249 | ENSMUST00000067218 | 430 | ) ) ) ) . ) ) ) |
| Pate3 | ENSMUSG00000094995 | ENSMUST00000178236 | 42 | . . . . . . . ) |
| Pate4 | ENSMUSG00000032099 | ENSMUST00000034610 | 20 | . . . . . . . . |
| Patl1 | ENSMUSG00000046139 | ENSMUST00000061618 | 143 | ) ) ) . . . . . |
| Pax2 | ENSMUSG00000004231 | ENSMUST00000174490 | 1265 | ) ) ) ) ) ) ) ) |
| Pax9 | ENSMUSG00000001497 | ENSMUST00000001538 | 170 | ) ) ) ) ) ) ) ) |
| Pbx3 | ENSMUSG00000038718 | ENSMUST00000113132 | 18 | . . . . . . . . |
| Pbx3 | ENSMUSG00000038718 | ENSMUST00000040638 | 24 | . . . . . . . . |
| Pcdh1 | ENSMUSG00000051375 | ENSMUST00000057185 | 141 | ) . ) ) ) ) ) ) |
| Pcdh10 | ENSMUSG00000049100 | ENSMUST00000170695 | 810 | ) ) ) . ( ( ( ( |
| Pcdh10 | ENSMUSG00000049100 | ENSMUST00000166126 | 812 | ) ) ) . ( ( ( ( |
| Pcdh10 | ENSMUSG00000049100 | ENSMUST00000171554 | 831 | ) ) ) ) ( ( ( ( |
| Pcdh10 | ENSMUSG00000049100 | ENSMUST00000193252 | 839 | ) ) . . ) ) ) ) |
| Pcdh11x | ENSMUSG00000034755 | ENSMUST00000113358 | 275 | . . . . . . . . |
| Pcdh11x | ENSMUSG00000034755 | ENSMUST00000113364 | 343 | . . . . . . . . |
| Pcdhga7 | ENSMUSG00000103472 | ENSMUST00000192511 | 84 | ) ) . ) ) ) . . |
| Pcsk1 | ENSMUSG00000021587 | ENSMUST00000022075 | 187 | ) ) ) . . . ) ) |
| Pdcl | ENSMUSG00000009030 | ENSMUST00000009174 | 114 | . ( ( ( ( ( . . |
| Pde4a | ENSMUSG00000032177 | ENSMUST00000003395 | 368 | . . . . . . . . |
| Pdlim1 | ENSMUSG00000055044 | ENSMUST00000068439 | 153 | . . . . . . ) . |
| Pdlim2 | ENSMUSG00000022090 | ENSMUST00000153735 | 90 | . . . . . . ( ( |
| Pdlim5 | ENSMUSG00000028273 | ENSMUST00000170361 | 0 | . . . . . . . . |
| Pdlim5 | ENSMUSG00000028273 | ENSMUST00000168967 | 0 | . . . . . . . . |
| Pdlim5 | ENSMUSG00000028273 | ENSMUST00000090134 | 22 | ( ( ( . . . . . |

| | | | | |
|---|---|---|---|---|
| Pdlim5 | ENSMUSG00000028273 | ENSMUST00000196908 | 33 | ( ( ( ( ( ( ( ( |
| Pdlim5 | ENSMUSG00000028273 | ENSMUST00000198381 | 64 | ( ( ( ( ( . . . |
| Pdlim5 | ENSMUSG00000028273 | ENSMUST00000029941 | 108 | ) ) ) ) ) ) ) ) |
| Pdzd4 | ENSMUSG00000002006 | ENSMUST00000002080 | 604 | . ) ) ) ) ) . . |
| Pdzd4 | ENSMUSG00000002006 | ENSMUST00000114438 | 604 | . ) ) ) ) ) . . |
| Pdzk1 | ENSMUSG00000038298 | ENSMUST00000107070 | 472 | ( . . ( ( ( ( ( |
| Pdzrn4 | ENSMUSG00000036218 | ENSMUST00000035399 | 227 | ) ) . . . . . . |
| Peg12 | ENSMUSG00000070526 | ENSMUST00000094339 | 74 | ( ( . . . . . . |
| Pelo | ENSMUSG00000042275 | ENSMUST00000109226 | 181 | ) . . ) ) ) ) ) |
| Perp | ENSMUSG00000019851 | ENSMUST00000019998 | 75 | ( . . ( ( . . . |
| Pex5 | ENSMUSG00000005069 | ENSMUST00000112532 | 0 | . . . . . . . . |
| Pfas | ENSMUSG00000020899 | ENSMUST00000021282 | 66 | . ( ( ( ( ( . . |
| Pfdn5 | ENSMUSG00000001289 | ENSMUST00000166658 | 61 | ) ) ) ) . . . . |
| Pfkfb2 | ENSMUSG00000026409 | ENSMUST00000189534 | 142 | . . . ( ( ( ( . |
| Pfkm | ENSMUSG00000033065 | ENSMUST00000051226 | 82 | . ( ( . . ( ( ( |
| Pgc | ENSMUSG00000023987 | ENSMUST00000024782 | 30 | . . . ) ) ) ) . |
| Pgd | ENSMUSG00000028961 | ENSMUST00000084124 | 76 | ) ) ) ) . . . . |
| Pgk1 | ENSMUSG00000062070 | ENSMUST00000081593 | 106 | ) ) ) . . . ) ) |
| Pgr | ENSMUSG00000031870 | ENSMUST00000189181 | 581 | ( ( ( . . . . . |
| Phactr1 | ENSMUSG00000054728 | ENSMUST00000148891 | 32 | . . . . . . . . |
| Phf12 | ENSMUSG00000037791 | ENSMUST00000049167 | 434 | ) ) ) ) . ) ) ) |
| Phyhip | ENSMUSG00000003469 | ENSMUST00000003561 | 288 | . . . . . . . . |
| Pid1 | ENSMUSG00000045658 | ENSMUST00000168574 | 209 | ) ) ) ) . ) ) ) |
| Pigf | ENSMUSG00000024145 | ENSMUST00000024957 | 79 | ) ) ) ) ) ) ) . |
| Pigq | ENSMUSG00000025728 | ENSMUST00000097368 | 78 | ( ( ( . . ( ( ( ( |
| Pigq | ENSMUSG00000025728 | ENSMUST00000026823 | 191 | ) ) ) ) . ( ( ( |
| Pigx | ENSMUSG00000023791 | ENSMUST00000189013 | 130 | ( ( ( . . . . . |
| Pigx | ENSMUSG00000023791 | ENSMUST00000096109 | 141 | ( ( ( . . . . . |
| Pih1d3 | ENSMUSG00000026063 | ENSMUST00000027230 | 41 | ) ) . . ) ) . ) |
| Pik3c3 | ENSMUSG00000033628 | ENSMUST00000115812 | 143 | ( ( ( . . . ( . |
| Pik3cg | ENSMUSG00000020573 | ENSMUST00000053215 | 85 | ) ) ) ) ) ) . . |
| Pik3r1 | ENSMUSG00000041417 | ENSMUST00000035532 | 298 | ) ) ) ) ) ) ) ) |
| Pip4k2b | ENSMUSG00000018547 | ENSMUST00000018691 | 71 | ) ) ) ) . . ) ) |
| Pitpnc1 | ENSMUSG00000040430 | ENSMUST00000103064 | 730 | ) ) ) ) ) ) . ) |
| Pkib | ENSMUSG00000019876 | ENSMUST00000177473 | 206 | . . ) ) ) ) ) ) |
| Pkib | ENSMUSG00000019876 | ENSMUST00000177325 | 221 | . . ) ) ) ) ) ) |
| Pkib | ENSMUSG00000019876 | ENSMUST00000075992 | 256 | . . ) ) ) ) ) ) |
| Pkib | ENSMUSG00000019876 | ENSMUST00000095668 | 326 | . . ) ) ) ) ) ) |
| Pkib | ENSMUSG00000019876 | ENSMUST00000175852 | 411 | . . ) ) ) ) ) ) |
| Pkib | ENSMUSG00000019876 | ENSMUST00000066028 | 520 | . . ) ) ) ) ) ) |
| Pla2g16 | ENSMUSG00000060675 | ENSMUST00000025925 | 82 | ) ) ) . ( ( ( ( |
| Pla2g2d | ENSMUSG00000041202 | ENSMUST00000105806 | 54 | . . . . ) ) ) ) |
| Pla2g4e | ENSMUSG00000050211 | ENSMUST00000090071 | 371 | . ( ( ( ( ( . . |
| Plagl1 | ENSMUSG00000019817 | ENSMUST00000121646 | 479 | . . . . . . . ( |
| Plcb2 | ENSMUSG00000040061 | ENSMUST00000102524 | 47 | ) . ) ) ) . . . |
| Plekha5 | ENSMUSG00000030231 | ENSMUST00000087622 | 52 | ( ( . ( ( . . . |
| Plekhd1 | ENSMUSG00000066438 | ENSMUST00000140770 | 231 | ( ( ( ( ( ( ( . |
| Plekhg6 | ENSMUSG00000038167 | ENSMUST00000042647 | 679 | ) ) ) ) ) ) ) . |
| Plekhm3 | ENSMUSG00000051344 | ENSMUST00000097713 | 321 | . ) ) ) . . ) ) |
| Plin2 | ENSMUSG00000028494 | ENSMUST00000000466 | 120 | . . . ( ( ( ( ( |
| Plod3 | ENSMUSG00000004846 | ENSMUST00000004968 | 575 | . . . . . . . . |
| Pms2 | ENSMUSG00000079109 | ENSMUST00000148011 | 28 | . ( ( ( ( ( ( ( |
| Pmvk | ENSMUSG00000027952 | ENSMUST00000184515 | 220 | ) ) ) ) ) ) ) ) |
| Pnmal2 | ENSMUSG00000070802 | ENSMUST00000094807 | 359 | . . . . . . . . |
| Pnpla5 | ENSMUSG00000018868 | ENSMUST00000019012 | 4 | ( ( ( . . . . . |
| Pnpla7 | ENSMUSG00000036833 | ENSMUST00000045295 | 216 | ) ) ) ) ) ) ) ) |
| Pnpo | ENSMUSG00000018659 | ENSMUST00000018803 | 32 | ) . . . . ) ) ) |
| Poll | ENSMUSG00000025218 | ENSMUST00000026239 | 8 | . . . . . . . . |

| | | | | |
|---|---|---|---|---|
| Polm | ENSMUSG00000020474 | ENSMUST00000020767 | 201 | ) ) ) ) ) . . . |
| Poln | ENSMUSG00000045102 | ENSMUST00000202638 | 82 | . . ) ) ) ) ) ) |
| Polr2k | ENSMUSG00000045996 | ENSMUST00000057177 | 31 | ( ( ( ( ( ( ( ( |
| Polr2k | ENSMUSG00000045996 | ENSMUST00000180159 | 54 | ) ) ) ) ) ) . . |
| Polr3b | ENSMUSG00000034453 | ENSMUST00000077175 | 152 | . ) ) ) . ) ) ) |
| Porcn | ENSMUSG00000031169 | ENSMUST00000077595 | 113 | ) . ) ) . . . . |
| Porcn | ENSMUSG00000031169 | ENSMUST00000089403 | 113 | ) . ) ) . . . . |
| Porcn | ENSMUSG00000031169 | ENSMUST00000089402 | 113 | ) . ) ) . . . . |
| Porcn | ENSMUSG00000031169 | ENSMUST00000082320 | 137 | ) . ) ) . . . . |
| Postn | ENSMUSG00000027750 | ENSMUST00000107985 | 4 | . ( ( . ( ( ( ( |
| Postn | ENSMUSG00000027750 | ENSMUST00000117373 | 4 | . ( ( . ( ( ( ( |
| Ppl | ENSMUSG00000039457 | ENSMUST00000035672 | 53 | . . . . . . . . |
| Ppm1b | ENSMUSG00000061130 | ENSMUST00000080217 | 372 | ) ) ) ) ) ) ) ) |
| Ppm1b | ENSMUSG00000061130 | ENSMUST00000112304 | 372 | ) ) ) ) ) ) ) ) |
| Ppm1e | ENSMUSG00000046442 | ENSMUST00000055438 | 86 | ( . . . . . . . |
| Ppm1k | ENSMUSG00000037826 | ENSMUST00000042766 | 190 | . . . . . . ) ) |
| Ppm1m | ENSMUSG00000020253 | ENSMUST00000076258 | 192 | . . . . . . . . |
| Ppp1r10 | ENSMUSG00000039220 | ENSMUST00000087210 | 453 | ) ) ) ) ) ) ) ) |
| Ppp1r18 | ENSMUSG00000034595 | ENSMUST00000122899 | 113 | . . . . . . . . |
| Ppp2r1a | ENSMUSG00000007564 | ENSMUST00000007708 | 113 | ) ) . . . . . . |
| Ppp2r2b | ENSMUSG00000024500 | ENSMUST00000025377 | 215 | ) ) . . ( ( ( ( |
| Ppp2r5e | ENSMUSG00000021051 | ENSMUST00000021447 | 545 | ) ) ) ) ) ) ) ) |
| Ppp6r3 | ENSMUSG00000024908 | ENSMUST00000172362 | 28 | ( . . . . . ( ( |
| Pqbp1 | ENSMUSG00000031157 | ENSMUST00000115655 | 179 | . . . . . . ( ( |
| Pqlc2 | ENSMUSG00000028744 | ENSMUST00000053862 | 254 | ) . . . . . . ) |
| Prcp | ENSMUSG00000061119 | ENSMUST00000076052 | 44 | ( . . . . . . . |
| Prdx1 | ENSMUSG00000028691 | ENSMUST00000135573 | 61 | . ) ) ) ) ) ) ) |
| Prdx2 | ENSMUSG00000005161 | ENSMUST00000005292 | 94 | ) ) ) ) . . . ) |
| Prkag2 | ENSMUSG00000028944 | ENSMUST00000076306 | 316 | . . . ) ) ) ) . |
| Prl7b1 | ENSMUSG00000021347 | ENSMUST00000080595 | 18 | ) . . . . ) ) ) |
| Prmt9 | ENSMUSG00000037134 | ENSMUST00000056237 | 97 | . . . ) ) ) ) ) |
| Procr | ENSMUSG00000027611 | ENSMUST00000029140 | 213 | ) ) . . ( ( ( ( |
| Prodh2 | ENSMUSG00000036892 | ENSMUST00000058280 | 25 | . . . . . . ( ( |
| Proser1 | ENSMUSG00000049504 | ENSMUST00000058577 | 510 | ( ( ( ( ( ( ( ( |
| Proser2 | ENSMUSG00000045319 | ENSMUST00000114942 | 175 | ) . ) ) ) ) ) ) |
| Proser2 | ENSMUSG00000045319 | ENSMUST00000054254 | 256 | . . . ) ) ) ) ( |
| Prpf18 | ENSMUSG00000039449 | ENSMUST00000035721 | 22 | ( ( ( ( . . ( ( |
| Prpf38a | ENSMUSG00000063800 | ENSMUST00000079213 | 143 | . . . . . . . . |
| Prpf38b | ENSMUSG00000027881 | ENSMUST00000029480 | 265 | . . . . . . . . |
| Prpf40b | ENSMUSG00000023007 | ENSMUST00000118287 | 40 | . . . . . . . . |
| Prpf40b | ENSMUSG00000023007 | ENSMUST00000145482 | 199 | ) ) ) ) ) ) ) ) |
| Prpf4b | ENSMUSG00000021413 | ENSMUST00000077853 | 275 | ( . . ( ( . ( ( |
| Prps1l1 | ENSMUSG00000092305 | ENSMUST00000134550 | 83 | . . . ( ( ( ( ( |
| Prps2 | ENSMUSG00000025742 | ENSMUST00000026839 | 115 | ) . . . . . . . |
| Prpsap2 | ENSMUSG00000020528 | ENSMUST00000168115 | 82 | ) ) . . . . . ) |
| Prr13 | ENSMUSG00000023048 | ENSMUST00000023810 | 160 | ( ( ( ( ( ( ( . |
| Prr14l | ENSMUSG00000054280 | ENSMUST00000120129 | 268 | . . ) ) ) ) ) ) |
| Prr3 | ENSMUSG00000038500 | ENSMUST00000165613 | 159 | ) . ) ) ) ) ) ) |
| Prr3 | ENSMUSG00000038500 | ENSMUST00000055454 | 947 | ) . ) ) ) ) ) ) |
| Prr5 | ENSMUSG00000036106 | ENSMUST00000171460 | 43 | . . . . . . ) ) |
| Prrt2 | ENSMUSG00000045114 | ENSMUST00000159916 | 231 | ) ) ) ) . . . . |
| Prrx1 | ENSMUSG00000026586 | ENSMUST00000174397 | 936 | ( ( ( ( ( ( ( ( |
| Prrx1 | ENSMUSG00000026586 | ENSMUST00000075805 | 940 | ( ( ( ( ( ( ( ( |
| Prrx1 | ENSMUSG00000026586 | ENSMUST00000027878 | 978 | ( ( ( ( ( ( ( ( |
| Prss3 | ENSMUSG00000071519 | ENSMUST00000096003 | 70 | . . . . . . . . |
| Prss32 | ENSMUSG00000048992 | ENSMUST00000061725 | 218 | ) ) ) ) ) ) ) ) |
| Prss37 | ENSMUSG00000029909 | ENSMUST00000031967 | 190 | ) ) ) ) ) ) ) ) |
| Prss54 | ENSMUSG00000048400 | ENSMUST00000052690 | 43 | ( . . . . . . . |

| | | | | |
|---|---|---|---|---|
| Prss8 | ENSMUSG00000030800 | ENSMUST00000032988 | 176 | . . . . . . . . |
| Psd3 | ENSMUSG00000030465 | ENSMUST00000098696 | 350 | . . ) ) . ) ) ) |
| Psg23 | ENSMUSG00000074359 | ENSMUST00000057810 | 156 | ) ( ( ( ( ( ( . |
| Psme1 | ENSMUSG00000022216 | ENSMUST00000174259 | 151 | ) ) ) . . . . . |
| Psmf1 | ENSMUSG00000032869 | ENSMUST00000042452 | 143 | . . . . . . . . |
| Ptbp1 | ENSMUSG00000006498 | ENSMUST00000095457 | 31 | . . . . . ) ) ) |
| Ptbp1 | ENSMUSG00000006498 | ENSMUST00000165704 | 70 | . . . . . ) ) ) |
| Ptbp1 | ENSMUSG00000006498 | ENSMUST00000172282 | 272 | ) . ) ) ) ) ) . |
| Pten | ENSMUSG00000013663 | ENSMUST00000013807 | 919 | . . . . . . . . |
| Ptgdr | ENSMUSG00000071489 | ENSMUST00000095959 | 94 | ( ( ( . ( ( ( ( |
| Ptgdr2 | ENSMUSG00000034117 | ENSMUST00000037261 | 237 | . . . . . . . . |
| Ptger2 | ENSMUSG00000037759 | ENSMUST00000046891 | 670 | ) . . . ) ) ) ) |
| Ptp4a1 | ENSMUSG00000026064 | ENSMUST00000027232 | 629 | . . . . . . ) ) |
| Ptp4a2 | ENSMUSG00000028788 | ENSMUST00000165853 | 646 | ) ) ) ) ) ) . ) |
| Ptp4a2 | ENSMUSG00000028788 | ENSMUST00000030578 | 935 | . . . . . . . . |
| Ptpn4 | ENSMUSG00000026384 | ENSMUST00000064091 | 671 | . . . . . . . . |
| Ptprg | ENSMUSG00000021745 | ENSMUST00000022264 | 691 | . . . . . ( ( ( |
| Ptx3 | ENSMUSG00000027832 | ENSMUST00000029421 | 109 | ( . . . . . . . |
| Pus3 | ENSMUSG00000032103 | ENSMUST00000034615 | 73 | ( ( ( ( ( . . . |
| Pycr1 | ENSMUSG00000025140 | ENSMUST00000170556 | 360 | . . . . . . . ) |
| Pycr2 | ENSMUSG00000026520 | ENSMUST00000027802 | 97 | . . . . ) ) ) ) |
| Pyroxd1 | ENSMUSG00000041671 | ENSMUST00000041852 | 11 | . . . . . . . . |
| Qdpr | ENSMUSG00000015806 | ENSMUST00000015950 | 52 | ) ) ) ) . . . ) |
| Qrsl1 | ENSMUSG00000019863 | ENSMUST00000020012 | 59 | . . . . ) ) ) ) |
| Rab28 | ENSMUSG00000029128 | ENSMUST00000201422 | 94 | ) ) ) ) . . . . |
| Rab28 | ENSMUSG00000029128 | ENSMUST00000031011 | 127 | ) ) ) ) . . ) ) |
| Rab28 | ENSMUSG00000029128 | ENSMUST00000202913 | 130 | ) ) ) ) ) . ) ) |
| Rab5b | ENSMUSG00000000711 | ENSMUST00000000727 | 109 | ) ) ) ) ) ) ) . |
| Rab5c | ENSMUSG00000019173 | ENSMUST00000019317 | 147 | . . . ) ) ) ) ) |
| Rab5c | ENSMUSG00000019173 | ENSMUST00000107364 | 147 | . . . ) ) ) ) ) |
| Rabggta | ENSMUSG00000040472 | ENSMUST00000169237 | 263 | . . . . . . . . |
| Rad17 | ENSMUSG00000021635 | ENSMUST00000177848 | 62 | ) ) . . . . ) ) |
| Rad50 | ENSMUSG00000020380 | ENSMUST00000020649 | 186 | ) ) ) . ) ) ) ) |
| Rad51d | ENSMUSG00000018841 | ENSMUST00000021033 | 71 | ) . . . . ( ( ( |
| Rad51d | ENSMUSG00000018841 | ENSMUST00000092844 | 71 | ) . . . . ( ( ( |
| Rad51d | ENSMUSG00000018841 | ENSMUST00000018985 | 96 | ) . . . . ( ( ( |
| Radil | ENSMUSG00000029576 | ENSMUST00000085758 | 77 | ) . . ) ) ) ) ) |
| Raet1d | ENSMUSG00000078452 | ENSMUST00000095795 | 49 | ) . . . . . ) ) |
| Raet1d | ENSMUSG00000078452 | ENSMUST00000182677 | 203 | . ) ) ) ) ) ) ) |
| Raet1e | ENSMUSG00000053219 | ENSMUST00000065527 | 49 | . . ) ) ) ) . ) |
| Ralbp1 | ENSMUSG00000024096 | ENSMUST00000024905 | 277 | ) ) ) ) . ) ) ) |
| Ralbp1 | ENSMUSG00000024096 | ENSMUST00000166543 | 601 | ) ) . ) ) ) ) ) |
| Ralgapa1 | ENSMUSG00000021027 | ENSMUST00000219432 | 261 | ) . . . ) ) ) ) |
| Ralgapa1 | ENSMUSG00000021027 | ENSMUST00000085385 | 331 | ) . . . ) ) ) ) |
| Raly | ENSMUSG00000027593 | ENSMUST00000116389 | 360 | ) ) . . . . . ) |
| Raly | ENSMUSG00000027593 | ENSMUST00000109701 | 374 | ) ) . . . . . ) |
| Ralyl | ENSMUSG00000039717 | ENSMUST00000108372 | 74 | ) ) ) ) ) ) ) ) |
| Ralyl | ENSMUSG00000039717 | ENSMUST00000192209 | 92 | . ) ) ) ) ) ) . |
| Rap2c | ENSMUSG00000050029 | ENSMUST00000053593 | 77 | . . . . . . . . |
| Rasa3 | ENSMUSG00000031453 | ENSMUST00000117551 | 81 | ) ) . ) . . ) ) |
| Rasal2 | ENSMUSG00000070565 | ENSMUST00000078308 | 598 | ) ) ) ) ) ) ) ) |
| Rassf2 | ENSMUSG00000027339 | ENSMUST00000028814 | 254 | ( ( ( . ( ( ( ( |
| Rassf9 | ENSMUSG00000044921 | ENSMUST00000219445 | 223 | ) ) . ( ( ( ( ( |
| Rbfox1 | ENSMUSG00000008658 | ENSMUST00000056416 | 97 | . . . . . ) ) . |
| Rbm41 | ENSMUSG00000031433 | ENSMUST00000033810 | 0 | . ( ( ( ( ( ( ( |
| Rbm41 | ENSMUSG00000031433 | ENSMUST00000113011 | 6 | ( ( ( . . ( ( ( |
| Rbm41 | ENSMUSG00000031433 | ENSMUST00000087400 | 44 | ( ( ( . . ( ( ( |
| Rbm48 | ENSMUSG00000040302 | ENSMUST00000042753 | 292 | ( . . ( ( ( ( . |

| | | | | |
|---|---|---|---|---|
| Rbms1 | ENSMUSG00000026970 | ENSMUST00000028347 | 413 | . ) ) ) ) ) ) . |
| Rbms2 | ENSMUSG00000040043 | ENSMUST00000092033 | 25 | . ( ( ( ( ( ( ( |
| Rc3h1 | ENSMUSG00000040423 | ENSMUST00000161609 | 342 | ) ) ) ) ) ) . . |
| Rcor3 | ENSMUSG00000037395 | ENSMUST00000110849 | 76 | . . . . . . . . |
| Rcor3 | ENSMUSG00000037395 | ENSMUST00000192491 | 92 | . . . . . . . . |
| Rcor3 | ENSMUSG00000037395 | ENSMUST00000073279 | 106 | . . . . . ) ) ) |
| Rcor3 | ENSMUSG00000037395 | ENSMUST00000192866 | 128 | . . . . . . . . |
| Rdh13 | ENSMUSG00000008435 | ENSMUST00000008579 | 120 | ) ) . . . . . ) |
| Rer1 | ENSMUSG00000029048 | ENSMUST00000030914 | 116 | ( ( ( ( ( ( ( ( |
| Retnlb | ENSMUSG00000022650 | ENSMUST00000023328 | 185 | ) ) ) ) ) ) ) . |
| Rfx3 | ENSMUSG00000040929 | ENSMUST00000174850 | 91 | . ( ( ( ( ( . . |
| Rfx4 | ENSMUSG00000020037 | ENSMUST00000060397 | 323 | . . . . . . . . |
| Rfx5 | ENSMUSG00000005774 | ENSMUST00000137088 | 91 | . . . . . . . . |
| Rfx6 | ENSMUSG00000019900 | ENSMUST00000050455 | 395 | ) ) ) . ) ) ) ) |
| Rfx7 | ENSMUSG00000037674 | ENSMUST00000163401 | 524 | ) ) ) ) ) ) ) ) |
| Rgl1 | ENSMUSG00000026482 | ENSMUST00000111859 | 420 | ) ) . . . . . . |
| Rgn | ENSMUSG00000023070 | ENSMUST00000023832 | 88 | ) ) . . . . ) ) |
| Rgs9 | ENSMUSG00000020599 | ENSMUST00000106706 | 21 | ) ) ) ) . ( ( ( |
| Rgs9 | ENSMUSG00000020599 | ENSMUST00000020920 | 84 | ) ) ) ) . ) ) ) |
| Rhebl1 | ENSMUSG00000023755 | ENSMUST00000024518 | 109 | ( ( . . . . . . |
| Rhoh | ENSMUSG00000029204 | ENSMUST00000201533 | 1160 | . . . . . . ) ) |
| Rhox13 | ENSMUSG00000050197 | ENSMUST00000152730 | 21 | ( . . . . . . . |
| Rhox3g | ENSMUSG00000080933 | ENSMUST00000119965 | 226 | . . ) ) ) ) ) ) |
| Ribc2 | ENSMUSG00000022431 | ENSMUST00000023067 | 125 | . . . . . . ) ) |
| Riiad1 | ENSMUSG00000028139 | ENSMUST00000029785 | 32 | ( ( . ( ( ( ( . |
| Riok3 | ENSMUSG00000024404 | ENSMUST00000025270 | 49 | ( . . . . . . . |
| Riox1 | ENSMUSG00000046791 | ENSMUST00000053744 | 27 | . . ) . ) ) ) ) |
| Rit2 | ENSMUSG00000057455 | ENSMUST00000153060 | 161 | . . . . . . . . |
| Rnase11 | ENSMUSG00000059648 | ENSMUST00000076106 | 21 | . . . . . . . . |
| Rnf103 | ENSMUSG00000052656 | ENSMUST00000064637 | 716 | ) ) . ) ) . . ) |
| Rnf11 | ENSMUSG00000028557 | ENSMUST00000030284 | 398 | . . . . . . . . |
| Rnf133 | ENSMUSG00000051956 | ENSMUST00000063548 | 157 | . . . . . . . . |
| Rnf14 | ENSMUSG00000060450 | ENSMUST00000072376 | 310 | . . . . . . . . |
| Rnf14 | ENSMUSG00000060450 | ENSMUST00000171461 | 623 | . . . . . ) ) ) |
| Rnf149 | ENSMUSG00000048234 | ENSMUST00000062525 | 101 | . . . ) ) ) ) ) |
| Rnf168 | ENSMUSG00000014074 | ENSMUST00000171474 | 608 | ) ) . . . . ) ) |
| Rnf181 | ENSMUSG00000055850 | ENSMUST00000154098 | 184 | ) . . ) ) . ) ) |
| Rnf208 | ENSMUSG00000044628 | ENSMUST00000114355 | 298 | . . . ) ) ) ) ) |
| Rnf220 | ENSMUSG00000028677 | ENSMUST00000030439 | 432 | ( . . . . . . . |
| Rnf38 | ENSMUSG00000035696 | ENSMUST00000045793 | 237 | . . ) ) ) ) ) . |
| Rnf38 | ENSMUSG00000035696 | ENSMUST00000098098 | 263 | ) ) ) ) . ) ) ) |
| Rnf4 | ENSMUSG00000029110 | ENSMUST00000030992 | 222 | ) . . . . ) ) ) |
| Rnf4 | ENSMUSG00000029110 | ENSMUST00000182047 | 283 | ) . . . . ) ) ) |
| Rnf4 | ENSMUSG00000029110 | ENSMUST00000182709 | 370 | ) . . . . ) ) ) |
| Rnh1 | ENSMUSG00000038650 | ENSMUST00000167493 | 51 | . . . . . . . . |
| Robo2 | ENSMUSG00000052516 | ENSMUST00000117785 | 59 | ) ) ) . . . . . |
| Robo4 | ENSMUSG00000032125 | ENSMUST00000214185 | 109 | ) ) ( ( ( . . . |
| Rorb | ENSMUSG00000036192 | ENSMUST00000112832 | 547 | . . . . ( ( ( ( |
| Rorc | ENSMUSG00000028150 | ENSMUST00000029795 | 12 | . . . . . . . . |
| Rp1 | ENSMUSG00000025900 | ENSMUST00000027032 | 82 | . . ( ( ( ( ( ( |
| Rpgr | ENSMUSG00000031174 | ENSMUST00000044598 | 128 | . . . . . . . . |
| Rpgr | ENSMUSG00000031174 | ENSMUST00000073392 | 128 | . . . . . . . . |
| Rpgr | ENSMUSG00000031174 | ENSMUST00000072393 | 128 | . . . . . . . . |
| Rpgr | ENSMUSG00000031174 | ENSMUST00000115532 | 128 | . . . . . . . . |
| Rpl10 | ENSMUSG00000008682 | ENSMUST00000008826 | 46 | ) ) ) ) . . ) ) |
| Rpl10a | ENSMUSG00000037805 | ENSMUST00000042334 | 113 | . . . . . . . . |
| Rpl11 | ENSMUSG00000059291 | ENSMUST00000102536 | 21 | . . . ) ) ) ) ) |
| Rpl13 | ENSMUSG00000000740 | ENSMUST00000000756 | 63 | ) ) . ) ) ) ) ) |

| | | | | |
|---|---|---|---|---|
| Rpl13a | ENSMUSG00000074129 | ENSMUST00000150350 | 23 | . . ) ) ) ) ) . |
| Rpl15 | ENSMUSG00000012405 | ENSMUST00000080281 | 83 | ) ) ) ) ) ) ) ) |
| Rpl17 | ENSMUSG00000062328 | ENSMUST00000079716 | 44 | . ) ) ) . . . . |
| Rpl18 | ENSMUSG00000059070 | ENSMUST00000072503 | 1 | . . . . . . . . |
| Rpl19 | ENSMUSG00000017404 | ENSMUST00000017548 | 36 | . . . . . . . . |
| Rpl21 | ENSMUSG00000041453 | ENSMUST00000035983 | 252 | . . . . . . . . |
| Rpl22l1 | ENSMUSG00000039221 | ENSMUST00000194649 | 41 | . . . . . . . . |
| Rpl22l1 | ENSMUSG00000039221 | ENSMUST00000043867 | 74 | . . . . . . . . |
| Rpl23 | ENSMUSG00000071415 | ENSMUST00000103146 | 62 | ( ( ( . . . . . |
| Rpl23a | ENSMUSG00000058546 | ENSMUST00000102483 | 26 | . . . . . . . . |
| Rpl24 | ENSMUSG00000098274 | ENSMUST00000023269 | 111 | ) ) . . . ) ) ) |
| Rpl26 | ENSMUSG00000060938 | ENSMUST00000073471 | 6 | . . . . . . . . |
| Rpl27 | ENSMUSG00000063316 | ENSMUST00000077856 | 125 | . ) ) ) ) . ) ) |
| Rpl27a | ENSMUSG00000046364 | ENSMUST00000143107 | 55 | . . . . . . . . |
| Rpl28 | ENSMUSG00000030432 | ENSMUST00000032597 | 91 | . . . . . ( ( ( |
| Rpl29 | ENSMUSG00000048758 | ENSMUST00000150576 | 83 | . . . . . . . . |
| Rpl30 | ENSMUSG00000058600 | ENSMUST00000079735 | 416 | . . . . . . . . |
| Rpl31 | ENSMUSG00000073702 | ENSMUST00000086535 | 68 | . . . . . . . . |
| Rpl32 | ENSMUSG00000057841 | ENSMUST00000081840 | 4 | . . . . . . . . |
| Rpl34 | ENSMUSG00000062006 | ENSMUST00000079085 | 56 | . . . . . . . . |
| Rpl35 | ENSMUSG00000062997 | ENSMUST00000080861 | 492 | ) ) ) ) . . ) ) |
| Rpl35a | ENSMUSG00000060636 | ENSMUST00000078804 | 29 | ( . ( . . . . ( |
| Rpl36a | ENSMUSG00000079435 | ENSMUST00000113211 | 32 | ( . . . . . . . |
| Rpl36al | ENSMUSG00000049751 | ENSMUST00000054544 | 45 | . . ) ) . . ) ) |
| Rpl38 | ENSMUSG00000057322 | ENSMUST00000077915 | 30 | . . . ) ) ) ) ) |
| Rpl38 | ENSMUSG00000057322 | ENSMUST00000106602 | 54 | . . . . . . . . |
| Rpl3l | ENSMUSG00000002500 | ENSMUST00000170239 | 37 | . . . . . . ) ) |
| Rpl5 | ENSMUSG00000058558 | ENSMUST00000082223 | 70 | . . . . . . . ( |
| Rpl6 | ENSMUSG00000029614 | ENSMUST00000031617 | 53 | . . . ) ) ) ) ) |
| Rpl7 | ENSMUSG00000043716 | ENSMUST00000058437 | 266 | . . . . ( ( ( ( |
| Rpl7a | ENSMUSG00000062647 | ENSMUST00000102898 | 39 | . . . . . . . . |
| Rpl8 | ENSMUSG00000003970 | ENSMUST00000004072 | 13 | . . ( ( ( . ( ( |
| Rplp2 | ENSMUSG00000025508 | ENSMUST00000084434 | 278 | . . . . . . . . |
| Rps10 | ENSMUSG00000052146 | ENSMUST00000114882 | 34 | ) ) ) . . . . ( |
| Rps11 | ENSMUSG00000003429 | ENSMUST00000003521 | 104 | ) ) ) ) . . . . |
| Rps12 | ENSMUSG00000061983 | ENSMUST00000218107 | 74 | ) . . . . . . . |
| Rps14 | ENSMUSG00000024608 | ENSMUST00000025511 | 141 | . . . . . . . . |
| Rps15 | ENSMUSG00000063457 | ENSMUST00000068408 | 6 | . . . . . . . . |
| Rps15a | ENSMUSG00000008683 | ENSMUST00000131374 | 27 | . . . . . . . . |
| Rps16 | ENSMUSG00000037563 | ENSMUST00000082134 | 98 | ) ) . . . . ) ) |
| Rps18 | ENSMUSG00000008668 | ENSMUST00000008812 | 53 | . . . . . . . ( |
| Rps2 | ENSMUSG00000044533 | ENSMUST00000170715 | 26 | ) . . . ) ) ) ) |
| Rps21 | ENSMUSG00000039001 | ENSMUST00000059080 | 37 | . . . . . ) ) ) |
| Rps23 | ENSMUSG00000049517 | ENSMUST00000051955 | 206 | . . . . . . . . |
| Rps24 | ENSMUSG00000025290 | ENSMUST00000169826 | 1 | . . . . . . . . |
| Rps24 | ENSMUSG00000025290 | ENSMUST00000225023 | 1 | . . . . . . . . |
| Rps25 | ENSMUSG00000009927 | ENSMUST00000080300 | 559 | . . . . . . . . |
| Rps26 | ENSMUSG00000025362 | ENSMUST00000026420 | 245 | ) ) . ) ) ) . ) |
| Rps27 | ENSMUSG00000090733 | ENSMUST00000170122 | 2 | ( ( . . . . . . |
| Rps27a | ENSMUSG00000020460 | ENSMUST00000102845 | 246 | . . . . . ) ) ) |
| Rps29 | ENSMUSG00000034892 | ENSMUST00000037023 | 95 | ( . ( ( . . . . |
| Rps3 | ENSMUSG00000030744 | ENSMUST00000032998 | 90 | . . . . . . . . |
| Rps4x | ENSMUSG00000031320 | ENSMUST00000033683 | 71 | ) . . . . . . . |
| Rps5 | ENSMUSG00000012848 | ENSMUST00000004554 | 27 | ( ( . . . . . ( |
| Rps6 | ENSMUSG00000028495 | ENSMUST00000102814 | 46 | ( ( . . . . . . |
| Rps7 | ENSMUSG00000061477 | ENSMUST00000074267 | 71 | . ) ) ) ) ) . . |
| Rps8 | ENSMUSG00000047675 | ENSMUST00000102696 | 159 | ) ) ) ) ) ) . ) |
| Rsad1 | ENSMUSG00000039096 | ENSMUST00000040487 | 58 | . . . . . . . . |

| | | | | |
|---|---|---|---|---|
| Rsph6a | ENSMUSG00000040866 | ENSMUST00000076887 | 33 | ) ) ) ) . ) ) ) |
| Rsph6a | ENSMUSG00000040866 | ENSMUST00000035521 | 42 | ) ) ) . . . ) ) |
| Rspo2 | ENSMUSG00000051920 | ENSMUST00000063492 | 810 | ) ) ) ) ) . . . |
| Rtcb | ENSMUSG00000001783 | ENSMUST00000001834 | 69 | . . . . . . . . |
| Rtl1 | ENSMUSG00000085925 | ENSMUST00000149046 | 129 | . . . . . . . . |
| Rtl1 | ENSMUSG00000098639 | ENSMUST00000184423 | 129 | . . . . . . . . |
| Rtn2 | ENSMUSG00000030401 | ENSMUST00000108468 | 18 | ( . . . . . ( . |
| Rtn4ip1 | ENSMUSG00000019864 | ENSMUST00000054418 | 420 | ( ( . . . . . . |
| Rwdd2b | ENSMUSG00000041079 | ENSMUST00000039101 | 54 | . . . . . . . . |
| Rxfp2 | ENSMUSG00000053368 | ENSMUST00000201612 | 116 | ) . . . . . . . |
| Rxfp2 | ENSMUSG00000053368 | ENSMUST00000065745 | 147 | ) ) . . . . ) ) |
| Rxfp2 | ENSMUSG00000053368 | ENSMUST00000110496 | 147 | ) ) . . . . ) ) |
| Rxrb | ENSMUSG00000039656 | ENSMUST00000116612 | 116 | ( ( ( ( ( . . . |
| Rxrb | ENSMUSG00000039656 | ENSMUST00000173354 | 142 | ( ( ( ( ( . . . |
| S100pbp | ENSMUSG00000040928 | ENSMUST00000106061 | 233 | ) . ) ) ) ) ) ) |
| S100pbp | ENSMUSG00000040928 | ENSMUST00000106059 | 248 | ) ) ) . ) ) ) ) |
| Samd5 | ENSMUSG00000060487 | ENSMUST00000100070 | 159 | . . . . . . . . |
| Samd9l | ENSMUSG00000047735 | ENSMUST00000120087 | 256 | ) ) ) ) ) ) ) ) |
| Sarm1 | ENSMUSG00000050132 | ENSMUST00000061174 | 214 | . . . . . . . . |
| Scaf8 | ENSMUSG00000046201 | ENSMUST00000076734 | 377 | ) ) ) ) ) ) ) ) |
| Scarb1 | ENSMUSG00000037936 | ENSMUST00000111390 | 169 | ) ) ) . . ( ( ( |
| Scarb1 | ENSMUSG00000037936 | ENSMUST00000086075 | 192 | ) ) ) . . ( ( ( |
| Scg3 | ENSMUSG00000032181 | ENSMUST00000213324 | 256 | ( ( ( ( ( ( . . |
| Scg3 | ENSMUSG00000032181 | ENSMUST00000034699 | 297 | ( ( ( ( ( ( . . |
| Schip1 | ENSMUSG00000027777 | ENSMUST00000029346 | 72 | . . . . . ) ) ) |
| Schip1 | ENSMUSG00000027777 | ENSMUST00000169909 | 72 | . . . . . ) ) ) |
| Sclt1 | ENSMUSG00000059834 | ENSMUST00000026866 | 426 | ) ) ) ) ) . . . |
| Scn2b | ENSMUSG00000070304 | ENSMUST00000093855 | 100 | ) ) . . . . . ) |
| Scn5a | ENSMUSG00000032511 | ENSMUST00000065196 | 1 | . . . ( ( ( ( ( |
| Scn5a | ENSMUSG00000032511 | ENSMUST00000117911 | 169 | ) ) . ( ( ( ( ( |
| Scoc | ENSMUSG00000063253 | ENSMUST00000212031 | 58 | ( ( . . . . . . |
| Scoc | ENSMUSG00000063253 | ENSMUST00000167525 | 60 | ( ( . . . . . . |
| Scrn3 | ENSMUSG00000008226 | ENSMUST00000090811 | 47 | ( . . . . ( ( ( |
| Sdc2 | ENSMUSG00000022261 | ENSMUST00000022871 | 451 | . ) ) ) ) ) ) ) |
| Sdf2 | ENSMUSG00000002064 | ENSMUST00000002133 | 184 | ) . . . ) ) ) ) |
| Sdf4 | ENSMUSG00000029076 | ENSMUST00000097734 | 76 | ) ) . ( ( ( ( ( |
| Sdf4 | ENSMUSG00000029076 | ENSMUST00000050078 | 335 | ( ( ( ( . ( ( . |
| Sdr16c6 | ENSMUSG00000071019 | ENSMUST00000108383 | 56 | . ( ( ( . . ( ( |
| Sdr9c7 | ENSMUSG00000040127 | ENSMUST00000047134 | 34 | . . . . . . . . |
| Sec24b | ENSMUSG00000001052 | ENSMUST00000001079 | 181 | ) . ) ) ) ) . ( |
| Sec61b | ENSMUSG00000053317 | ENSMUST00000065678 | 52 | . . . . . . . . |
| Selenon | ENSMUSG00000050989 | ENSMUST00000060435 | 7 | ( ( . . . ( ( ( |
| Sema3c | ENSMUSG00000028780 | ENSMUST00000030568 | 670 | ) ) ) ) . ) ) . |
| Sema3g | ENSMUSG00000021904 | ENSMUST00000090180 | 39 | . . . . . . . . |
| Sema6b | ENSMUSG00000001227 | ENSMUST00000167545 | 231 | ) ) ) ) . ) ) ) |
| Senp7 | ENSMUSG00000052917 | ENSMUST00000049128 | 56 | ( ( . . . . . . |
| Serpina3a | ENSMUSG00000041536 | ENSMUST00000109965 | 55 | . ( ( ( ( . . ( |
| Serpina9 | ENSMUSG00000058260 | ENSMUST00000058464 | 56 | ) ) ) ) ) ) ) ) |
| Serpinb1b | ENSMUSG00000051029 | ENSMUST00000016951 | 45 | . . . . ) ) ) ) |
| Serpinb1c | ENSMUSG00000079049 | ENSMUST00000021834 | 50 | . . . . ( ( . . |
| Serpinb5 | ENSMUSG00000067006 | ENSMUST00000086701 | 39 | ) ) . . . . . . |
| Serpinb6d | ENSMUSG00000047889 | ENSMUST00000059637 | 171 | . . . . . ) ) ) |
| Set | ENSMUSG00000054766 | ENSMUST00000102866 | 258 | ( . . . . . ( ( |
| Setbp1 | ENSMUSG00000024548 | ENSMUST00000025430 | 326 | . . . . . . . . |
| Sf3a2 | ENSMUSG00000020211 | ENSMUST00000148665 | 609 | ( ( ( ( . . . . |
| Sgf29 | ENSMUSG00000030714 | ENSMUST00000032956 | 153 | . . . . . . . . |
| Sgk1 | ENSMUSG00000019970 | ENSMUST00000020145 | 63 | ) ) ) ) . . ) ) |
| Sgk1 | ENSMUSG00000019970 | ENSMUST00000100036 | 81 | ( ( ( ( . . . . |

| | | | | |
|---|---|---|---|---|
| Sh3bgr | ENSMUSG00000040666 | ENSMUST00000048770 | 216 | . . . . . . . . |
| Sh3bgrl | ENSMUSG00000031246 | ENSMUST00000033598 | 182 | ) ) ) ) . . . . |
| Sh3bp5 | ENSMUSG00000021892 | ENSMUST00000100730 | 41 | ( ( . . . . . . |
| Sh3bp5l | ENSMUSG00000013646 | ENSMUST00000116376 | 491 | ) ) ) . . ) ) ) |
| Sh3yl1 | ENSMUSG00000020669 | ENSMUST00000110880 | 481 | ( ( ( ( ( ( ( . |
| Sh3yl1 | ENSMUSG00000020669 | ENSMUST00000020997 | 505 | ( ( ( ( ( ( ( ( |
| Shisa4 | ENSMUSG00000041889 | ENSMUST00000041240 | 234 | ) ) . . . ) ) ) |
| Sirpb1a | ENSMUSG00000095788 | ENSMUST00000192700 | 39 | ) ) ) . . . . . |
| Sirpb1b | ENSMUSG00000095028 | ENSMUST00000091319 | 39 | ) ) . . . ) ) ) |
| Sirpb1c | ENSMUSG00000074677 | ENSMUST00000050623 | 39 | ) ) . . . ) ) ) |
| Sirt4 | ENSMUSG00000029524 | ENSMUST00000112066 | 304 | . . . . . . . . |
| Sirt6 | ENSMUSG00000034748 | ENSMUST00000042923 | 171 | . ) ) ) ) ) ) ) |
| Six3 | ENSMUSG00000038805 | ENSMUST00000176081 | 379 | . . . . . . . . |
| Skil | ENSMUSG00000027660 | ENSMUST00000029194 | 649 | . ( ( ( . . ( ( |
| Skil | ENSMUSG00000027660 | ENSMUST00000118470 | 652 | . ( ( ( . . ( ( |
| Slain1 | ENSMUSG00000055717 | ENSMUST00000069443 | 27 | ) ) ) . ( ( ( ( |
| Slain2 | ENSMUSG00000036087 | ENSMUST00000143829 | 191 | ) ) ) ) ) ) ) ) |
| Slain2 | ENSMUSG00000036087 | ENSMUST00000144843 | 256 | ) ) ) ) ) ) ) ) |
| Slamf1 | ENSMUSG00000015316 | ENSMUST00000015460 | 8 | ( ( ( ( ( . . . |
| Slamf8 | ENSMUSG00000053318 | ENSMUST00000065679 | 59 | . . . . . ) ) ) |
| Slc13a2 | ENSMUSG00000001095 | ENSMUST00000001122 | 0 | . . . . . . . . |
| Slc13a5 | ENSMUSG00000020805 | ENSMUST00000021161 | 81 | . . . . ) ) ) ) |
| Slc15a2 | ENSMUSG00000022899 | ENSMUST00000164579 | 99 | . . . . . ) ) ) |
| Slc15a2 | ENSMUSG00000022899 | ENSMUST00000023616 | 121 | . . . . . ) ) ) |
| Slc16a11 | ENSMUSG00000040938 | ENSMUST00000171032 | 321 | . ( ( ( ( ( ( ( |
| Slc16a13 | ENSMUSG00000044367 | ENSMUST00000060010 | 306 | ) ) ) ) ) ) ) . |
| Slc16a5 | ENSMUSG00000045775 | ENSMUST00000092445 | 71 | ) ) ) ) ) ) . ( |
| Slc16a6 | ENSMUSG00000041920 | ENSMUST00000070872 | 338 | . . . . . ) ) ) |
| Slc23a1 | ENSMUSG00000024354 | ENSMUST00000025212 | 66 | ( . . . . . . . |
| Slc24a4 | ENSMUSG00000041771 | ENSMUST00000079020 | 187 | . . . . . ) ) ) |
| Slc24a5 | ENSMUSG00000035183 | ENSMUST00000070353 | 18 | ( ( . . . . . . |
| Slc25a15 | ENSMUSG00000031482 | ENSMUST00000033871 | 173 | ) ) ) ) ) ) ) ) |
| Slc25a53 | ENSMUSG00000044348 | ENSMUST00000171738 | 67 | . . . . . . ) ) |
| Slc26a3 | ENSMUSG00000001225 | ENSMUST00000001254 | 175 | . . . ) ) ) ) . |
| Slc2a4 | ENSMUSG00000018566 | ENSMUST00000018710 | 121 | . . . . . . . . |
| Slc2a9 | ENSMUSG00000005107 | ENSMUST00000067886 | 97 | ) . . ( ( ( ( ( |
| Slc30a3 | ENSMUSG00000029151 | ENSMUST00000202740 | 252 | . ( ( ( . . . . |
| Slc31a1 | ENSMUSG00000066150 | ENSMUST00000084526 | 136 | . . . ) ) ) . . |
| Slc32a1 | ENSMUSG00000037771 | ENSMUST00000045738 | 449 | . . . . . . ) ) |
| Slc35e2 | ENSMUSG00000042202 | ENSMUST00000105608 | 391 | . . . . . . . . |
| Slc41a1 | ENSMUSG00000013275 | ENSMUST00000086559 | 955 | . ) ) ) ) ) ) ) |
| Slc43a1 | ENSMUSG00000027075 | ENSMUST00000028469 | 75 | ( . ( ( ( ( . . |
| Slc43a1 | ENSMUSG00000027075 | ENSMUST00000111624 | 209 | ) ) ) ) ) ) ) ) |
| Slc4a10 | ENSMUSG00000026904 | ENSMUST00000054484 | 36 | ) ) ) ) ) ) ) ) |
| Slc4a10 | ENSMUSG00000026904 | ENSMUST00000102735 | 159 | ) ) ) ) ) ) ) ) |
| Slc4a11 | ENSMUSG00000074796 | ENSMUST00000099362 | 245 | . . . ) ) ) ) . |
| Slc5a12 | ENSMUSG00000041644 | ENSMUST00000111026 | 207 | . ) ) ) . ) ) ) |
| Slc5a12 | ENSMUSG00000041644 | ENSMUST00000045972 | 216 | . ) ) ) . ) ) ) |
| Slc6a13 | ENSMUSG00000030108 | ENSMUST00000064580 | 43 | ) ) ) . . . ) ) |
| Slc6a15 | ENSMUSG00000019894 | ENSMUST00000179636 | 287 | ( ( ( . . . . . |
| Slc6a15 | ENSMUSG00000019894 | ENSMUST00000074204 | 384 | ) ) ) ) ) ) ) ) |
| Slc6a2 | ENSMUSG00000055368 | ENSMUST00000072939 | 246 | . . . . . . . . |
| Slc7a11 | ENSMUSG00000027737 | ENSMUST00000029297 | 315 | . . . . . . . . |
| Slc7a15 | ENSMUSG00000020600 | ENSMUST00000036938 | 279 | . . . . ( ( ( . |
| Slc7a2 | ENSMUSG00000031596 | ENSMUST00000057784 | 151 | . . . . . . . . |
| Slc7a2 | ENSMUSG00000031596 | ENSMUST00000098816 | 167 | . . . . . . . . |
| Slc7a8 | ENSMUSG00000022180 | ENSMUST00000022787 | 564 | ) ) ) ) ) ) ) ) |
| Slc8a2 | ENSMUSG00000030376 | ENSMUST00000168693 | 6 | . . . . . . . . |

| | | | | |
|---|---|---|---|---|
| Slc8a2 | ENSMUSG00000030376 | ENSMUST00000211649 | 528 | . . . . . . . . |
| Slitrk6 | ENSMUSG00000045871 | ENSMUST00000078386 | 471 | . . . . . ) ) ) |
| Slx1b | ENSMUSG00000059772 | ENSMUST00000144897 | 115 | ) ) ( ( ( ( ( . |
| Smad9 | ENSMUSG00000027796 | ENSMUST00000029371 | 283 | . . . . . . . . |
| Smarca1 | ENSMUSG00000031099 | ENSMUST00000077569 | 83 | . . . . ) ) ) ) |
| Smarca1 | ENSMUSG00000031099 | ENSMUST00000101616 | 83 | . . . . ) ) ) ) |
| Smarcb1 | ENSMUSG00000000902 | ENSMUST00000121304 | 177 | . . . . . ) ) ) |
| Smarcb1 | ENSMUSG00000000902 | ENSMUST00000000925 | 180 | . . . . . . . ( |
| Smchd1 | ENSMUSG00000024054 | ENSMUST00000127430 | 162 | ) . . . . . . . |
| Smco1 | ENSMUSG00000046345 | ENSMUST00000093183 | 122 | ) . . . . . . . |
| Smim19 | ENSMUSG00000031534 | ENSMUST00000033935 | 373 | . ) ) ) ) . ) ) |
| Smim20 | ENSMUSG00000061461 | ENSMUST00000147148 | 93 | ) ) ) ) . . . . |
| Smim3 | ENSMUSG00000038059 | ENSMUST00000042710 | 513 | ( ( . . . . . . |
| Smim6 | ENSMUSG00000075420 | ENSMUST00000132961 | 346 | . . . ) ) ) ) ) |
| Smug1 | ENSMUSG00000036061 | ENSMUST00000064067 | 74 | . . . . . . . . |
| Snca | ENSMUSG00000025889 | ENSMUST00000114268 | 213 | ) ) ) ) . ) ) ) |
| Snhg11 | ENSMUSG00000044349 | ENSMUST00000109488 | 345 | ( ( ( ( ( . . . |
| Snrpa | ENSMUSG00000061479 | ENSMUST00000163311 | 185 | ) . . . . . ) ) |
| Snrpn | ENSMUSG00000102252 | ENSMUST00000059305 | 397 | ( ( ( . . . . . |
| Snrpn | ENSMUSG00000102252 | ENSMUST00000098402 | 548 | . . . . . . . . |
| Sntb1 | ENSMUSG00000060429 | ENSMUST00000039769 | 259 | ) ) ) . . . . ) |
| Snu13 | ENSMUSG00000063480 | ENSMUST00000080622 | 32 | ( . . ( ( ( ( ( |
| Snx33 | ENSMUSG00000032733 | ENSMUST00000050916 | 1553 | . . . . . . . . |
| Snx5 | ENSMUSG00000027423 | ENSMUST00000028909 | 97 | ) . ) ) . . ) ) |
| Sod1 | ENSMUSG00000022982 | ENSMUST00000023707 | 25 | ) ) ) ) ( ( ( ( |
| Sox5 | ENSMUSG00000041540 | ENSMUST00000111748 | 43 | ) . ) ) ) ) ) ) |
| Spa17 | ENSMUSG00000001948 | ENSMUST00000002013 | 182 | ) ) ) ) ) ) ) . |
| Spata7 | ENSMUSG00000021007 | ENSMUST00000101146 | 31 | ( ( ( ( ( ( . . |
| Spata7 | ENSMUSG00000021007 | ENSMUST00000101144 | 61 | . . . . . . . . |
| Spata7 | ENSMUSG00000021007 | ENSMUST00000048402 | 71 | ( ( ( ( ( ( . . |
| Spatc1 | ENSMUSG00000049653 | ENSMUST00000074173 | 68 | ( ( ( ( . . . . |
| Spats2 | ENSMUSG00000051934 | ENSMUST00000063517 | 545 | ) ) ) ) ) ) ) ) |
| Spc25 | ENSMUSG00000005233 | ENSMUST00000112320 | 236 | ( ( ( ( ( ( . . |
| Speg | ENSMUSG00000026207 | ENSMUST00000113589 | 190 | . . . . . . . . |
| Spesp1 | ENSMUSG00000046846 | ENSMUST00000056949 | 66 | . ( ( ( ( . . . |
| Sphk2 | ENSMUSG00000057342 | ENSMUST00000072836 | 577 | . . . . . ( ( ( |
| Spink14 | ENSMUSG00000051050 | ENSMUST00000060328 | 15 | . . . . . ( ( ( |
| Spink8 | ENSMUSG00000050074 | ENSMUST00000198988 | 196 | ) ) . ) ) ) . ) |
| Spns1 | ENSMUSG00000030741 | ENSMUST00000032994 | 318 | ( ( ( ( . . . . |
| Spred3 | ENSMUSG00000037239 | ENSMUST00000048923 | 62 | . . . . . . . . |
| Spsb4 | ENSMUSG00000046997 | ENSMUST00000055433 | 783 | ) ) ) . . . . . |
| Sptan1 | ENSMUSG00000057738 | ENSMUST00000100225 | 50 | . . . . . . . . |
| Sptan1 | ENSMUSG00000057738 | ENSMUST00000113717 | 170 | . . . . . . . . |
| Sptbn4 | ENSMUSG00000011751 | ENSMUST00000108362 | 180 | ) ) ) ) ) ) ) ) |
| Srprb | ENSMUSG00000032553 | ENSMUST00000035157 | 70 | . . . . . . . . |
| Srsf1 | ENSMUSG00000018379 | ENSMUST00000079866 | 72 | . . ) ) ) ) ) ) |
| Srsf1 | ENSMUSG00000098301 | ENSMUST00000193491 | 72 | . . ) ) ) ) ) ) |
| Srsf1 | ENSMUSG00000018379 | ENSMUST00000139129 | 417 | . . ) ) ) ) ) ) |
| Srsf1 | ENSMUSG00000098301 | ENSMUST00000183352 | 417 | . . ) ) ) ) ) ) |
| Ssbp1 | ENSMUSG00000029911 | ENSMUST00000121360 | 590 | ( ( ( ( . . . ( |
| Ssc5d | ENSMUSG00000035279 | ENSMUST00000057612 | 131 | . . . . . . . . |
| Ssr2 | ENSMUSG00000041355 | ENSMUST00000035785 | 197 | . . . . ) ) ) ) |
| Ssr4 | ENSMUSG00000002014 | ENSMUST00000002090 | 151 | ( ( ( ( ( ( ( ( |
| Ssr4 | ENSMUSG00000002014 | ENSMUST00000166518 | 185 | ( ( ( ( ( ( ( ( |
| Ssxb1 | ENSMUSG00000079705 | ENSMUST00000037297 | 107 | . ( ( ( ( ( . . |
| Ssxb10 | ENSMUSG00000068219 | ENSMUST00000103003 | 108 | . . . . ) ) ) ) |
| Ssxb2 | ENSMUSG00000023165 | ENSMUST00000103000 | 107 | ( ( ( ( . . . . |
| Ssxb9 | ENSMUSG00000068218 | ENSMUST00000115584 | 108 | ( ( ( ( . . . . |

| | | | | |
|---|---|---|---|---|
| St13 | ENSMUSG00000022403 | ENSMUST00000172107 | 360 | . . ( ( . . . . |
| St3gal1 | ENSMUSG00000013846 | ENSMUST00000092640 | 133 | . . . . . ) ) ) |
| St7 | ENSMUSG00000029534 | ENSMUST00000115417 | 328 | ) ) ) ) ) . . . |
| St7 | ENSMUSG00000029534 | ENSMUST00000053148 | 374 | . . . . . . . . |
| Stac2 | ENSMUSG00000017400 | ENSMUST00000017544 | 458 | . . . . . . . . |
| Stam | ENSMUSG00000026718 | ENSMUST00000102960 | 124 | ) ) ) . ) ) ) ) |
| Steap3 | ENSMUSG00000026389 | ENSMUST00000112641 | 389 | ) ) ) ) ( ( ( ( |
| Stk19 | ENSMUSG00000061207 | ENSMUST00000077477 | 40 | ( ( ( . . . . . |
| Stk3 | ENSMUSG00000022329 | ENSMUST00000018476 | 232 | . . . . . . . . |
| Stk32a | ENSMUSG00000039954 | ENSMUST00000045477 | 190 | ) ) ) ) ) ) ) . |
| Stk33 | ENSMUSG00000031027 | ENSMUST00000106745 | 455 | ( ( . ( ( ( ( . |
| Stk36 | ENSMUSG00000033276 | ENSMUST00000087183 | 220 | . . ( ( ( ( ( ( |
| Stkld1 | ENSMUSG00000049897 | ENSMUST00000055406 | 48 | ( ( ( ( . . . . |
| Stox2 | ENSMUSG00000038143 | ENSMUST00000211737 | 1305 | ) . . . ) ) ) ) |
| Stox2 | ENSMUSG00000038143 | ENSMUST00000079195 | 1345 | ) . . . ) ) ) ) |
| Stradb | ENSMUSG00000026027 | ENSMUST00000027185 | 391 | . . . . . ( ( ( |
| Stx12 | ENSMUSG00000028879 | ENSMUST00000030698 | 50 | ) ) . . . . . ( |
| Styx | ENSMUSG00000053205 | ENSMUST00000226873 | 305 | ) ) ) ) ) . ) ) |
| Sugt1 | ENSMUSG00000022024 | ENSMUST00000054908 | 64 | ) ) ) . . . ) ) |
| Sult1c2 | ENSMUSG00000023122 | ENSMUST00000023886 | 573 | ) . . ) ) ) ) ) |
| Sult1e1 | ENSMUSG00000029272 | ENSMUST00000031201 | 75 | . . . . ( ( ( ( |
| Susd1 | ENSMUSG00000038578 | ENSMUST00000040166 | 230 | ( ( . ( ( ( ( ( |
| Syce3 | ENSMUSG00000078938 | ENSMUST00000109314 | 8 | ) ) ) ) ) . . ( |
| Syngap1 | ENSMUSG00000067629 | ENSMUST00000194598 | 155 | . . . . . . . . |
| Sypl | ENSMUSG00000020570 | ENSMUST00000020885 | 91 | ( ( . . . . . . |
| Syt6 | ENSMUSG00000027849 | ENSMUST00000118563 | 370 | ( . . . . . . . |
| Syt8 | ENSMUSG00000031098 | ENSMUST00000122393 | 61 | ) ) ) ) ) ) ) ) |
| T | ENSMUSG00000062327 | ENSMUST00000074667 | 16 | ( ( ( ( ( ( ( ( |
| Taf1 | ENSMUSG00000031314 | ENSMUST00000149274 | 50 | ) ) ) ) . . ) ) |
| Taf1d | ENSMUSG00000031939 | ENSMUST00000034415 | 47 | . ( ( ( ( ( ( ( |
| Tagln | ENSMUSG00000032085 | ENSMUST00000034590 | 195 | . . . ( ( ( . . |
| Tagln3 | ENSMUSG00000022658 | ENSMUST00000096057 | 290 | ) ) ) ) . . . . |
| Tanc1 | ENSMUSG00000035168 | ENSMUST00000037526 | 224 | ) ) ) ) ) ) ) ) |
| Tanc1 | ENSMUSG00000035168 | ENSMUST00000112568 | 226 | ) ) ) ) ) ) ) ) |
| Tap1 | ENSMUSG00000037321 | ENSMUST00000170086 | 246 | . ( ( ( . . . . |
| Tardbp | ENSMUSG00000041459 | ENSMUST00000105699 | 27 | . . ) ) ) ) ) ) |
| Tardbp | ENSMUSG00000041459 | ENSMUST00000105702 | 265 | . ) ) ) ) ) ) ) |
| Tardbp | ENSMUSG00000041459 | ENSMUST00000172073 | 265 | . ) ) ) ) ) ) ) |
| Tardbp | ENSMUSG00000041459 | ENSMUST00000165113 | 265 | . ) ) ) ) ) ) ) |
| Tardbp | ENSMUSG00000041459 | ENSMUST00000084125 | 288 | . ) ) ) ) ) ) ) |
| Tas1r1 | ENSMUSG00000028950 | ENSMUST00000030792 | 84 | ( ( ( ( ( . . ( |
| Tasp1 | ENSMUSG00000039033 | ENSMUST00000099304 | 67 | ) ) ) ) ) ) ) ) |
| Tasp1 | ENSMUSG00000039033 | ENSMUST00000110079 | 87 | ) ) . ( ( ( ( ( |
| Tasp1 | ENSMUSG00000039033 | ENSMUST00000046656 | 309 | . . . . . ) ) ) |
| Tbc1d19 | ENSMUSG00000039178 | ENSMUST00000037337 | 81 | ( ( ( ( ( ( ( ( |
| Tbc1d21 | ENSMUSG00000036244 | ENSMUST00000040217 | 58 | ( ( ( ( ( ( ( ( |
| Tbc1d24 | ENSMUSG00000036473 | ENSMUST00000168378 | 249 | ) ) ) ) ) ) . ) |
| Tbc1d24 | ENSMUSG00000036473 | ENSMUST00000097376 | 271 | ) ) ) ) ) ) ) . |
| Tbc1d24 | ENSMUSG00000036473 | ENSMUST00000171189 | 421 | ) . . . . ) ) ) |
| Tbc1d8 | ENSMUSG00000003134 | ENSMUST00000054462 | 105 | . . ( ( ( . . . |
| Tbca | ENSMUSG00000042043 | ENSMUST00000046644 | 14 | ( . . . . . . . |
| Tbcd | ENSMUSG00000039230 | ENSMUST00000103013 | 50 | ) ) . . ) ) ) ) |
| Tbk1 | ENSMUSG00000020115 | ENSMUST00000020316 | 108 | ( ( ( ( . . . . |
| Tbpl2 | ENSMUSG00000061809 | ENSMUST00000080453 | 10 | . . . . . ( ( ( |
| Tbrg4 | ENSMUSG00000000384 | ENSMUST00000189268 | 209 | ) . ) ) . . . ) |
| Tbxas1 | ENSMUSG00000029925 | ENSMUST00000003017 | 129 | ) ) ) ) ) ) ) ) |
| Tcaf1 | ENSMUSG00000036667 | ENSMUST00000045054 | 56 | ( ( . . . . ( ( |
| Tcf4 | ENSMUSG00000053477 | ENSMUST00000114985 | 494 | ) ) . ) ) ) ) ) |

| | | | | |
|---|---|---|---|---|
| Tcf7l2 | ENSMUSG00000024985 | ENSMUST00000111659 | 249 | ( ( ( ( . ( ( ( |
| Tcf7l2 | ENSMUSG00000024985 | ENSMUST00000111658 | 423 | . . . . . . . . |
| Tcf7l2 | ENSMUSG00000024985 | ENSMUST00000041717 | 487 | . . ) ) ) ) ) . |
| Tcf7l2 | ENSMUSG00000024985 | ENSMUST00000061496 | 497 | . . ) ) ) ) ) . |
| Tcf7l2 | ENSMUSG00000024985 | ENSMUST00000111657 | 497 | . . ) ) ) ) ) . |
| Tchhl1 | ENSMUSG00000027908 | ENSMUST00000029516 | 55 | ) . . . . . . . |
| Tcn2 | ENSMUSG00000020432 | ENSMUST00000109990 | 79 | . ) ) ) ) . . ) |
| Tcn2 | ENSMUSG00000020432 | ENSMUST00000109991 | 108 | ) ) ) ) ) . . . |
| Tcn2 | ENSMUSG00000020432 | ENSMUST00000109989 | 114 | ) ) ) ) ) ) . . |
| Tcp11l1 | ENSMUSG00000027175 | ENSMUST00000028597 | 115 | ) ) ) . . . . ) |
| Tcp11l2 | ENSMUSG00000020034 | ENSMUST00000020223 | 453 | . . . . . . . . |
| Tctn1 | ENSMUSG00000038593 | ENSMUST00000111738 | 1 | . . . . . . . . |
| Tctn3 | ENSMUSG00000025008 | ENSMUST00000025981 | 15 | ) ) ) ) . . . . |
| Tdgf1 | ENSMUSG00000032494 | ENSMUST00000035075 | 281 | . ) ) ) ) . . . |
| Tdrd1 | ENSMUSG00000025081 | ENSMUST00000111604 | 317 | ) ) ) ) ) ) . . |
| Tdrd3 | ENSMUSG00000022019 | ENSMUST00000168275 | 87 | . . . . . . . . |
| Tdrd3 | ENSMUSG00000022019 | ENSMUST00000169504 | 87 | . . . . . . . . |
| Tekt4 | ENSMUSG00000024175 | ENSMUST00000025002 | 101 | . ( ( ( . . . ( |
| Tenm1 | ENSMUSG00000016150 | ENSMUST00000016294 | 61 | . ) ) ) ) ) ) . |
| Tesmin | ENSMUSG00000024905 | ENSMUST00000025840 | 0 | . . . . . . . . |
| Tet3 | ENSMUSG00000034832 | ENSMUST00000186548 | 507 | ( . . . . . . . |
| Tex10 | ENSMUSG00000028345 | ENSMUST00000164866 | 303 | . . . . . . ) ) |
| Tex22 | ENSMUSG00000012211 | ENSMUST00000012355 | 614 | ( ( ( ( ( ( . . |
| Tex33 | ENSMUSG00000062154 | ENSMUST00000169575 | 57 | . . ) ) ) . . . |
| Tex44 | ENSMUSG00000036574 | ENSMUST00000046004 | 6 | ( . ( ( ( . ( . |
| Tfap2a | ENSMUSG00000021359 | ENSMUST00000021787 | 337 | ) ) ) ) . ) ) ) |
| Tfap2b | ENSMUSG00000025927 | ENSMUST00000064976 | 333 | . . . . . ) ) ) |
| Tfap2d | ENSMUSG00000042596 | ENSMUST00000037294 | 172 | ) ) ) ) ) ) ) ) |
| Tfeb | ENSMUSG00000023990 | ENSMUST00000113288 | 216 | . . ) ) ) ) ) ) |
| Tfec | ENSMUSG00000029553 | ENSMUST00000031533 | 245 | ) . ) ) ) ) ) ) |
| Tfg | ENSMUSG00000022757 | ENSMUST00000065515 | 267 | . . . . . . ) ) |
| Tgfa | ENSMUSG00000029999 | ENSMUST00000032066 | 197 | . . . . . . . . |
| Tgfb1i1 | ENSMUSG00000030782 | ENSMUST00000070656 | 1266 | ) . . ) ) ) ) ) |
| Tgfb2 | ENSMUSG00000039239 | ENSMUST00000195201 | 908 | ) ) . . ) ) ) ) |
| Tgfb2 | ENSMUSG00000039239 | ENSMUST00000045288 | 1197 | . . . . . . . . |
| Tgfb3 | ENSMUSG00000021253 | ENSMUST00000003687 | 1040 | ( ( ( ( . . . . |
| Tgm7 | ENSMUSG00000079103 | ENSMUST00000110675 | 224 | ) ) ( ( . . . . |
| Thbs3 | ENSMUSG00000028047 | ENSMUST00000029682 | 12 | ( . . . ( ( ( . |
| Tial1 | ENSMUSG00000030846 | ENSMUST00000106226 | 445 | ) ) ) ) . . . . |
| Tiam1 | ENSMUSG00000002489 | ENSMUST00000164263 | 244 | ) ) ) ) ) ) ) ) |
| Timm29 | ENSMUSG00000048429 | ENSMUST00000062125 | 181 | ( ( . . . . . . |
| Tlcd1 | ENSMUSG00000019437 | ENSMUST00000092880 | 360 | ( . ( ( . ( ( ( |
| Tll1 | ENSMUSG00000053626 | ENSMUST00000066166 | 588 | ( ( ( ( ( . . . |
| Tlr8 | ENSMUSG00000040522 | ENSMUST00000112170 | 204 | ( ( . ( ( ( . . |
| Tlx2 | ENSMUSG00000068327 | ENSMUST00000089641 | 178 | . . . ) ) ) ) . |
| Tm4sf5 | ENSMUSG00000018919 | ENSMUST00000019063 | 35 | ( ( ( ( . . . . |
| Tm9sf2 | ENSMUSG00000025544 | ENSMUST00000026624 | 176 | . . . . . ) ) ) |
| Tmbim6 | ENSMUSG00000023010 | ENSMUST00000159209 | 99 | ) . ) ) ) ) ) . |
| Tmbim7 | ENSMUSG00000014529 | ENSMUST00000198739 | 147 | . ( ( ( ( ( ( . |
| Tmbim7 | ENSMUSG00000014529 | ENSMUST00000014673 | 323 | . ( ( ( ( ( ( . |
| Tmcc2 | ENSMUSG00000042066 | ENSMUST00000142609 | 78 | ( . . . . . . . |
| Tmco1 | ENSMUSG00000052428 | ENSMUST00000195015 | 412 | ) ) . . . . ) ) |
| Tmem101 | ENSMUSG00000020921 | ENSMUST00000021296 | 48 | . . . . . . . . |
| Tmem14a | ENSMUSG00000025933 | ENSMUST00000027064 | 184 | ) ) ) . . ) ) ) |
| Tmem14a | ENSMUSG00000025933 | ENSMUST00000027065 | 271 | ) ) ) . . ) ) ) |
| Tmem150a | ENSMUSG00000055912 | ENSMUST00000069695 | 152 | ) . . . ) ) ) ) |
| Tmem150a | ENSMUSG00000102455 | ENSMUST00000193693 | 152 | ) . . . ) ) ) ) |
| Tmem165 | ENSMUSG00000029234 | ENSMUST00000031144 | 226 | ) . . ) ) ) ) ) |

| | | | | |
|---|---|---|---|---|
| Tmem167 | ENSMUSG00000012422 | ENSMUST00000161568 | 603 | ) ) ) . ) ) . ) |
| Tmem171 | ENSMUSG00000052485 | ENSMUST00000064347 | 93 | . . . ) ) ) ) ) |
| Tmem175 | ENSMUSG00000013495 | ENSMUST00000078323 | 77 | . ) ) ) . . ) ) |
| Tmem175 | ENSMUSG00000013495 | ENSMUST00000063272 | 140 | . ) ) ) . . ) ) |
| Tmem182 | ENSMUSG00000079588 | ENSMUST00000114765 | 72 | . ) ) ) ) ) . . |
| Tmem198 | ENSMUSG00000051703 | ENSMUST00000113575 | 373 | ) . . . . . . . |
| Tmem211 | ENSMUSG00000066964 | ENSMUST00000086615 | 196 | . . . . . . . . |
| Tmem213 | ENSMUSG00000029829 | ENSMUST00000031851 | 72 | ) ) . . . . . . |
| Tmem258 | ENSMUSG00000036372 | ENSMUST00000040372 | 165 | . . . . . ) ) ) |
| Tmem29 | ENSMUSG00000041353 | ENSMUST00000059256 | 329 | ) ) ) ) ) ) ) ) |
| Tmem29 | ENSMUSG00000041353 | ENSMUST00000173996 | 360 | ) ) ) ) . ) ) ) |
| Tmem45a2 | ENSMUSG00000046748 | ENSMUST00000067173 | 113 | ( ( . . . . . . |
| Tmem74b | ENSMUSG00000044364 | ENSMUST00000060196 | 470 | ) ) ) ) ) . . . |
| Tmem79 | ENSMUSG00000001420 | ENSMUST00000001456 | 85 | ) ) ) ) . . . ( |
| Tmem8b | ENSMUSG00000078716 | ENSMUST00000167153 | 571 | ) ) ) ) ) . . . |
| Tmem94 | ENSMUSG00000020747 | ENSMUST00000093912 | 173 | . ) ) ) ) ) ) ) |
| Tmprss9 | ENSMUSG00000059406 | ENSMUST00000105333 | 49 | . . . . . . . ( |
| Tmsb4x | ENSMUSG00000049775 | ENSMUST00000112172 | 210 | . . . . . . ) ) |
| Tmx3 | ENSMUSG00000024614 | ENSMUST00000025515 | 95 | . . . . ) ) ) ) |
| Tnf | ENSMUSG00000024401 | ENSMUST00000167924 | 112 | . . . . ) ) ) ) |
| Tnf | ENSMUSG00000024401 | ENSMUST00000025263 | 115 | . . . . ) ) ) ) |
| Tnfaip3 | ENSMUSG00000019850 | ENSMUST00000105527 | 158 | ) ) ) . ) ) ) ) |
| Tnfaip8l2 | ENSMUSG00000013707 | ENSMUST00000013851 | 53 | ) ) ) . . . . . |
| Tnfrsf14 | ENSMUSG00000042333 | ENSMUST00000123514 | 57 | . . ) ) ) ) ) ) |
| Tnfrsf17 | ENSMUSG00000022496 | ENSMUST00000023140 | 117 | . . . . . . . ) |
| Tnfsf14 | ENSMUSG00000005824 | ENSMUST00000005976 | 83 | ( ( ( . . . . . |
| Tnip3 | ENSMUSG00000044162 | ENSMUST00000114236 | 152 | . . . ) ) ) ) ) |
| Tnn | ENSMUSG00000026725 | ENSMUST00000039178 | 117 | ) ) ) . ) ) ) ) |
| Tomm22 | ENSMUSG00000022427 | ENSMUST00000023062 | 3 | ( . . . . . . ( |
| Tomm7 | ENSMUSG00000028998 | ENSMUST00000030851 | 23 | ( ( ( ( . . . . |
| Top1 | ENSMUSG00000070544 | ENSMUST00000109468 | 291 | ) . . . . . . . |
| Toporsl | ENSMUSG00000028314 | ENSMUST00000029995 | 302 | ) ) . . ) ) ) ) |
| Toporsl | ENSMUSG00000028314 | ENSMUST00000107671 | 336 | ) ) . . ) ) ) ) |
| Tor1aip1 | ENSMUSG00000026466 | ENSMUST00000169241 | 146 | . . . . ) ) ) ) |
| Tox | ENSMUSG00000041272 | ENSMUST00000039987 | 260 | ( ( ( ( ( . . . |
| Tpo | ENSMUSG00000020673 | ENSMUST00000021005 | 47 | . . ) ) ) ) . . |
| Tpp2 | ENSMUSG00000041763 | ENSMUST00000188313 | 42 | . . . . . . . . |
| Tpp2 | ENSMUSG00000041763 | ENSMUST00000087933 | 50 | ) ) . . . . . . |
| Tpt1 | ENSMUSG00000060126 | ENSMUST00000110894 | 237 | ( ( ( . . . . . |
| Tra2a | ENSMUSG00000029817 | ENSMUST00000204189 | 39 | . ) ) ) ) . . ) |
| Tra2a | ENSMUSG00000029817 | ENSMUST00000031841 | 59 | . ) ) ) ) . . ) |
| Trak2 | ENSMUSG00000026028 | ENSMUST00000027186 | 360 | . . . ) ) ) ) ) |
| Trappc1 | ENSMUSG00000049299 | ENSMUST00000102602 | 67 | . . . . . . . . |
| Trappc11 | ENSMUSG00000038102 | ENSMUST00000039061 | 88 | ) ) . . . . . . |
| Trappc12 | ENSMUSG00000020628 | ENSMUST00000168129 | 100 | . ( ( . . ( ( ( |
| Trappc13 | ENSMUSG00000021711 | ENSMUST00000022224 | 203 | . ) ) ) ) . . . |
| Trappc13 | ENSMUSG00000021711 | ENSMUST00000141557 | 205 | . ) ) ) ) . . . |
| Trappc13 | ENSMUSG00000021711 | ENSMUST00000144060 | 249 | . ) ) ) ) . . . |
| Trappc5 | ENSMUSG00000040236 | ENSMUST00000044857 | 272 | ) . . . . . . ) |
| Trappc9 | ENSMUSG00000047921 | ENSMUST00000023276 | 8 | . . . . . . . . |
| Trem2 | ENSMUSG00000023992 | ENSMUST00000024791 | 76 | . . ( ( ( ( ( ( |
| Treml4 | ENSMUSG00000051682 | ENSMUST00000136272 | 3 | . . . . . . . . |
| Treml4 | ENSMUSG00000051682 | ENSMUST00000154335 | 3 | . . . . . . . . |
| Treml4 | ENSMUSG00000051682 | ENSMUST00000059873 | 45 | ) ) ) ) . . ) ) |
| Trim17 | ENSMUSG00000036964 | ENSMUST00000075141 | 58 | ( ( ( . . . . . |
| Trim2 | ENSMUSG00000027993 | ENSMUST00000107695 | 174 | ) ) ) ) . . . . |
| Trim25 | ENSMUSG00000000275 | ENSMUST00000107896 | 29 | . ( ( ( ( . ( ( |
| Trim6 | ENSMUSG00000072244 | ENSMUST00000098180 | 92 | ( ( ( ( ( ( ( ( |

| | | | | |
|---|---|---|---|---|
| Trim72 | ENSMUSG00000042828 | ENSMUST00000081042 | 88 | ) ) ) ) . ) ) ) |
| Trmt112 | ENSMUSG00000038812 | ENSMUST00000088257 | 349 | ) ) ) ) ) ) ( ( |
| Trmt112 | ENSMUSG00000038812 | ENSMUST00000116551 | 398 | . . ( ( . . . . |
| Trp53 | ENSMUSG00000059552 | ENSMUST00000108658 | 118 | . . . . . ) ) ) |
| Trp53 | ENSMUSG00000059552 | ENSMUST00000171247 | 118 | . . . . . ) ) ) |
| Trpv5 | ENSMUSG00000036899 | ENSMUST00000031901 | 68 | ) . . . . . . . |
| Trrap | ENSMUSG00000045482 | ENSMUST00000100467 | 38 | ) ) ) ) ) ) ) ) |
| Tsc22d1 | ENSMUSG00000022010 | ENSMUST00000022587 | 198 | . . . . . . . . |
| Tsc22d2 | ENSMUSG00000027806 | ENSMUST00000199164 | 958 | ) ) . . . . . . |
| Tsc22d2 | ENSMUSG00000027806 | ENSMUST00000099090 | 963 | . . . . . . . . |
| Tshr | ENSMUSG00000020963 | ENSMUST00000021343 | 3 | . ( ( ( ( ( . . |
| Tshr | ENSMUSG00000020963 | ENSMUST00000021346 | 9 | . ( ( ( ( ( . . |
| Tssk5 | ENSMUSG00000060794 | ENSMUST00000071119 | 187 | ) ) ) ) ) ) . . |
| Ttbk2 | ENSMUSG00000090100 | ENSMUST00000028740 | 54 | ) ) ) ) ) . ) ) |
| Ttc25 | ENSMUSG00000006784 | ENSMUST00000092684 | 150 | ) ) ) ) ) ) ) ) |
| Ttc37 | ENSMUSG00000033991 | ENSMUST00000091466 | 334 | ) ) ) ) ) ) ) ) |
| Ttc41 | ENSMUSG00000044937 | ENSMUST00000075632 | 344 | . . ( ( ( ( . . |
| Ttc41 | ENSMUSG00000044937 | ENSMUST00000061458 | 420 | . . ( ( ( ( . . |
| Ttc9c | ENSMUSG00000071660 | ENSMUST00000096751 | 303 | . . . . . ) ) ) |
| Ttll1 | ENSMUSG00000022442 | ENSMUST00000016897 | 304 | ( ( ( ( . . ( ( |
| Tuba1c | ENSMUSG00000043091 | ENSMUST00000058914 | 485 | . . ) ) ) ) ) ) |
| Tubb5 | ENSMUSG00000001525 | ENSMUST00000001566 | 189 | ) ) ) ) ) ) ) . |
| Tulp2 | ENSMUSG00000023467 | ENSMUST00000085331 | 115 | ( . . ( ( ( ( ( |
| Tulp2 | ENSMUSG00000023467 | ENSMUST00000107759 | 231 | ) ) . . . . . . |
| Tulp2 | ENSMUSG00000023467 | ENSMUST00000210813 | 232 | ) ) . . . . . . |
| Txlnb | ENSMUSG00000039891 | ENSMUST00000037964 | 153 | . . . . . . . . |
| Txndc15 | ENSMUSG00000021497 | ENSMUST00000021959 | 82 | . . . . . . . . |
| Txnip | ENSMUSG00000038393 | ENSMUST00000049093 | 228 | . . . . . ) ) . |
| Txnip | ENSMUSG00000038393 | ENSMUST00000074519 | 231 | . . . . . ) ) . |
| Tyw5 | ENSMUSG00000048495 | ENSMUST00000162686 | 562 | ( ( ( ( ( ( ( . |
| Uba3 | ENSMUSG00000030061 | ENSMUST00000089287 | 13 | ( . . . . . ( ( |
| Uba52 | ENSMUSG00000090137 | ENSMUST00000081940 | 49 | ) ) ) ) ) . ) ) |
| Ubd | ENSMUSG00000035186 | ENSMUST00000038844 | 29 | . . . . . ) ) ) |
| Ube2b | ENSMUSG00000020390 | ENSMUST00000020657 | 113 | ) ) ) ) ) ) ) ) |
| Ube2c | ENSMUSG00000001403 | ENSMUST00000088248 | 38 | . . . . . . . . |
| Ube2d2a | ENSMUSG00000091896 | ENSMUST00000170693 | 366 | ) ) . ) ) ) ) ) |
| Ube2d2b | ENSMUSG00000063447 | ENSMUST00000072578 | 348 | . . . . . . . ) |
| Ube2q2 | ENSMUSG00000032307 | ENSMUST00000121677 | 203 | . ) ) ) ( ( ( ( |
| Ube2q2 | ENSMUSG00000032307 | ENSMUST00000059555 | 744 | ) ) ) ( ( ( ( ( |
| Ube2t | ENSMUSG00000026429 | ENSMUST00000027687 | 267 | ) ) ) ) ) ) ) ) |
| Ube2w | ENSMUSG00000025939 | ENSMUST00000117146 | 115 | ) ) ) . . . ) ) |
| Ube4a | ENSMUSG00000059890 | ENSMUST00000117506 | 181 | ) . . . . . . . |
| Ubl3 | ENSMUSG00000001687 | ENSMUST00000079324 | 600 | . . . . . . . . |
| Ubl7 | ENSMUSG00000055720 | ENSMUST00000163329 | 57 | ) . . ( ( ( ( ( |
| Ubox5 | ENSMUSG00000027300 | ENSMUST00000028761 | 251 | ) ) ) ) ) ) ) ) |
| Ubqln2 | ENSMUSG00000050148 | ENSMUST00000060714 | 161 | ) ) ) ) ) ) ) . |
| Ubtf | ENSMUSG00000020923 | ENSMUST00000107119 | 106 | . . . ) ) ) ) ) |
| Ubxn6 | ENSMUSG00000019578 | ENSMUST00000019722 | 22 | . . . . . . . . |
| Ugt2b1 | ENSMUSG00000035836 | ENSMUST00000031183 | 18 | . . . . . . . . |
| Uhrf1bp1 | ENSMUSG00000039512 | ENSMUST00000114849 | 193 | ) ) . . . ( ( ( |
| Uimc1 | ENSMUSG00000025878 | ENSMUST00000026997 | 24 | . ( ( ( ( ( . . |
| Uimc1 | ENSMUSG00000025878 | ENSMUST00000099496 | 29 | . ( ( ( ( ( . . |
| Ulbp1 | ENSMUSG00000079685 | ENSMUST00000177585 | 241 | ) ) ) ) ) ) ) ) |
| Unc119 | ENSMUSG00000002058 | ENSMUST00000002127 | 9 | . . ( ( ( ( ( ( |
| Unc5c | ENSMUSG00000059921 | ENSMUST00000075282 | 113 | . . . . . . . . |
| Unc5c | ENSMUSG00000059921 | ENSMUST00000106236 | 471 | . . ) ) ) . ) ) |
| Unc79 | ENSMUSG00000021198 | ENSMUST00000101099 | 610 | ) ) ) ( ( ( ( ( |
| Unc93a | ENSMUSG00000067049 | ENSMUST00000084966 | 253 | ) ) ) ) . . . . |

| | | | | |
|---|---|---|---|---|
| Upp2 | ENSMUSG00000026839 | ENSMUST00000102755 | 151 | . ) ) ) ) ) ) ) |
| Upp2 | ENSMUSG00000026839 | ENSMUST00000059102 | 389 | ) ) ) ) . . ) ) |
| Uprt | ENSMUSG00000073016 | ENSMUST00000087867 | 77 | ) ) ) ) ) . . ) |
| Use1 | ENSMUSG00000002395 | ENSMUST00000019169 | 6 | . . . . . . . . |
| Ush1g | ENSMUSG00000045288 | ENSMUST00000103037 | 85 | ) ) ) ) ) . . . |
| Usp14 | ENSMUSG00000047879 | ENSMUST00000116669 | 122 | . . . . . . . . |
| Usp14 | ENSMUSG00000047879 | ENSMUST00000092096 | 196 | . ) ) ) . . . . |
| Usp17la | ENSMUSG00000054568 | ENSMUST00000067695 | 82 | . . . ) ) ) ) ) |
| Usp2 | ENSMUSG00000032010 | ENSMUST00000114830 | 34 | ( ( ( ( ( ( ( ( |
| Usp54 | ENSMUSG00000034235 | ENSMUST00000035340 | 707 | . . . . . . . . |
| Usp9x | ENSMUSG00000031010 | ENSMUST00000089302 | 317 | . . . . . . . . |
| Ust | ENSMUSG00000047712 | ENSMUST00000061601 | 381 | . . . . . ) ) ) |
| Utp20 | ENSMUSG00000004356 | ENSMUST00000004470 | 104 | ) . . . ) ) ) ) |
| Vamp1 | ENSMUSG00000030337 | ENSMUST00000100942 | 131 | ) ) ) ) ) ) ) ) |
| Vamp1 | ENSMUSG00000030337 | ENSMUST00000032487 | 161 | ) ) ) ) ) ) ) ) |
| Vars2 | ENSMUSG00000038838 | ENSMUST00000043674 | 299 | . ) ) ) ) ) ) . |
| Vax1 | ENSMUSG00000006270 | ENSMUST00000172821 | 10 | . . . . . . . . |
| Vdac2 | ENSMUSG00000021771 | ENSMUST00000022293 | 403 | . . . . . . . . |
| Vil1 | ENSMUSG00000026175 | ENSMUST00000027366 | 47 | ) . . . . . . . |
| Vldlr | ENSMUSG00000024924 | ENSMUST00000047645 | 169 | . . . . . . . . |
| Vldlr | ENSMUSG00000024924 | ENSMUST00000167487 | 169 | . . . . . . . . |
| Vldlr | ENSMUSG00000024924 | ENSMUST00000172302 | 506 | . . ) ) ) . . ) ) |
| Vma21 | ENSMUSG00000073131 | ENSMUST00000114577 | 434 | . . ) ) ) ) ) ) |
| Vmn1r18 | ENSMUSG00000091382 | ENSMUST00000164732 | 37 | . . . . . . . ) |
| Vmn1r185 | ENSMUSG00000091924 | ENSMUST00000171039 | 6 | . . . ( ( ( ( ( |
| Vmn1r189 | ENSMUSG00000099611 | ENSMUST00000186062 | 41 | . . . . . ) ) ) |
| Vmn1r19 | ENSMUSG00000115799 | ENSMUST00000089830 | 622 | ) ) ) ) ) ) ) ) |
| Vmn1r216 | ENSMUSG00000115697 | ENSMUST00000080253 | 59 | . . . . . . . ) |
| Vmn1r44 | ENSMUSG00000068234 | ENSMUST00000089420 | 77 | . . . . . . ) ) |
| Vmn1r53 | ENSMUSG00000057697 | ENSMUST00000076086 | 52 | . . ) ) ) ) ) ) |
| Vmn1r71 | ENSMUSG00000059206 | ENSMUST00000079113 | 684 | . . . . . . . ) |
| Vnn3 | ENSMUSG00000020010 | ENSMUST00000020190 | 59 | . ( ( ( ( ( ( . |
| Vps13d | ENSMUSG00000020220 | ENSMUST00000036579 | 140 | ) ) . . . ) ) ) |
| Vsig8 | ENSMUSG00000049598 | ENSMUST00000061835 | 65 | ( . . . . . . . |
| Vsir | ENSMUSG00000020101 | ENSMUST00000020301 | 138 | . . . . . . . ) |
| Vsir | ENSMUSG00000020101 | ENSMUST00000105460 | 138 | . . . . . . . ) |
| Vtn | ENSMUSG00000017344 | ENSMUST00000017488 | 155 | . . . . . . . . |
| Wasf2 | ENSMUSG00000028868 | ENSMUST00000084241 | 225 | ( ( ( ( ( . . . |
| Wdr5 | ENSMUSG00000026917 | ENSMUST00000113952 | 152 | ) ) ) ) ) . . ( |
| Wdr53 | ENSMUSG00000022787 | ENSMUST00000023474 | 231 | . . . . . . . . |
| Wdr63 | ENSMUSG00000043020 | ENSMUST00000160285 | 195 | ) ) ) ) ) ) ) ) |
| Wdr72 | ENSMUSG00000044976 | ENSMUST00000055879 | 340 | ) ) ) ) ) ) ) . |
| Wfdc13 | ENSMUSG00000067704 | ENSMUST00000088260 | 30 | . ( ( . . . . . |
| Wfdc3 | ENSMUSG00000076434 | ENSMUST00000103096 | 216 | . . . . . . . . |
| Wipf1 | ENSMUSG00000075284 | ENSMUST00000102679 | 208 | ) ) ) . . ) ) ) |
| Wnk1 | ENSMUSG00000045962 | ENSMUST00000088644 | 971 | ) ) ) ) ) ) . . |
| Wnk1 | ENSMUSG00000045962 | ENSMUST00000177761 | 971 | ) ) ) ) ) ) . . |
| Wnk1 | ENSMUSG00000045962 | ENSMUST00000060043 | 988 | . . . . . . . . |
| Wscd2 | ENSMUSG00000063430 | ENSMUST00000094452 | 947 | . . . . . . . . |
| Wt1 | ENSMUSG00000016458 | ENSMUST00000143043 | 170 | ) ) ) ) . ) ) ) |
| Xkr8 | ENSMUSG00000037752 | ENSMUST00000045550 | 14 | ( ( ( ( ( ( ( ( |
| Xpa | ENSMUSG00000028329 | ENSMUST00000030013 | 66 | ( ( ( . . . . . |
| Xpo6 | ENSMUSG00000000131 | ENSMUST00000009344 | 491 | ) ) . ) ) ) ) ) |
| Xpo6 | ENSMUSG00000000131 | ENSMUST00000168189 | 584 | . . ) ) ) ) ) . |
| Yeats4 | ENSMUSG00000020171 | ENSMUST00000020382 | 180 | ( ( ( ( . . . . |
| Yipf1 | ENSMUSG00000057375 | ENSMUST00000139527 | 65 | . ( ( ( ( ( . . |
| Yipf1 | ENSMUSG00000057375 | ENSMUST00000075693 | 86 | . ( ( ( ( ( . . |
| Yme1l1 | ENSMUSG00000026775 | ENSMUST00000028117 | 264 | ) ) ) ) ) ) ) ) |

| | | | | |
|---|---|---|---|---|
| Ywhag | ENSMUSG00000051391 | ENSMUST00000055808 | 156 | . . . . . . . . |
| Yy2 | ENSMUSG00000091736 | ENSMUST00000065806 | 138 | . . ( ( . . . . |
| Zbed3 | ENSMUSG00000041995 | ENSMUST00000045909 | 40 | . . . . ) ) ) ) |
| Zbtb18 | ENSMUSG00000063659 | ENSMUST00000193480 | 92 | . . . . . . . . |
| Zbtb18 | ENSMUSG00000063659 | ENSMUST00000094276 | 450 | . . . . . . . . |
| Zbtb18 | ENSMUSG00000063659 | ENSMUST00000077225 | 1256 | . ) ) ) ) . ) ) |
| Zbtb22 | ENSMUSG00000051390 | ENSMUST00000053429 | 156 | ) ) ) ) . . ) ) |
| Zbtb5 | ENSMUSG00000049657 | ENSMUST00000055028 | 91 | ( ( ( ( ( ( ( ( |
| Zbtb6 | ENSMUSG00000066798 | ENSMUST00000053098 | 35 | ( ( ( ( . . ( ( |
| Zbtb7b | ENSMUSG00000028042 | ENSMUST00000029677 | 271 | ) ) ) ) ) ) ) ) |
| Zbtb7b | ENSMUSG00000028042 | ENSMUST00000107435 | 781 | . ) ) ) ) ) ) ) |
| Zbtb9 | ENSMUSG00000079605 | ENSMUST00000120016 | 197 | . . . . . ) ) ) |
| Zc3h11a | ENSMUSG00000102976 | ENSMUST00000191896 | 248 | . . ) ) ) ) ) ) |
| Zc3h11a | ENSMUSG00000102976 | ENSMUST00000027736 | 256 | . . ) ) ) ) ) ) |
| Zdhhc12 | ENSMUSG00000015335 | ENSMUST00000081838 | 12 | ( ( ( ( . ( ( ( |
| Zdhhc12 | ENSMUSG00000015335 | ENSMUST00000102865 | 25 | ( ( ( . . . . . |
| Zdhhc5 | ENSMUSG00000034075 | ENSMUST00000035840 | 1149 | ) ) . ) ) . . . |
| Zfand2b | ENSMUSG00000026197 | ENSMUST00000027394 | 162 | ) ) ) ) ) ) ) ) |
| Zfat | ENSMUSG00000022335 | ENSMUST00000160248 | 124 | . . . . . . . . |
| Zfat | ENSMUSG00000022335 | ENSMUST00000162054 | 142 | . . . . . . . . |
| Zfhx3 | ENSMUSG00000038872 | ENSMUST00000043896 | 648 | ( ( ( ( . . ( ( |
| Zfp112 | ENSMUSG00000052675 | ENSMUST00000005413 | 99 | ) . . . . . . ) |
| Zfp131 | ENSMUSG00000094870 | ENSMUST00000177916 | 15 | ( . . . . . . . |
| Zfp131 | ENSMUSG00000094870 | ENSMUST00000178271 | 16 | ( . . . . . . . |
| Zfp148 | ENSMUSG00000022811 | ENSMUST00000165418 | 433 | ) ) ) ) ) ) ) ) |
| Zfp207 | ENSMUSG00000017421 | ENSMUST00000053740 | 17 | . . . . . . . ( |
| Zfp207 | ENSMUSG00000017421 | ENSMUST00000017567 | 111 | . . ) ) . . ) ) |
| Zfp207 | ENSMUSG00000017421 | ENSMUST00000165565 | 135 | . . ) ) . . ) ) |
| Zfp219 | ENSMUSG00000049295 | ENSMUST00000226522 | 119 | ( ( ( ( ( ( ( ( |
| Zfp219 | ENSMUSG00000049295 | ENSMUST00000067549 | 253 | ( ( ( ( ( ( ( ( |
| Zfp219 | ENSMUSG00000049295 | ENSMUST00000166169 | 341 | ) ) ) ) ) ) ) ) |
| Zfp277 | ENSMUSG00000055917 | ENSMUST00000069637 | 121 | ) . ) ) ) . . . |
| Zfp280d | ENSMUSG00000038535 | ENSMUST00000098576 | 98 | . . . . . . . . |
| Zfp324 | ENSMUSG00000004500 | ENSMUST00000038701 | 314 | ) ) ) . ( ( ( ( |
| Zfp354a | ENSMUSG00000020364 | ENSMUST00000109119 | 241 | . . . . . . . ) |
| Zfp429 | ENSMUSG00000078994 | ENSMUST00000109732 | 7 | ( ( ( ( ( ( . . |
| Zfp454 | ENSMUSG00000048728 | ENSMUST00000050595 | 133 | ) ) ) ) ) ) ) . |
| Zfp456 | ENSMUSG00000078995 | ENSMUST00000057070 | 56 | ( ( ( ( ( ( . . |
| Zfp532 | ENSMUSG00000042439 | ENSMUST00000049016 | 306 | . . . . . . . . |
| Zfp536 | ENSMUSG00000043456 | ENSMUST00000056338 | 482 | ( ( ( ( . . ( . |
| Zfp593 | ENSMUSG00000028840 | ENSMUST00000030644 | 37 | . . . . . . . ) |
| Zfp597 | ENSMUSG00000039789 | ENSMUST00000090522 | 293 | ) ) ) ) ) ) ) ) |
| Zfp608 | ENSMUSG00000052713 | ENSMUST00000064763 | 446 | ) ) ) ) ) ) ) ) |
| Zfp7 | ENSMUSG00000033669 | ENSMUST00000023179 | 48 | ) ) ) ) . ) ) ) |
| Zfp706 | ENSMUSG00000062397 | ENSMUST00000078976 | 201 | ) . ) ) ) ) ) ) |
| Zfp747 | ENSMUSG00000054381 | ENSMUST00000067425 | 128 | ( ( . . . ( ( ( |
| Zfp764 | ENSMUSG00000045757 | ENSMUST00000059199 | 122 | . ( ( ( ( ( ( ( |
| Zfp790 | ENSMUSG00000011427 | ENSMUST00000032796 | 235 | . . . . . . ) ) |
| Zfp82 | ENSMUSG00000098022 | ENSMUST00000182546 | 290 | . . ( ( ( ( ( ( |
| Zfp82 | ENSMUSG00000098022 | ENSMUST00000183190 | 335 | ( ( ( ( ( ( ( ( |
| Zfp82 | ENSMUSG00000098022 | ENSMUST00000080834 | 416 | . . ( ( ( ( ( ( |
| Zfp821 | ENSMUSG00000031728 | ENSMUST00000212000 | 432 | ( ( ( ( . ( ( ( |
| Zfp821 | ENSMUSG00000031728 | ENSMUST00000034163 | 478 | ( ( ( ( . ( ( ( |
| Zfp865 | ENSMUSG00000074405 | ENSMUST00000085427 | 94 | . . . . . . . . |
| Zfp865 | ENSMUSG00000074405 | ENSMUST00000076251 | 288 | . . . . . . . . |
| Zfp935 | ENSMUSG00000113450 | ENSMUST00000221747 | 24 | ( ( ( ( ( . . . |
| Zfp946 | ENSMUSG00000071266 | ENSMUST00000088763 | 430 | ) ) ) . ) ) ) ) |
| Zfp946 | ENSMUSG00000071266 | ENSMUST00000120222 | 439 | ) ) ) . ) ) ) ) |

| | | | | |
|---|---|---|---|---|
| Zfp949 | ENSMUSG00000032425 | ENSMUST00000161458 | 405 | ) ) ) . ) ) ) ) |
| Zfp949 | ENSMUSG00000032425 | ENSMUST00000162827 | 714 | ) ) ) . ) ) ) ) |
| Zfp966 | ENSMUSG00000089756 | ENSMUST00000109018 | 5 | ( ( ( ( . . . ( |
| Zfyve27 | ENSMUSG00000018820 | ENSMUST00000169536 | 196 | ) ) ) . . . . . |
| Zic3 | ENSMUSG00000067860 | ENSMUST00000088627 | 449 | . . . . . . ( ( |
| Zic4 | ENSMUSG00000036972 | ENSMUST00000172646 | 196 | . ) ) . . . . . |
| Zmynd8 | ENSMUSG00000039671 | ENSMUST00000177633 | 63 | ) ) . ( ( . . ( |
| Znrd1 | ENSMUSG00000036315 | ENSMUST00000113669 | 175 | ) ) ) ) ) ) ) ) |
| Znrf3 | ENSMUSG00000041961 | ENSMUST00000172492 | 264 | . . . . . . . . |
| Zscan12 | ENSMUSG00000036721 | ENSMUST00000225545 | 27 | . . ) ) . . ) ) |
| Zxdb | ENSMUSG00000073062 | ENSMUST00000101388 | 14 | ( ( ( ( ( . . . |

**Regulated_protein_syntesi**

X. laevis proteins whose translation was significantly changed by any cue-stimulation tested

Netrin_5min_average_ratio: Average log2 SILAC ratios of Netrin-1-stimulation (5min) vs. control stimulation. NaN = Not a number.

Netrin_5min_p.adj: Adjusted p.value, determined by LIMMA

| Gene names | Xenbase gene name | Netrin5min_average_ ratio | Netrin5min_p.adj | Netrin30min_average_ ratio | Netrin30min_p.adj | BDNF 5min_average_ ratio | BDNF 5min_p.adj | BDNF 30min_average_ ratio | BDNF 30min p.adj | Sema3a 5min_average_ ratio | Sema3a 5min_p.adj | Sema3a 30min_average_ ratio | Sema3a 30min_ p.adj |
|---|---|---|---|---|---|---|---|---|---|---|---|---|---|
| 3a;MASP | unnamed | 1.52 | 0 | 1.59 | NaN | -0.38 | 0.29 | -0.49 | 0.27 | -1.88 | 0 | NaN | NaN |
| acad9-prov | acad9 | 0.95 | 0.05 | 0.96 | 0.03 | 0.98 | 0.04 | 0.55 | 0.26 | 0.79 | 0.17 | NaN | NaN |
| act2;actc1;act3;MGC64484;MGC79012;acta2 | actc1;acta1;acta2;act3 | -0.28 | NaN | -0.11 | NaN | -0.05 | NaN | NaN | NaN | -1.53 | 0 | -0.86 | 0.2 |
| actb | actb | -1.34 | 0 | -0.14 | NaN | -0.53 | NaN | -0.01 | NaN | -2.27 | NaN | -1.91 | 0.12 |
| adss | adss | NaN | NaN | NaN | NaN | 1.1 | 0.04 | 0.24 | NaN | NaN | NaN | NaN | NaN |
| anp32e | anp32e | -0.58 | 0.22 | -1.35 | 0 | -0.08 | NaN | 0.96 | 0.17 | 0.11 | NaN | -0.86 | 0.3 |
| anxa7 | | 0.24 | NaN | 0.3 | NaN | 0.15 | NaN | -0.09 | NaN | 1.58 | 0.01 | 1.79 | 0.19 |
| aplp2 B | | 0.08 | NaN | 0.86 | 0.05 | -0.42 | 0.26 | -0.71 | 0.13 | -0.41 | 0.39 | 0.58 | 0.37 |
| arf1;MGC80261;arf-1 | arf1;arf3;arf5 | -0.46 | 0.33 | -1.45 | 0.04 | -0.07 | NaN | 0.44 | 0.25 | -0.24 | NaN | NaN | NaN |
| atp1b3 | atp1b3 | NaN | NaN | 1.59 | 0.02 | 0.15 | NaN | 0.59 | 0.22 | NaN | NaN | NaN | NaN |
| atp5b | atp5b | 0.29 | NaN | 0.18 | NaN | 0.37 | NaN | 0.65 | 0.2 | 1.26 | 0.03 | 0.47 | 0.27 |
| atp6v0d1 | atp6v0d1 | 0.63 | NaN | 1.29 | NaN | 0.32 | NaN | -0.38 | 0.45 | 3.6 | 0 | NaN | NaN |
| atp6v1g1 | atp6v1g1 | 0.92 | 0.22 | 4.37 | 0 | NaN | NaN | NaN | NaN | 0.41 | 0.38 | NaN | NaN |
| bdnf;bndf | bdnf | NaN | NaN | NaN | NaN | 0.87 | 0.04 | 0.17 | NaN | NaN | NaN | NaN | NaN |
| c9 | c9 | -0.39 | 0.37 | -0.23 | NaN | -0.95 | NaN | -1.35 | 0.11 | -2.35 | 0 | NaN | NaN |
| calm1;Cam | | 0.39 | 0.33 | 0.46 | 0.27 | -0.03 | NaN | -0.86 | 0.16 | 1.36 | 0 | 2.11 | NaN |
| calu | calu | 1.2 | 0.05 | 0.24 | NaN | 1.21 | 0.03 | 1.46 | 0.11 | NaN | NaN | NaN | NaN |
| canx | canx | 1.66 | NaN | 1.32 | 0.01 | -0.67 | NaN | -0.01 | NaN | NaN | NaN | NaN | NaN |
| cct5 | cct5 | 0.34 | NaN | -1 | 0.04 | -0.11 | NaN | NaN | NaN | NaN | NaN | NaN | NaN |
| CD38;LOC100036901 | cd38 | NaN | NaN | NaN | NaN | -1.09 | 0.1 | NaN | NaN | NaN | NaN | NaN | NaN |
| cd81 | cd81 | 2.44 | NaN | 4.31 | 0 | NaN | NaN | NaN | NaN | NaN | NaN | NaN | NaN |
| cdc42 | cdc42 | -0.34 | NaN | -1.6 | 0.02 | -0.99 | 0.06 | NaN | NaN | 0.54 | 0.3 | NaN | NaN |
| cdh4 | cdh4 | NaN | NaN | NaN | NaN | 1.54 | 0.02 | NaN | NaN | 1.45 | 0 | NaN | NaN |
| cfl1-a | cfl1 | 0.4 | 0.33 | 0.3 | NaN | 0.03 | NaN | -0.17 | NaN | 0.82 | 0.09 | 0.8 | 0.19 |
| cfl1-b | cfl1 | -0.49 | 0.27 | 0.11 | NaN | -0.33 | NaN | NaN | NaN | -1.69 | 0 | -0.69 | 0.2 |
| cirbp-a | cirbp | -1.2 | 0.01 | -0.73 | 0.1 | -0.11 | NaN | NaN | NaN | -0.13 | NaN | -0.55 | 0.2 |
| cltb-prov | cltb | NaN | NaN | 1.44 | 0.04 | -1.97 | 0 | NaN | NaN | -1.87 | 0 | -0.95 | 0.26 |
| cndp2-prov | cndp2 | -0.2 | NaN | 0.23 | NaN | 0.37 | NaN | 0.08 | NaN | -1.09 | 0.07 | -0.51 | 0.29 |

| | | | | | | | | | | | | | |
|---|---|---|---|---|---|---|---|---|---|---|---|---|---|
| col18a1 | col18a1 | 0 | NaN | -0.88 | 0.04 | -0.09 | NaN | -0.1 | NaN | 0.98 | 0.1 | 0.25 | NaN |
| col2a1 | col2a1 | -0.41 | NaN | -1.4 | 0 | -0.86 | 0.06 | -0.69 | 0.2 | NaN | NaN | NaN | NaN |
| copa-prov | copa | 0.8 | 0.27 | -0.52 | 0.23 | -1.23 | 0.06 | -1.46 | NaN | NaN | NaN | NaN | NaN |
| copb2 | copb2 | 0.27 | NaN | -0.79 | 0.08 | -0.09 | NaN | 0.17 | NaN | NaN | NaN | NaN | NaN |
| crc | calr | 1.17 | 0.01 | 0.88 | 0.04 | -0.01 | NaN | -0.52 | 0.26 | NaN | NaN | NaN | NaN |
| cst3 | cst3 | 0.08 | NaN | 1.05 | NaN | -1.13 | 0.04 | NaN | NaN | 0.24 | NaN | NaN | NaN |
| ctnnb1 | ctnnb1;ctnnb1 | 1.69 | 0.02 | 1.59 | NaN | 0.33 | NaN | NaN | NaN | NaN | NaN | NaN | NaN |
| cyp | | -0.55 | 0.25 | -0.07 | NaN | -1.01 | 0.03 | 0.28 | NaN | -0.4 | 0.32 | 0.18 | NaN |
| dagfl5;LOC398500 | dag1 | NaN | NaN | NaN | NaN | 2.48 | 0 | NaN | NaN | NaN | NaN | NaN | NaN |
| ddb1 | ddb1 | NaN | NaN | -1.49 | 0.03 | 1.56 | 0.02 | 0.84 | 0.14 | NaN | NaN | NaN | NaN |
| ddx39a | ddx39a | 1.06 | 0.09 | -0.2 | NaN | NaN | NaN | NaN | NaN | NaN | NaN | NaN | NaN |
| dfna20 | | -0.86 | 0.07 | -0.76 | 0.26 | NaN | NaN | NaN | NaN | NaN | NaN | NaN | NaN |
| dynll1 | dynll1 | -1.29 | 0.09 | NaN | NaN | NaN | NaN | NaN | NaN | NaN | NaN | NaN | NaN |
| eef1as | eef1a1 | 0.98 | NaN | 0.23 | NaN | 1.19 | 0.01 | 0.75 | 0.18 | -0.66 | 0.16 | -0.45 | 0.46 |
| EEF1D | | 0.36 | NaN | 0.51 | 0.23 | 0.95 | 0.04 | 0.2 | NaN | 0.49 | 0.28 | 0 | NaN |
| eef2.1 | eef2.1 | -0.79 | 0.11 | -1.76 | 0 | -0.12 | NaN | -0.27 | NaN | NaN | NaN | NaN | NaN |
| eif3e-b;eif3e-a | eif3e;eif3e | -1.75 | 0.02 | NaN | NaN | NaN | NaN | NaN | NaN | NaN | NaN | NaN | NaN |
| eif5a | eif5a | 0.12 | NaN | -0.77 | 0.08 | 0.48 | 0.25 | 0.35 | NaN | -0.25 | NaN | -2.26 | 0.12 |
| ero1l | | -2.27 | 0 | NaN | NaN | NaN | NaN | NaN | NaN | NaN | NaN | NaN | NaN |
| fkbp2;MGC80429 | fkbp2;fkbp2 | -0.05 | NaN | 0.24 | NaN | -2.53 | 0 | 0.72 | NaN | -0.1 | NaN | NaN | NaN |
| gapdh | | -0.3 | NaN | -0.44 | 0.29 | -0.44 | 0.25 | -0.64 | 0.2 | -0.87 | 0.07 | 0.17 | NaN |
| glul | glul | NaN | NaN | 1.05 | 0.07 | 1.2 | 0.03 | -0.14 | NaN | NaN | NaN | NaN | NaN |
| gpm6a | gpm6a | -0.12 | NaN | NaN | NaN | 0.85 | 0.07 | -1.24 | 0.15 | 1.21 | 0.04 | NaN | NaN |
| gyg | | -0.11 | NaN | -1.22 | 0.01 | -0.63 | 0.22 | -0.26 | NaN | 0.06 | NaN | NaN | NaN |
| H2B | hist1h2bj | -0.15 | NaN | 0.43 | 0.29 | 0.86 | 0.07 | 0 | NaN | 1.09 | 0.02 | 1.49 | 0.19 |
| hnrnph1 | hnrnph1 | 1.01 | 0.06 | 1.51 | 0.01 | 2.15 | NaN | 1.85 | NaN | NaN | NaN | NaN | NaN |
| hnrpa2b1 | hnrnpa2b1 | 1.68 | NaN | 0.93 | NaN | 0.69 | 0.1 | 1.16 | 0.13 | NaN | NaN | NaN | NaN |
| Hspd1 | hspd1 | 0.17 | NaN | -0.05 | NaN | 1.22 | 0.03 | NaN | NaN | NaN | NaN | NaN | NaN |
| IsoT | usp5 | NaN | NaN | NaN | NaN | 1.12 | 0.09 | 0.32 | NaN | NaN | NaN | NaN | NaN |
| klc1 | klc1 | 1.66 | 0.03 | NaN | NaN | 0.32 | NaN | -0.83 | 0.16 | NaN | NaN | NaN | NaN |
| ldha | ldhb | 1.97 | 0 | 2.23 | NaN | 1.09 | 0.01 | 0.96 | 0.13 | 2.07 | 0 | 3.4 | 0.12 |
| ldlr2-a;ldlr-b | ldlr | NaN | NaN | -1.17 | 0.09 | 0.43 | 0.33 | NaN | NaN | NaN | NaN | NaN | NaN |
| lgals3 | lgals3 | -0.36 | NaN | -0.86 | 0.05 | -1.09 | 0.04 | NaN | NaN | 0.57 | 0.21 | -0.16 | NaN |
| lin7c | lin7c | NaN | NaN | NaN | NaN | -1.33 | 0.01 | NaN | NaN | NaN | NaN | NaN | NaN |
| LOC100037025 | ganab | -0.14 | NaN | NaN | NaN | 0.97 | 0.04 | 0.12 | NaN | NaN | NaN | NaN | NaN |
| LOC100037195 | mccc2 | -0.81 | 0.27 | NaN | NaN | -1.74 | 0 | 1.23 | 0.16 | NaN | NaN | NaN | NaN |
| LOC100127277 | unnamed | 0.13 | NaN | 1 | 0.02 | 0.46 | 0.39 | NaN | NaN | NaN | NaN | NaN | NaN |
| LOC100127332 | | 1.39 | NaN | 0.63 | NaN | -0.77 | 0.06 | 0.89 | 0.13 | 0.47 | 0.29 | -0.4 | 0.31 |
| LOC100137679 | | NaN | NaN | -2.52 | 0 | NaN | NaN | NaN | NaN | NaN | NaN | NaN | NaN |
| LOC100158365 | usp7 | NaN | NaN | NaN | NaN | 1.84 | 0 | NaN | NaN | NaN | NaN | NaN | NaN |
| LOC397710 | cntn1 | NaN | NaN | NaN | NaN | 1.5 | 0.03 | NaN | NaN | NaN | NaN | NaN | NaN |
| LOC397716 | psmb6 | -0.16 | NaN | 1.75 | 0 | 0.27 | NaN | 0.81 | 0.13 | -0.94 | 0.05 | -0.54 | 0.26 |
| LOC397732;atp5a | atp5a1 | 0.02 | NaN | 1.45 | 0 | 0.02 | NaN | 0.5 | 0.19 | 0.72 | NaN | NaN | NaN |
| LOC397751;hnrnpa1 | | 0.57 | 0.27 | 1.32 | NaN | 1.68 | 0 | 1.28 | 0.11 | -0.43 | NaN | NaN | NaN |
| LOC397895 | gsn | -0.01 | NaN | 0.91 | 0.04 | 0.15 | NaN | 0.1 | NaN | 0.14 | NaN | 0.97 | 0.29 |
| LOC397911 | lmnb1 | -0.46 | 0.43 | -1.14 | 0.04 | 0.07 | NaN | -1.29 | NaN | NaN | NaN | NaN | NaN |

| | | | | | | | | | | | | | |
|---|---|---|---|---|---|---|---|---|---|---|---|---|---|
| LOC397931 | | 0.19 | NaN | 0.81 | 0.07 | 0.06 | NaN | 0.02 | NaN | -1.11 | 0.02 | -0.4 | 0.3 |
| LOC397939 | scg2 | NaN | NaN | NaN | NaN | NaN | NaN | NaN | NaN | 1.81 | 0 | NaN | NaN |
| LOC398217 | gpm6b | -1.42 | 0.06 | 0.07 | NaN | 0.85 | 0.11 | NaN | NaN | NaN | NaN | NaN | NaN |
| LOC398287 | olfm1 | -1.29 | 0.01 | -0.34 | NaN | -0.16 | NaN | -0.17 | NaN | 0.5 | 0.32 | NaN | NaN |
| LOC398455;hnrnpa0 | hnrnpa0 | 1.35 | 0.03 | 1.13 | NaN | -0.65 | 0.27 | NaN | NaN | NaN | NaN | NaN | NaN |
| LOC398472 | anxa5 | 0.34 | NaN | 1.13 | 0.02 | 0.69 | 0.25 | NaN | NaN | NaN | NaN | NaN | NaN |
| LOC398563 | | 1.15 | 0.06 | 0.5 | 0.32 | -0.51 | 0.35 | NaN | NaN | NaN | NaN | NaN | NaN |
| LOC414678 | myl6 | 1.16 | NaN | 0.05 | NaN | 0.83 | 0.08 | 0.18 | NaN | -1.31 | 0.01 | -0.78 | 0.19 |
| LOC443721 | ctsd | -0.55 | 0.27 | -0.43 | 0.46 | 0.28 | NaN | 1.49 | 0.11 | -1.56 | 0.01 | NaN | NaN |
| LOC445859 | | -1.26 | 0.04 | -0.49 | 0.33 | -2.07 | 0 | NaN | NaN | -1 | 0.09 | NaN | NaN |
| LOC446231;ube2l3 | ube2l3 | 0.25 | NaN | NaN | NaN | 1.49 | 0.03 | NaN | NaN | NaN | NaN | NaN | NaN |
| LOC446265 | psmd1 | 0.36 | NaN | -0.12 | NaN | 1.25 | 0.01 | -0.3 | NaN | NaN | NaN | NaN | NaN |
| LOC494658 | hnrnph3 | 1.51 | 0.01 | NaN | NaN | 0.22 | NaN | -0.44 | NaN | NaN | NaN | NaN | NaN |
| LOC494665 | rab14 | 0.64 | 0.27 | 1.73 | 0.01 | 0.35 | NaN | -0.52 | NaN | 0.48 | 0.33 | NaN | NaN |
| LOC494725 | nudc | -1.31 | 0.03 | -1.14 | NaN | -0.73 | NaN | 0.21 | NaN | -1.13 | 0.06 | NaN | NaN |
| LOC494763 | slc2a1 | 2.93 | 0 | 0.59 | 0.27 | 0 | NaN | 1.4 | 0.15 | 1.94 | 0 | NaN | NaN |
| LOC495025 | uchl3 | -0.03 | NaN | NaN | NaN | -0.25 | NaN | -0.35 | NaN | -1.05 | 0.08 | NaN | NaN |
| LOC495169 | pdia4 | 1.51 | 0.05 | 0.84 | NaN | NaN | NaN | NaN | NaN | NaN | NaN | NaN | NaN |
| LOC495270 | ppib | -1.24 | NaN | 0.6 | 0.18 | 1.46 | 0.03 | NaN | NaN | 1.63 | NaN | NaN | NaN |
| LOC495277 | psma6 | -0.52 | 0.27 | 0.03 | NaN | -1.17 | 0.01 | 0.23 | NaN | -0.22 | NaN | -0.1 | NaN |
| LOC495300 | pdcd5 | 0.04 | NaN | -1.98 | 0 | NaN | NaN | NaN | NaN | 0.1 | NaN | NaN | NaN |
| LOC495318 | lsm4 | NaN | NaN | 4.15 | 0 | NaN | NaN | NaN | NaN | NaN | NaN | NaN | NaN |
| LOC495940 | pcsk5 | 0.83 | 0.09 | 0.2 | NaN | -0.36 | NaN | 0.28 | NaN | -0.56 | 0.29 | NaN | NaN |
| LOC495981 | mgst3 | 1.51 | 0 | -0.56 | NaN | -0.04 | NaN | -0.62 | 0.21 | 0.75 | 0.18 | NaN | NaN |
| LOC496060 | fabp7 | 1.14 | 0.07 | 2.69 | NaN | -0.09 | NaN | -0.12 | NaN | 0.76 | 0.11 | 1.09 | 0.18 |
| LOC496072 | cetp | -0.93 | 0.14 | -1.59 | 0.01 | 0.06 | NaN | -0.32 | NaN | 0.34 | NaN | NaN | NaN |
| LOC733431 | | 0.82 | 0.09 | 0.54 | 0.26 | NaN | NaN | NaN | NaN | NaN | NaN | NaN | NaN |
| LOC734152 | | 0.35 | NaN | NaN | NaN | -0.37 | NaN | NaN | NaN | 3.17 | 0 | NaN | NaN |
| MARCKS | | 0.45 | 0.36 | NaN | NaN | 0.43 | 0.25 | NaN | NaN | 2.1 | 0 | NaN | NaN |
| mdh2 | mdh2 | -0.22 | NaN | -0.85 | 0.09 | -0.17 | NaN | NaN | NaN | -0.69 | 0.14 | NaN | NaN |
| mdk-b | mdk | -0.78 | 0.11 | -0.44 | 0.29 | 0.55 | 0.21 | -0.38 | 0.53 | -1.29 | 0.03 | NaN | NaN |
| mec-12 | tuba4b | -0.26 | NaN | -0.07 | NaN | -0.89 | 0.06 | NaN | NaN | -1.02 | 0.03 | -0.49 | 0.31 |
| metrnl | metrnl | 0.47 | 0.32 | 0.91 | 0.04 | -0.95 | 0.04 | -1.2 | 0.11 | 1.16 | 0.05 | NaN | NaN |
| MGC114675 | atp6v1a | -0.36 | NaN | 1.14 | 0.04 | 0.59 | 0.19 | 0.64 | 0.15 | NaN | NaN | NaN | NaN |
| MGC114755 | cisd1 | 1.34 | 0.01 | 0.08 | NaN | -0.15 | NaN | NaN | NaN | 0.55 | NaN | NaN | NaN |
| MGC114839 | uroc1 | -0.33 | NaN | -0.88 | 0.12 | -0.41 | 0.35 | -0.96 | 0.13 | -1.08 | 0.07 | NaN | NaN |
| MGC114846 | aldh16a1 | NaN | NaN | -1.73 | 0.01 | NaN | NaN | NaN | NaN | NaN | NaN | NaN | NaN |
| MGC114910;LOC733147;MGC82058 | ezr;msn | NaN | NaN | NaN | NaN | -1.3 | 0.05 | 0.61 | 0.27 | NaN | NaN | NaN | NaN |
| MGC115336 | aplp1 | -0.17 | NaN | 0.41 | 0.34 | 0.08 | NaN | 0.59 | NaN | -1 | 0.09 | -0.68 | 0.26 |
| MGC130849 | gpx4 | NaN | NaN | NaN | NaN | -0.88 | 0.06 | NaN | NaN | NaN | NaN | NaN | NaN |
| MGC131357 | ndufa8 | NaN | NaN | -1.69 | 0.01 | NaN | NaN | NaN | NaN | NaN | NaN | NaN | NaN |
| MGC132155 | nasp | 0.12 | NaN | -0.77 | 0.08 | -0.63 | 0.17 | -0.81 | 0.18 | NaN | NaN | 0.38 | 0.48 |
| MGC132157 | anp32c | 0.93 | 0.05 | 0.89 | 0.07 | NaN | NaN | NaN | NaN | 0.64 | 0.25 | 0.93 | 0.19 |
| MGC132180 | | 0.85 | NaN | 1.57 | NaN | 1.3 | 0 | 0.91 | NaN | 1.5 | 0 | 1.92 | 0.19 |
| MGC132184 | basp1 | NaN | NaN | -0.06 | NaN | -1.55 | 0.02 | NaN | NaN | 0.07 | NaN | NaN | NaN |
| MGC132191 | uchl1 | NaN | NaN | NaN | NaN | -1.97 | 0 | NaN | NaN | NaN | NaN | NaN | NaN |

| | | | | | | | | | | | | | |
|---|---|---|---|---|---|---|---|---|---|---|---|---|---|
| MGC52616 | hspa9 | -1.39 | 0.07 | 0 | NaN | NaN | NaN | 0.43 | 0.38 | NaN | NaN | NaN | NaN |
| MGC64292 | pcolce | 0.9 | NaN | 0.91 | 0.04 | -0.03 | NaN | 0.46 | 0.27 | 0.37 | NaN | NaN | NaN |
| MGC64541 | psap | 0.21 | NaN | -1.04 | 0.04 | 0.31 | NaN | 0.39 | 0.37 | NaN | NaN | NaN | NaN |
| MGC68473 | epb41l3 | -1.5 | 0.05 | NaN | NaN | 0.27 | NaN | 0.14 | NaN | NaN | NaN | NaN | NaN |
| MGC68479 | | 1.46 | 0 | -0.4 | NaN | 2.03 | NaN | NaN | NaN | -0.08 | NaN | -0.15 | NaN |
| MGC68491 | pmp2 | NaN | NaN | -1.29 | 0.06 | -0.97 | NaN | 0.09 | NaN | 0.43 | 0.38 | NaN | NaN |
| MGC68523;rab7-prov | rab7a;rab7a | -0.37 | NaN | 0.82 | 0.15 | 1.09 | 0.02 | 1.01 | 0.11 | NaN | NaN | NaN | NaN |
| MGC68529;rps12;LOC100126614 | rps12;rps12 | -0.19 | NaN | 1.47 | 0 | -0.72 | 0.24 | NaN | NaN | -1.15 | 0.02 | -0.87 | 0.19 |
| MGC68629 | rab10 | -2.91 | 0 | NaN | NaN | -1.09 | 0.04 | -0.95 | 0.15 | NaN | NaN | NaN | NaN |
| MGC68767 | napsa | -1.18 | 0.01 | -1.35 | 0.01 | -0.97 | 0.04 | -0.31 | NaN | 0.2 | NaN | NaN | NaN |
| MGC68785 | tkt | 0.4 | NaN | NaN | NaN | 1.13 | 0.09 | NaN | NaN | NaN | NaN | NaN | NaN |
| MGC68971 | pc.1 | -0.29 | NaN | NaN | NaN | 1.37 | 0.04 | 1.55 | 0.13 | NaN | NaN | NaN | NaN |
| MGC78867 | lta4h | 0.39 | NaN | 1.71 | 0.01 | 0.98 | NaN | -0.74 | 0.2 | 0.78 | 0.17 | NaN | NaN |
| MGC79030 | hspe1 | -1.4 | 0.06 | NaN | NaN | NaN | NaN | NaN | NaN | -1.52 | NaN | NaN | NaN |
| MGC80011 | pon2;pon2 | -0.16 | NaN | 0.22 | NaN | -0.93 | 0.09 | 0.14 | NaN | NaN | NaN | NaN | NaN |
| MGC80163 | | -0.37 | NaN | 0.22 | NaN | -1.52 | 0 | NaN | NaN | 0.29 | NaN | NaN | NaN |
| MGC80186 | g3bp1 | -0.79 | 0.27 | -1.45 | 0.04 | NaN | NaN | NaN | NaN | NaN | NaN | NaN | NaN |
| MGC80200 | nrcam | NaN | NaN | NaN | NaN | 1.27 | 0.06 | NaN | NaN | NaN | NaN | NaN | NaN |
| MGC80245 | | NaN | NaN | NaN | NaN | -1.35 | 0.04 | NaN | NaN | NaN | NaN | NaN | NaN |
| MGC80370 | kazald1 | 3.9 | 0 | NaN | NaN | NaN | NaN | NaN | NaN | NaN | NaN | NaN | NaN |
| MGC80718 | gpi | -0.5 | 0.31 | -0.7 | 0.22 | -0.86 | 0.07 | 0.42 | 0.35 | NaN | NaN | NaN | NaN |
| MGC80961 | cttn | NaN | NaN | 0.58 | NaN | -1.4 | 0 | -0.38 | 0.35 | NaN | NaN | NaN | NaN |
| MGC81140 | lap3 | -1.4 | NaN | -1.91 | 0.01 | -0.66 | 0.26 | -0.82 | 0.13 | NaN | NaN | NaN | NaN |
| MGC81156 | gstt1 | 0.24 | NaN | 1.31 | 0.01 | 0.52 | 0.27 | 0.32 | NaN | -0.37 | NaN | -0.39 | 0.29 |
| MGC81323 | tuba4a | 0.75 | 0.13 | 0.59 | 0.22 | -0.8 | 0.09 | 0.68 | NaN | -0.24 | NaN | -0.97 | 0.2 |
| MGC81824 | dcn | 0.69 | 0.15 | 1.08 | 0.01 | 0.46 | 0.24 | 0.81 | 0.16 | -0.33 | NaN | NaN | NaN |
| MGC81848 | | -2.4 | NaN | -2.63 | 0 | -0.04 | NaN | -0.44 | 0.27 | NaN | NaN | NaN | NaN |
| MGC81889;MGC80109;MGC154789 | uba52;rps27a;ubc;ubb | -0.77 | 0.11 | -0.75 | 0.09 | 0.6 | NaN | -0.83 | 0.13 | -0.44 | 0.3 | -0.25 | NaN |
| MGC81911 | ech1 | NaN | NaN | NaN | NaN | NaN | NaN | NaN | NaN | -2.22 | 0 | NaN | NaN |
| MGC82200 | snx1 | NaN | NaN | NaN | NaN | 2.08 | 0 | NaN | NaN | NaN | NaN | NaN | NaN |
| MGC82306 | rps18 | -1.09 | 0.08 | 0.46 | 0.35 | -1.25 | 0.06 | -0.89 | 0.16 | NaN | NaN | NaN | NaN |
| MGC82327 | | -0.43 | NaN | 0.57 | NaN | -0.84 | 0.07 | -1.52 | 0.13 | -0.78 | 0.17 | -0.13 | NaN |
| MGC82428 | cox6b1 | NaN | NaN | NaN | NaN | NaN | NaN | NaN | NaN | -0.85 | 0.08 | -0.9 | 0.2 |
| MGC82602 | | 1.52 | 0.05 | NaN | NaN | 3.13 | NaN | 2.2 | NaN | 1.07 | 0.07 | 0.38 | 0.29 |
| MGC82702 | hba-l5 | -0.32 | NaN | 0.99 | 0.08 | 1.01 | 0.03 | 0.79 | NaN | -0.35 | NaN | -0.03 | NaN |
| MGC82894 | psmd3 | 0.45 | 0.33 | 0.03 | NaN | 0.3 | NaN | 0.13 | NaN | 1.22 | 0.04 | NaN | NaN |
| MGC82927 | f7 | 1.97 | 0.01 | NaN | NaN | NaN | NaN | NaN | NaN | NaN | NaN | NaN | NaN |
| MGC82947 | bola2 | 2.08 | 0 | NaN | NaN | NaN | NaN | NaN | NaN | NaN | NaN | NaN | NaN |
| MGC83065 | cand1 | -0.29 | NaN | NaN | NaN | 1.49 | 0.03 | NaN | NaN | NaN | NaN | NaN | NaN |
| MGC83078;PRDX2 | prdx2 | -1.35 | 0 | -0.72 | 0.15 | NaN | NaN | NaN | NaN | 0.17 | NaN | 0.75 | 0.2 |
| MGC83495 | vwa5a.2 | -1.04 | 0.17 | -1.47 | 0.01 | -0.64 | NaN | -0.88 | NaN | -1.02 | 0.08 | NaN | NaN |
| MGC83546 | amph | NaN | NaN | NaN | NaN | 1.6 | 0.02 | NaN | NaN | NaN | NaN | NaN | NaN |
| MGC83563 | nnt | NaN | NaN | -0.92 | 0.18 | -0.98 | 0.04 | 0.05 | NaN | NaN | NaN | NaN | NaN |
| MGC83669 | ppa2 | -1.88 | 0.01 | NaN | NaN | NaN | NaN | NaN | NaN | NaN | NaN | NaN | NaN |
| MGC83716 | fkbp4 | NaN | NaN | NaN | NaN | 1.98 | 0 | NaN | NaN | NaN | NaN | NaN | NaN |
| MGC83971 | rab6b | -1.16 | 0.06 | -1.85 | NaN | 0.3 | NaN | -0.12 | NaN | 0.89 | 0.07 | 1.02 | 0.2 |

| | | | | | | | | | | | | | |
|---|---|---|---|---|---|---|---|---|---|---|---|---|---|
| MGC84055 | pacsin1 | NaN | NaN | NaN | NaN | -2.1 | 0 | NaN | NaN | NaN | NaN | NaN | NaN |
| MGC84105 | prkcsh | -0.25 | NaN | 1.14 | 0.02 | -0.2 | NaN | 1.29 | NaN | NaN | NaN | NaN | NaN |
| MGC84194 | mfge8 | -1.13 | 0.14 | -1.21 | 0.01 | 0.33 | NaN | NaN | NaN | NaN | NaN | NaN | NaN |
| MGC84281 | rbp4l | -0.92 | 0.05 | -0.3 | NaN | -0.7 | 0.09 | -0.59 | NaN | 0.81 | 0.09 | 0.19 | NaN |
| MGC84451 | ywhah | NaN | NaN | NaN | NaN | -1.71 | 0 | -1.1 | 0.15 | NaN | NaN | NaN | NaN |
| MGC84683 | spock2 | -0.48 | 0.28 | 0.21 | NaN | 0.71 | 0.09 | 0.03 | NaN | 0.77 | 0.18 | NaN | NaN |
| MGC85130 | atp6v1h | -1.88 | 0.01 | NaN | NaN | 0.45 | 0.27 | 0.61 | NaN | NaN | NaN | NaN | NaN |
| MGC85281 | pmp2 | -0.43 | 0.34 | 0.34 | NaN | -0.34 | NaN | NaN | NaN | 1.37 | 0.02 | NaN | NaN |
| MGC85310 | rpl11 | 1.19 | 0.05 | NaN | NaN | -1.73 | 0.01 | -0.44 | NaN | NaN | NaN | NaN | NaN |
| MGC85348 | rpl23a | 0.13 | NaN | NaN | NaN | 1.17 | 0.07 | 0.26 | NaN | NaN | NaN | NaN | NaN |
| MGC86492 | loc100498624 | -0.65 | 0.17 | -1.29 | 0 | -0.5 | 0.2 | -0.58 | 0.27 | -1.25 | 0.01 | -1.1 | 0.19 |
| mt-nd5;nad5 | nd5 | NaN | NaN | NaN | NaN | 1.25 | 0.06 | NaN | NaN | NaN | NaN | NaN | NaN |
| mvp | mvp | NaN | NaN | -1.65 | 0 | NaN | NaN | NaN | NaN | NaN | NaN | NaN | NaN |
| N1 | nasp | -0.45 | 0.33 | -0.3 | NaN | -0.62 | 0.18 | -0.47 | 0.33 | 1.31 | 0.03 | NaN | NaN |
| naca | naca.2 | 0.81 | NaN | 0.8 | 0.07 | 0.22 | NaN | 0.59 | 0.21 | NaN | NaN | NaN | NaN |
| ndrg2 | ndrg2 | -0.06 | NaN | 1.85 | 0.01 | -0.18 | NaN | 0.68 | 0.19 | NaN | NaN | NaN | NaN |
| ndrg4-a | ndrg3 | NaN | NaN | NaN | NaN | 2 | 0 | NaN | NaN | NaN | NaN | NaN | NaN |
| nell2 | nell2 | -1.69 | NaN | -1.36 | 0.05 | -0.04 | NaN | -0.04 | NaN | NaN | NaN | NaN | NaN |
| Nogo;RTN4 | rtn4 | -0.07 | NaN | 0.58 | 0.28 | 0.2 | NaN | -0.43 | 0.28 | 1.08 | 0.07 | 0.35 | NaN |
| npm1 | npm1 | 0.92 | NaN | 0.07 | NaN | 0.76 | 0.16 | NaN | NaN | 0.98 | 0.04 | 0.84 | 0.19 |
| nrn1-a;nrn1-b | nrn1;nrn1 | 2.88 | 0 | NaN | NaN | NaN | NaN | NaN | NaN | NaN | NaN | NaN | NaN |
| pafah1b1 | pafah1b1 | -1.03 | 0.03 | 0.17 | NaN | -0.57 | 0.17 | -0.51 | 0.2 | -0.43 | 0.3 | -0.17 | NaN |
| Park7;SP22;MGC84701 | park7;park7 | NaN | NaN | -2.1 | 0 | NaN | NaN | NaN | NaN | -1.28 | 0.03 | NaN | NaN |
| pcca | pcca | -1.43 | 0.06 | 0.76 | NaN | -0.48 | 0.3 | 0.17 | NaN | NaN | NaN | NaN | NaN |
| PdhE1beta-1 | pdhb | 0.67 | 0.32 | 1.45 | 0 | NaN | NaN | NaN | NaN | NaN | NaN | NaN | NaN |
| pgd | pgd | NaN | NaN | NaN | NaN | 0.96 | 0.07 | NaN | NaN | NaN | NaN | NaN | NaN |
| Pin1;pin1 | pin1;pin1 | 1.38 | 0.07 | NaN | NaN | NaN | NaN | NaN | NaN | NaN | NaN | NaN | NaN |
| pkm | pkm | -0.1 | NaN | 0.75 | 0.09 | 0.8 | 0.06 | -0.12 | NaN | 0.24 | NaN | -1.27 | 0.19 |
| plscr2 | plscr2 | 0.93 | 0.22 | 1.53 | 0 | 2.04 | 0 | NaN | NaN | 0.3 | NaN | NaN | NaN |
| PP2A | ppp2r2a | 0.61 | NaN | 1.87 | 0.01 | -0.36 | NaN | NaN | NaN | NaN | NaN | NaN | NaN |
| ppp3r1 | ppp3r1 | 1.2 | 0.05 | 1.13 | NaN | NaN | NaN | NaN | NaN | NaN | NaN | NaN | NaN |
| Prom-1;Proml-1 | | -0.55 | 0.24 | 0.33 | NaN | -0.01 | NaN | 0.37 | NaN | -1.87 | 0 | NaN | NaN |
| psma3 | psma3 | 1.04 | 0.03 | 1.1 | 0.01 | 0.29 | NaN | 0.18 | NaN | 0.47 | 0.29 | 0.32 | NaN |
| psma4 | psma4 | -2.48 | 0 | -2.86 | 0 | NaN | NaN | NaN | NaN | -0.01 | NaN | NaN | NaN |
| psma7-b;psma7-a | psma7;psma7 | -0.42 | 0.32 | -0.18 | NaN | -0.55 | 0.18 | -0.09 | NaN | -0.87 | 0.07 | -0.51 | 0.26 |
| psmb1 | psmb1 | 0.08 | NaN | -0.03 | NaN | -0.85 | 0.07 | 0.55 | 0.25 | -0.8 | 0.16 | 0.9 | NaN |
| psmb2 | psmb2 | NaN | NaN | NaN | NaN | 0.3 | NaN | -0.49 | NaN | 1.36 | 0.02 | -0.45 | 0.41 |
| psmc3-a;psmc3-b | nthl1;psmc3 | NaN | NaN | NaN | NaN | -1.12 | 0.04 | NaN | NaN | NaN | NaN | NaN | NaN |
| ptma-b | ptma | NaN | NaN | NaN | NaN | -1.13 | 0.09 | 0.86 | 0.16 | NaN | NaN | NaN | NaN |
| rab5a | rab5a;rab5c | -1.86 | 0 | NaN | NaN | -0.01 | NaN | -0.85 | 0.16 | NaN | NaN | NaN | NaN |
| ran | ran;ran | 1.12 | 0.02 | NaN | NaN | 1.35 | 0.04 | -0.05 | NaN | NaN | NaN | NaN | NaN |
| rbbp4-b | rbbp4 | NaN | NaN | NaN | NaN | 1.5 | 0.03 | -0.22 | NaN | NaN | NaN | NaN | NaN |
| rbm8a-b | rbm8a | NaN | NaN | NaN | NaN | 2.45 | 0 | NaN | NaN | NaN | NaN | NaN | NaN |
| RG | | 1.71 | 0 | 1.62 | 0 | -0.82 | 0.08 | -0.05 | NaN | NaN | NaN | NaN | NaN |
| rpl18;rpl18-b | rpl18 | 1.34 | 0.03 | 2.38 | 0 | NaN | NaN | NaN | NaN | -0.4 | NaN | NaN | NaN |
| rpl27 | rpl27 | NaN | NaN | NaN | NaN | NaN | NaN | NaN | NaN | 1.35 | 0.02 | NaN | NaN |

| | | | | | | | | | | | | | |
|---|---|---|---|---|---|---|---|---|---|---|---|---|---|
| rpl31 | rpl31 | NaN | NaN | NaN | NaN | 1.95 | 0 | 0.87 | 0.16 | NaN | NaN | NaN | NaN |
| rpl4-a | rpl4 | -0.15 | NaN | -0.22 | NaN | -0.89 | 0.1 | -0.4 | 0.37 | NaN | NaN | NaN | NaN |
| rpl5-a | rpl5 | 1.25 | 0.05 | 1.08 | 0.03 | 0.5 | NaN | -0.33 | NaN | NaN | NaN | NaN | NaN |
| rplp0 | rplp0 | 0.25 | NaN | 0.53 | 0.3 | 1.52 | 0.03 | 0.95 | NaN | NaN | NaN | NaN | NaN |
| rps11 | rps11 | NaN | NaN | NaN | NaN | 2.2 | 0 | NaN | NaN | NaN | NaN | NaN | NaN |
| rps14 | rps14 | 1.37 | 0.01 | 1.29 | 0.01 | NaN | NaN | NaN | NaN | NaN | NaN | NaN | NaN |
| rps16;MGC80065 | rps16 | 1.81 | 0.01 | NaN | NaN | -0.14 | NaN | NaN | NaN | NaN | NaN | NaN | NaN |
| rps4 | rps4x;rps4x | 3.2 | NaN | 5.32 | 0 | 0.22 | NaN | -0.51 | 0.26 | NaN | NaN | NaN | NaN |
| rpsa | rpsa | 0.42 | 0.32 | 0.69 | 0.12 | 0.31 | NaN | 1.27 | 0.11 | 1.32 | 0 | 0.54 | 0.22 |
| RRM1 | rrm1 | -0.62 | 0.24 | 1.29 | 0 | 0.2 | NaN | 0.14 | NaN | NaN | NaN | NaN | NaN |
| rtn3-a | rtn3 | 2.1 | 0 | 0.94 | 0.18 | 0.28 | NaN | -1.13 | 0.13 | -0.79 | NaN | NaN | NaN |
| RTN4 | rtn4 | 0.08 | NaN | -1.06 | 0.06 | -0.12 | NaN | -0.04 | NaN | NaN | NaN | NaN | NaN |
| sars | sars | NaN | NaN | NaN | NaN | -2.23 | 0 | NaN | NaN | NaN | NaN | NaN | NaN |
| sdha-a | sdha | -0.96 | NaN | -0.01 | NaN | -0.79 | 0.09 | -0.64 | 0.29 | NaN | NaN | NaN | NaN |
| serpin I1;LOC734183;MGC84260 | serpini1;serpini1 | NaN | NaN | NaN | NaN | 1.11 | 0.09 | NaN | NaN | NaN | NaN | NaN | NaN |
| serpinb6 | serpinb6 | 0.08 | NaN | 0.18 | NaN | 1.06 | 0.03 | 0.73 | 0.2 | NaN | NaN | NaN | NaN |
| Sfrs1 | | -0.66 | 0.32 | 0.58 | 0.28 | 1.14 | 0.04 | 0.56 | 0.18 | NaN | NaN | NaN | NaN |
| sgne1 | | 0.43 | 0.32 | 2.25 | NaN | 1.92 | NaN | NaN | NaN | 1.12 | 0.02 | 1.27 | 0.19 |
| skp1 | skp1 | -0.61 | 0.25 | -1.61 | 0 | -0.68 | 0.19 | -1.11 | NaN | 0.87 | 0.13 | NaN | NaN |
| sod1-a | sod1 | -0.27 | NaN | NaN | NaN | 0.34 | NaN | NaN | NaN | 0.94 | 0.05 | NaN | NaN |
| srsf2 | srsf2 | -1.31 | NaN | -1.21 | 0.01 | 0.36 | NaN | 0.28 | NaN | NaN | NaN | NaN | NaN |
| st3Gal-VI | st3gal6 | -0.22 | NaN | -1.25 | 0.01 | -0.24 | NaN | -0.05 | NaN | NaN | NaN | NaN | NaN |
| stip1 | stip1 | 1.35 | 0.01 | 0.35 | NaN | 0.13 | NaN | -0.02 | NaN | 0.59 | 0.28 | NaN | NaN |
| syp | syp | -0.03 | NaN | 0.48 | 0.26 | -0.69 | 0.1 | -0.92 | 0.18 | 1.06 | 0.03 | 0.94 | 0.26 |
| tagln2 | tagln2 | 1.05 | 0.09 | NaN | NaN | NaN | NaN | NaN | NaN | NaN | NaN | NaN | NaN |
| thbs1;LOC779026 | thbs1 | -0.67 | 0.21 | -0.96 | 0.03 | -0.6 | 0.19 | 0.27 | NaN | NaN | NaN | NaN | NaN |
| thbs4-prov | thbs4 | NaN | NaN | -1.09 | 0.12 | -1.45 | 0 | -0.12 | NaN | NaN | NaN | NaN | NaN |
| tuba3c | tuba3c | -0.78 | 0.11 | -0.95 | 0.03 | -0.46 | 0.24 | NaN | NaN | -1.56 | 0 | -1.86 | 0.12 |
| tubb | tubb | 0.17 | NaN | 1.17 | 0.02 | 1.09 | NaN | 0.21 | NaN | 0.66 | 0.16 | 1.08 | 0.19 |
| tubb4a | tubb4a | -0.3 | NaN | -1.11 | 0.02 | -0.21 | NaN | 0.59 | 0.18 | -0.08 | NaN | -0.65 | NaN |
| vamp2-a;vamp2;LOC495063;xsybi | vamp3;vamp2;vamp2;vamp1 | 1.4 | NaN | 1.22 | NaN | 0.9 | 0.06 | 0.76 | 0.14 | 0.87 | 0.07 | 1.1 | 0.2 |
| vdac1 | vdac1 | 0.92 | 0.22 | NaN | NaN | -1.44 | 0 | -0.11 | NaN | NaN | NaN | NaN | NaN |
| vdac2 | vdac2 | 1.64 | 0 | 1.42 | NaN | 0.31 | NaN | 1.86 | NaN | NaN | NaN | NaN | NaN |
| vha55 | atp6v1b2 | 0.45 | 0.3 | 1.49 | 0 | 0.53 | 0.22 | 0.83 | 0.16 | NaN | NaN | NaN | NaN |
| Xcad-11 | cdh11 | -0.2 | NaN | 0.88 | 0.12 | 1 | 0.06 | 0.54 | 0.25 | NaN | NaN | NaN | NaN |
| XDRP1 | ubqln4 | NaN | NaN | 1.3 | 0.06 | NaN | NaN | NaN | NaN | NaN | NaN | NaN | NaN |
| xFRP | | 0.9 | 0.06 | 0.29 | NaN | 0.36 | NaN | 0.21 | NaN | -0.46 | 0.29 | 0.08 | NaN |
| XPTP-LAR | ptprf | a | 0.27 | 1.82 | NaN | 0.35 | NaN | NaN | NaN | 1.01 | 0.09 | NaN | NaN |
| xrpn10;Xrpn10c | psmd4 | 1.87 | 0 | 0.28 | NaN | -0.17 | NaN | 0.19 | NaN | NaN | NaN | NaN | NaN |
| yes;MGC115231;yes1 | yes1 | -0.09 | NaN | NaN | NaN | 0.97 | 0.04 | 1.54 | 0.11 | NaN | NaN | NaN | NaN |
| | hist1h4a | 0.66 | 0.17 | 0.63 | NaN | -0.21 | NaN | -0.14 | NaN | 1.13 | 0.06 | 0.91 | 0.19 |
| | nme2 | NaN | NaN | 1.76 | 0 | -0.54 | NaN | 0.46 | 0.25 | -1.4 | 0.02 | NaN | NaN |
| | rbp4 | 0.55 | 0.27 | -1.35 | 0.01 | -1.82 | 0 | -1.23 | 0.13 | NaN | NaN | NaN | NaN |
| | apoa1 | 0.44 | 0.31 | 0.07 | NaN | -0.33 | NaN | -0.1 | NaN | -1.6 | 0 | -0.6 | 0.22 |
| | hnrnpc | -4.73 | NaN | NaN | NaN | -3.37 | 0 | NaN | NaN | NaN | NaN | NaN | NaN |

### Enriched_GOs

GO terms enriched in gene groups whose translation was signifiacntly regulatd by cue stimulations

Regulated_by_any_cue: p-value (DAVID) for enrichment of GO in genes whose translation was signifiacntly regulatd by any of three different cues

Netrin1_up: p-value (DAVID) for enrichment of GO in genes whose translation was signifiacntly increased by Netrin-1 stimulation

| Term | Regulated_by_any_cue | Netrin1_up | Netrin1_down | BDNF_up | BDNF_down | Sema_up | Sema_down |
|---|---|---|---|---|---|---|---|
| GO:0004298~threonine-type endopeptidase activity | 1.E-07 | 2.E-01 | 1.E+00 | 1.E-01 | 1.E-01 | 8.E-02 | 7.E-02 |
| GO:0006511~ubiquitin-dependent protein catabolic process | 5.E-07 | 4.E-01 | 3.E-01 | 5.E-03 | 3.E-01 | 1.E+00 | 2.E-01 |
| GO:0005840~ribosome | 7.E-07 | 1.E-04 | 1.E+00 | 2.E-02 | 1.E-02 | 1.E+00 | 1.E+00 |
| GO:0019773~proteasome core complex, alpha-subunit complex | 4.E-06 | 1.E+00 | 1.E+00 | 6.E-02 | 1.E+00 | 1.E+00 | 1.E+00 |
| GO:0005737~cytoplasm | 1.E-05 | 2.E-01 | 1.E-02 | 4.E-02 | 3.E-02 | 3.E-01 | 5.E-03 |
| GO:0003735~structural constituent of ribosome | 7.E-05 | 1.E-03 | 1.E+00 | 2.E-01 | 5.E-02 | 4.E-01 | 1.E+00 |
| GO:0006412~translation | 8.E-05 | 8.E-04 | 1.E+00 | 2.E-01 | 1.E-01 | 3.E-01 | 1.E+00 |
| GO:0007017~microtubule-based process | 1.E-04 | 2.E-01 | 2.E-02 | - | 1.E-01 | - | 4.E-03 |
| GO:0005839~proteasome core complex | 3.E-04 | 1.E+00 | - | 1.E+00 | 1.E+00 | 1.E+00 | 4.E-02 |
| GO:0006457~protein folding | 3.E-04 | 4.E-01 | 6.E-02 | 3.E-01 | 3.E-01 | - | - |
| GO:0005200~structural constituent of cytoskeleton | 5.E-04 | 2.E-01 | 2.E-01 | - | 1.E-01 | - | 3.E-03 |
| GO:0015991~ATP hydrolysis coupled proton transport | 7.E-04 | 2.E-02 | 1.E+00 | - | - | 7.E-02 | - |
| GO:0003924~GTPase activity | 2.E-03 | 5.E-02 | 2.E-01 | 4.E-01 | 4.E-01 | - | 4.E-02 |
| GO:0005509~calcium ion binding | 3.E-03 | 8.E-03 | 6.E-01 | 8.E-02 | 1.E+00 | 6.E-01 | 6.E-01 |
| GO:0005525~GTP binding | 1.E-02 | 4.E-01 | 4.E-03 | 2.E-01 | 2.E-01 | 1.E+00 | 3.E-01 |
| GO:0003755~peptidyl-prolyl cis-trans isomerase activity | 2.E-02 | 1.E+00 | - | 1.E-01 | 1.E+00 | - | - |
| GO:0005856~cytoskeleton | 3.E-02 | 5.E-01 | 1.E-01 | - | 3.E-01 | - | 2.E-01 |
| GO:0051603~proteolysis involved in cellular protein catabolic process | 3.E-02 | 1.E+00 | - | - | 1.E+00 | 1.E+00 | 1.E+00 |
| GO:0005215~transporter activity | 4.E-02 | 1.E+00 | 8.E-02 | - | 5.E-02 | 2.E-02 | - |
| GO:0005874~microtubule | 4.E-02 | 5.E-01 | 1.E-01 | - | 4.E-01 | - | 3.E-02 |
| GO:0008289~lipid binding | 5.E-02 | 1.E+00 | 3.E-01 | - | - | 1.E+00 | 1.E+00 |
| GO:0007155~cell adhesion | 5.E-02 | 1.E+00 | 1.E+00 | 1.E+00 | 2.E-01 | 1.E+00 | 1.E+00 |
| GO:0000166~nucleotide binding | 6.E-02 | 3.E-01 | 5.E-01 | 1.E-01 | 1.E+00 | - | - |
| GO:0005198~structural molecule activity | 6.E-02 | 6.E-01 | 3.E-02 | - | 4.E-01 | - | 1.E+00 |
| GO:0009409~response to cold | 7.E-02 | 1.E+00 | 1.E+00 | - | 1.E+00 | 1.E+00 | - |
| GO:0007016~cytoskeletal anchoring at plasma membrane | 7.E-02 | - | 1.E+00 | 1.E+00 | - | - | - |
| GO:0046961~proton-transporting ATPase activity, rotational mechanism | 8.E-02 | 1.E-01 | 1.E+00 | - | - | - | - |
| GO:0004075~biotin carboxylase activity | 8.E-02 | - | 1.E+00 | 1.E+00 | - | - | - |
| GO:0008308~voltage-gated anion channel activity | 8.E-02 | 1.E+00 | - | - | 1.E+00 | - | - |

| | | | | | | | |
|---|---|---|---|---|---|---|---|
| GO:0005743~mitochondrial inner membrane | 9.E-02 | 1.E+00 | 1.E+00 | 1.E+00 | 3.E-01 | - | - |
| GO:0030042~actin filament depolymerization | 1.E-01 | - | - | - | - | 1.E+00 | 1.E+00 |
| GO:0050660~flavin adenine dinucleotide binding | 1.E-01 | 1.E+00 | 1.E+00 | 1.E+00 | 1.E+00 | 1.E+00 | - |
| GO:0003746~translation elongation factor activity | 1.E-01 | - | 1.E+00 | 1.E-01 | - | - | - |
| GO:0005501~retinoid binding | 1.E-01 | - | 3.E-02 | - | 2.E-02 | 1.E+00 | - |
| GO:0016820~hydrolase activity, acting on acid anhydrides, catalyzing transmembrane movement of substances | 1.E-01 | 3.E-02 | - | - | - | - | - |
| GO:0005576~extracellular region | 1.E-01 | 1.E-01 | 8.E-01 | 8.E-01 | 7.E-01 | 7.E-01 | 2.E-01 |
| GO:0030141~secretory granule | 1.E-01 | - | - | - | - | 2.E-02 | - |
| GO:0097433~dense body | 1.E-01 | - | 3.E-02 | - | - | - | - |
| GO:0042176~regulation of protein catabolic process | 1.E-01 | - | - | 1.E+00 | - | 1.E+00 | - |
| GO:0015992~proton transport | 1.E-01 | 1.E+00 | - | - | 1.E+00 | - | - |
| GO:0007399~nervous system development | 1.E-01 | 4.E-01 | 4.E-01 | - | - | - | 1.E+00 |
| GO:0008201~heparin binding | 1.E-01 | 1.E+00 | - | - | - | - | 7.E-02 |
| GO:0046914~transition metal ion binding | 1.E-01 | 1.E+00 | - | - | - | - | 1.E+00 |
| GO:0045454~cell redox homeostasis | 1.E-01 | 1.E+00 | 3.E-01 | - | 1.E+00 | - | - |
| GO:0045261~proton-transporting ATP synthase complex, catalytic core F(1) | 1.E-01 | 1.E+00 | - | - | - | 1.E+00 | - |
| GO:0005911~cell-cell junction | 1.E-01 | 1.E+00 | 1.E+00 | - | 1.E+00 | - | 1.E+00 |
| GO:0016192~vesicle-mediated transport | 1.E-01 | 5.E-01 | 1.E+00 | 1.E+00 | 3.E-01 | 1.E+00 | 1.E+00 |
| GO:0016579~protein deubiquitination | 2.E-01 | - | - | 1.E-01 | 1.E+00 | - | - |
| GO:0051015~actin filament binding | 2.E-01 | - | - | - | - | 1.E+00 | 1.E+00 |
| GO:0022604~regulation of cell morphogenesis | 2.E-01 | - | - | - | - | 1.E+00 | 1.E+00 |
| GO:0007264~small GTPase mediated signal transduction | 2.E-01 | 7.E-01 | 2.E-02 | 6.E-01 | 6.E-01 | 1.E+00 | - |
| GO:0003756~protein disulfide isomerase activity | 2.E-01 | 1.E+00 | 1.E+00 | - | - | - | - |
| GO:0000502~proteasome complex | 2.E-01 | - | - | 1.E+00 | - | 1.E+00 | - |
| GO:0004843~thiol-dependent ubiquitin-specific protease activity | 2.E-01 | - | - | 1.E+00 | 1.E+00 | - | 1.E+00 |
| GO:0005623~cell | 2.E-01 | - | 6.E-02 | - | 1.E+00 | - | - |
| GO:0030234~enzyme regulator activity | 2.E-01 | - | - | 1.E+00 | - | 1.E+00 | - |
| GO:0046034~ATP metabolic process | 2.E-01 | 7.E-02 | - | - | - | - | - |
| GO:0005581~collagen trimer | 2.E-01 | - | 6.E-02 | - | 1.E+00 | - | - |
| GO:0016363~nuclear matrix | 2.E-01 | - | - | - | - | 1.E+00 | 1.E+00 |
| GO:0010628~positive regulation of gene expression | 2.E-01 | - | 1.E+00 | - | - | - | 1.E+00 |
| GO:0042742~defense response to bacterium | 2.E-01 | 1.E+00 | - | - | - | - | 1.E+00 |
| GO:0006094~gluconeogenesis | 2.E-01 | - | - | 1.E+00 | 1.E+00 | - | - |
| GO:0046933~proton-transporting ATP synthase activity, rotational mechanism | 2.E-01 | 1.E+00 | - | - | - | 1.E+00 | - |
| GO:0019898~extrinsic component of membrane | 3.E-01 | - | 1.E+00 | - | 1.E+00 | - | - |
| GO:0033180~proton-transporting V-type ATPase, V1 domain | 3.E-01 | 8.E-02 | - | - | - | - | - |
| GO:0015629~actin cytoskeleton | 3.E-01 | - | - | - | - | 1.E+00 | 1.E+00 |
| GO:0030246~carbohydrate binding | 3.E-01 | 1.E+00 | 1.E+00 | 1.E+00 | 1.E+00 | - | 1.E+00 |
| GO:0006886~intracellular protein transport | 3.E-01 | 1.E+00 | 1.E+00 | 1.E+00 | 4.E-01 | 1.E+00 | 1.E+00 |
| GO:0005833~hemoglobin complex | 3.E-01 | 9.E-02 | - | 1.E+00 | - | 1.E+00 | - |
| GO:0000910~cytokinesis | 3.E-01 | - | - | - | - | 1.E+00 | 1.E+00 |
| GO:0003995~acyl-CoA dehydrogenase activity | 3.E-01 | 1.E+00 | - | 1.E+00 | - | 1.E+00 | - |
| GO:0005925~focal adhesion | 3.E-01 | - | 1.E-01 | - | - | - | - |
| GO:0008601~protein phosphatase type 2A regulator activity | 3.E-01 | 1.E-01 | - | - | - | - | - |
| GO:0030529~intracellular ribonucleoprotein complex | 3.E-01 | - | 1.E+00 | 2.E-01 | - | - | - |
| GO:0070469~respiratory chain | 3.E-01 | - | 1.E+00 | 1.E+00 | - | - | - |

| | | | | | | | |
|---|---|---|---|---|---|---|---|
| GO:0005578~proteinaceous extracellular matrix | 4.E-01 | - | 3.E-01 | 1.E+00 | 1.E+00 | - | - |
| GO:0005344~oxygen transporter activity | 4.E-01 | 1.E-01 | - | 1.E+00 | - | 1.E+00 | - |
| GO:0050661~NADP binding | 4.E-01 | - | - | - | 1.E+00 | - | 1.E+00 |
| GO:0004867~serine-type endopeptidase inhibitor activity | 4.E-01 | 1.E+00 | - | 1.E+00 | - | - | - |
| GO:0004190~aspartic-type endopeptidase activity | 4.E-01 | - | 1.E+00 | - | 1.E+00 | - | 1.E+00 |
| GO:0019825~oxygen binding | 4.E-01 | 1.E-01 | - | 1.E+00 | - | 1.E+00 | - |
| GO:0006869~lipid transport | 4.E-01 | - | 1.E+00 | - | - | - | 1.E+00 |
| GO:0001558~regulation of cell growth | 4.E-01 | 1.E+00 | 1.E+00 | - | - | - | - |
| GO:0030145~manganese ion binding | 4.E-01 | - | 1.E+00 | - | 1.E+00 | - | - |
| GO:0019843~rRNA binding | 4.E-01 | 1.E+00 | - | 1.E+00 | - | - | - |
| GO:0005938~cell cortex | 4.E-01 | - | - | - | - | 1.E+00 | 1.E+00 |
| GO:0015986~ATP synthesis coupled proton transport | 4.E-01 | 1.E+00 | - | - | - | 1.E+00 | - |
| GO:0000287~magnesium ion binding | 4.E-01 | 1.E+00 | 1.E+00 | 4.E-01 | 1.E+00 | - | - |
| GO:0008237~metallopeptidase activity | 4.E-01 | 1.E+00 | - | - | - | - | 1.E+00 |
| GO:0007010~cytoskeleton organization | 4.E-01 | - | - | - | - | 1.E+00 | 1.E+00 |
| GO:0006099~tricarboxylic acid cycle | 4.E-01 | - | 1.E+00 | - | 1.E+00 | - | - |
| GO:0005544~calcium-dependent phospholipid binding | 4.E-01 | 1.E+00 | - | - | - | 1.E+00 | - |
| GO:0006096~glycolytic process | 5.E-01 | - | - | - | 1.E+00 | - | 1.E+00 |
| GO:0005789~endoplasmic reticulum membrane | 5.E-01 | 7.E-01 | 6.E-01 | 1.E+00 | - | 1.E+00 | - |
| GO:0005764~lysosome | 6.E-01 | - | 1.E+00 | - | - | - | 1.E+00 |
| GO:0030496~midbody | 6.E-01 | - | - | - | - | 1.E+00 | 1.E+00 |
| GO:0003729~mRNA binding | 6.E-01 | - | 1.E+00 | 1.E+00 | - | - | - |
| GO:0005524~ATP binding | 6.E-01 | 3.E-01 | 7.E-01 | 5.E-01 | 1.E+00 | 1.E+00 | 9.E-01 |
| GO:0005975~carbohydrate metabolic process | 6.E-01 | 1.E+00 | 1.E+00 | 3.E-01 | - | 1.E+00 | - |
| GO:0008152~metabolic process | 6.E-01 | - | - | 1.E+00 | 1.E+00 | - | 1.E+00 |
| GO:0000786~nucleosome | 6.E-01 | - | - | 1.E+00 | - | 1.E-01 | - |
| GO:0005741~mitochondrial outer membrane | 6.E-01 | 1.E+00 | - | - | 1.E+00 | - | - |
| GO:0005815~microtubule organizing center | 6.E-01 | - | 1.E+00 | 1.E+00 | - | - | - |
| GO:0005829~cytosol | 6.E-01 | 1.E+00 | - | 2.E-01 | - | - | - |
| GO:0005886~plasma membrane | 7.E-01 | 1.E+00 | 5.E-01 | 8.E-01 | - | 1.E+00 | - |
| GO:0003676~nucleic acid binding | 7.E-01 | 5.E-01 | 1.E+00 | 8.E-01 | 1.E+00 | 1.E+00 | - |
| GO:0016569~covalent chromatin modification | 7.E-01 | - | 1.E+00 | 1.E+00 | - | - | - |
| GO:0005654~nucleoplasm | 7.E-01 | - | 1.E+00 | - | - | 1.E+00 | - |
| GO:0004252~serine-type endopeptidase activity | 7.E-01 | 1.E-01 | - | - | - | - | 1.E+00 |
| GO:0005615~extracellular space | 7.E-01 | 1.E+00 | - | 4.E-01 | 1.E+00 | 1.E+00 | - |
| GO:0006810~transport | 8.E-01 | - | 3.E-01 | - | - | - | - |
| GO:0006260~DNA replication | 8.E-01 | 1.E+00 | - | 1.E+00 | - | - | - |
| GO:0005794~Golgi apparatus | 8.E-01 | 1.E+00 | 1.E+00 | 1.E+00 | - | - | - |
| GO:0006281~DNA repair | 9.E-01 | - | 1.E+00 | 1.E+00 | - | 1.E+00 | - |
| GO:0005783~endoplasmic reticulum | 9.E-01 | 4.E-01 | - | - | - | - | - |
| GO:0016874~ligase activity | 9.E-01 | - | - | - | 3.E-01 | - | - |
| GO:0005622~intracellular | 9.E-01 | - | 3.E-01 | 1.E+00 | 5.E-01 | 1.E+00 | 1.E+00 |
| GO:0016020~membrane | 9.E-01 | 1.E+00 | - | - | 1.E+00 | 1.E+00 | 3.E-01 |
| GO:0016567~protein ubiquitination | 9.E-01 | - | 4.E-01 | 1.E+00 | - | - | - |
| GO:0016055~Wnt signaling pathway | 9.E-01 | 5.E-01 | - | - | 1.E+00 | - | 1.E+00 |
| GO:0000139~Golgi membrane | 9.E-01 | 1.E+00 | - | - | 1.E+00 | - | - |

| | | | | | | | |
|---|---|---|---|---|---|---|---|
| GO:0003723~RNA binding | 9.E-01 | - | 8.E-01 | 8.E-01 | 1.E+00 | 1.E+00 | - |
| GO:0020037~heme binding | 9.E-01 | 6.E-01 | - | 1.E+00 | - | 1.E+00 | - |
| GO:0030154~cell differentiation | 9.E-01 | 1.E+00 | 1.E+00 | - | - | - | - |
| GO:0005506~iron ion binding | 9.E-01 | 6.E-01 | - | 1.E+00 | - | 1.E+00 | - |
| GO:0005634~nucleus | 1.E+00 | 1.E+00 | 1.E+00 | 8.E-01 | 9.E-01 | 6.E-01 | 1.E+00 |
| GO:0003677~DNA binding | 1.E+00 | - | 1.E+00 | 8.E-01 | - | 9.E-01 | - |
| GO:0046872~metal ion binding | 1.E+00 | 1.E+00 | 1.E+00 | 1.E+00 | 1.E+00 | 1.E+00 | 1.E+00 |
| GO:0016021~integral component of membrane | 1.E+00 | 1.E+00 | 1.E+00 | 1.E+00 | 1.E+00 | 9.E-01 | 1.E+00 |
| GO:0008270~zinc ion binding | 1.E+00 | 1.E+00 | - | 1.E+00 | - | - | - |
| GO:0006351~transcription, DNA-templated | 1.E+00 | 1.E+00 | - | 1.E+00 | 1.E+00 | - | 1.E+00 |
| GO:0006452~translational frameshifting | 1.E+00 | - | 1.E+00 | - | - | - | - |
| GO:0045901~positive regulation of translational elongation | 1.E+00 | - | 1.E+00 | - | - | - | - |
| GO:0050731~positive regulation of peptidyl-tyrosine phosphorylation | 1.E+00 | 1.E+00 | - | - | - | - | - |
| GO:0019430~removal of superoxide radicals | 1.E+00 | - | 1.E+00 | - | - | - | - |
| GO:0045087~innate immune response | 1.E+00 | - | - | 1.E+00 | - | - | - |
| GO:0034063~stress granule assembly | 1.E+00 | - | 1.E+00 | - | - | - | - |
| GO:0000132~establishment of mitotic spindle orientation | 1.E+00 | - | 1.E+00 | - | - | - | - |
| GO:0006414~translational elongation | 1.E+00 | - | - | - | 1.E+00 | - | - |
| GO:0050819~negative regulation of coagulation | 1.E+00 | 1.E+00 | - | - | - | - | - |
| GO:0007528~neuromuscular junction development | 1.E+00 | - | - | - | - | - | 1.E+00 |
| GO:0006796~phosphate-containing compound metabolic process | 1.E+00 | - | 1.E+00 | - | - | - | - |
| GO:0048255~mRNA stabilization | 1.E+00 | - | 1.E+00 | - | - | - | - |
| GO:0019752~carboxylic acid metabolic process | 1.E+00 | 1.E+00 | - | 1.E+00 | - | 1.E+00 | - |
| GO:2000507~positive regulation of energy homeostasis | 1.E+00 | 1.E+00 | - | - | 1.E+00 | 1.E+00 | - |
| GO:0060212~negative regulation of nuclear-transcribed mRNA poly(A) tail shortening | 1.E+00 | - | 1.E+00 | - | - | - | - |
| GO:0048793~pronephros development | 1.E+00 | - | 1.E+00 | - | - | - | - |
| GO:0006334~nucleosome assembly | 1.E+00 | - | - | - | - | 1.E+00 | - |
| GO:0090131~mesenchyme migration | 1.E+00 | - | - | - | - | - | 1.E+00 |
| GO:0000055~ribosomal large subunit export from nucleus | 1.E+00 | 1.E+00 | - | 1.E+00 | - | - | - |
| GO:0006338~chromatin remodeling | 1.E+00 | - | - | 1.E+00 | - | - | - |
| GO:0051016~barbed-end actin filament capping | 1.E+00 | 1.E+00 | - | - | - | - | - |
| GO:0001502~cartilage condensation | 1.E+00 | 1.E+00 | - | - | 1.E+00 | - | 1.E+00 |
| GO:0008380~RNA splicing | 1.E+00 | - | - | 1.E+00 | - | - | - |
| GO:0048738~cardiac muscle tissue development | 1.E+00 | 1.E+00 | - | - | 1.E+00 | - | 1.E+00 |
| GO:0097320~membrane tubulation | 1.E+00 | - | - | - | 1.E+00 | - | - |
| GO:0030336~negative regulation of cell migration | 1.E+00 | 1.E+00 | - | - | 1.E+00 | - | 1.E+00 |
| GO:0006417~regulation of translation | 1.E+00 | - | - | 1.E+00 | - | - | - |
| GO:0060271~cilium morphogenesis | 1.E+00 | 1.E+00 | - | - | - | - | - |
| GO:0032456~endocytic recycling | 1.E+00 | 1.E+00 | - | - | - | - | - |
| GO:0006813~potassium ion transport | 1.E+00 | 1.E+00 | - | - | - | - | - |
| GO:0008284~positive regulation of cell proliferation | 1.E+00 | 1.E+00 | - | - | - | - | - |
| GO:0042773~ATP synthesis coupled electron transport | 1.E+00 | - | - | 1.E+00 | - | - | - |
| GO:0015031~protein transport | 1.E+00 | 1.E+00 | - | - | - | - | - |
| GO:0006006~glucose metabolic process | 1.E+00 | - | - | - | - | - | 1.E+00 |
| GO:0000028~ribosomal small subunit assembly | 1.E+00 | - | - | - | - | 1.E+00 | - |
| GO:0006606~protein import into nucleus | 1.E+00 | 1.E+00 | - | 1.E+00 | - | - | - |

| | | | | | | | |
|---|---|---|---|---|---|---|---|
| GO:0090336~positive regulation of brown fat cell differentiation | 1.E+00 | 1.E+00 | - | - | 1.E+00 | 1.E+00 | - |
| GO:0009948~anterior/posterior axis specification | 1.E+00 | 1.E+00 | - | - | 1.E+00 | - | 1.E+00 |
| GO:0006665~sphingolipid metabolic process | 1.E+00 | - | 1.E+00 | - | - | - | - |
| GO:0042147~retrograde transport, endosome to Golgi | 1.E+00 | - | - | 1.E+00 | - | - | - |
| GO:0014850~response to muscle activity | 1.E+00 | 1.E+00 | - | - | 1.E+00 | 1.E+00 | - |
| GO:0006461~protein complex assembly | 1.E+00 | 1.E+00 | - | - | - | - | - |
| GO:0000381~regulation of alternative mRNA splicing, via spliceosome | 1.E+00 | - | - | 1.E+00 | - | - | - |
| GO:0045893~positive regulation of transcription, DNA-templated | 1.E+00 | 1.E+00 | - | - | 1.E+00 | - | 1.E+00 |
| GO:0007049~cell cycle | 1.E+00 | - | - | 1.E+00 | - | - | - |
| GO:0045727~positive regulation of translation | 1.E+00 | - | 1.E+00 | - | - | - | - |
| GO:0007218~neuropeptide signaling pathway | 1.E+00 | - | - | - | - | 1.E+00 | - |
| GO:0022008~neurogenesis | 1.E+00 | - | 1.E+00 | - | - | - | - |
| GO:0006108~malate metabolic process | 1.E+00 | - | 1.E+00 | - | - | - | - |
| GO:0019370~leukotriene biosynthetic process | 1.E+00 | 1.E+00 | - | - | - | - | - |
| GO:0055114~oxidation-reduction process | 1.E+00 | - | 1.E+00 | - | - | - | - |
| GO:0009617~response to bacterium | 1.E+00 | - | - | - | - | - | 1.E+00 |
| GO:0009411~response to UV | 1.E+00 | - | 1.E+00 | - | - | - | - |
| GO:0060317~cardiac epithelial to mesenchymal transition | 1.E+00 | - | 1.E+00 | - | - | - | - |
| GO:0006955~immune response | 1.E+00 | - | - | - | - | - | 1.E+00 |
| GO:0006105~succinate metabolic process | 1.E+00 | - | - | - | 1.E+00 | - | - |
| GO:0000187~activation of MAPK activity | 1.E+00 | 1.E+00 | - | - | - | - | - |
| GO:0006491~N-glycan processing | 1.E+00 | 1.E+00 | - | - | - | - | - |
| GO:0006879~cellular iron ion homeostasis | 1.E+00 | - | - | - | 1.E+00 | - | - |
| GO:0044208~'de novo' AMP biosynthetic process | 1.E+00 | - | - | 1.E+00 | - | - | - |
| GO:0006189~'de novo' IMP biosynthetic process | 1.E+00 | 1.E+00 | - | - | - | - | - |
| GO:0016486~peptide hormone processing | 1.E+00 | - | - | - | - | 1.E+00 | - |
| GO:0016477~cell migration | 1.E+00 | - | 1.E+00 | - | - | - | - |
| GO:0010629~negative regulation of gene expression | 1.E+00 | - | 1.E+00 | - | - | - | - |
| GO:0045892~negative regulation of transcription, DNA-templated | 1.E+00 | 1.E+00 | - | - | 1.E+00 | - | 1.E+00 |
| GO:0042254~ribosome biogenesis | 1.E+00 | - | - | 1.E+00 | - | - | - |
| GO:0051781~positive regulation of cell division | 1.E+00 | - | - | - | - | - | 1.E+00 |
| GO:0007156~homophilic cell adhesion via plasma membrane adhesion molecules | 1.E+00 | - | - | 1.E+00 | - | - | - |
| GO:0050873~brown fat cell differentiation | 1.E+00 | 1.E+00 | - | - | 1.E+00 | 1.E+00 | - |
| GO:0006486~protein glycosylation | 1.E+00 | - | 1.E+00 | - | - | - | - |
| GO:0000184~nuclear-transcribed mRNA catabolic process, nonsense-mediated decay | 1.E+00 | - | - | 1.E+00 | - | - | - |
| GO:0045010~actin nucleation | 1.E+00 | 1.E+00 | - | - | - | - | - |
| GO:0006897~endocytosis | 1.E+00 | - | 1.E+00 | - | - | - | - |
| GO:0051028~mRNA transport | 1.E+00 | - | - | 1.E+00 | - | - | - |
| GO:0008203~cholesterol metabolic process | 1.E+00 | - | 1.E+00 | - | - | - | - |
| GO:0042157~lipoprotein metabolic process | 1.E+00 | - | - | - | - | - | 1.E+00 |
| GO:0031076~embryonic camera-type eye development | 1.E+00 | 1.E+00 | - | - | 1.E+00 | - | 1.E+00 |
| GO:0006956~complement activation | 1.E+00 | 1.E+00 | - | - | - | - | 1.E+00 |
| GO:0072004~kidney field specification | 1.E+00 | - | 1.E+00 | - | - | - | - |
| GO:0048168~regulation of neuronal synaptic plasticity | 1.E+00 | - | - | - | 1.E+00 | 1.E+00 | - |
| GO:0006542~glutamine biosynthetic process | 1.E+00 | 1.E+00 | - | 1.E+00 | - | - | - |
| GO:0043161~proteasome-mediated ubiquitin-dependent protein catabolic process | 1.E+00 | - | 1.E+00 | 1.E+00 | - | - | - |

| | | | | | | | |
|---|---|---|---|---|---|---|---|
| GO:0006090~pyruvate metabolic process | 1.E+00 | - | - | 1.E+00 | - | - | - |
| GO:0006814~sodium ion transport | 1.E+00 | 1.E+00 | - | - | - | - | - |
| GO:0022904~respiratory electron transport chain | 1.E+00 | - | - | - | 1.E+00 | - | - |
| GO:0006183~GTP biosynthetic process | 1.E+00 | 1.E+00 | - | - | - | - | 1.E+00 |
| GO:0010718~positive regulation of epithelial to mesenchymal transition | 1.E+00 | - | 1.E+00 | - | - | - | - |
| GO:0042026~protein refolding | 1.E+00 | - | - | 1.E+00 | - | - | - |
| GO:0032071~regulation of endodeoxyribonuclease activity | 1.E+00 | - | - | - | - | 1.E+00 | - |
| GO:0007015~actin filament organization | 1.E+00 | - | - | - | 1.E+00 | - | - |
| GO:0006396~RNA processing | 1.E+00 | 1.E+00 | - | - | - | - | - |
| GO:0006241~CTP biosynthetic process | 1.E+00 | 1.E+00 | - | - | - | - | 1.E+00 |
| GO:0007596~blood coagulation | 1.E+00 | 1.E+00 | - | - | - | - | - |
| GO:0032331~negative regulation of chondrocyte differentiation | 1.E+00 | 1.E+00 | - | - | 1.E+00 | - | 1.E+00 |
| GO:0001654~eye development | 1.E+00 | 1.E+00 | - | 1.E+00 | - | - | - |
| GO:0061037~negative regulation of cartilage development | 1.E+00 | 1.E+00 | - | - | 1.E+00 | - | 1.E+00 |
| GO:0000056~ribosomal small subunit export from nucleus | 1.E+00 | 1.E+00 | - | 1.E+00 | - | - | - |
| GO:0043691~reverse cholesterol transport | 1.E+00 | - | 1.E+00 | - | - | - | - |
| GO:0007165~signal transduction | 1.E+00 | - | - | 1.E+00 | - | - | - |
| GO:0042060~wound healing | 1.E+00 | 1.E+00 | - | - | 1.E+00 | - | 1.E+00 |
| GO:0007067~mitotic nuclear division | 1.E+00 | - | 1.E+00 | - | - | - | - |
| GO:0050728~negative regulation of inflammatory response | 1.E+00 | 1.E+00 | - | - | 1.E+00 | 1.E+00 | - |
| GO:0006098~pentose-phosphate shunt | 1.E+00 | - | - | 1.E+00 | - | - | - |
| GO:0006352~DNA-templated transcription, initiation | 1.E+00 | - | - | - | - | 1.E+00 | - |
| GO:0051301~cell division | 1.E+00 | - | 1.E+00 | - | - | - | - |
| GO:0006895~Golgi to endosome transport | 1.E+00 | 1.E+00 | - | - | - | - | - |
| GO:0002062~chondrocyte differentiation | 1.E+00 | 1.E+00 | - | - | 1.E+00 | - | 1.E+00 |
| GO:0046883~regulation of hormone secretion | 1.E+00 | - | - | - | - | 1.E+00 | - |
| GO:0006434~seryl-tRNA aminoacylation | 1.E+00 | - | - | - | 1.E+00 | - | - |
| GO:0048666~neuron development | 1.E+00 | - | - | - | 1.E+00 | - | - |
| GO:0006413~translational initiation | 1.E+00 | - | 1.E+00 | - | - | - | - |
| GO:0034333~adherens junction assembly | 1.E+00 | 1.E+00 | - | - | 1.E+00 | - | 1.E+00 |
| GO:0006086~acetyl-CoA biosynthetic process from pyruvate | 1.E+00 | 1.E+00 | - | - | - | - | - |
| GO:0008104~protein localization | 1.E+00 | 1.E+00 | - | - | - | - | - |
| GO:0007369~gastrulation | 1.E+00 | - | 1.E+00 | - | - | - | - |
| GO:0006446~regulation of translational initiation | 1.E+00 | - | 1.E+00 | - | - | - | - |
| GO:0030866~cortical actin cytoskeleton organization | 1.E+00 | - | 1.E+00 | - | - | - | - |
| GO:0090382~phagosome maturation | 1.E+00 | 1.E+00 | - | - | - | - | - |
| GO:0060699~regulation of endoribonuclease activity | 1.E+00 | - | - | - | - | 1.E+00 | - |
| GO:0006397~mRNA processing | 1.E+00 | - | - | 1.E+00 | - | - | - |
| GO:0019521~D-gluconate metabolic process | 1.E+00 | - | - | 1.E+00 | - | - | - |
| GO:0009790~embryo development | 1.E+00 | - | 1.E+00 | - | - | - | - |
| GO:0043433~negative regulation of sequence-specific DNA binding transcription factor activity | 1.E+00 | 1.E+00 | - | - | 1.E+00 | - | 1.E+00 |
| GO:0030163~protein catabolic process | 1.E+00 | - | - | - | 1.E+00 | - | - |
| GO:0001731~formation of translation preinitiation complex | 1.E+00 | - | 1.E+00 | - | - | - | - |
| GO:0003190~atrioventricular valve formation | 1.E+00 | - | 1.E+00 | - | - | - | - |
| GO:0006826~iron ion transport | 1.E+00 | - | - | - | 1.E+00 | - | - |
| GO:0006228~UTP biosynthetic process | 1.E+00 | 1.E+00 | - | - | - | - | 1.E+00 |

| | | | | | | | |
|---|---|---|---|---|---|---|---|
| GO:0031497~chromatin assembly | 1.E+00 | - | - | 1.E+00 | - | - | - |
| GO:0006355~regulation of transcription, DNA-templated | 1.E+00 | - | - | 1.E+00 | - | - | - |
| GO:0043486~histone exchange | 1.E+00 | - | 1.E+00 | - | - | - | - |
| GO:0051014~actin filament severing | 1.E+00 | 1.E+00 | - | - | - | - | - |
| GO:0043128~positive regulation of 1-phosphatidylinositol 4-kinase activity | 1.E+00 | 1.E+00 | - | - | - | - | - |
| GO:0031175~neuron projection development | 1.E+00 | - | - | 1.E+00 | - | - | - |
| GO:0030041~actin filament polymerization | 1.E+00 | - | - | - | 1.E+00 | - | - |
| GO:0045905~positive regulation of translational termination | 1.E+00 | - | 1.E+00 | - | - | - | - |
| GO:0031398~positive regulation of protein ubiquitination | 1.E+00 | 1.E+00 | - | - | 1.E+00 | - | 1.E+00 |
| GO:0005788~endoplasmic reticulum lumen | 1.E+00 | 1.E+00 | - | - | - | - | - |
| GO:0030132~clathrin coat of coated pit | 1.E+00 | 1.E+00 | - | - | 1.E+00 | - | 1.E+00 |
| GO:0030027~lamellipodium | 1.E+00 | - | - | - | - | - | 1.E+00 |
| GO:0033186~CAF-1 complex | 1.E+00 | - | - | 1.E+00 | - | - | - |
| GO:0000159~protein phosphatase type 2A complex | 1.E+00 | 1.E+00 | - | - | - | - | - |
| GO:0043231~intracellular membrane-bounded organelle | 1.E+00 | 1.E+00 | - | - | - | - | - |
| GO:0005882~intermediate filament | 1.E+00 | 1.E+00 | - | - | - | - | - |
| GO:0031465~Cul4B-RING E3 ubiquitin ligase complex | 1.E+00 | - | 1.E+00 | 1.E+00 | - | - | - |
| GO:0005905~clathrin-coated pit | 1.E+00 | - | 1.E+00 | - | - | - | - |
| GO:0005890~sodium:potassium-exchanging ATPase complex | 1.E+00 | 1.E+00 | - | - | - | - | - |
| GO:0019005~SCF ubiquitin ligase complex | 1.E+00 | - | 1.E+00 | - | - | - | - |
| GO:0030286~dynein complex | 1.E+00 | - | 1.E+00 | - | - | - | - |
| GO:0016607~nuclear speck | 1.E+00 | - | - | 1.E+00 | - | - | - |
| GO:0046930~pore complex | 1.E+00 | 1.E+00 | - | - | - | - | - |
| GO:0005802~trans-Golgi network | 1.E+00 | 1.E+00 | - | - | - | - | - |
| GO:0005852~eukaryotic translation initiation factor 3 complex | 1.E+00 | - | 1.E+00 | - | - | - | - |
| GO:0045178~basal part of cell | 1.E+00 | 1.E+00 | - | - | 1.E+00 | - | 1.E+00 |
| GO:0033018~sarcoplasmic reticulum lumen | 1.E+00 | 1.E+00 | - | 1.E+00 | - | - | - |
| GO:0000812~Swr1 complex | 1.E+00 | - | 1.E+00 | - | - | - | - |
| GO:0022624~proteasome accessory complex | 1.E+00 | - | - | - | 1.E+00 | - | - |
| GO:0008021~synaptic vesicle | 1.E+00 | - | - | - | 1.E+00 | 1.E+00 | - |
| GO:0033179~proton-transporting V-type ATPase, V0 domain | 1.E+00 | - | - | - | - | 1.E+00 | - |
| GO:0035098~ESC/E(Z) complex | 1.E+00 | - | - | 1.E+00 | - | - | - |
| GO:0042470~melanosome | 1.E+00 | 1.E+00 | - | 1.E+00 | - | - | - |
| GO:0005638~lamin filament | 1.E+00 | - | 1.E+00 | - | - | - | - |
| GO:0055037~recycling endosome | 1.E+00 | 1.E+00 | - | - | - | - | - |
| GO:0030175~filopodium | 1.E+00 | - | - | - | - | - | 1.E+00 |
| GO:0005579~membrane attack complex | 1.E+00 | - | - | - | - | - | 1.E+00 |
| GO:0016471~vacuolar proton-transporting V-type ATPase complex | 1.E+00 | 1.E+00 | - | - | - | - | - |
| GO:0005887~integral component of plasma membrane | 1.E+00 | 1.E+00 | - | - | - | - | - |
| GO:0044297~cell body | 1.E+00 | - | - | - | - | - | 1.E+00 |
| GO:0010494~cytoplasmic stress granule | 1.E+00 | - | 1.E+00 | - | - | - | - |
| GO:0034364~high-density lipoprotein particle | 1.E+00 | - | 1.E+00 | - | - | - | - |
| GO:0005853~eukaryotic translation elongation factor 1 complex | 1.E+00 | - | - | 1.E+00 | - | - | - |
| GO:0030130~clathrin coat of trans-Golgi network vesicle | 1.E+00 | 1.E+00 | - | - | 1.E+00 | - | 1.E+00 |
| GO:0097443~sorting endosome | 1.E+00 | - | 1.E+00 | - | - | - | - |
| GO:0000221~vacuolar proton-transporting V-type ATPase, V1 domain | 1.E+00 | - | 1.E+00 | - | - | - | - |

| | | | | | | | |
|---|---|---|---|---|---|---|---|
| GO:0005875~microtubule associated complex | 1.E+00 | - | 1.E+00 | - | - | - | - |
| GO:0030117~membrane coat | 1.E+00 | - | 1.E+00 | - | - | - | - |
| GO:0030904~retromer complex | 1.E+00 | - | - | 1.E+00 | - | - | - |
| GO:0005730~nucleolus | 1.E+00 | - | - | - | - | 1.E+00 | - |
| GO:0016010~dystrophin-associated glycoprotein complex | 1.E+00 | - | - | 1.E+00 | - | - | - |
| GO:0048471~perinuclear region of cytoplasm | 1.E+00 | - | - | 1.E+00 | - | - | - |
| GO:0031467~Cul7-RING ubiquitin ligase complex | 1.E+00 | - | 1.E+00 | - | - | - | - |
| GO:0033290~eukaryotic 48S preinitiation complex | 1.E+00 | - | 1.E+00 | - | - | - | - |
| GO:0080008~Cul4-RING E3 ubiquitin ligase complex | 1.E+00 | - | 1.E+00 | 1.E+00 | - | - | - |
| GO:0022627~cytosolic small ribosomal subunit | 1.E+00 | - | - | - | - | 1.E+00 | - |
| GO:0005637~nuclear inner membrane | 1.E+00 | - | 1.E+00 | - | - | - | - |
| GO:0030126~COPI vesicle coat | 1.E+00 | - | - | - | 1.E+00 | - | - |
| GO:0008540~proteasome regulatory particle, base subcomplex | 1.E+00 | 1.E+00 | - | - | - | - | - |
| GO:0016282~eukaryotic 43S preinitiation complex | 1.E+00 | - | 1.E+00 | - | - | - | - |
| GO:0031225~anchored component of membrane | 1.E+00 | 1.E+00 | - | - | - | - | - |
| GO:0005739~mitochondrion | 1.E+00 | - | - | - | - | - | 1.E+00 |
| GO:0005635~nuclear envelope | 1.E+00 | 1.E+00 | - | 1.E+00 | - | - | - |
| GO:0005871~kinesin complex | 1.E+00 | 1.E+00 | - | - | - | - | - |
| GO:0045335~phagocytic vesicle | 1.E+00 | 1.E+00 | - | - | - | - | - |
| GO:0031464~Cul4A-RING E3 ubiquitin ligase complex | 1.E+00 | - | 1.E+00 | 1.E+00 | - | - | - |
| GO:0034362~low-density lipoprotein particle | 1.E+00 | - | 1.E+00 | - | - | - | - |
| GO:0004802~transketolase activity | 1.E+00 | - | - | 1.E+00 | - | - | - |
| GO:0004364~glutathione transferase activity | 1.E+00 | - | - | - | 1.E+00 | - | - |
| GO:0004177~aminopeptidase activity | 1.E+00 | - | 1.E+00 | - | - | - | - |
| GO:0005528~FK506 binding | 1.E+00 | - | - | 1.E+00 | - | - | - |
| GO:0004463~leukotriene-A4 hydrolase activity | 1.E+00 | 1.E+00 | - | - | - | - | - |
| GO:0016671~oxidoreductase activity, acting on a sulfur group of donors, disulfide as acceptor | 1.E+00 | - | 1.E+00 | - | - | - | - |
| GO:0005179~hormone activity | 1.E+00 | 1.E+00 | - | - | 1.E+00 | 1.E+00 | - |
| GO:0051287~NAD binding | 1.E+00 | - | - | - | - | - | 1.E+00 |
| GO:0070181~small ribosomal subunit rRNA binding | 1.E+00 | - | 1.E+00 | - | - | - | - |
| GO:0016787~hydrolase activity | 1.E+00 | - | - | - | 1.E+00 | - | - |
| GO:0004784~superoxide dismutase activity | 1.E+00 | - | - | - | - | 1.E+00 | - |
| GO:0004616~phosphogluconate dehydrogenase (decarboxylating) activity | 1.E+00 | - | - | 1.E+00 | - | - | - |
| GO:0019899~enzyme binding | 1.E+00 | - | 1.E+00 | - | - | - | - |
| GO:0004639~phosphoribosylaminoimidazolesuccinocarboxamide synthase activity | 1.E+00 | 1.E+00 | - | - | - | - | - |
| GO:0003824~catalytic activity | 1.E+00 | - | - | - | - | - | 1.E+00 |
| GO:0005319~lipid transporter activity | 1.E+00 | - | - | - | 1.E+00 | - | - |
| GO:0005055~laminin receptor activity | 1.E+00 | - | - | - | - | 1.E+00 | - |
| GO:0004743~pyruvate kinase activity | 1.E+00 | 1.E+00 | - | 1.E+00 | - | - | - |
| GO:0005094~Rho GDP-dissociation inhibitor activity | 1.E+00 | 1.E+00 | - | - | 1.E+00 | - | - |
| GO:0035091~phosphatidylinositol binding | 1.E+00 | - | - | 1.E+00 | - | - | - |
| GO:0016805~dipeptidase activity | 1.E+00 | - | - | - | - | - | 1.E+00 |
| GO:0030060~L-malate dehydrogenase activity | 1.E+00 | - | 1.E+00 | - | - | - | - |
| GO:0004828~serine-tRNA ligase activity | 1.E+00 | - | - | - | 1.E+00 | - | - |
| GO:0015035~protein disulfide oxidoreductase activity | 1.E+00 | - | 1.E+00 | - | - | - | - |
| GO:0004553~hydrolase activity, hydrolyzing O-glycosyl compounds | 1.E+00 | - | - | 1.E+00 | - | - | - |

| | | | | | | | |
|---|---|---|---|---|---|---|---|
| GO:0015078~hydrogen ion transmembrane transporter activity | 1.E+00 | - | - | - | - | 1.E+00 | - |
| GO:0008199~ferric iron binding | 1.E+00 | - | - | - | 1.E+00 | - | - |
| GO:0009374~biotin binding | 1.E+00 | - | - | 1.E+00 | - | - | - |
| GO:0004459~L-lactate dehydrogenase activity | 1.E+00 | 1.E+00 | - | 1.E+00 | - | 1.E+00 | - |
| GO:0004869~cysteine-type endopeptidase inhibitor activity | 1.E+00 | - | - | - | 1.E+00 | - | - |
| GO:0016301~kinase activity | 1.E+00 | 1.E+00 | - | 1.E+00 | - | - | - |
| GO:0050135~NAD(P)+ nucleosidase activity | 1.E+00 | - | - | - | 1.E+00 | - | - |
| GO:0008379~thioredoxin peroxidase activity | 1.E+00 | - | 1.E+00 | - | - | - | - |
| GO:0016757~transferase activity, transferring glycosyl groups | 1.E+00 | - | 1.E+00 | - | - | - | - |
| GO:0004356~glutamate-ammonia ligase activity | 1.E+00 | 1.E+00 | - | 1.E+00 | - | - | - |
| GO:0043022~ribosome binding | 1.E+00 | - | 1.E+00 | - | - | - | - |
| GO:0004347~glucose-6-phosphate isomerase activity | 1.E+00 | - | - | - | 1.E+00 | - | - |
| GO:0004064~arylesterase activity | 1.E+00 | - | - | - | 1.E+00 | - | - |
| GO:0004019~adenylosuccinate synthase activity | 1.E+00 | - | - | 1.E+00 | - | - | - |
| GO:0045735~nutrient reservoir activity | 1.E+00 | - | - | - | 1.E+00 | - | - |
| GO:0008750~NAD(P)+ transhydrogenase (AB-specific) activity | 1.E+00 | - | - | - | 1.E+00 | - | - |
| GO:0004365~glyceraldehyde-3-phosphate dehydrogenase (NAD+) (phosphorylating) activity | 1.E+00 | - | - | - | - | - | 1.E+00 |
| GO:0003743~translation initiation factor activity | 1.E+00 | - | 1.E+00 | - | - | - | - |
| GO:0004722~protein serine/threonine phosphatase activity | 1.E+00 | - | - | - | 1.E+00 | - | - |
| GO:0008177~succinate dehydrogenase (ubiquinone) activity | 1.E+00 | - | - | - | 1.E+00 | - | - |
| GO:0008373~sialyltransferase activity | 1.E+00 | - | 1.E+00 | - | - | - | - |
| GO:0016620~oxidoreductase activity, acting on the aldehyde or oxo group of donors, NAD or NADP as acceptor | 1.E+00 | - | 1.E+00 | - | - | - | - |
| GO:0004748~ribonucleoside-diphosphate reductase activity, thioredoxin disulfide as acceptor | 1.E+00 | 1.E+00 | - | - | - | - | - |
| GO:0004791~thioredoxin-disulfide reductase activity | 1.E+00 | - | 1.E+00 | - | - | - | - |
| GO:0036094~small molecule binding | 1.E+00 | - | 1.E+00 | - | 1.E+00 | 1.E+00 | - |
| GO:0030371~translation repressor activity | 1.E+00 | - | 1.E+00 | - | - | - | - |
| GO:0003777~microtubule motor activity | 1.E+00 | 1.E+00 | - | - | - | - | - |
| GO:0004550~nucleoside diphosphate kinase activity | 1.E+00 | 1.E+00 | - | - | - | - | 1.E+00 |
| GO:0004725~protein tyrosine phosphatase activity | 1.E+00 | - | - | - | - | 1.E+00 | - |
| GO:0030955~potassium ion binding | 1.E+00 | 1.E+00 | - | 1.E+00 | - | - | - |
| GO:0003730~mRNA 3'-UTR binding | 1.E+00 | - | 1.E+00 | - | - | - | - |
| GO:0004715~non-membrane spanning protein tyrosine kinase activity | 1.E+00 | - | - | 1.E+00 | - | - | - |
| GO:0036459~thiol-dependent ubiquitinyl hydrolase activity | 1.E+00 | - | - | 1.E+00 | - | - | - |
| GO:0008097~5S rRNA binding | 1.E+00 | 1.E+00 | - | - | - | - | - |
| GO:0009055~electron carrier activity | 1.E+00 | - | 1.E+00 | - | - | - | - |
| GO:0043395~heparan sulfate proteoglycan binding | 1.E+00 | - | - | - | - | - | 1.E+00 |
| GO:0051082~unfolded protein binding | 1.E+00 | - | - | - | - | 1.E+00 | - |
| GO:0004857~enzyme inhibitor activity | 1.E+00 | - | - | - | - | 1.E+00 | - |
| GO:0004739~pyruvate dehydrogenase (acetyl-transferring) activity | 1.E+00 | 1.E+00 | - | - | - | - | - |
| GO:0008137~NADH dehydrogenase (ubiquinone) activity | 1.E+00 | - | - | 1.E+00 | - | - | - |
| GO:0004427~inorganic diphosphatase activity | 1.E+00 | - | 1.E+00 | - | - | - | - |
| GO:0003953~NAD+ nucleosidase activity | 1.E+00 | - | - | - | 1.E+00 | - | - |
| GO:0004736~pyruvate carboxylase activity | 1.E+00 | - | - | 1.E+00 | - | - | - |
| GO:0015288~porin activity | 1.E+00 | 1.E+00 | - | - | - | - | - |
| GO:0022891~substrate-specific transmembrane transporter activity | 1.E+00 | 1.E+00 | - | - | - | 1.E+00 | - |
| GO:0004129~cytochrome-c oxidase activity | 1.E+00 | - | - | - | - | - | 1.E+00 |

| | | | | | | | |
|---|---|---|---|---|---|---|---|
| GO:0005102~receptor binding | 1.E+00 | - | - | - | - | 1.E+00 | - |
| GO:0004871~signal transducer activity | 1.E+00 | 1.E+00 | - | - | 1.E+00 | - | 1.E+00 |
| GO:0008235~metalloexopeptidase activity | 1.E+00 | - | 1.E+00 | - | - | - | - |
| GO:0017127~cholesterol transporter activity | 1.E+00 | - | 1.E+00 | - | - | - | - |
| GO:0051537~2 iron, 2 sulfur cluster binding | 1.E+00 | 1.E+00 | - | - | - | - | - |
| GO:0042393~histone binding | 1.E+00 | - | 1.E+00 | - | - | - | - |
| GO:0005201~extracellular matrix structural constituent | 1.E+00 | - | 1.E+00 | - | 1.E+00 | - | - |
| GO:0016153~urocanate hydratase activity | 1.E+00 | - | - | - | - | - | 1.E+00 |

Axonal_RibosomeAssemblyFactors: All genes annotated with GO termes "nucleolus" or "ribosome biogenesis" that are detected in the axonal transcritome

| Xenbase_ID | mouse gene name | GO term "nucleolus" | GO term "ribosome biogenesis" | Xlaevis axonal transcriptome | Xlaevis axonal proteome | mouse axon translatome |
|---|---|---|---|---|---|---|
| XB-GENE-1000251 | hmgb2 | + | - | + | + | - |
| XB-GENE-1000491 | fbxw7 | + | - | + | - | + |
| XB-GENE-1000765 | ddx17 | + | + | + | - | - |
| XB-GENE-1000959 | vmp1 | + | - | + | - | - |
| XB-GENE-1000983 | rpl11 | + | + | + | + | + |
| XB-GENE-1001390 | tyms | + | - | - | + | - |
| XB-GENE-1001971 | rpl31 | + | - | + | + | - |
| XB-GENE-1002501 | nono | + | - | + | - | - |
| XB-GENE-1003294 | rps6 | + | + | + | + | - |
| XB-GENE-1003623 | rps2 | - | + | + | + | - |
| XB-GENE-1004641 | nup88 | - | + | + | - | - |
| XB-GENE-1005073 | ltv1 | - | + | + | - | - |
| XB-GENE-1006309 | rpl18 | + | - | + | + | + |
| XB-GENE-1007936 | rps21 | - | + | - | + | + |
| XB-GENE-1008656 | frg1 | + | + | + | - | - |
| XB-GENE-1008861 | rpl12 | + | + | + | + | - |
| XB-GENE-1008948 | dhx15 | + | - | - | + | - |
| XB-GENE-1010464 | snrpb2 | + | - | + | - | - |
| XB-GENE-1013231 | asna1 | + | - | - | + | - |
| XB-GENE-1013309 | uso1 | + | - | - | + | + |
| XB-GENE-1014730 | cask | + | - | - | + | - |
| XB-GENE-1015264 | ola1 | + | - | + | - | - |
| XB-GENE-1016064 | rpl35a | - | + | + | + | - |
| XB-GENE-1016064 | rpl35a | - | + | + | + | - |
| XB-GENE-1016264 | exosc7 | + | + | - | + | - |
| XB-GENE-1016377 | rps17 | - | + | + | - | + |
| XB-GENE-1016477 | mrps15 | + | - | + | - | - |
| XB-GENE-1188424 | sub1 | + | - | + | + | - |
| XB-GENE-17330372 | xrn2 | + | - | - | + | - |
| XB-GENE-17330417 | ube2n | + | - | + | + | - |
| XB-GENE-17330554 | ddx3x | - | + | + | + | - |
| XB-GENE-17331043 | kif2a | + | - | + | - | - |
| XB-GENE-17331209 | rpl7a | - | + | + | + | - |
| XB-GENE-17331340 | nuak1 | + | - | + | - | - |
| XB-GENE-17331553 | wbp11 | - | + | + | - | - |
| XB-GENE-17332277 | dctn3 | + | - | - | + | + |
| XB-GENE-17332836 | rpap2 | + | - | + | - | - |
| XB-GENE-17334360 | fcf1 | + | + | + | - | + |
| XB-GENE-17334580 | tsc1 | - | + | + | - | - |

| | | | | | | |
|---|---|---|---|---|---|---|
| XB-GENE-17334901 | smad7 | + | - | + | - | - |
| XB-GENE-17334929 | rbm4b | + | - | + | + | - |
| XB-GENE-17335149 | xpo1 | + | + | + | + | - |
| XB-GENE-17335914 | nhp2 | + | + | + | - | - |
| XB-GENE-17335931 | fbxo11 | + | - | + | - | - |
| XB-GENE-17337446 | rplp0 | - | + | + | + | - |
| XB-GENE-17337654 | aamp | - | + | - | + | + |
| XB-GENE-17338571 | dedd2 | + | - | + | - | - |
| XB-GENE-17339479 | cdc14b | + | - | + | - | - |
| XB-GENE-17339580 | plrg1 | + | - | + | - | - |
| XB-GENE-17339884 | pus1 | + | - | + | - | - |
| XB-GENE-17340555 | ddx5 | + | - | - | + | - |
| XB-GENE-17340776 | rps13 | + | - | + | + | - |
| XB-GENE-17340849 | srsf5 | + | - | + | + | - |
| XB-GENE-17341054 | parn | + | - | + | - | - |
| XB-GENE-17342307 | nucks1 | + | - | + | - | - |
| XB-GENE-17342370 | rai14 | + | - | + | - | - |
| XB-GENE-17343779 | jpt1 | + | - | + | - | - |
| XB-GENE-17343925 | edf1 | + | - | - | + | - |
| XB-GENE-17344173 | sub1 | + | - | + | + | - |
| XB-GENE-17344257 | ddx6 | + | - | + | + | + |
| XB-GENE-17344335 | rbm14 | + | - | + | + | - |
| XB-GENE-17344569 | sdha | + | - | - | + | + |
| XB-GENE-17346030 | abce1 | - | + | + | + | - |
| XB-GENE-17346538 | ddx23 | + | - | + | - | - |
| XB-GENE-492329 | tra2a | + | - | - | + | - |
| XB-GENE-492785 | cirbp | + | - | + | + | - |
| XB-GENE-493737 | eif3l | + | - | - | + | + |
| XB-GENE-5738730 | gnai3 | + | - | - | + | - |
| XB-GENE-5786746 | rps16 | - | + | + | - | - |
| XB-GENE-5800445 | zc3h15 | + | - | + | - | + |
| XB-GENE-5819322 | rpl14 | - | + | + | + | + |
| XB-GENE-5828312 | leo1 | + | - | + | - | + |
| XB-GENE-5833061 | rps15 | - | + | + | + | + |
| XB-GENE-5865181 | macrod2 | + | - | - | + | - |
| XB-GENE-5871876 | ckap5 | + | - | - | + | - |
| XB-GENE-5891407 | rpl23a | - | + | + | + | - |
| XB-GENE-5900967 | kdm7a | + | - | + | - | - |
| XB-GENE-5919816 | srp19 | + | - | - | + | - |
| XB-GENE-5929894 | acat2 | + | - | - | + | + |
| XB-GENE-5930659 | rps9 | + | - | + | + | + |
| XB-GENE-5935325 | rps8 | - | + | + | + | + |
| XB-GENE-5945253 | map2 | + | - | - | + | - |
| XB-GENE-5948460 | ppp1ca | + | - | + | - | - |
| XB-GENE-5952259 | rabgef1 | + | - | + | - | + |
| XB-GENE-6042311 | pim1 | + | - | + | - | - |

| | | | | | | |
|---|---|---|---|---|---|---|
| XB-GENE-6053022 | myc | + | - | + | - | - |
| XB-GENE-6068534 | gtpbp4 | + | + | + | - | - |
| XB-GENE-6077488 | slbp | + | - | + | - | - |
| XB-GENE-6078160 | bcas2 | + | - | - | + | + |
| XB-GENE-6251644 | timm13 | + | - | - | + | + |
| XB-GENE-6251695 | mtdh | + | - | + | - | + |
| XB-GENE-6251778 | pin4 | + | + | - | + | - |
| XB-GENE-6251827 | rpl5 | + | + | + | + | - |
| XB-GENE-6251888 | rps3 | + | - | + | + | - |
| XB-GENE-6252077 | arl2 | + | - | + | - | - |
| XB-GENE-6252283 | mettl1 | + | - | + | - | - |
| XB-GENE-6252609 | ybx2 | + | - | - | + | - |
| XB-GENE-6252872 | ccnd2 | + | - | + | - | + |
| XB-GENE-6253275 | tcof1 | + | - | + | - | + |
| XB-GENE-6253626 | snu13 | + | + | + | + | - |
| XB-GENE-6253776 | rps27 | - | + | + | + | - |
| XB-GENE-6253891 | grb2 | + | - | + | - | - |
| XB-GENE-6253983 | cdkn2aipnl | + | - | - | + | + |
| XB-GENE-6254047 | en2 | + | - | + | - | - |
| XB-GENE-6254049 | ubtf | + | - | + | - | + |
| XB-GENE-6254080 | mphosph6 | + | + | - | + | - |
| XB-GENE-6254218 | fgfr1 | + | - | + | - | - |
| XB-GENE-6254357 | pinx1 | + | - | + | - | - |
| XB-GENE-6254410 | etv4 | + | - | + | - | - |
| XB-GENE-6254472 | h1fx | + | - | + | - | - |
| XB-GENE-6254475 | klf6 | + | - | + | - | - |
| XB-GENE-6254478 | rps14 | + | + | + | + | - |
| XB-GENE-6254553 | npm3 | + | + | + | + | - |
| XB-GENE-6254630 | sumo1 | + | - | + | - | - |
| XB-GENE-6254911 | polr2e | + | - | - | + | - |
| XB-GENE-6254950 | txn2 | + | - | - | + | + |
| XB-GENE-6255612 | wdr82 | + | - | + | - | - |
| XB-GENE-6255773 | rpl18 | + | - | + | + | + |
| XB-GENE-6255941 | mrpl1 | - | + | + | - | - |
| XB-GENE-6256032 | ilf3 | + | - | - | + | - |
| XB-GENE-6256184 | plscr1 | + | - | - | + | - |
| XB-GENE-6256274 | homez | + | - | + | - | - |
| XB-GENE-6256327 | fbl | + | + | - | + | - |
| XB-GENE-6256351 | ctsb | + | - | + | + | + |
| XB-GENE-6256575 | myc | + | - | + | - | - |
| XB-GENE-6256630 | rps6 | + | + | + | + | - |
| XB-GENE-6256647 | cirbp | + | - | + | + | - |
| XB-GENE-6485730 | chd2 | + | - | + | - | - |
| XB-GENE-6488161 | tsc1 | - | + | + | - | - |
| XB-GENE-865142 | nsa2 | + | + | + | - | - |
| XB-GENE-865341 | eef1a1 | + | - | + | + | + |

| | | | | | | |
|---|---|---|---|---|---|---|
| XB-GENE-865449 | prdx1 | + | - | - | + | - |
| XB-GENE-865923 | habp4 | + | - | - | + | - |
| XB-GENE-865972 | tbca | + | - | - | + | - |
| XB-GENE-866018 | nucks1 | + | - | + | - | - |
| XB-GENE-866053 | srsf9 | + | - | + | + | - |
| XB-GENE-866094 | polr2h | + | - | - | + | + |
| XB-GENE-866129 | bccip | - | + | - | + | - |
| XB-GENE-866205 | idh3g | + | - | - | + | + |
| XB-GENE-866214 | sf3b4 | + | - | - | + | - |
| XB-GENE-866350 | rbm4b | + | - | + | + | - |
| XB-GENE-866600 | ppid | + | - | - | + | + |
| XB-GENE-868348 | eif6 | + | + | - | + | + |
| XB-GENE-876487 | ilf2 | + | - | + | + | + |
| XB-GENE-877259 | nkrf | + | - | + | - | - |
| XB-GENE-921816 | rps25 | + | + | + | + | - |
| XB-GENE-922237 | ddx6 | + | - | + | + | + |
| XB-GENE-940111 | rps13 | + | - | + | + | - |
| XB-GENE-941790 | gemin2 | + | - | + | - | - |
| XB-GENE-941933 | zpr1 | + | - | - | + | + |
| XB-GENE-942100 | gnai1 | + | - | - | + | - |
| XB-GENE-943140 | pa2g4 | + | + | + | + | - |
| XB-GENE-943755 | ube2n | + | - | + | + | - |
| XB-GENE-950166 | mob1a | + | - | + | - | - |
| XB-GENE-952676 | pdha1 | + | - | - | + | - |
| XB-GENE-952935 | fbxo11 | + | - | + | - | - |
| XB-GENE-952982 | lipa | + | - | - | + | - |
| XB-GENE-954740 | rpl7 | - | + | + | + | + |
| XB-GENE-955429 | hspa8 | + | - | + | + | - |
| XB-GENE-958073 | jpt1 | + | - | + | - | - |
| XB-GENE-958287 | rpl6 | + | + | + | + | - |
| XB-GENE-958312 | rpl23 | + | - | + | + | - |
| XB-GENE-959327 | rpl7a | - | + | + | + | - |
| XB-GENE-960234 | hspa9 | + | - | - | + | + |
| XB-GENE-961676 | ppp1cb | + | - | + | + | - |
| XB-GENE-962272 | cdk8 | + | - | + | - | - |
| XB-GENE-963406 | cpt2 | + | - | + | + | - |
| XB-GENE-963927 | rpl24 | - | + | + | + | - |
| XB-GENE-964826 | rpl13 | + | - | + | - | - |
| XB-GENE-966363 | naa10 | + | - | - | + | + |
| XB-GENE-967651 | rps24 | - | + | + | + | - |
| XB-GENE-967901 | rpl38 | - | + | + | + | - |
| XB-GENE-967940 | ppp1cc | + | - | + | - | - |
| XB-GENE-968803 | rps19 | + | + | + | + | + |
| XB-GENE-969783 | rps10 | + | + | + | - | - |
| XB-GENE-969834 | rpl35 | - | + | + | + | + |
| XB-GENE-970184 | top2a | + | - | - | + | - |

| | | | | | | |
|---|---|---|---|---|---|---|
| XB-GENE-970516 | rpl10 | - | + | + | + | - |
| XB-GENE-971707 | rpl3 | + | + | + | + | + |
| XB-GENE-972241 | rpsa | - | + | + | + | + |
| XB-GENE-974023 | ube2i | + | - | + | - | + |
| XB-GENE-976216 | rplp0 | - | + | + | + | - |
| XB-GENE-977079 | apex1 | + | - | - | + | - |
| XB-GENE-977331 | rps7 | + | + | + | + | + |
| XB-GENE-977778 | exosc2 | + | + | + | - | - |
| XB-GENE-978304 | rps3 | + | - | + | + | - |
| XB-GENE-978496 | sumo1 | + | - | + | - | - |
| XB-GENE-978890 | pwp1 | + | + | + | + | - |
| XB-GENE-979830 | rsl24d1 | + | + | + | - | - |
| XB-GENE-980044 | cmpk1 | + | - | + | + | - |
| XB-GENE-980281 | cct5 | + | - | + | + | + |
| XB-GENE-981648 | cox7a2l | + | - | + | - | + |
| XB-GENE-982565 | blm | + | - | - | + | - |
| XB-GENE-983608 | pcdh1 | + | - | - | + | - |
| XB-GENE-983917 | rpl5 | + | + | + | + | - |
| XB-GENE-986099 | grwd1 | + | - | - | + | + |
| XB-GENE-987148 | stk24 | + | - | + | - | - |
| XB-GENE-987378 | sirt2 | + | - | - | + | - |
| XB-GENE-987869 | gpatch2 | + | - | + | - | - |
| XB-GENE-988734 | anp32b | + | - | + | + | - |
| XB-GENE-989990 | ncl | + | + | + | + | - |
| XB-GENE-990494 | rrn3 | + | + | - | + | - |
| XB-GENE-991563 | wdr43 | + | + | + | - | - |
| XB-GENE-992535 | lyar | + | + | - | + | - |
| XB-GENE-993334 | rtf1 | + | - | + | - | - |
| XB-GENE-993907 | srp68 | + | - | + | - | + |
| XB-GENE-994299 | ssrp1 | + | - | + | - | - |
| XB-GENE-994401 | eif3a | + | - | + | + | - |
| XB-GENE-995327 | mrps9 | + | + | + | - | - |
| XB-GENE-996300 | rcc2 | + | - | + | - | - |
| XB-GENE-997476 | utp6 | + | + | + | - | - |
| XB-GENE-997561 | senp3 | + | - | + | - | - |
| XB-GENE-998249 | ciapin1 | + | - | - | + | - |
| XB-GENE-999065 | ell3 | + | - | + | - | - |
| XB-GENE-999176 | ppm1b | + | - | + | + | - |
| XB-GENE-999337 | rpl27 | - | + | + | + | - |
| XB-GENE-999551 | npm2 | + | - | - | + | - |
| XB-GENE-999738 | ddx3x | - | + | + | + | - |

Axonal_RibosomeAssemblyFactors: All genes annotated with GO termes "nucleolus" or "ribosome biogenesis" that are detected in the axonal transcritome

| Xenbase_ID | mouse gene name | GO term "nucleolus" | GO term "ribosome biogenesis" | Xlaevis axonal transcriptome | Xlaevis axonal proteome | mouse axon translatome |
|---|---|---|---|---|---|---|
| XB-GENE-1000251 | hmgb2 | + | - | + | + | - |
| XB-GENE-1000491 | fbxw7 | + | - | + | - | + |
| XB-GENE-1000765 | ddx17 | + | + | + | - | - |
| XB-GENE-1000959 | vmp1 | + | - | + | - | - |
| XB-GENE-1000983 | rpl11 | + | + | + | + | + |
| XB-GENE-1001390 | tyms | + | - | - | + | - |
| XB-GENE-1001971 | rpl31 | + | - | + | + | - |
| XB-GENE-1002501 | nono | + | - | + | - | - |
| XB-GENE-1003294 | rps6 | + | + | + | + | - |
| XB-GENE-1003623 | rps2 | - | + | + | + | - |
| XB-GENE-1004641 | nup88 | - | + | + | - | - |
| XB-GENE-1005073 | ltv1 | - | + | + | - | - |
| XB-GENE-1006309 | rpl18 | + | - | + | + | + |
| XB-GENE-1007936 | rps21 | - | + | - | + | + |
| XB-GENE-1008656 | frg1 | + | + | + | - | - |
| XB-GENE-1008861 | rpl12 | + | + | + | + | - |
| XB-GENE-1008948 | dhx15 | + | - | - | + | - |
| XB-GENE-1010464 | snrpb2 | + | - | + | - | - |
| XB-GENE-1013231 | asna1 | + | - | - | + | - |
| XB-GENE-1013309 | uso1 | + | - | - | + | + |
| XB-GENE-1014730 | cask | + | - | - | + | - |
| XB-GENE-1015264 | ola1 | + | - | + | - | - |
| XB-GENE-1016064 | rpl35a | - | + | + | + | - |
| XB-GENE-1016064 | rpl35a | - | + | + | + | - |
| XB-GENE-1016264 | exosc7 | + | + | - | + | - |
| XB-GENE-1016377 | rps17 | - | + | + | - | + |
| XB-GENE-1016477 | mrps15 | + | - | + | - | - |
| XB-GENE-1188424 | sub1 | + | - | + | + | - |
| XB-GENE-17330372 | xrn2 | + | - | - | + | - |
| XB-GENE-17330417 | ube2n | + | - | + | + | - |
| XB-GENE-17330554 | ddx3x | - | + | + | + | - |
| XB-GENE-17331043 | kif2a | + | - | + | - | - |
| XB-GENE-17331209 | rpl7a | - | + | + | + | - |
| XB-GENE-17331340 | nuak1 | + | - | + | - | - |
| XB-GENE-17331553 | wbp11 | - | + | + | - | - |
| XB-GENE-17332277 | dctn3 | + | - | - | + | + |
| XB-GENE-17332836 | rpap2 | + | - | + | - | - |
| XB-GENE-17334360 | fcf1 | + | + | + | - | + |
| XB-GENE-17334580 | tsc1 | - | + | + | - | - |

| | | | | | | |
|---|---|---|---|---|---|---|
| XB-GENE-17334901 | smad7 | + | - | + | - | - |
| XB-GENE-17334929 | rbm4b | + | - | + | + | - |
| XB-GENE-17335149 | xpo1 | + | + | + | + | - |
| XB-GENE-17335914 | nhp2 | + | + | + | - | - |
| XB-GENE-17335931 | fbxo11 | + | - | + | - | - |
| XB-GENE-17337446 | rplp0 | - | + | + | + | - |
| XB-GENE-17337654 | aamp | - | + | - | + | + |
| XB-GENE-17338571 | dedd2 | + | - | + | - | - |
| XB-GENE-17339479 | cdc14b | + | - | + | - | - |
| XB-GENE-17339580 | plrg1 | + | - | + | - | - |
| XB-GENE-17339884 | pus1 | + | - | + | - | - |
| XB-GENE-17340555 | ddx5 | + | - | - | + | - |
| XB-GENE-17340776 | rps13 | + | - | + | + | - |
| XB-GENE-17340849 | srsf5 | + | - | + | + | - |
| XB-GENE-17341054 | parn | + | - | + | - | - |
| XB-GENE-17342307 | nucks1 | + | - | + | - | - |
| XB-GENE-17342370 | rai14 | + | - | + | - | - |
| XB-GENE-17343779 | jpt1 | + | - | + | - | - |
| XB-GENE-17343925 | edf1 | + | - | - | + | - |
| XB-GENE-17344173 | sub1 | + | - | + | + | - |
| XB-GENE-17344257 | ddx6 | + | - | + | + | + |
| XB-GENE-17344335 | rbm14 | + | - | + | + | - |
| XB-GENE-17344569 | sdha | + | - | - | + | + |
| XB-GENE-17346030 | abce1 | - | + | + | + | - |
| XB-GENE-17346538 | ddx23 | + | - | + | - | - |
| XB-GENE-492329 | tra2a | + | - | - | + | - |
| XB-GENE-492785 | cirbp | + | - | + | + | - |
| XB-GENE-493737 | eif3l | + | - | - | + | + |
| XB-GENE-5738730 | gnai3 | + | - | - | + | - |
| XB-GENE-5786746 | rps16 | - | + | + | - | - |
| XB-GENE-5800445 | zc3h15 | + | - | + | - | + |
| XB-GENE-5819322 | rpl14 | - | + | + | + | + |
| XB-GENE-5828312 | leo1 | + | - | + | - | + |
| XB-GENE-5833061 | rps15 | - | + | + | + | + |
| XB-GENE-5865181 | macrod2 | + | - | - | + | - |
| XB-GENE-5871876 | ckap5 | + | - | - | + | - |
| XB-GENE-5891407 | rpl23a | - | + | + | + | - |
| XB-GENE-5900967 | kdm7a | + | - | + | - | - |
| XB-GENE-5919816 | srp19 | + | - | - | + | - |
| XB-GENE-5929894 | acat2 | + | - | - | + | + |
| XB-GENE-5930659 | rps9 | + | - | + | + | + |
| XB-GENE-5935325 | rps8 | - | + | + | + | + |
| XB-GENE-5945253 | map2 | + | - | - | + | - |
| XB-GENE-5948460 | ppp1ca | + | - | + | - | - |
| XB-GENE-5952259 | rabgef1 | + | - | + | - | + |
| XB-GENE-6042311 | pim1 | + | - | + | - | - |

| | | | | | | |
|---|---|---|---|---|---|---|
| XB-GENE-6053022 | myc | + | - | + | - | - |
| XB-GENE-6068534 | gtpbp4 | + | + | + | - | - |
| XB-GENE-6077488 | slbp | + | - | + | - | - |
| XB-GENE-6078160 | bcas2 | + | - | - | + | + |
| XB-GENE-6251644 | timm13 | + | - | - | + | + |
| XB-GENE-6251695 | mtdh | + | - | + | - | + |
| XB-GENE-6251778 | pin4 | + | + | - | + | - |
| XB-GENE-6251827 | rpl5 | + | + | + | + | - |
| XB-GENE-6251888 | rps3 | + | - | + | + | - |
| XB-GENE-6252077 | arl2 | + | - | + | - | - |
| XB-GENE-6252283 | mettl1 | + | - | + | - | - |
| XB-GENE-6252609 | ybx2 | + | - | - | + | - |
| XB-GENE-6252872 | ccnd2 | + | - | + | - | + |
| XB-GENE-6253275 | tcof1 | + | - | + | - | + |
| XB-GENE-6253626 | snu13 | + | + | + | + | - |
| XB-GENE-6253776 | rps27 | - | + | + | + | - |
| XB-GENE-6253891 | grb2 | + | - | + | - | - |
| XB-GENE-6253983 | cdkn2aipnl | + | - | - | + | + |
| XB-GENE-6254047 | en2 | + | - | + | - | - |
| XB-GENE-6254049 | ubtf | + | - | + | - | + |
| XB-GENE-6254080 | mphosph6 | + | + | - | + | - |
| XB-GENE-6254218 | fgfr1 | + | - | + | - | - |
| XB-GENE-6254357 | pinx1 | + | - | + | - | - |
| XB-GENE-6254410 | etv4 | + | - | + | - | - |
| XB-GENE-6254472 | h1fx | + | - | + | - | - |
| XB-GENE-6254475 | klf6 | + | - | + | - | - |
| XB-GENE-6254478 | rps14 | + | + | + | + | - |
| XB-GENE-6254553 | npm3 | + | + | + | + | - |
| XB-GENE-6254630 | sumo1 | + | - | + | - | - |
| XB-GENE-6254911 | polr2e | + | - | - | + | - |
| XB-GENE-6254950 | txn2 | + | - | - | + | + |
| XB-GENE-6255612 | wdr82 | + | - | + | - | - |
| XB-GENE-6255773 | rpl18 | + | - | + | + | + |
| XB-GENE-6255941 | mrpl1 | - | + | + | - | - |
| XB-GENE-6256032 | ilf3 | + | - | - | + | - |
| XB-GENE-6256184 | plscr1 | + | - | - | + | - |
| XB-GENE-6256274 | homez | + | - | + | - | - |
| XB-GENE-6256327 | fbl | + | + | - | + | - |
| XB-GENE-6256351 | ctsb | + | - | + | + | + |
| XB-GENE-6256575 | myc | + | - | + | - | - |
| XB-GENE-6256630 | rps6 | + | + | + | + | - |
| XB-GENE-6256647 | cirbp | + | - | + | + | - |
| XB-GENE-6485730 | chd2 | + | - | + | - | - |
| XB-GENE-6488161 | tsc1 | - | + | + | - | - |
| XB-GENE-865142 | nsa2 | + | + | + | - | - |
| XB-GENE-865341 | eef1a1 | + | - | + | + | + |

| | | | | | | |
|---|---|---|---|---|---|---|
| XB-GENE-865449 | prdx1 | + | - | - | + | - |
| XB-GENE-865923 | habp4 | + | - | - | + | - |
| XB-GENE-865972 | tbca | + | - | - | + | - |
| XB-GENE-866018 | nucks1 | + | - | + | - | - |
| XB-GENE-866053 | srsf9 | + | - | + | + | - |
| XB-GENE-866094 | polr2h | + | - | - | + | + |
| XB-GENE-866129 | bccip | - | + | - | + | - |
| XB-GENE-866205 | idh3g | + | - | - | + | + |
| XB-GENE-866214 | sf3b4 | + | - | - | + | - |
| XB-GENE-866350 | rbm4b | + | - | + | + | - |
| XB-GENE-866600 | ppid | + | - | - | + | + |
| XB-GENE-868348 | eif6 | + | + | - | + | + |
| XB-GENE-876487 | ilf2 | + | - | + | + | + |
| XB-GENE-877259 | nkrf | + | - | + | - | - |
| XB-GENE-921816 | rps25 | + | + | + | + | - |
| XB-GENE-922237 | ddx6 | + | - | + | + | + |
| XB-GENE-940111 | rps13 | + | - | + | + | - |
| XB-GENE-941790 | gemin2 | + | - | + | - | - |
| XB-GENE-941933 | zpr1 | + | - | - | + | + |
| XB-GENE-942100 | gnai1 | + | - | - | + | - |
| XB-GENE-943140 | pa2g4 | + | + | + | + | - |
| XB-GENE-943755 | ube2n | + | - | + | + | - |
| XB-GENE-950166 | mob1a | + | - | + | - | - |
| XB-GENE-952676 | pdha1 | + | - | - | + | - |
| XB-GENE-952935 | fbxo11 | + | - | + | - | - |
| XB-GENE-952982 | lipa | + | - | - | + | - |
| XB-GENE-954740 | rpl7 | - | + | + | + | + |
| XB-GENE-955429 | hspa8 | + | - | + | + | - |
| XB-GENE-958073 | jpt1 | + | - | + | - | - |
| XB-GENE-958287 | rpl6 | + | + | + | + | - |
| XB-GENE-958312 | rpl23 | + | - | + | + | - |
| XB-GENE-959327 | rpl7a | - | + | + | + | - |
| XB-GENE-960234 | hspa9 | + | - | - | + | + |
| XB-GENE-961676 | ppp1cb | + | - | + | + | - |
| XB-GENE-962272 | cdk8 | + | - | + | - | - |
| XB-GENE-963406 | cpt2 | + | - | + | + | - |
| XB-GENE-963927 | rpl24 | - | + | + | + | - |
| XB-GENE-964826 | rpl13 | + | - | + | - | - |
| XB-GENE-966363 | naa10 | + | - | - | + | + |
| XB-GENE-967651 | rps24 | - | + | + | + | - |
| XB-GENE-967901 | rpl38 | - | + | + | + | - |
| XB-GENE-967940 | ppp1cc | + | - | + | - | - |
| XB-GENE-968803 | rps19 | + | + | + | + | + |
| XB-GENE-969783 | rps10 | + | + | + | - | - |
| XB-GENE-969834 | rpl35 | - | + | + | + | + |
| XB-GENE-970184 | top2a | + | - | - | + | - |

| | | | | | | |
|---|---|---|---|---|---|---|
| XB-GENE-970516 | rpl10 | - | + | + | + | - |
| XB-GENE-971707 | rpl3 | + | + | + | + | + |
| XB-GENE-972241 | rpsa | - | + | + | + | + |
| XB-GENE-974023 | ube2i | + | - | + | - | + |
| XB-GENE-976216 | rplp0 | - | + | + | + | - |
| XB-GENE-977079 | apex1 | + | - | - | + | - |
| XB-GENE-977331 | rps7 | + | + | + | + | + |
| XB-GENE-977778 | exosc2 | + | + | + | - | - |
| XB-GENE-978304 | rps3 | + | - | + | + | - |
| XB-GENE-978496 | sumo1 | + | - | + | - | - |
| XB-GENE-978890 | pwp1 | + | + | + | + | - |
| XB-GENE-979830 | rsl24d1 | + | + | + | - | - |
| XB-GENE-980044 | cmpk1 | + | - | + | + | - |
| XB-GENE-980281 | cct5 | + | - | + | + | + |
| XB-GENE-981648 | cox7a2l | + | - | + | - | + |
| XB-GENE-982565 | blm | + | - | - | + | - |
| XB-GENE-983608 | pcdh1 | + | - | - | + | - |
| XB-GENE-983917 | rpl5 | + | + | + | + | - |
| XB-GENE-986099 | grwd1 | + | - | - | + | + |
| XB-GENE-987148 | stk24 | + | - | + | - | - |
| XB-GENE-987378 | sirt2 | + | - | - | + | - |
| XB-GENE-987869 | gpatch2 | + | - | + | - | - |
| XB-GENE-988734 | anp32b | + | - | + | + | - |
| XB-GENE-989990 | ncl | + | + | + | + | - |
| XB-GENE-990494 | rrn3 | + | + | - | + | - |
| XB-GENE-991563 | wdr43 | + | + | + | - | - |
| XB-GENE-992535 | lyar | + | + | - | + | - |
| XB-GENE-993334 | rtf1 | + | - | + | - | - |
| XB-GENE-993907 | srp68 | + | - | + | - | + |
| XB-GENE-994299 | ssrp1 | + | - | + | - | - |
| XB-GENE-994401 | eif3a | + | - | + | + | - |
| XB-GENE-995327 | mrps9 | + | + | + | - | - |
| XB-GENE-996300 | rcc2 | + | - | + | - | - |
| XB-GENE-997476 | utp6 | + | + | + | - | - |
| XB-GENE-997561 | senp3 | + | - | + | - | - |
| XB-GENE-998249 | ciapin1 | + | - | - | + | - |
| XB-GENE-999065 | ell3 | + | - | + | - | - |
| XB-GENE-999176 | ppm1b | + | - | + | + | - |
| XB-GENE-999337 | rpl27 | - | + | + | + | - |
| XB-GENE-999551 | npm2 | + | - | - | + | - |
| XB-GENE-999738 | ddx3x | - | + | + | + | - |
